# Measuring carbohydrate recognition profile of lectins on live cells using liquid glycan array (LiGA)

**DOI:** 10.1101/2023.10.04.559794

**Authors:** Mirat Sojitra, Edward N. Schmidt, Guilherme M. Lima, Eric J. Carpenter, Kelli A. McCord, Alexey Atrazhev, Matthew S. Macauley, Ratmir Derda

**Affiliations:** Department of Chemistry, University of Alberta, Edmonton, AB T6G 2G2, Canada; Department of Medical Microbiology and Immunology, University of Alberta, Edmonton, AB T6G 2G2, Canada

## Abstract

Glycans constitute a significant fraction of biomolecular diversity on the surface of cells across all the species in all kingdoms of life. As the structure of glycans is not encoded by the DNA of the host organisms, it is impossible to use cutting-edge DNA technology to study the role of cellular glycosylation or to understand how cell-surface glycome is recognized by glycan-binding proteins (GBPs). To address this gap, we recently described a genetically-encoded liquid glycan array (LiGA) platform that allows profiling of glycan:GBP interactions on the surface of live cells *in vitro* and *in vivo* using next-generation sequencing (NGS). LiGA is a library of DNA-barcoded bacteriophages coated with 5-1500 copies of a glycan; the DNA barcode inside each bacteriophage encodes the structure and density of the displayed glycans. Deep sequencing of the glycophages associated with live cells yields a glycan-binding profile of GBPs displayed on the surface of such cells. This protocol provides detailed instructions of using LiGA to probe cell surface receptors and includes information on the preparation of glycophages, analysis by MALDI-TOF MS, the assembly of a LiGA library, and its deep-sequencing. Using the protocol detailed in this report, we measure a glycan-binding profile of the immunomodulatory SiglecLJ1, -2, -6, -7, and -9 expressed on the surface of different cell types and uncover previously unknown environment-dependent recognition of glycans by Siglec-receptors on the surface of live cells. Protocols similar to the one described in this report will make it possible to measure the precise glycan-binding profile of any GPBs displayed on the surface of any cell types.

## Introduction

Across the entire phylogenetic tree, glycosylation (carbohydrates) is found among all the species. Glycan–GBP (glycan binding protein) interactions are ubiquitous on the surface of cells and viruses, and mediate many important physiological processes, such as cellular communication, host-pathogen recognition, and immune responses.^1,2^ In a cell–cell interactions, for example, the glycocalyx of one cell is recognized by GBPs extended from the opposing cell surface resulting in an exchange of informational cues. Many of the modern diagnostics and therapeutic applications of glycans emerged from an understanding of how GBPs on a cell surface interact with glycans on the surface of cells or invading pathogens.^3^ Thus, the development of a tool that rapidly profiles the glycan binding specificity of a lectin expressed on the surface of live cells provides a means to better understand the role of saccharides in biological systems.

High throughput glycan binding assays must recapitulate the multivalent presentation of saccharides for effective mimicking of the natural display of glycans found on the surface of cells. For example, the widely used, glycan arrays mimic the multivalent display of glycans by printing synthetic glycans on the surface of the glass.^4^ Despite their utility in identifying ligand specificity of a lectin, such ‘glass’ glycan arrays cannot effectively assay GBPs in their natural environment: on the surface of live cells. Isolating GBPs from their natural environment (e.g., cell surface) can lead to misleading results due to confounding factors. For example, the sialic acid binding lectin hCD22 expressed on the B-cell surface is often bound to sialic acid present on the same cell surface and as a result, these *cis*-interactions attenuate the interaction of hCD22 with exogenous glycans (*trans*-binding).^5,6^ Due to the ubiquitous presence of diverse glycans on the surface of cells, a conceptually similar *cis*-inhibition may modulate glycan-binding preferences of many other classes of cell-surface GBPs. Glycan probes such as glyco-polymers,^7^ glyco-liposomes,^8^ and glycosylated viral capsids^9^ have been used extensively to study the multivalent specificity of GBPs on live cells and cellular response, however, unlike the canonical glycan arrays, these synthetic glycan nano-scale probes do not encode the identity of glycans and as a result, the utility of such probes is limited to the lower throughput “one-experiment-one-glycan” measurements. There is an unmet need for technology that can test the biological role of multivalent glycan display in live cells *in vitro* and *in vivo* in a multiplexed fashion: multiple glycans and their presentations in a single experiment.

To address this need, we developed the Liquid Glycan Array (LiGA)^10^ which is a collection of silently DNA barcoded^11^ M13 phage particles that are chemically modified by glycans to yield a multivalent display of glycans on a monodisperse 700 nm x 5 nm M13 virion. We demonstrated that LiGA can be constructed using pre-synthesized glycans^10^ and customized using enzymatic modifications directly on phage.^12^ Inspired by DNA-encoded libraries (DEL), DNA-displayed glycan libraries have been developed;^13–15^ however, LiGA offers several advantages over such display of glycans on ‘naked’ DNA (i) The internal location of the DNA barcode inside M13 virion in LiGA protects this DNA in demanding biological environment and thus make it possible to investigate the association of LiGA with GBPs on live cells both *in vitro* and *in vivo*;^10,12^ (ii) Examples of display of glycans on DNA are either mandatory monovalent display on 5’ end of DNA strand,^13,14^ or fixed trivalent display on DNA hairpin^15^ whereas number of glycans on phage can be controlled from 20 to 1500 copies per 700 nm-long virion. This density can also be encoded in DNA,^10^ and it has a significant effect on interactions of glycans with target GBPs.^10,12,16^ In three previous reports, we employed LiGA to profile the binding preferences of three receptors: DC-SIGN^10,12^, hCD22,^10,12^ and Siglec-1^16^ on the surface of live cells. In this report, we describe the systematic application of a LiGA library of ∼100 glycophage conjugates (LiGA-100) and a focused ganglioside LiGA (LiGA-g) to profile a subset of human Siglec family of GBPs expressed on the surface of cells. We demonstrated the applicability of LiGA technology by utilizing distinct lineages of cells with various levels of intrinsic glycosylation on the surface to profile the specificity of Siglec-1, -2, -6, -7, and -9 and inactive arginine mutants of most of these receptors.

Among the 15 human Siglec cell surface receptors, the majority of the Siglecs function as an immune receptor. Siglecs exhibit a relatively weak affinity for sialo-glycans, with *K*_d_ ranging from 0.05 to 5 mM.^17^ The major determinants of their ligand specificity are combination of several factors: (i) the glycosidic linkage between the sialic acid and the galactose/GalNAc (ex. α2-6 or α2-3), (ii) presence of additional subunits of sialic acid (ex. α2-8), and (iii) composition of on the glycan core preceding the sialic acid (e.g., monosaccharide composition,^18^ presence of sulfation,^19^ etc.). Despite the weak affinity, interactions of Siglecs on immune cells and sialic acids in the glycocalyx of other cells control numerous immunological and oncological processes. For example, hyper-sialylated cancer cell surfaces engage the Siglec-7 and -9 on natural killer (NK) cells to evade the immune response by recruiting the Siglecs receptors to the immune synapses.^20,21,22^ Blocking these Siglec-sialo glycan interactions is being actively pursued as a strategy to increase the ability of the immune system or chemotherapy to eradicate tumor cells.^3,23,24^ Understanding glycan binding specificities of Siglec immunomodulators is critical in the discovery of lead compounds for the next generation of carbohydrate-based therapeutics. This study showcases the ability of LiGA technology to measure the glycan-binding profile of Siglec receptors displayed on the surface of distinct mammalian cell types. Following the protocols described in this report, any GBP expressed on the cell surface of cells can be profiled using the same strategy.

### Design of DNA barcoded M13 phages

The DNA-encoded M13 phages are scalable and stable nano-carriers which are compatible with a wide range of cloning methods as well as chemical and enzymatic conjugation strategies.^25^ Glycans are chemically conjugated on the surface of M13 phages and the internal location of the DNA barcode not only protects the integrity of DNA but also avoids non-specific interactions between charged DNA strands and glycans, glycan-binding proteins (GBPs) or cells displaying such GBPs. For the DNA barcode, we used a SDB (silent double barcode) created by cloning SB1 (silent barcode 1)^11^ and SB2 (silent barcode 2) into the pIII-leader region of M13KE vector (**Extended Data Figure 1a**). Building on previously developed cloning vectors,^11^ the SB2 region was cloned as a sequence of degenerate DNA codons all encoding SVEKNDQKTYHAGGG (SVEK) peptide displayed on pIII protein. Although the SVEK peptide has no intrinsic function, it contains an N-terminal serine residue that could be used for site-selective modification, such as biotinylation.^25^ SVEK on pIII peptide can be also replaced by any desired functional peptide like FLAG tag or Spy-Tag peptide.^25–27^ Together, the cloning of SB1 and SB2 degenerate nucleotide sequences offered 10^10^ theoretically possible combinations and yielded a library of ‘silently’ double-barcoded (SDB) vectors that encode chemically identical phage particles (**Extended Data Figure 1a**). To isolate clonal phages, we used classical ‘plaque picking’ from agar overlay plated with a low dilution of the SDB-SVEK-M13 phage library (**Extended Data Figure 1b**). We retained phage clones that contained a unique DNA barcode sequence separated from one another by at least three nucleotide substitutions (Hamming distance, H > 3). The H > 3 design allowed for the correction of point mutations, single nucleotide insertions or deletions, and other mistakes that often occur in next-generation DNA sequencing. Every phage was amplified in mid-scale culture (25 mL) and purified by PEG precipitation to yield a homogeneous solution of phage at high concentration (10^13^ pfu/mL). To remove the LPS (lipopolysaccharides) endotoxins present in the amplified clonal phages, the phages were further subject to standard Triton X-100 purification (**Extended Data Figure 1b**);^28,29^ this LPS-removal protocol has been employed by Krag and co-workers to prepare clinical-grade M13 phage particles for injections into human patients.^30^ Post purification, the clonal phages were stored in 1:1 PBS–glycerol at -20 °C until conjugation with glycans.

### Modification of prospectively DNA barcoded phages by glycan

Strain-promoted azide–alkyne cycloaddition (SPAAC)^10,12^ ligates readily available saccharides with alkyl-azido linkers to dibenzocyclooctyne (DBCO)-modified phages. We employ the same strategy to assemble glycan-coated phages used in this report. For synthetic glycans, approximately 63 azido-glycans were obtained as part of the public Consortium for Functional Glycomics (CFG) collection,^31^ and an additional 12 azido-glycans were synthesized as described in a previous report.^10^ Ligation of glycans to purified phage clones begins with acylation of a subset of the 2700 major coat protein pVIII protein of the M13 phage by DBCO-NHS ester (**Extended Data Figure 1c-d**). Varying the concentration of DBCO–NHS ester from 0.5 to 2 mM enabled installing the desired number of conjugation sites (DBCO) on a single phage virion and offers precise control over the density of displayed glycan. Measurements of peak intensities in MALDI-TOF of unmodified pVIII and modified, pVIII–DBCO peak yielded an estimate of the fraction of pVIII proteins modified by DBCO (**Extended Data Figure 1e**). Subsequent addition of 1 mM azido-glycan quantitatively consumed the strained alkynes and ligated the glycan to pVIII, as evidenced by the appearance of pVIII–DBCO–glycan peak and the concomitant disappearance of the pVIII–DBCO peak (**Extended Data Figure 1e**). Mixing these differentially barcoded glycophages produced a LiGA library that contains any desired combinations of glycans. Each LiGA mixture is accompanied by a data table (LiGA dictionary) listing the SDB barcode and the corresponding glycan.

### Development of the protocol

A typical binding assay between cells and LiGA library consists of four critical steps: i) *incubation* of LiGA with cells, ii) *washing* away non-binding phages, iii) *elution* of the bound phage, and iv) *PCR* amplification of DNA of the eluted phage and next-generation sequencing by Illumina Sequencer (**Figure 1a-d**). First, the confluent GBP^+^ and GBP^−^ mammalian cells are resuspended in the incubation buffer (HEPES buffer supplemented with 1% BSA) at ∼4 × 10^6^ cells/mL to maintain conditions compatible with live cells (**Figure 1b**). The cells are aliquoted (250 μL each) into FACS tubes (Corning, #352054), which afforded ∼1 million cells per FACS tube for replicate samples. To recover enough phage DNA for a reliable PCR, 100 phages for every 1 cell were mixed and incubated at 37 °C for 1 h (**Figure 1b**). Specifically, 10^6^ cells are mixed with 10^8^ PFU of LiGA, and in such LiGA there are on average 10^6^ copies of each clonal glycophage. In previous reports, incubation of LiGA with cells was performed at 4 °C to primarily focus on binding and to avoid the internalization of the glycophage.^10^ In this report, we used both at 4 °C and 37 °C conditions and observed a significant increase in the number of phages retained by cells at 37 °C. The increased number of retained phage particles, in turn, improved the reliability of downstream PCR procedures (**Extended Data Figure 2a**).

**Figure 1:**
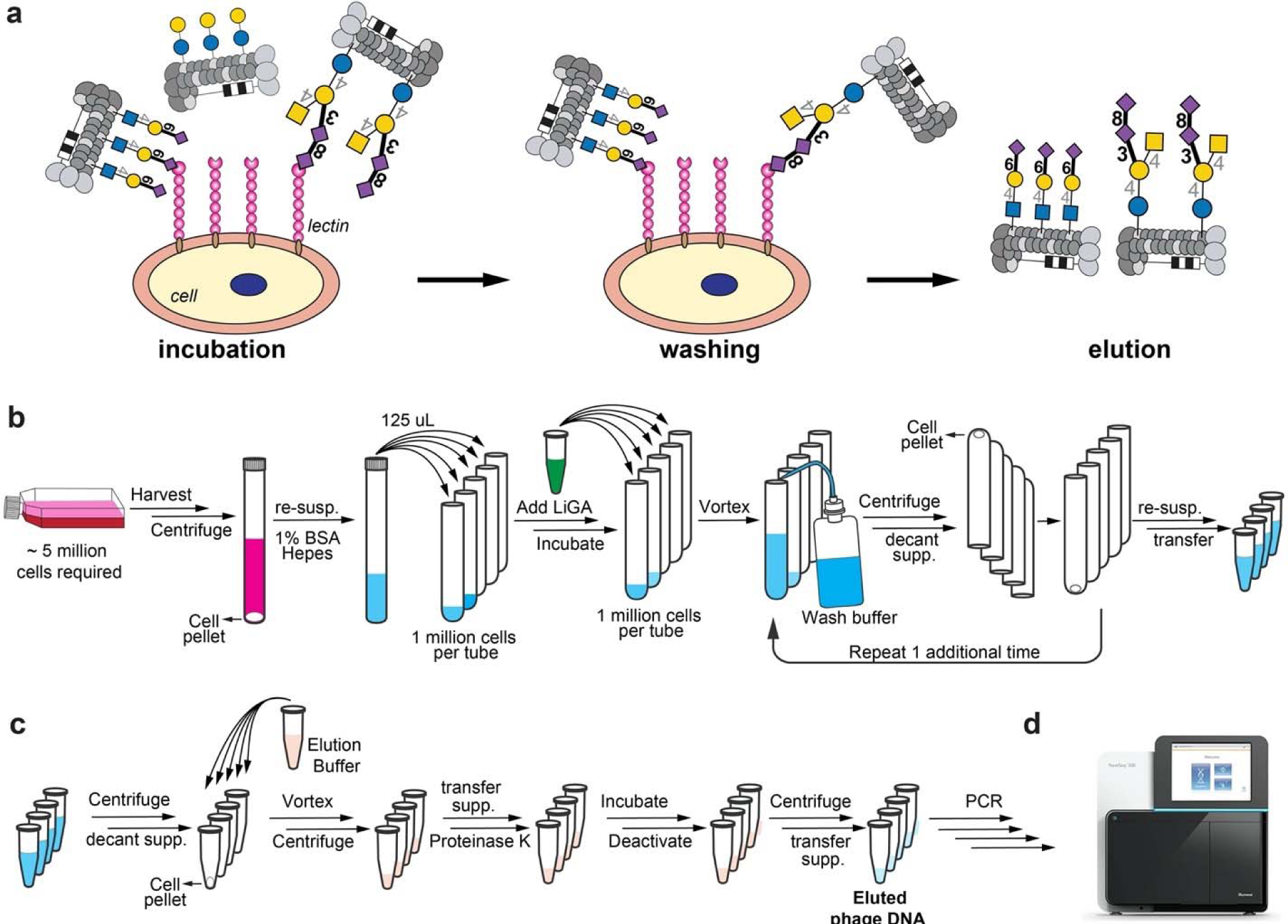
Describes the workflow for cell-surface binding assay using LiGA. **a)** Schematic showing the sequence of LiGA binding assay. **b)** LiGA library is incubated with cells in a HEPES buffered solution supplemented with BSA. Typically, GBP+ and isogenic GBP-cells are processed with LiGA in parallel on the same day with five replicates each. Cells with bound phages are washed three times to remove non-specific binding glycosylated phages present within LiGA. **c)** The bound phages are eluted using a combination of cell lysing and RNase A treatment, followed by Proteinase K to release DNA material from the phage. **d)** The eluted DNA solution is used as a PCR template to prepare for deep sequencing.

Glycophages associated with the cells via weak or non-specific interactions should be washed away by repeated low-speed centrifugation of the cell suspension (**Figure 1b**); for reference, the parameters of such washes (speed, duration, buffer, tubes) are identical to those used for preparation of cell suspension for flow-cytometry experiments. We considered three additional parameters to maximize the specific signal and minimize non-specific binding of the LiGA: (i) washing of the cells must be done in a buffer capable of maintaining the integrity of cells and glycan–GBP interactions (HBS is supplemented with divalent cations and with 0.1% BSA to maintain viscosity and osmolality values compatible with live cells), (ii) centrifugation speed must be low (<300 ×g) to avoid cell compaction and lysis, (iii) number of washes should be balanced to retain specific binding while removing non-specific binding. To monitor the stringency of washes, as a model, we used weak yet specific interactions between αMan and Dendritic Cell-Specific Intercellular adhesion molecule-3-Grabbing Non-integrin (DC-SIGN) with *K*_d_ = 2300 μM^32^. We used a positive control phage clone that transduces mNeonGreen (green plaques) decorated alpha-Mannose and a negative control clone that transduces mCherry reporter (red plaques) and is decorated by lactose (Galβ1-4Glc1β-) and wild type unmodified phage that transduces no reporter (no color). The mannose-(green), lactose-(red) and unmodified-(white) were combined with LiGA built from M13 clones that transduce β-galactosidase (blue plaques). Plating the cell suspension after each wash on agar overlay made it possible to count the plaque-forming units (PFU) for each glycosylated phage and monitor the retention (or recovery, R = PFU^output^LJ÷LJPFU^input^) of each subpopulation of phage by DC-SIGN^+^ cells. We observed higher retention of mannose-(green) than lactose-(red) or unmodified-(white) phage by DC-SIGN^+^ cells over four washes of the same cell pellet (**Extended Data Figure 2b**). The recovery of LiGA-YY (blue) and Manα-(green) stabilized at 0.1–1% at 3–4 washes thus indicating that three washes are sufficient to remove phages associating via non-specific interaction and retain weak, specific interaction. Optimization of wash using colorimetric reporter phage is not mandatory, but it is a convenient feature that allows simultaneous measurement of 4 distinct phage clones in the same population without sequencing. Similar optimization of the wash can be performed by any other technique that can measure the input and output number of phage (e.g., qPCR used in **Extended Data Fig. 2d**).

In first report of LiGA,^10^ we eluted the DNA barcodes from cell-bound phages by resuspending the cell–LiGA pellet in H_2_O and incubating it at 95 °C to release phage particles / phage DNA into the supernatant. We subsequently observed that such heat-based elution also releases a lot of cellular debris that complicated downstream PCR at a low copy number of eluted phage. Specifically, heat elution yielded DNA and downstream PCR product in interactions of LiGA with GBP^+^ cells, which retained 10^3^-10^5^ phage particles; however, the same heat elution frequently failed to produce sufficient quality of DNA from LiGA associated with control, GBP^-^ cells (i.e., cells that retained <<10^3^ phage particles). We reasoned that the chromatin and nucleic acids from the cells interfere with PCR by non-specific primer annealing, especially when the template (phage DNA) is present in low copy number (**Extended Data Figure 2c**). To remove the contaminating cellular material, we utilized a sequential treatment by RNase A and Proteinase K, to specifically isolate the phage DNA from the cells (**Figure 1c**). **Extended Data Figure 2c-d** compares “heat elution” and the “enzymatic elution” of LiGA bound to Siglec-1^+^ Chinese Hamster Ovary (CHO) cells; enzymatic elution yielded less cellular genomic materials on an agarose gel and a higher amount of phage DNA measured using qPCR.

To assess the glycan binding profile, next-generation sequencing (NGS) quantifies the relative abundance of each DNA barcode (i.e., glycophage) associated with the GBP^+^ and GBP^-^ mammalian cells. We sequence SDB barcodes using Illumina sequencing paired end 2x75 bp kit, but we anticipate any NGS platform can be used. In previous reports, we optimized a one-step PCR procedure to convert phage ssDNA to dsDNA product that contains unique multiplexing index barcodes and the Illumina adapters for binding to flow cell and bridge amplification (**Figure 2b and Supplementary Table 1**).^10^ One-step strategy, however, requires primers with large “over-hangs” which produced significant off-target amplification in PCR with low copy number of template (**Extended Data Figure 2e**). To mitigate this problem for low copy number samples, we employed a “two-step” or nested PCR method (**Figure 2a**). The first step uses robust PCR primers with no overhangs and the ample amount of dsDNA product of the first step PCR serves as an abundant template for the second PCR step which appends multiplexing barcodes and the Illumina adapters (**Figure 2a**). The two-step PCR method increased the reliability of PCR from eluted DNA samples containing as few as ∼10^1^ PFU (**Extended Data Figure 2e**).

**Figure 2:**
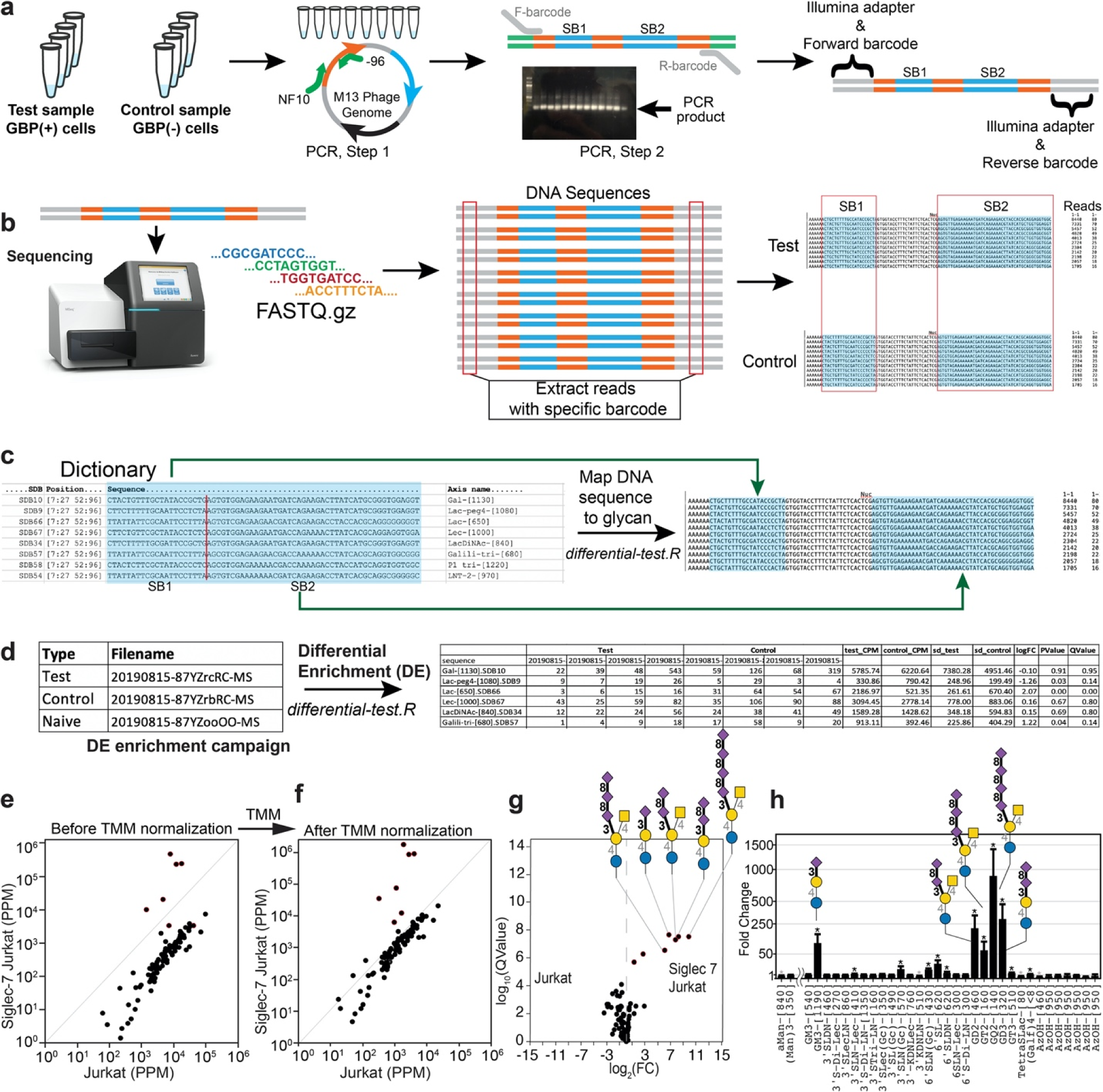
The workflow for PCR, deep-seq, and bioinformatics. **a)** Eluted samples of GBP+ and isogenic GBP-cell line are PCR amplified using primers NF10 and -96. Then, the PCR product is amplified further with primers containing Illumina adapters and multiplexing barcodes. **b)** Illumina sequencing of the PCR product. The content of the FASTQ file is converted to count tables by extracting specific reads based on the multiplexing barcodes. **c)** Glycans are assigned to each DNA read based on the SDB barcode that is stored in a dictionary. **d)** A comparison campaign is generated, and a differential enrichment algorithm is applied to determine the fold change (FC) and q-values. **e)** Scatter plot of normalized DNA reads for each glycan recovered from test and control cell line; normalization was performed simply by division by total read depth. **f)** Scatter plot of TMM normalized DNA reads. Unlike (**e**), the majority of reads are equalized between test and control cells in (**f**). **g)** A Volcano plot visualization of glycans that are enriched significantly in GBP+ but not control cell lines. **h)** Condensed bar chart version of the plot (**g**) in which the FC value is plotted for each glycan and the significance is denoted as * for q-value < 0.05. The results of the entire LiGA-100 binding are presented in **Figure 3e**.

**Figure 3:**
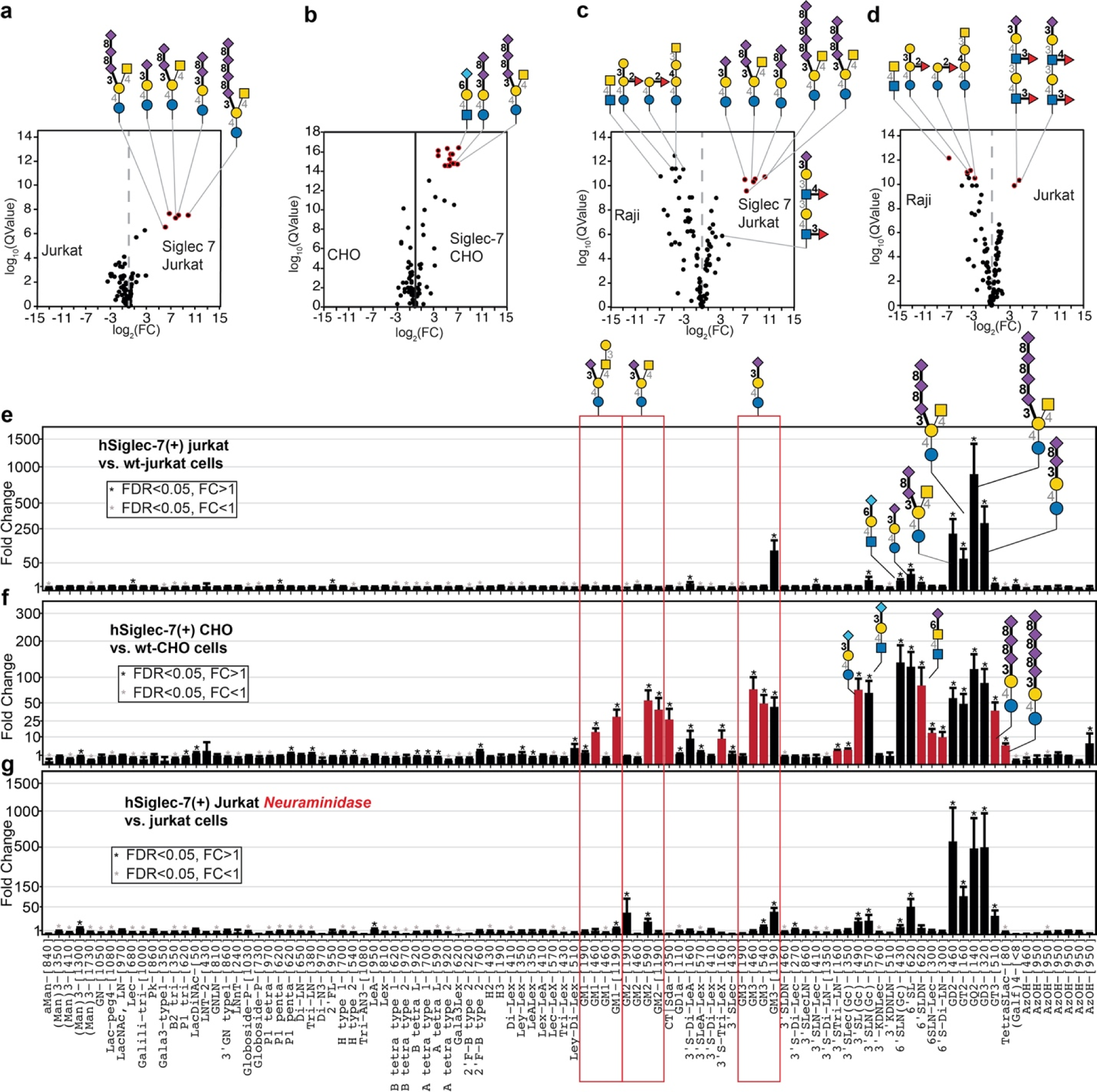
Siglec 7 enrichment profile on different cell milieu. **a-d)** Volcano plot of TMMnormalized sequencing data with Benjamini–Hochberg FDR correction of different cell lines probed with LiGA-100. Wild-type Jurkat and Raji cell comparison shown to demonstrate the endogenous glycan-binding preferences of different cell lines. **e)** LiGA binding to Siglec-7 expressed on Jurkat cells, and **f)** LiGA binding to Siglec-7 expressed on CHO cells. **g)** LiGA binding to Siglec-7 expressing Jurkat cells treated with neuraminidase. Neuraminidase treatment of Jurkat cells was not able to significantly alter the binding profile of Siglec-7. In **b-d**, the FC was calculated by EdgeR DE analysis using a negative binomial model, TMM normalization, and BH correction for FDR. Error bars represent s.d. propagated from the variance of the TMM19 normalized sequencing data.

The PCR product of samples containing unique multiplexing index barcodes are mixed and prepared for Illumina sequencing. Using 20 forward and 20 reverse barcoded primers (**Supplementary Table 1**), a total of 400 distinct multiplexed PCR products can be analyzed in one Illumina run. To ensure that each PCR product received similar sequencing coverage, we often run all PCR products on a 2% agarose gel and pool 20 ng of each DNA sample in a single mixture. The pooled DNA is further separated on 4% agarose gel and the cut-out DNA band is purified using a commercially available agarose gel purification kit. The pooled sample is processed as outlined in the Denature and Dilute Libraries Guide for NextSeq 500 with two exceptions; i) the pooled sample concentration is diluted to 1.2 pM and ii) the pooled sample is mixed with 1.2 pM of denatured PhiX. The resulting mixture of library and PhiX is loaded on the NextSeq 500/550 v2.5 Kit (2 × 75 Cycles) and the sample is sequenced using the Illumina NextSeq500 system.

We sequenced 3-7 replicate samples of the SDB phages associated with GBP^+^ and GBP^-^ cell pellets and used a LiGA-specific lookup table to assign copy numbers to each glycophage and determine fold change (FC) enrichment for each glycan. At the high level, the DE procedure can be approximated as a “*division*” with copy numbers of phage associated with GBP^-^ cells in a formal “*denominator*”. As the presence of near zero numbers in the denominator is undesired, we recommend running more replicate samples for GBP^-^ cells: these samples retain fewer phage particles and they are more likely to produce low quality / low copy number NGS data; in turn, low copy number reads in “denominator” inflate the noise of the DE procedure. Specific steps of NGS/DE analysis can be outlined as demultiplexing, read alignment, filtering, dictionary matching, grouping into “test” and “control” samples, normalization, and DE analysis. The sequencing data is de-multiplexed using the barcode sequences at both ends of the molecule identifying the original source sample. The paired-end sequencing used in this report gives an overlapping sequence from both ends of the molecule. These were merged into one sequence and the barcodes and primer binding sequences were removed from the ends. To filter out problematic DNA sequences, we eliminate the reads that i) do not match an expected barcode, ii) contain uncalled bases, or iii) contain sequence mismatch between the two ends. The rationale for these filtering steps were detailed in previous publications.^10,33^ Next, each DNA read is matched with DNA sequences of SDBs in the LiGA dictionary. We allowed for one base substitution between the observed DNA sequence and the DNA sequences in the Dictionary (Hamming distance 1, or H=1) and discarded all other reads as “unmapped”; the mapped H=1 variants are pooled and translated to a glycan-[density] using the Dictionary (**Supplementary Table 4-7 and Figure 2c**). Samples from multiple replicates of phages associated with GBP^+^ and GBP^-^ cells are merged in one table for differential enrichment (DE) analysis.

Quantification of binding of each glycan employs the DE pipeline from EdgeR adapted from DE analysis of gene expression.^34,35^ DE pipeline compares the normalized copy number of each glycophage associated with GBP^+^ (e.g., Siglec^+^ CHO), and GBP^-^ cells (e.g., Siglec^-^ CHO) over multiple replicates (**Figure 2d**). For DE-analysis, three factors were considered: (i) inferring the abundance of each SDB using a negative binomial distribution of the observed copy numbers; (ii) normalization of data across multiple samples using the Trimmed Mean of M-values (TMM) normalization (**Figure 2e-f**); and (iii) Benjamini–Hochberg (BH)^36^ correction to control the false discovery rate (FDR) at αLJ≤LJ0.05.^37^ Using this analysis, the final output of the DE pipeline is a table containing the following information; SDB, glycan, glycan-density, respective SDB abundances for each replicate of both GBP^+^ and GBP^-^ cells, the TMM normalized abundance, fold change (FC), p-value and the FDR corrected q-value (**Figure 2d).** The FC served as a metric for the binding of glycophage to GBP displayed on cells and the q-value estimates the statistical significance of the FC for each SDB.

**Figures 2e-g** describe the role of TMM normalization in data: the TMM normalization establishes a pseudo-baseline between the GBP^+^ and GBP^-^ cells (**Figure 2f**); without normalization, non-binding phage are depleted in GBP^+^ population and, hence, appear to be enriched in GBP^-^ cells (**Figure 2e**). Originally designed to analyze changes in gene expression from sequencing data, the TMM normalization assumes that the majority of the glycophages within a LiGA do not bind to GBP (i.e., they have indistinguishable binding to the GBP^+^ and GBP^-^ cells). If necessary, normalization can be adjusted to set a specific set of phage clones to FC=1. To this end, we modified multiple phage clones by DBCO and capped by azidoethanol; the normalization and DE algorithms can be run to set the population of “AzOH” clones to FC=1. Once FC and q-values are calculated by the DE pipeline, volcano plots can be used to visualize the FC respective and q-values for each glycophages, (**Figure 2h**) but our preference is to use bar charts to visualize the FC values along with statistical significance as (*). Such bar chart representation is used because it is identical to the format used to display binding data obtained using other glycan array experiments (**Figure 2h**). In a specific example in **Figure 2h**, the bar chart of DE analysis highlights seven glycophages significantly associating with Siglec-7^+^ Jurkat cells when compared to Siglec-7^-^ Jurkat cells (binding of the full array are represented in **Figure 3e**). The error bars in the bar charts display an estimate of variance propagated from the variance of reads in the replicates of test and control samples. The error bar is intended merely as a visualization tool and is not used for statistical significance analysis. The statistical test produced by the DE-pipeline is indicated by the asterisks which show the positive and negative samples have statistically significant difference FC with a q-value calculated using Benjamini–Hochberg (BH) correction for false discovery rate (FDR) below 5%.

### Application of protocol

LiGA, in principle, can be applied to any cell type to determine a cell surface receptor’s glycan binding preference. In this report, we focus on the cell-surface immunomodulatory receptors, Siglec-1, -2, -6, -7, and -9, and their corresponding arginine mutants^38^ using a LiGA containing 100 distinctly barcoded phages (termed LiGA-100). The LiGA-100 contains 78 different glycans with six of these glycans displayed at 2–4 different densities. To dissect Siglec-7 recognition, we incubated the LiGA-100 with Jurkat cells expressing Siglec-7 and wild-type Jurkat cells (**Figure 3a and Figure 3e**). DE-analysis uncovered seven glycans enriched by Siglec-7: GM3, 3’SLN, 6’SLN, GD2, GT2, GQ2, and GD3. The LiGA-100 contains other known glycan ligands of Siglec-7, however, we were unable to detect their binding to Siglec-7^+^ Jurkat cells.^39–42^ For example, the GT3 binds to Siglec-7 with relatively high affinity (*K*_d_ = 0.184 mM),^43^ yet the GT3-[510] glycophage was not enriched by Siglec-7 Jurkat cells. To test whether Jurkat cell glycocalyx^44^ influences the Siglec-7:glycan recognition, we measured the binding profile of Siglec-7 expression in another cell line (Siglec-7^+^ CHO, Chinese Hamster Ovary) and used isogenic Siglec-7^-^ CHO cells as reference (**Figure 3b and Figure 3f**). The DE analysis identified 15 additional (22 in total) saccharides that were significantly enriched in Siglec-7^+^ CHO, one of which was GT3-[510] (red bars, **Figure 3f**). These additional glycans were reported as Siglec-7 ligands using other binding techniques. For example, the enrichment of 6’SLDN decorated glycophages by Siglec-7 expressing CHO cells was in line with results determined using multivalent glycan probes by Paulson and co-workers.^45^ The differences in recognition profiles of Siglec-7^+^ CHO and Siglec-7^+^ Jurkat cells may result from *cis*-masking with sialic acids on the Jurkat cell surface. To test this hypothesis, we de-sialylated the Siglec-7^+^ Jurkat cells using Neuraminidase-A treatment but observed only a minor change in the binding of LiGA-100 to treated and untreated Siglec-7^+^ Jurkat cells (**Figure 3g**). Only a few glycans, such as the glycan 3’-SL (Gc), showed an increase in enrichment after treatment of cells by neuraminidase (**Figure 3g**). Based on the LiGA binding result of Neuraminidase-treated Siglec-7^+^ Jurkat cells, the differences between CHO and Jurkat cells cannot be fully explained by *cis*-inhibition on the surface of Jurkat cells. One possible explanation is that the 5-fold higher expression of Siglec-7^+^ on the surface of CHO cells likely provides a higher avidity to engage more ligands (**Supporting Figure 1**). Thus, the recognition of glycans by Siglec-7 on the cell surface is context dependent and we are not aware of any other systematic study that have observed such phenomena.

To test whether cell-line dependent recognition translates to other Siglec proteins, we tested the binding of LiGA-100 to Siglec-9 expressed on CHO cell and U937 that only express Siglec-9, which were created by generating a triple mutant with Siglec-3, -5, and -7 disrupted by CRISPR-CAS9 followed by lentiviral-mediated transduction with Siglec-9 (Sigelc-9^+^ U937^TKO^ cells).^46^ DE-analysis of LiGA-100 binding to Siglec-9^+^ CHO cells (compared to wild-type CHO cells) confirmed enrichment of 19 glycans and recognized an additional 4-7 glycans compared to binding to Sigelc-9^+^ U937^TKO^ cells; there was a significantly stronger overall recognition profile with lower baseline in Siglec9^+^ CHO data when compared to Siglec9^+^ U937^TKO^ cells (**Extended Data Fig. 3a-b**). Cell-line dependence of Siglec-glycan interactions, hence, is a recurrent observation.

From the 19 glycans enriched by Siglec-9^+^ U937^TKO^ (**Figure 4b**), some were also enriched by Siglec-7^+^ cells (**Figure 4b**) or CD22^+^ CHO cells (**Figure 4d**). For example, GM3-[1190] was enriched by Siglec-7^+^ Jurkat and CHO cells and Siglec-9^+^ U937^TKO^ and CHO cells. A minor preference of Siglec-9 for α2-6-linked sialic acid compared to α2-3-linked sialic acid was observed among the enriched sialyl glycans (**Figure 4b**). For example, the 6’ SLDN was enriched with a higher FC value compared to the GM3 by Siglec-7^+^ U937^TKO^ cells. As shown in **Figure 4b**, the Siglec-9^+^ U937^TKO^ cells enriched both α2–3- and α2–6-linked sialic acids, in line with previously reported ligand specificities of Siglec-9.^45,47^ We also observed that within the LiGA-100 binding, Siglec-9^+^ cells exhibited a minor preference for the α2–3-linked sialyl glycans linked to Galβ1–4Glc (GM3) compared to the α2–3-linked to GalNAcβ1–4GlcNAc (3’ SLDN) (**Figure 4b**). Comparing the enrichment profile of Siglec-9 and other platforms, we noted differences in binding. For example, the binding of GD1a to Siglec-9 was confirmed in a glycolipid microplate array in an *in vitro* assay, but only modest enrichment (FC = ∼2) above the baseline (FC = 1) was observed in LiGA.^48^ These results critically emphasize that Siglec-glycan recognition is context dependent and platform dependent. LiGA enables evaluation of this recognition in the most biologically relevant environment, on the surface of live cells.

**Figure 4:**
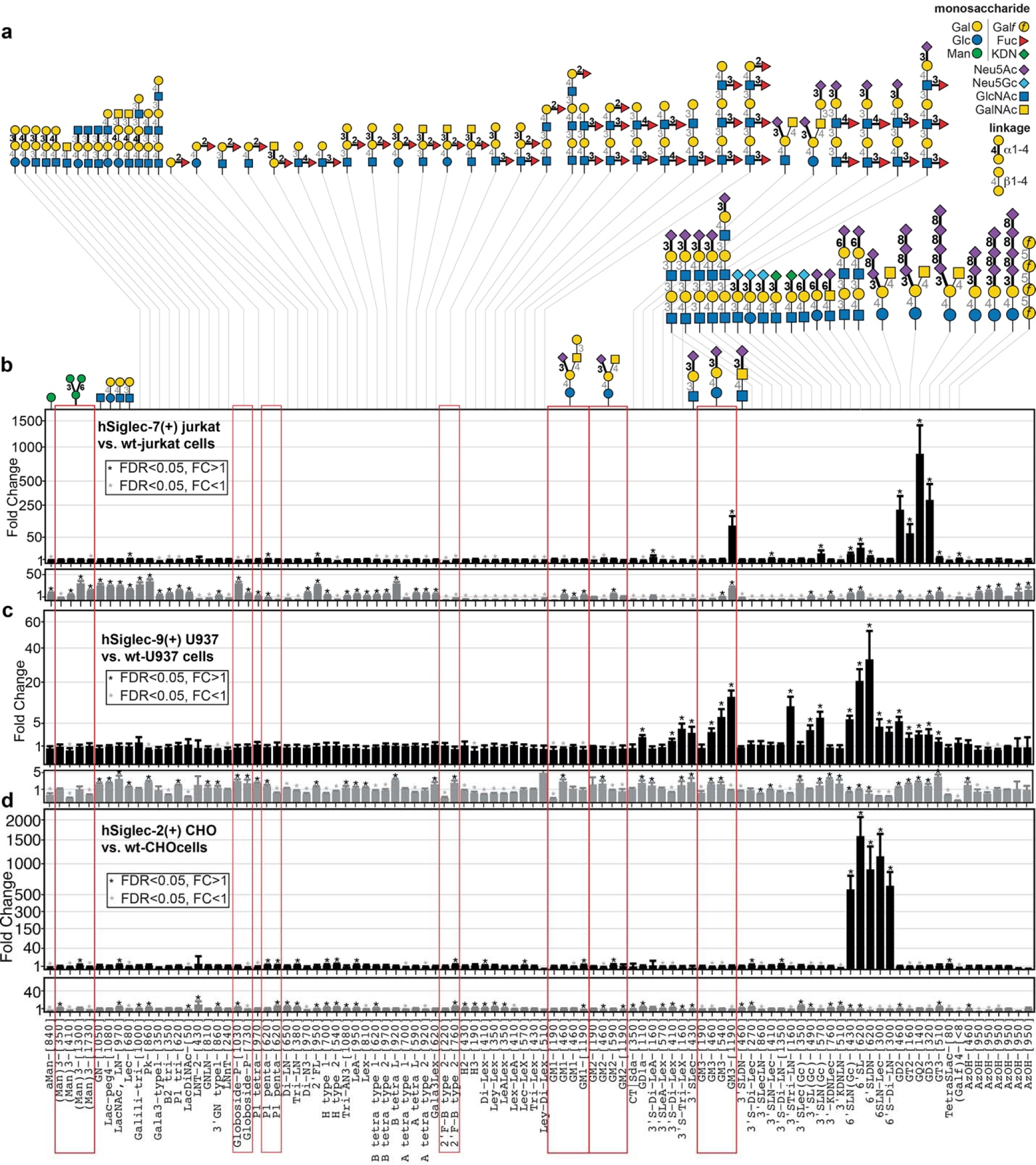
LiGA measures the binding specificity of Siglec-7, -9, hCD22 on distinct cell surfaces. **a**) Visual representation of the glycans in the LiGA. **b)** binding of LiGA to Siglec-7+ Jurkat cells normalized by binding to Jurkat cells. **c)** Binding of LiGA to Siglec-9+ U937 cells normalized by binding to U937 cells. **d)** Binding of LiGA to hCD22+ CHO cells normalized by binding to CHO cells. In **b**–**d**, the response of respective arginine mutant Siglec is represented below each bar chart. In **b-d**, the FC was calculated by EdgeR DE analysis using a negative binomial model, TMM normalization, and BH correction for FDR. Error bars represent s.d. propagated from the variance of the TMM-normalized sequencing data.

The guanidine functional group of the arginine side chain is crucial in the Siglec family of lectins for sialic acid binding. Accordingly, the arginine residue present within the ligand binding site in the V-set domain was mutated for Siglec-7, -9, and -2.^49^ The CHO cells expressing the arginine mutant Siglecs were incubated with LiGA-100, and the enrichment of each glycophage was measured using wildtype CHO cells as control. The binding of all sialo-glycans was abrogated when the essential arginine residue of Siglec-7, -9, and -2 was mutated (**Extended Data Figure 3b-c and Figure 4**). This observation confirms that the interaction occurs with a canonical binding pocket on the v-set domain, and the Siglec-expressing cells do not engage other classes of cell surface receptors for interaction with glycophages.

Unlike the relatively defined recognition profiles of Siglec-7, -9, and -2 favoring sialo-glycans compared to asialo glycans, the recognition profiles of Siglec-1 and -6 on the surface of CHO cells were significantly more complex (**Extended Data Figure 5 and Figure 5**). Incubation of LiGA-100 with CHO cells expressing Siglec-1 and isogenic CHO cells yielded a modest enrichment of six sialo glycans and three asialo-glycans (**Figure 5b**). One possible explanation for the modest enrichment is the low affinity of Siglec-1 for its cognate targets; the reported *K*_d_ of Siglec-1 and gangliosides are 1.2 mM (GM1), 0.9 mM (GM2), 0.5 mM (GM3), 1.3 mM (GD1a).^16^ Since the Siglec-1–ganglioside interaction has been established using other methods,^50,51^ we explored this interaction further on the surface of CHO cells using a focused, density scan LiGA. Five clonal phages, each with a unique SDB were mixed to create a multi-SDB (MSDB) and the MSDB was incubated with DBCO-NHS at a range of concentrations to pre-position a desired number of DBCO on each phage. Subsequently, azido-gangliosides were added to the MSDB-DBCO solution to create a MSDB-glycophages mixture. We termed the library LiGA-g which was created by mixing these individual MSDB-gangliosides that enabled 5 distinct binding event measurements within a single assay (**Figure 5c-d**). The ability to observe multiple technical replicates of glycophages within the same LiGA-g enabled us to rule out artifacts that may originate from minor preferences during the amplification or sequencing of individual DNA barcodes.

**Figure 5:**
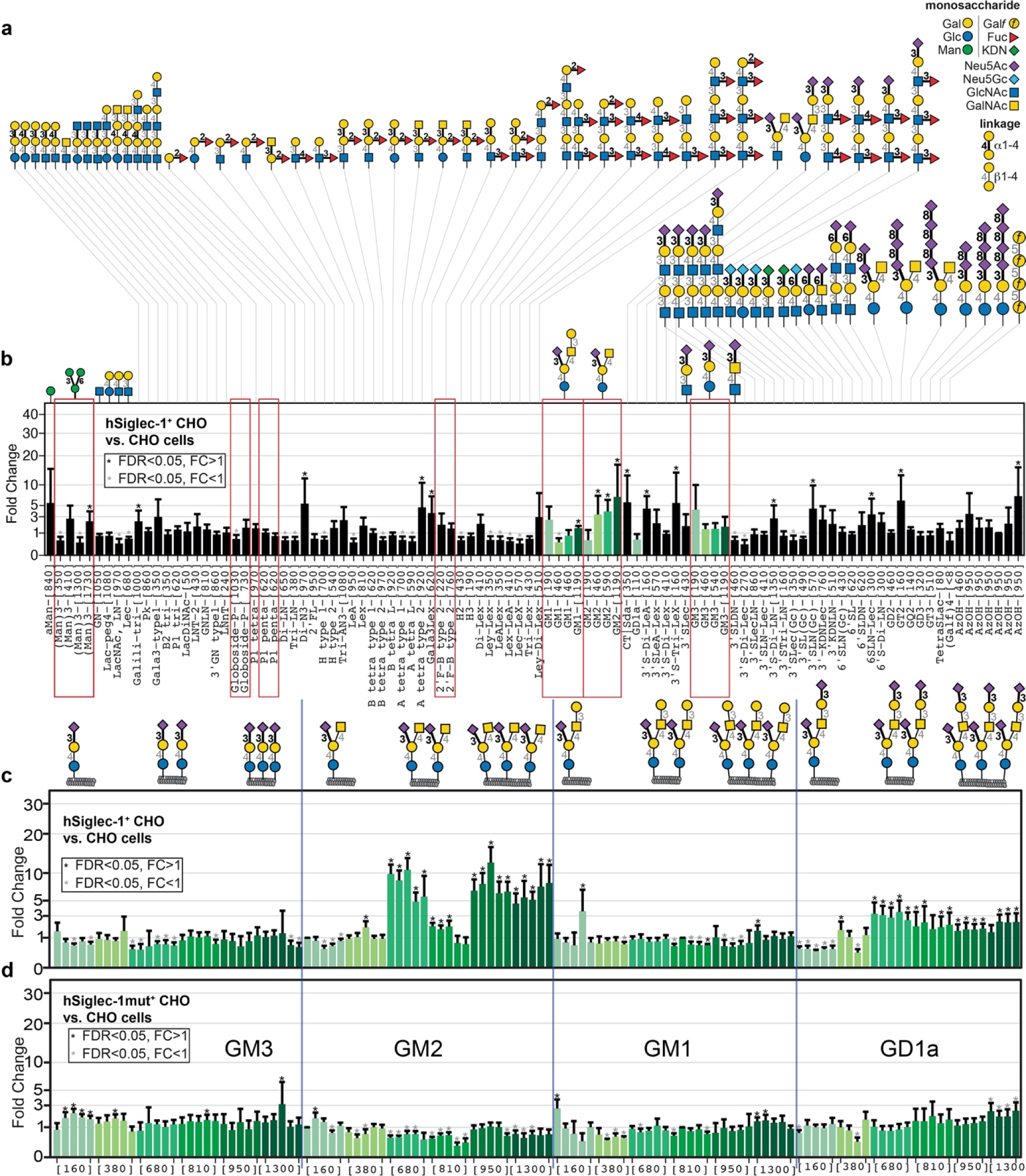
LiGA profiling of Siglec-1 displayed on CHO cells. **a**) Visual representation of the glycans in the LiGA. **b**) Siglec-1+ expressing CHO cells binding assay with LiGA-100. **c**) Enrichment of GM1, GM2, GM3, and GD1a to Siglec-1 expressing CHO cells. Binding was density dependent for GM2 and GD1a, while GM1 and GM3 did not show any binding. **d)** Siglec-1+ R116A mutant showed abrogation of binding for each of the ganglioside GM1, GM2, GM3 and GD1a. In **b-d**, the FC was calculated by EdgeR DE analysis using a negative binomial model, TMM normalization, and BH correction for FDR. Error bars represent s.d. propagated from the variance of the TMM-normalized sequencing data.

We measured the binding of LiGA to Siglec-1^+^ CHO cells, using the CHO cells as a control and deep sequenced the MSDB associated with the Siglec-1^+^ and Siglec^−^ cells. The resulting enrichment profile from Siglec-1^+^ cells uncovered a density dependence on the binding of the gangliosides GM2 and GD1a (**Figure 5c**). Phage particles that displayed >680 copies exhibited significantly higher binding than those with <680 copies of GM2 and GD1a. The enrichment level of GM2 plateaued at higher than 680 copy number, suggesting that a critical density of presentation of 680 copies is required for Siglec-1 recognition. We also observed a noticeable reduction in the enrichment of phage displaying 810 copies of GM2 when compared to 680 and 950 copies. The significance of this observation was reproducible in multiple biological replicates and was confirmed by 5 independent DNA barcodes. Similarly, GM1 and GD1a exhibit comparable monovalent affinities however,^16^ within LiGA-g we could only detect the binding of GD1a to Siglec-1^+^ CHO cells and not GM1 (**Figure 5c**). A minor decrease of enrichment was observed as density was gradually increased from 680 copies to 1350 copies of GD1a; which may be the result of steric occlusion. In a previous publication, we juxtaposed the binding of Siglec-1^+^ CHO cells to LiGA and liposomes decorated with GM1, GM2, GM3, and GD1a and observed a similar bimodal response with LiGA and liposomes. The liposomes decorated with GM1, GM2, GM3, and GD1a to Siglec-1^+^ CHO; notably, the liposomes modified with GM3 showed the lowest level of binding despite its highest monovalent affinity among the four tested gangliosides.^179^ This discrepancy in observations between LiGA, glycan-liposome, and monovalent affinity measured by mass spectrometry, highlights the subtle differences in presentation, linker structure, and mobility of gangliosides. To validate the binding of glycophages with the canonical ligand binding sites, we incubated the same LiGA-g with CHO cells expressing Siglec-1 R116A mutant, which abrogated the binding of all the glycophages modified with GM1, GM2, GM3, and GD1a (**Figure 5d**). These experiments highlight the ability of LiGA to measure robustly multiplexed measurements of both affinity- and avidity-based glycan–lectin interactions between glycans displayed on M13 bacteriophage capsids and lectins on the surface of mammalian cells.

Recognition of LiGA-100 by Siglec-6^+^ CHO cell was dramatically different from Siglec-1, -2, -7, and -9 expressing CHO cells. We observed 14 sialoglycans and 24 asialoglycans significantly enriched by Siglec-6^+^ CHO cells compared to the parental CHO cells (**Extended Data Figure 5b**). In line with a previous study that reported GM3 as a ligand of Siglec-6,^16^ we observed a significant enrichment of GM3 using LiGA-100 with the highest level of enrichment observed for phage modified with 1350 copies. Surprisingly, the FC of phages decorated with asialoglycan Le^y^-di-Le^x^ and Man_3_ was at a comparable FC of GM3 (**Extended Data Figure 5b**). To analyze Siglec-6 specificity in different milieu, we explored the LiGA-100 binding profile of purified Siglec-6 and Siglec-6 R122A mutant coated on wells (**Extended Data Figure 5c**). We observed negligible enrichment glycophages present within LiGA-100 by the wild-type and R122A Siglec-6 purified proteins. In contrast with the previously reported binding specificity of Siglec-6, the sialyl-Tn (Neu5Acα2-6GalNAcα-) was not enriched within the LiGA-100.^52–54^ We would like to note that our results do not preclude the possibility that the glycan ‘display’ format of LiGA limits Sigec-6 engagement, and the lipid hypothesis described in a recent publication^16^ corroborates these observations.

### Comparison to other technologies and limitations

The unexplored aspects of glycan’s role in biology have led to the convergence of several new platforms that utilize pre-synthesized glycans and bioanalytical techniques such as flow cytometry, chromatography, ELISA, and NGS to evaluate the binding profile of GBPs. LiGA, built using barcoded M13 phage uniquely combines multiplexing, encoding, monodisperse nano-carrier, and *in vivo*/*in vitro* compatibility. Liposomes displaying multivalent glycans have been used extensively to study cell surface receptors in live animals in vivo. A minor drawback of liposomes is the lack of encoding and multiplexing capabilities. In contrast, solid polymer microspheres, such as the Luminex beads can encode glycan identity using fluorescence;^55^ however, there are no reports of glycan-Luminex bead arrays used to study GBPs on live cells; solid microbeads, unlike liposomes and phage are unlikely to be compatible with injections *in vivo*.

Prior to LiGA, the display of *N*-glycans on the pIII protein of phagemid was achieved by Aebi and DeLisa research groups using bacterial glycosylation machinery to create biosynthetic ‘glyco-phagemid’.^56,57^ The use of such ‘glyco-phagemid’ was limited to the study of low valency interactions as only one glycan can be displayed per monovalent phagemid construct. The coupling of glycan synthesis with bacterial glycosylation machinery also restricts glycophage to display of the naturally produced glycans. Previous reports have utilized the ligation of glycans to DNA to create a library of glycans DNA-encoded library of glycans, however such a library lacked the ability to control the valency of presentation: a critical feature required to mimic multivalent glycan presentation on the surface of cells.^13,14^ To partially mitigate this issue, Wang and co-workers displayed three copies of glycans on a library of DNA hairpins.^15^ Nevertheless, the exposed DNA barcode is a liability, as it could be degraded by nucleases in a complex biological milieu and interfere with the binding of GBPs. Viral particles such as Qβ have unique advantages such as monodisperse morphology and size; they have been modified to display multiple copies of glycans to study glycan-GBP interactions in vitro and in vivo.^58–60^ However, Qβ particles used in such studies lack nucleic acids and cannot encode the glycan structure. It should be noted that the shape of Qβ (icosahedron) is different from M13 (rod-like); it may be beneficial to explore how these differences in shape influence the recognition of glycans by GBPs on the surface of cells. Therefore, future viral-particle-based glycan arrays could use DNA or RNA-barcoded viral capsids of different shapes to dissect the influence of multivalent architecture on glycan recognition.^61^

LiGA combines all critical features of the aforementioned glycan probes; the multivalent presentation and the biocompatibility of liposomes and Qβ phages, encoding of glycan structure, and multiplexing of Luminex beads. The capsid of the M13 phage protects the DNA barcode within the structure of the virion and enables the profiling of ligand–receptor interactions on the surface of live cells and in live animals. LiGA, like DEL, Luminex glycan arrays, and traditional glass-based arrays, is created by “brute force” parallel conjugation of one-glycan-at-a-time to a prospectively barcoded carrier; however, parallel conjugation of several glycans to several hundred carriers is a challenge solved in previous versions of glycan arrays and it is neither unique nor new to LiGA. Like glycans on DNA,^13^ LiGA can also be upgraded using the chemoenzymatic synthesis directly on phage using glycosyl transferases or glycosidases.^12^

The ‘unnatural’ linkage between the glycan and phages could be a potential weakness of the LiGA platform; the hydrophobic DBCO and triazole linking phage to pVIII may result in non-specific binding. We routinely subtract the effect of the DBCO linker using DBCO-phages capped with azido ethanol as a reference admixed to the same LiGA (**Figure 2-5**, “AzOH”). However, this strategy does not fully recapitulate the presentation of glycans on the surface of cells. Although our approach of LiGA can successfully decipher glycan-GBP interactions on the cell surface, future versions of LiGA can use a library of glycophages that display not only the glycan but also other classes of biological molecules that are naturally displayed by cells (ex. antigen, lipid chains and other receptor ligands).

### Conclusion

Here, we report the use of LiGA-100 and a focused glycan library LiGA-g to decipher the glycan binding specificity of five Siglec members and their Arg-mutants expressed on cell lines of distinct lineages. Since our initial report of LiGA, we have optimized critical parameters from the protocol for cell surface glycan-GBP profiling. The use of MSDB enabled us to measure multiple technical replicates within a single assay to decouple weak interactions of glycan to GBP on the cell surface. Using deep sequencing as a bioanalytical tool to measure glycan binding interaction is cost effective, it is widely available, and the data can be readily shared within the glycobiology community.

### Experimental design

The overall flow of the experiment is shown in **Figure 2**. The data processing flow is shown in **Figure 3**.

## Biological Materials

### Bacterial strains required for cloning and plaque isolation

- Bacterial strains: *E.coli* SS320 (Lucigen)
- *E. coli* K12 ER2738 (NEB)

## Reagents

### Common Reagents

- Tetracycline (Tc) (Sigma-Aldrich, cat. no. 87128)
- Tryptone (Gibco, cat. no. 211705)
- Yeast extract (Gibco, cat. no. 211929)
- Agar, bacteriological (Sigma-Aldrich, cat. no. A5306)
- EtOH, analytical grade (Sigma-Aldrich, cat. no. E7023)
- Qubit dsDNA (double-stranded DNA) HS (high sensitivity) assay kit (Thermo Fisher Scientific, cat. no. Q32851)
- Milli-Q H_2_O
- PhiX Control v3 (Illumina, cat. no. FC-110-3001)
- Orange DNA loading dye (6x) (Thermo Fisher, cat. no. R0631)
- NextSeq 500/500 high output kit v2.5 (75 cycles) (Illumina, cat. no. 20024906)
- SYBR™ Safe DNA Gel Stain (Invitrogen cat no. S33102)
- Bovine Serum Albumin (Sigma-Aldrich Cat No. A7906)
- *N,N*-Dimethylformamide (FischerSci cat no. D119-4)

## Reagent setup

### Lysogeny Broth (LB)

Add 10 g/liter tryptone, 5 g/liter yeast extract, and 5 g/liter NaCl to a 2 L Erlenmeyer flask; add Milli-Q H_2_O to desired volume and autoclave-sterilize. Store at room temperature (18–24 °C) for ≤1 month.

### LB agar (1.5% (wt/vol) agar)

Add 10 g/liter tryptone, 5 g/liter yeast extract, 5 g/liter NaCl, and 15 g/liter agar to a 2 L Erlenmeyer flask; add Milli-Q H_2_O to the desired volume and autoclave-sterilize the solution. Incubate at room temperature until it is 55 °C and add 40 mg/L IPTG and 50 mg/L X-Gal and mix the solution. Immediately, pour 20 mL per petri dish. Typically, 1 L of LB agar solution is enough for 40 petri dishes.

### PBS (10x)

The solution is composed of NaCl (1.37 M), KCl (27 mM), Na_2_HPO_4_ (100 mM), KH_2_PO_4_ (18 mM). For 1 L, mix 17.8 g of Na_2_HPO_4_, 2.4 g of KH_2_PO_4_, 80 g of NaCl, and 2 g of KCl in 700 mL of Milli-Q H_2_O. Once the salts have dissolved, volume to 1 L and autoclave-sterilize the solution.

### 50% (vol/vol) glycerol (100 mL)

Mix 50 mL of glycerol and 50 mL of autoclave sterilized Milli-Q H_2_O in a 200-mL glass bottle in a biological hood. Store at room temperature for ≤1 year. We use a fresh bottle of glycerol to make this solution and do not autoclave the solution to avoid glycerol decomposition.

### HEPES buffer (10x)

The solution is composed of HEPES (200 mM), NaCl (1.5 M), and CaCl_2_ (2 mM). Typically, 300 mL of the buffer is created by mixing 21.5 g of HEPES, 26.3 g of NaCl, and 0.7 g of CaCl_2_ in 200 mL of autoclaved sterilized Milli-Q H_2_O.

### Equipment

- 1.5-mL microcentrifuge tubes
- 0.7-mL microcentrifuge tubes
- Bacterial culture tubes
- Erlenmeyer flasks, baffled, 250 mL and 2 liters
- Petri dishes (FisherSci cat no. FB0875712)
- qPCR tubes, 8-Strips, 0.2 mL (BioRad cat no. TLS0801)
- qPCR caps, 8-Strips, (BioRad cat no. TCS0803)
- Falcon tubes, polypropylene, conical, 15 and 50 mL
- FACS tubes, polypropylene, round bottom, sterile, 5 mL (Corning, cat. no. 352063)
- Filter pipette tips, sterile, volumes: 1-1,000 uL
- Pipette tips, sterile, volumes: 1-1,000 uL
- Vortex mixer
- Thermocycler with heated lid
- Incubator (static) for bacterial plates
- Incubator (shaking) for liquid bacterial cultures
- Heat block, digital, 1.5-2.0-mL microcentrifuge tube capacity
- Benchtop microcentrifuge
- Qubit Flex Fluorometric (Thermo Fisher, cat. no. Q33327)

### Software

- (Recommended) RStudio, an integrated development environment for R (https://rstudio.com/products/rstudio/download/)
- R packages, including edgeR for differential analysis binding (https://bioconductor.org/packages/release/bioc/html/edgeR.html), as well as readxl, writexl, tidyverse, scales, ggplot2, ggrepel, ggiraph and dependencies.
- The differential-test.R script and “differential-testing” folder for identifying significantly enriched glycans. (https://github.com/derdalab/liga/tree/master/differential-testing).

## Experimental Procedure

### Phage library cloning

1. A library of degenerate codons, termed silent double barcodes (SDB), was created in the phage genome at a position proximal to the gIII cloning site starting from a M13KE vector containing the stuffer sequence CAGTTTACGTAGCTGCATCAGGGTGGAGGT corresponding to the peptide QFT*LHQ, with * representing an amber stop codon.
2. The insert fragment was PCR amplified using the primers **P1** and **P2**, and the vector fragment was PCR amplified using primers **P3** and **P4**. Sequences: **P1, P2**, **P3**, **P4**. Sequences of primers **P1–P5** are provided in the **Supplementary Table 3**.
3. PCR was performed using 50 ng phage dsDNA with 1 mM dNTPs, 0.5 µM primers, and 0.5 µL Phusion High Fidelity DNA polymerase in 1x PCR buffer (NEB #B0518S) in a total volume of 50 μL.
4. The temperature cycling protocol was performed as follows: a) 98 °C 3 min, b) 98 °C 30 s, c) 60 °C 30 s, d) 72 °C 4 min, e) repeat b)–d) for 35 cycles, f) 72 °C 10 min, g) 4 °C hold. The parent DNA template was removed via enzymatic digestion with restriction enzyme *Dpn*I (NEB #R0176S) and the PCR amplified fragments were gel purified. Next, NEBuilder Hifi DNA assembly was carried out following the manufacturer protocols by mixing 100 ng of vector, 4 ng insert, 10 μL of NEBuilder Hifi DNA assembly master mix (NEB #E2621S), and deionized H_2_O up to a total volume of 20 μL.
5. The resulting ligated DNA was transformed into *E. coli* K12 ER2738 and propagated overnight at 37 °C.
6. The overnight culture was centrifuged to separate the bacteriophage from the host cells and the supernatant was supplemented with 5% PEG-8000, 0.5 M NaCl to precipitate the phages.
7. The solution is incubated for 8 h at 4 °C followed by a 15 min centrifugation at 13, 000 ×g and the resulting phage pellet was resuspended in 50% glycerol-PBS solution.
8. The resulting phage library with SB1 QFT*LHQ was PCR and analyzed using Illumina sequencing.

**Note:** Deep sequencing of the resulting SB1 QFT*LHQ cloning vector with 6,144 theoretical sequence combinations in the leader region is available at: https://48hd.cloud/file/20161105-68OOooIC-NB.

9. Into the resulting vector SB1-QFT*LHQ, the degenerate codon library of the SVEKNDQKTYHAGGG peptide was cloned to produce a vector containing SB1-SVEKNDQKTYHAGG sequence.
10. The vector was PCR amplified using primers **P4** and **P6**, and the insert was amplified from annealed primers **P5** and **P2**.
11. PCR fragments were processed using the NEBuilder Hifi DNA assembly kit (NEB #E2621S) using the steps described above.
12. The resulting ligated DNA was transformed into electrocompetent cells *E.coli* SS320 (Lucigen) and propagated overnight.
13. The culture was centrifuged in 50 mL to remove host cells and incubated with 5% PEG-8000, 0.5 M NaCl for 8 h at 4 °C, followed by 15 min centrifugation at 13,000 ×g.
14. The PEG precipitated phage was re-suspended in PBS-Glycerol 50% and stored at –20 °C.
15. The resulting M13-SDB-SVEKY is a library of chemically identical phages with 6,144 theoretical redundant sequence combinations in the SB1 region (∼200 practically observed) and 2.1 × 10^6^ theoretical redundant sequence combinations in the SB2 region, yielding between 10^8^ to 10^10^ possible sequence combinations (**Extended Data Figure 1**).
16. The sequence of the M13-SDB-SVEKY vector is available on GeneBank (#MN865131). Deep sequencing of the M13-SDB-SVEKY and SB1-QFT*LHQ libraries is available at: https://48hd.cloud/file/20161215-67OOooOO-NB and https://48hd.cloud/file/20161105-68OOooIC-NB.

### SDB clone isolation and amplification

17. **Day 1**: Set a liquid culture of *E. coli* K12 ER2738 by inoculating a colony in 25 mL of LB in a 125 mL Erlenmeyer flask. Incubate the flask overnight at 37 °C with shaking at 250 RPM.
18. **Day 2:** Serially dilute a 10 µL aliquot of the M13-SDB-SVEKY library to 10^4^ PFU/mL solution in LB.
19. Mix the 10 uL of 10^4^ PFU/mL M13-SDB-SVEK library with 200 μL of *E. coli* K12 ER2738, and mix with 3 mL of top agar and immediately pour it on top of an agar plate.
20. Let the top agar solidify and incubate the plate in the 37 °C static incubator for 8-10 h.
21. Isolate single clones of phage by picking a plaque using a sterile p200 tip and resuspending the plaque in 0.5 mL of 50% glycerol-PBS solution in a sterile microcentrifuge tube (1.5 mL).
22. Incubate the tube at 55 °C for 15 min to kill any E. *coli*.
23. Use 10 μL from step 22 to inoculate 5 mL of LB with 0.5 mL of log phase *E. coli* K12 ER2738 in a 10 mL culture tube for 4.5 h with vigorous shaking.
24. Pellet cells by centrifugation at 4500 g for 10 min.
25. Phage DNA was extracted from the pellet using a GeneJET Plasmid Miniprep kit (Thermo Fisher, #K0502) and analyzed by Sanger sequencing.

**Note:** Sanger sequencing is optional to analyze the SDB sequence. Illumina sequencing can be used to detect a mixture of sequences that may be present within an isolated clone.

26. The supernatant was incubated with 5% PEG-8000, 0.5 M NaCl for 8 h at 4 °C, followed by 15 min centrifugation at 13000 ×g.
27. The pelleted phage particle was resuspended in 1 mL PBS-Glycerol 50%, titered, and stored at –20 °C.
28. The clonal phage was PCR amplified using a protocol described in “*PCR Protocol and Illumina sequencing*” and analyzed by Illumina sequencing. Data is available at https://48hd.cloud/ by searching for a specific SDB number. An example of searching for “SDB50” can be found at: https://48hd.cloud/search/SDB50/1.
29. We selected the clones that contained three or more base pair substitutions from one another (i.e., Hamming distance, H ≥ 3). H ≥ 3 permits the correction of any point mutations that may have arisen during the analysis by deep sequencing (**Supplementary Table 4**).

### Modification of Phage Clones with Glycans

30. A solution of each SDB phage clone (10^12^–10^13^ PFU/mL in PBS) was combined with DCBO-NHS (20 mM in DMF) to afford a 1 mM concentration of DCBO-NHS in the reaction mixture.

**Note:** This typically yields 25% of pVIII modification after 45 min incubation.

31. After reaction with DBCO-NHS, the SDB phage clone was purified on a Zeba™ Spin Desalting column (7K MWCO, 0.5 mL, Thermo Fisher, #89882).
32. The filtrate was evaluated for DBCO modification using MALDI.
33. Solutions of azido-glycans (10 mM stock in H_2_O) were added to the filtrates to afford a 2 mM concentration of glycan-azide, and the solutions were incubated further overnight at 4 °C.
34. The reaction progress was evaluated using MALDI.

**Critical Step:** If the reactions were incomplete and a residual pVIII-DBCO peak was detected in MALDI, we added an additional amount of azido glycan and extended the incubation further overnight at 4 °C. After this overnight incubation, we supplemented the reaction with 5 mM of azidoethanol to cap any unreacted DBCO.

35. The conjugates were purified by Zeba column and supplemented by and stored as 50% glycerol stock at –20 °C.

### Analysis of Glycosylation of Phage Samples by MALDI-TOF MS

36. The sinapinic acid matrix^62^ (Sigma, #D7927) was formed by deposition of two layers: layer 1 (10 mg/mL sinapinic acid in acetone-methanol, 4:1) and layer 2 (10 mg/mL sinapinic acid in acetonitrile:water, 1:1, with 0. 1% TFA).
37. In a typical sample preparation, 2 μL of phage solution in PBS was combined with 4 μL of layer 2, then a mixture of 1:1 layer 1: layer 2+phage was deposited, in that order, onto the MALDI inlet plate, ensuring that layer 1 is completely dry before adding layer 2+phage.
38. The spots were washed with 10 μL of water with 0.1% TFA. MS-MALDI-TOF spectra were recorded on AB Sciex Voyager Elite MALDI, a mass spectrometer equipped with MALDI-TOF pulsed nitrogen laser (337 nm) (3 ns pulse up to 300 µJ/pulse) operating in full scan MS in positive ionization mode.
39. Raw MALDI *.txt files were plotted by plotMALDI.m MatLab script, available in the SI/Maldi data folder. These steps (39-40) are optional but convenient for analyzing a large number of samples.
40. An auxiliary drawGlycan.m script used glycan names in a short-form IUPAC notation used by CFG to draw CFG representation of glycans, calculate the molecular weights of glycans, and calculate the MW of species anticipated to be observed in MALDI: (i) pVIII protein; (ii) DBCO-pVIII; (iii) glycosylated pVIII; (iv) glycosylated pVIII with lost sialic acid (for sialylated glycans only).

**Note**: The script fits the peaks in the areas around those masses to *h* exp(– (*m* – *m*_0_)^2^) + *am* + *b* (Gaussian peak with a linear baseline correction). The parameter *h* was used as a simple estimate of the peak height. To estimate the fraction of glycosylated pVIII proteins, let *a* be the height of the non-glycosylated pVIII peak and *b* that of the glycosylated pVIII peak. Then, the total amount of pVIII is *a* + *b*, and *b* / (*a* + *b*) is the fraction of glycosylated pVIII molecules. For multiple peaks (e.g., sialylated glycans) the *b* is the sum of peak heights for all the decomposition products. The absolute copy number of glycosylated pVIII proteins was calculated by multiplying the glycosylated pVIII fraction by 2700 and rounded to the nearest ten.

### Preparation of LiGA from Glycosylated Clones

41. LiGA was prepared by mixing equal volumes of 50% glycerol glycosylated clones obtained in step 35 in a single microcentrifuge tube.
42. The tube was stored at –20 °C and given a two-letter unique identified (e.g., EF), which corresponds to a “dictionary” (EF.xlsx), a table that describes the correspondence between DNA barcodes and the glycan structure and its density of presentation (**Supplementary Table 5-7**).

**Note:** These dictionaries were used to translate DNA in the deep-sequencing files to the corresponding glycans and their density.

43. The mixture was characterized by titering.

### Cell Panning Procedure

44. Confluent GBP^+^ and GBP^-^ lectin cells are resuspended in an incubation buffer at a final concentration of 4 × 10^6^ cells/mL by pipetting up and down.

**Critical Step:** It is recommended to have five replicates of each cell type for better statistics during differential enrichment analysis.

45. Label FACS tubes (Corning, #352054) with a sharpie (e.g., “hCD22-CHO” and “CHO”).
46. Pipette 1 × 10^6^ cells (250 µL) into each FACS tube.
47. Add LiGA to each FACS tube such that N × 10^6^ PFU (N = glycan-phage conjugates) is incubated with 1 × 10^6^ cells.
48. Incubate the cells with LiGA for 1 hour at 37 °C.

**Critical Step:** A gentle vortex every 15 min during the incubation is recommended.

49. After incubation, add 1 mL of wash buffer into each FACS tube containing cells with a squeeze bottle (use of a serological pipette is an alternative method).
50. After incubation, add 1 mL of wash buffer into each FACS tube containing cells with a squeeze bottle (use of a serological pipette is an alternative method).

**Critical Step:** Avoid pipetting directly into the tube and aim for the walls of the FACS tube to minimize foaming and bubbles.

51. Gently vortex the cells to create a homogenous mixture with the wash buffer.
52. Add 2 mL of wash buffer into each FACS tube to a final volume of 3 mL washing buffer.
53. Centrifuge the tubes at 220 ×g for 5 min at 4 °C in a swinging bucket rotor.
54. Remove the supernatant by inverting the tubes over a liquid disposal container and counting to 5.
55. Then, press the rim edge of the tube while inverted against the Kimwipe to remove any residual liquid.
56. Place the tubes back upright into a rack.
57. Repeat steps 53-57 for a second wash.
58. Resuspend the pellet in 1 mL of HEPES buffer and transfer the resuspended cells to a 1.7 mL labeled microcentrifuge tube.
59. Centrifuge the tubes at 220 ×g for 5 min at 4 °C in a swinging bucket rotor.

### Elution of DNA from Bound Phage (Proteinase K Treatment)

**Note:** This method supersedes the boiling method reported in previous publications.

60. Re-suspend the cell pellet from step 50 in 50 µL of 1x HF buffer (Thermo Scientific, Cat# F518L) with 0.1 mg/mL RNase A.
61. Vortex the resuspended cells for 1 min at high speed.

**Note**: The cells will lyse, and their nuclei will stick together to make visible chromatin flakes.

62. Centrifuge the cell suspension for 3 min at 500 ×g.
63. Transfer the supernatant to a labeled 1.7 mL tube.
64. Add 1 µL of 1 mg/mL Proteinase K and incubate for 10 min at 55 °C in a heat block.
65. Incubate the tubes at 97 °C for 3 min to deactivate the Proteinase K.
66. Centrifuge the tubes at 21,000 ×g (14,000 rpm) in a microcentrifuge for 5 min.
67. Transfer the supernatant to a new tube and use the solution for PCR. Perform the first step of PCR with this supernatant on the same day.

**Pause point:** The product of the PCR can be stored for a long term at –20 °C.

### PCR –Single step

**Caution**: the single step PCR is a legacy procedure used in previous publications; it is recommended to use only for high-copy number samples!

68. Reaction mixture:

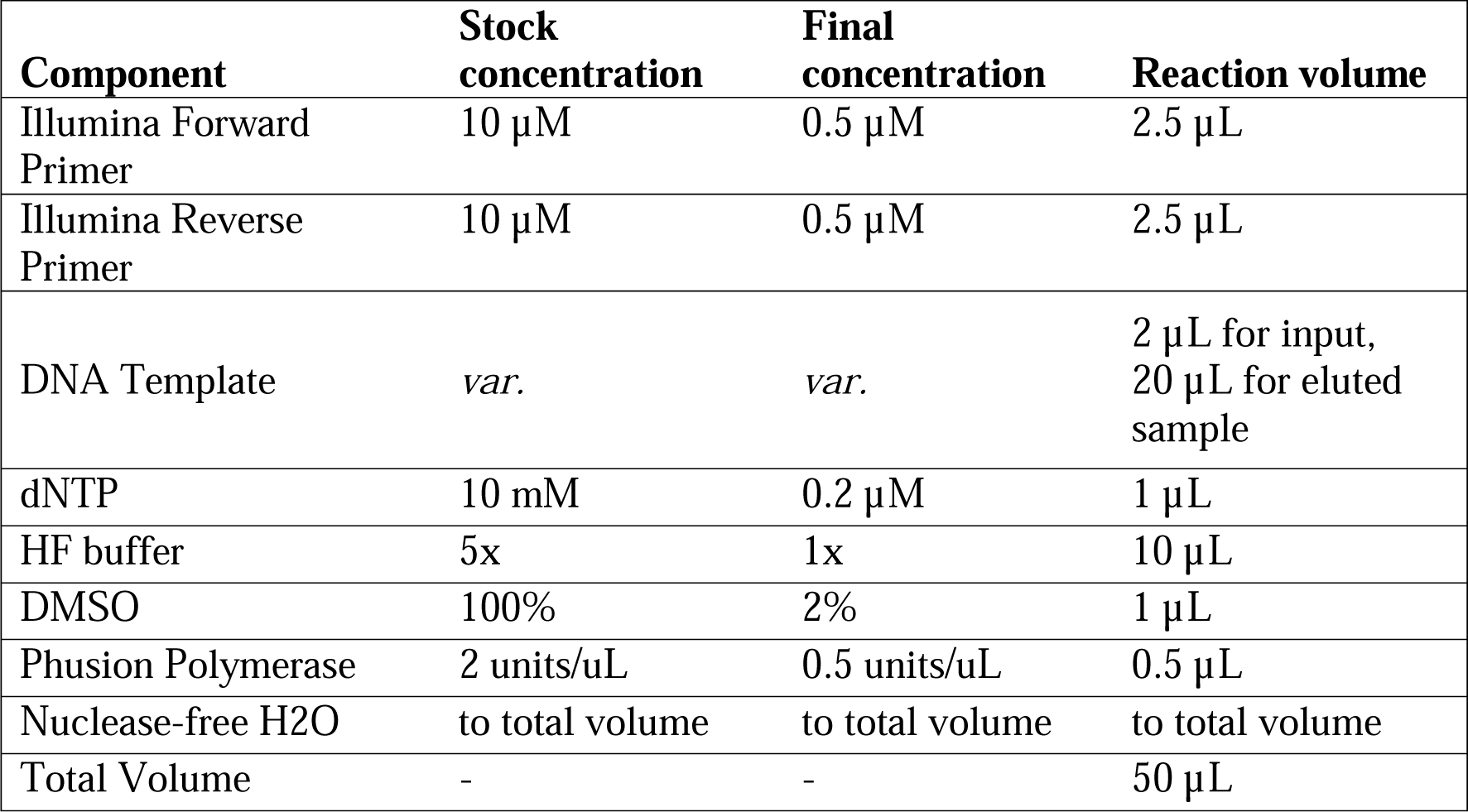
69. Thermocycler settings for 1-step PCR procedure:

69. Thermocycler settings for 1-step PCR procedure:
| Temperature | Time | No of cycles |
| --- | --- | --- |
| 95 °C | 5 min | 1 |
| 95 °C | 10 sec | 35 |
| 50 °C | 30 sec |  |
| 72 °C | 20 sec |  |
| 72 °C | 5 min | 1 |
| 4 °C | *Hold* | *Hold* |

### PCR – 2 Step procedure

**Note:** This method supersedes the single-step PCR method outlined in steps 68-69.

70. The reaction mixture for the first step:

71. Thermocycler settings for the first step:

| Component | Stock concentration | Final concentration | Reaction volume |
| --- | --- | --- | --- |
| NF10 (Forward Primer) | 10 μM | 0.5 μM | 2.5 μL |
| -96 (Reverse Primer) | 10 μM | 0.5 μM | 2.5 μL |
| DNA Template | *var.* | *var.* | 2 μL for input,<br>20 μL for eluted sample |
| dNTP | 10 mM | 0.2 μM | 1 μL |
| HF buffer | 5x | 1x | 10 μL |
| DMSO | 100% | 2% | 1 μL |
| Phusion Polymerase | 2 units/μL | 0.5 units/μL | 0.5 μL |
| Nuclease-free H<sub>2</sub>O | to final volume | to final volume |  |
| Total | - | - | 50 μL |

71. Thermocycler settings for the first step:
| Temperature | Time | No of cycles |
| --- | --- | --- |
| 95 °C | 5 min | 1 |
| 95 °C | 10 sec | 30 |
| 50 °C | 30 sec |  |
| 72 °C | 20 sec |  |
| 72 °C | 5 min | 1 |
| 4 °C | *Hold* | *Hold* |

**Pause Point:** The PCR product of step 71 can be stored at –20 °C until the next steps.

72. Reaction mixture for the second step:

73. Thermocycler settings for the second step:

**Pause Point:** The PCR product of step 71 can be stored at –20 °C until the next steps. 72. Reaction mixture for the second step:
| **Component** | **Stock concentration** | **Final concentration** | **Reaction volume** |
| --- | --- | --- | --- |
| Illumina Forward Primer | 10 uM | 0.5 μM | 2.5 μL |
| Illumina Reverse Primer | 10 uM | 0.5 μM | 2.5 μL |
| DNA Template | *var.* | *var.* | 5 μL of PCR mixture from "step 71" |
| dNTP | 10 mM | 0.2 μM | 1 μL |
| HF buffer | 5x | 1x | 10 μL |
| Phusion Polymerase | 2 units/uL | 0.5 units/uL | 0.5 μL |
| Nuclease-free H<sub>2</sub>O | to final volume | - | 28.5 μL |
| Total | - | - | 50 μL |

73. Thermocycler settings for the second step:
| **Temperature** | **Time** | **No of cycles** |
| --- | --- | --- |
| 95 °C | 5 min | 1 |
| 95 °C | 10 sec | 15 |
| 50 °C | 30 sec |  |
| 72 °C | 20 sec |  |
| 72 °C | 5 min | 1 |
| 4 °C | *Hold* | *Hold* |

### Preparing of DNA agarose gel

74. Prepare a 2% agarose gel by mixing 1 g of agarose with 50 mL of TAE (1x) buffer in a 125 mL Erlenmeyer flask.
75. Heat the mixture in a microwave. Commonly, the mixture is heated for 20 s intervals and mixed by swirling the flask.
76. Once the agarose has dissolved, take the flask out of the microwave and let it cool on a bench top until the temperature reaches 65 °C.
77. During the time agarose gel is cooling, place the gel tray into the casting apparatus and place a comb into the gel mold to create the wells. Alternatively, one may also use tape on the open edges of a gel tray to create a gel mold. We typically use a 15-well comb to set up one gel.
78. Add the SYBR Safe DNA gel stain to the agarose mixture according to the manufacturer’s instructions and swirl the flask to mix. Typically, we use 5 μL of the SYBR™ Safe DNA Gel Stain to a 50 mL solution of agarose.
79. Pour the agarose mixture into the gel mold and let it cool to temperature. The gel should solidify within 15 min at room temperature. To preserve the integrity of the SYBR safe DNA stain, we cover the gel mold with a tin foil to avoid photo-bleaching.

### Gel quantification of DNA Fragments

80. Aliquot 5 μL of the PCR samples from step 73 into a new PCR tube and add the 6x loading dye to the solution. We add 1 μL of the 6x Orange G DNA loading dye to the 5 μL of the PCR samples.
81. Transfer the 6 μL of the mixture into a gel well.
82. For quantification, mix 1 μL of the 50 basepair DNA ladder with 1 μL of the 6x Orange DNA loading dye and add to the gel.
83. Program the power supply to the desired voltage and monitor the progress of the band with Orange Dye progressing down the gel.
84. Once the DNA loading dye band has reached the end of the gel, quantify the gel band using a gel imaging system, with the DNA ladder band being the control. The **Extended Data Figure 2e** shows a representation of the DNA fragment because of the PCR.
85. Using the information from step 84, mix 40 ng of each sample into a tube and process it further for gel purification.

**Pause Point:** The pooled library can be stored at –20 °C until ready to proceed to purification and sequencing.

**Note**: To ensure proper purification, we restricted the pooling of libraries of the same size together. Over 3-4 years (30 sequencing runs, >5000 samples), we observed that optimal PCR protocols almost always produce PCR product bands with indistinguishable (high) intensity. For such samples ∼2 μL of unpurified PCR product from each sample can be simply mixed without prior quantification by gel. In rare cases, PCR products produce low-intensity bands; we frequently attempted to “rescue” such samples—if band intensity is 10× lower than average, we supplied a 10× higher volume of sample. These attempts were generally unsuccessful and produced few reliable reads in downstream NGS. If low intensity PCR product bands are observed, we recommend rerunning the procedure because it is likely that DNA associated with such bands cannot be reliably sequenced.

### Gel Purification of the DNA fragment

86. Prepare a 4% agarose gel by mixing 2 g of agarose with 50 mL of TAE (1x) buffer in a 125 mL Erlenmeyer flask.
87. Follow steps 75 to 79 to setup a gel with one modification: 8-well comb is used to accommodate a larger volume of sample loading.

### PCR product purification and Qubit quantification

88. The pooled library sample from Step 85 is mixed with 6x Orange G DNA loading dye and loaded onto the 4% agarose gel setup in Step 87.
89. Program the power supply to the desired voltage and monitor the progress of the band with Orange Dye progressing down the gel.
90. Once the DNA loading dye band has reached the end of the gel, use a commercially available gel purification kit to isolate the DNA band corresponding to the SDB barcode.
91. The protocol outlined in the kit is followed except for one exception: the DNA is eluted using two volumes of 20 μL of Nuclease free H_2_O.

**Pause Point:** Store DNA at –20 °C until ready to proceed to Qubit quantification and sequencing.

92. The purified samples are quantified using a Qubit dsDNA HS assay kit according to the manufacturer’s instructions.

**Critical Step:** To ensure proper quantification of the pooled sample library, three replicates are averaged for sample concentration measured using Qubit Fluorometric Quantification with recalibration performed after each replicate.

93. The pooled sample is diluted to 4 nM and quantified using the Qubit dsDNA HS assay kit according to the manufacturer’s instructions.
94. The library is further diluted to 1.2 pM following the procedure outlined in the NextSeq 500 and NextSeq 550 Sequencing Systems Denature and Dilute Libraries Guide.

**Note:** The loading concentration of the library is 1.8 pM according to the procedure outlined in the library guide, however in this protocol we load 1.2 pM due to the low diversity of the library.

95. Sequence the (double-ended) for 75 cycles. After sequencing, FASTQ files are automatically generated and uploaded to Illumina BaseSpace.

### Data Analysis

96. The Gzip compressed FASTQ files were downloaded from BaseSpace™ Sequence Hub.
97. The processing starts from the alignment of sequences from Forward and Reverse reads and mapping the multiplexing barcode sequences that correspond to individual experiments.
98. Within these files, the F and R strands are identified and aligned, allowing for no more than one point mutation per adapter region.
99. The adapter and multiplexing regions are trimmed leaving behind the DNA sequence flanking the adapter sequences.
100. The final output of the pipeline is plain-text-based lists of DNA sequences of the SDB region and their abundances. Technical replicates are often combined into a single text file.

**Note:** Intermediate data files that contain F and R aligned sequences with untrimmed adapters can be used to evaluate the overall fidelity of sequencing and processing (e.g., depth of sequences and number of reads that match the pattern that corresponds to the SDB insert).

101. DNA sequences are compared to the sequence of known SDB barcodes, and each sequence is converted to a recognizable SDB allowing for one substitution within the SDB sequence (Hamming distance = 1, H=1). All unrecognized sequences are marked as unknown; copy numbers of recognized SDB are pooled; and SDB that are present in the LiGA dictionary are converted to glycan-[density].
102. In preparation for DE analysis, the table is created that combines glycan-[density] data from the “test” experiment (binding of LiGA to GBP+ cell), “control” experiment (binding of LiGA to GBP-cells), and input (composition of LiGA before binding to cells).

**Note:** The copy number of each glycophage can be inspected to assess whether data is suitable for the calculation of the enrichment of glycans. We recommend looking for problematic traits associated with the reproducibility of replicates: (i) near zero copy numbers for many glycans in GBP^-^ samples (possibly due to low read depth); (ii) extreme deviation of reads in some replicates from reads all others replicates; (iii) significant fraction of “unmapped reads” and contaminating DNA sequences in some replicates. If these problems are detected, a decision must be made whether these “damaged” replicates should be removed prior to DE analysis. We also recommend examining the performance of each glycan/SDB and flag glycans/SBD which contain near zero copy numbers in input LiGA and LiGA affiliated with GBP^-^ cells.

Interpretation of enrichment for such glycans might not be possible; the samples might need to be sequenced at higher depth or the LiGA might have to be re-mixed to include a higher amount of the low abundance glycan/SDB.

### Differential enrichment (DE) analysis

103. The DE enrichment pipeline starts with setting up a campaign file which outlines the comparison to perform for each glycan-[density] copy numbers for “test” (e.g. Siglec-7^+^ CHO cells), “control” (e.g. Siglec-7^-^ CHO cells) and input library (**Figure 2d**).
104. Using the EdgeR package, the counts are modeled using a negative binomial function and normalized using TMM normalization across multiple replicates.
105. Appropriate modeling of the counts using a negative binomial model.
106. The fold-change (FC) is determined as a ratio of the normalized count of an SDB in the “test” data set and the normalized count of the same SDB in the “control” data set.
107. The P-value is calculated for the statistical significance of normalized counts in the “test” and “control” data sets. The Benjamini–Hochberg (BH) correction is applied to the P-value to control the false discovery rate (FDR) at α = 0.05 Q-value.
108. The FC plotted as a bar plot with * indicating significant differences between the “test” and “control” data set.

## Reagents Required

### Primers Required

Note: All DNA primers were obtained from Integrated DNA Technologies Inc. The sequences of primers are listed in **Supplementary Table 1-3**.

### Equipment required for binding assay

A. Pipette (Gilson, p1000, p100, p10)
B. Squeeze bottle (FischerSci Cat# 03-409-10D)
C. 15 mL Falcon tubes (FischerSci Cat# 14-959-70C)
D. 50 mL Falcon tubes (FischerSci Cat# 14-959-49A)
E. 1.5 mL microcentrifuge tubes (FischerSci Cat# 14-222-155)
F. FACS tubes (FisherSci Cat# 14-959-2A)
G. PCR machine (Biorad C1000 Touch thermal cycler)
H. qPCR machine (Biorad CFX96 Real-Time system)
I. qPCR tubes (Biorad Cat# TCS0803)
J. Benchtop centrifuge (Eppendorf centrifuge Cat# 5920R)

### Software required for analysis

A. R (programming language, R version 4.2.1)
B. RStudio (Version 2023.06.1+524)
C. GitHub master (https://github.com/derdalab/liga)
D. Differential Enrichment (https://github.com/derdalab/liga/tree/master/differential-testing)

### Reagents required

A. 1× HEPES buffer (20 mM HEPES, 150 mM NaCl, 2 mM CaCl_2_, pH 7.4, sterile filter using 0.22 µm bottle top filter)
B. RNase A (SigmaAldrich Cat# 556746)
C. Prepare 0.1 mg/mL solution of RNase A in 1× HF buffer (Cat #B0518S)
D. Proteinase-K (Thermo Fischer Cat# 25530049)
E. Prepare a 10 mg/mL solution by mixing the stock solution with 1× HEPES buffer in a 1:1 ratio.
F. 50 BP DNA ladder (NEB Cat# N3236S)

### General Procedure

#### Cell Culture Methods

Full-length hCD22, Siglec-7, and respective mutants were transfected in the pcDNA5/FRT vector and then were transfected stably into the CHO cell line through the Flp-In^TM^ (ThermoFisher, #K601001) system under selection with 0.5 mg/mL hygromycin-B (ThermoFisher, #10687010) for two weeks. The cells were passaged using TrypLE Express (ThermoFischer, # 12605036) as a dissociation solution and maintained in DMEM-F12 medium supplemented with 10% Fetal Bovine Serum and 1% Penicillin-Streptomycin (ThermoFisher #11330057, #26140079, #15140122). Untransfected CHO cells were maintained in the same conditions and used as a control. Confluent mammalian cells Siglec^+^ and CHO-wt cells were detached from the culture flask using TrypLE Express and centrifuged for 5 min at 300 ×g. The cell pellet was resuspended in incubation buffer.

The Siglec-7^+^ Jurkat and Siglec-9^+^ U937^TKO^ cells were maintained in RPMI medium supplemented with 10% Fetal Bovine Serum and 1% Penicillin-Streptomycin (Thermo Fisher #12633012, #26140079, #15140122). Wild-type Jurkat and Siglec-9^-^ U937^TKO^ cells were maintained in the same conditions and used as a control. Confluent mammalian cells were centrifuged for 5 min at 300 ×g, and the resulting cell pellet was resuspended in incubation buffer.

### Illumina Sequencing Data Processing

The Gzip compressed FASTQ files were downloaded from BaseSpace™ Sequence Hub. The files were converted into tables of DNA sequences and their counts per experiment, essentially as previously described.^33^ Briefly, FASTQ files were parsed into separate files based on unique multiplexing barcodes within the reads. Reads that did not contain an identifiable multiplex barcode were discarded. Reads that contained a Phred = 0 quality score in any position also were discarded. Additional quality control was based on: (i) mapping the forward and reverse primer regions, allowing no more than one base substitution each, (ii) alignment of the forward and reverse read-end overlap, allowing no mismatches between forward and reverse read in the overlap region. Reads that did not match criteria (i) and (ii) were discarded. The two ends of read-pairs that pass the filtering criteria were joined and trimmed to the DNA sequences located between the priming regions; the reads were organized in a tab-delimited text file containing the unique DNA sequences and their copy numbers. Technical replicates often were combined in the same file.

The files with DNA reads, raw counts, and mapped glycans were uploaded to http://ligacloud.ca/ server. Each experiment has a unique alphanumeric name (e.g., 20180711-87YOrdRB-JM) and unique static URL: for example, http://ligacloud.ca/searchLibInfo?f=0&b=0&d=20180711-87YOrdRB-JM. Data files also can be searched for by any metadata associated with the experiment (e.g., target name, experimental conditions).

## Supporting information

File contains supplementary tables and figure

Zipped folder containing all the software, MALDI TOF spectra and analyzed files

---

**Extended Data Figure 1:**
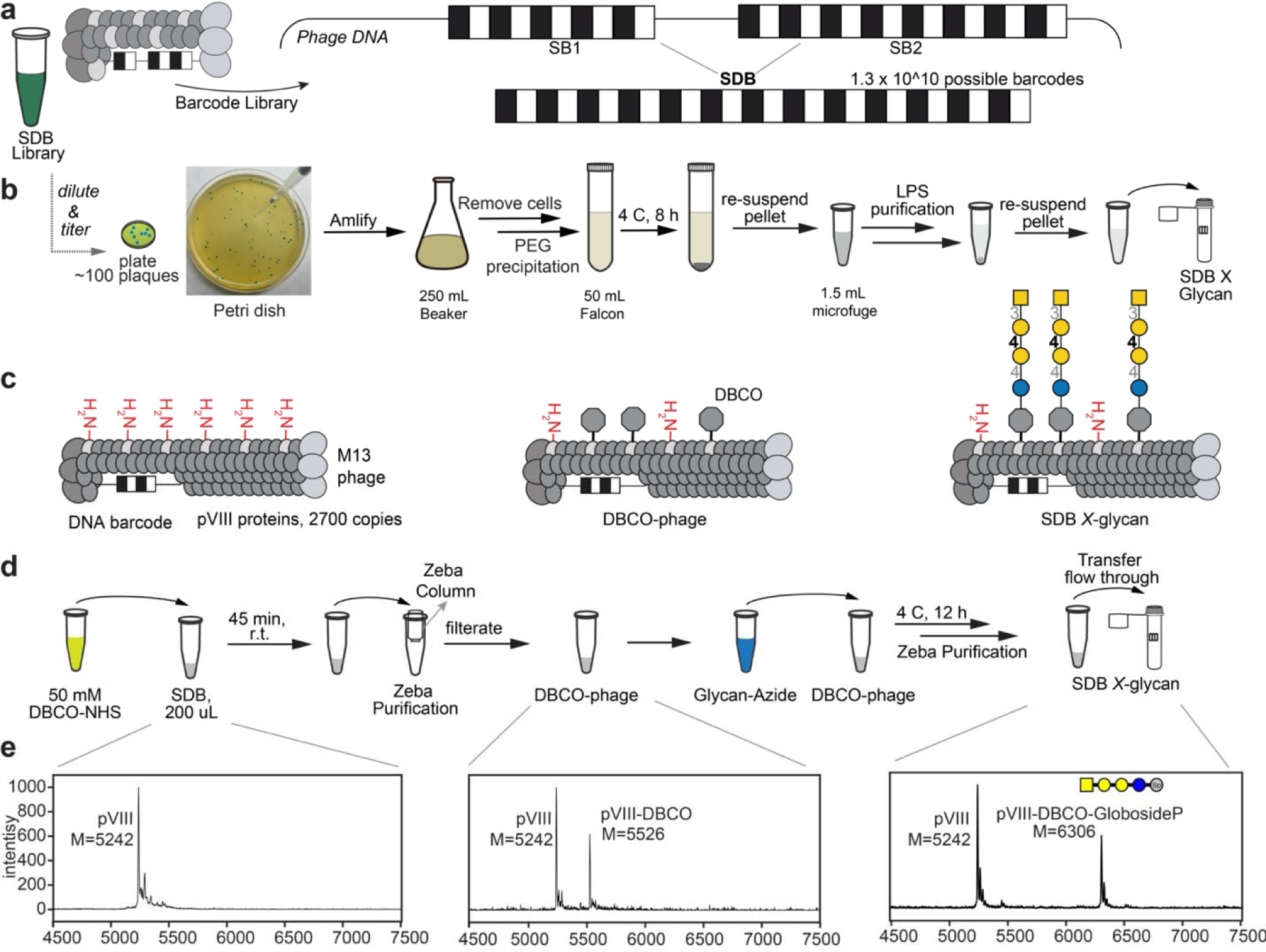
Describes the workflow for generating the encoding glycan using SDB M13 phage. **a)** The M13KE SDB SVEK library is a combination of two degenerate codon regions in the phage DNA: SB1 and SB2. The combination of SB1 and SB2 yields a total of 1.3 × 1010 DNA possible barcoded phages that are phenotypically identical. **b)** The M13KE SDB SVEK library was plated at ∼100 plaques per plate. Each plaque (clone) was isolated, individually amplified, and purified with Triton X-100 and PEG to remove LPS. **c)** Schematic showing unmodified phage, DBCO-modified phage, and phage with azido-glycan. **d)** Workflow for modification of clonal phage with a distinct barcode. First, the phage is reacted with DBCONHS and verified by MALDI-TOF. Azide glycan is ligated with the DBCO on the phage. **e)** Typical MALDI-TOF spectra of unmodified phage, phage modified with DBCO, and after cycloaddition of azide glycan.

**Extended Data Fig. 2:**
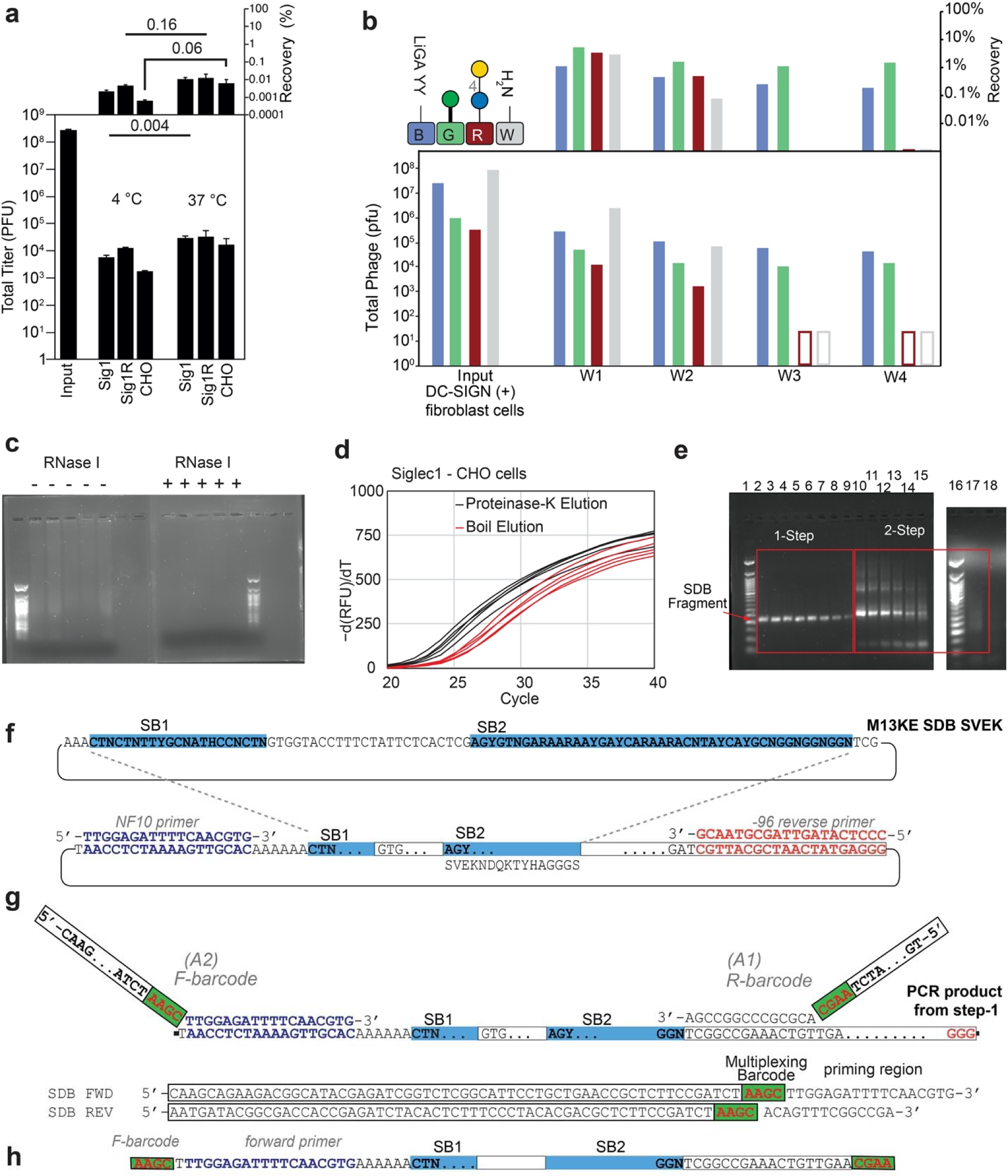
Optimizing LiGA binding, washing, elution, and PCR. **a**) Incubation at 37 °C consistently showed higher amounts of eluted phage particles. **b**) Titer results describe the binding of a specific reporter phage decorated with DC-SIGN binding glycan (αMan-green) over multiple washes. **c**) RNase I treatment to degrade cellular RNA. **d)** The amount of phage eluted was consistently higher with Proteinase K treatment compared to boiling the cell pellet. **e)** Gel describes a one-step and two-step PCR product. Lane 1 and 16 are DNA 50 bp ladder. Lanes 2-9 are PCR of phage sample in decreasing concentration of phage from 108 to 101 PFU using NF10 and -96 primers. Lanes 10-18 are PCR of phage sample in decreasing concentration of phage from 108 to 101 PFU using F1 and R1 primers. **f)** PCR using no-overhang primers annealing to regions outside of the SB1 and SB2 regions. **g)** Second PCR appends the Illumina adapters and multiplexing barcodes to the PCR product from the first step (**f**). **h)** Sequence of the final DNA fragment.

**Extended Data Fig. 3:**
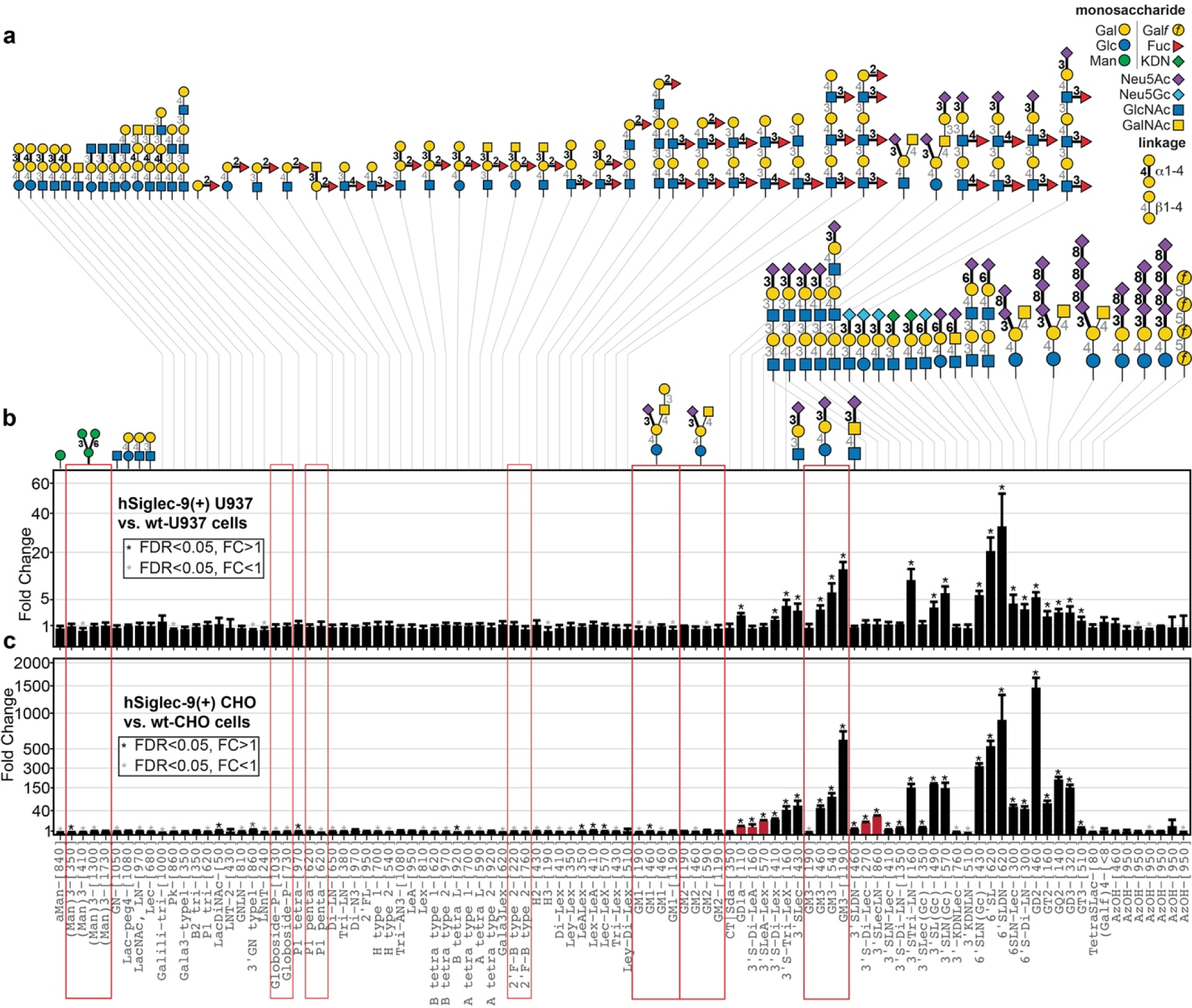
LiGA profiling of hSiglec-9 on the surface of U937 and CHO cells. **a**) Visual representation of the glycans in the LiGA. **b**) Siglec-9 expressing U937 cells with LiGA-100 enriched broad range of sialylated glycans. **c**) Siglec-9 expressing CHO cells showed a higher level of enrichment of the sialylated glycans and minor preference for a2-6 linked and poly-sialylated glycans. In **b-c**, the FC was calculated by Bioconductor edgeR DE analysis using the negative binomial model, TMM normalization, and BH correction for FDR (n= 5 for each cell type). Error bars represent s.d. propagated from the variance of the TMM-normalized sequencing data.

**Extended Data Figure 4:**
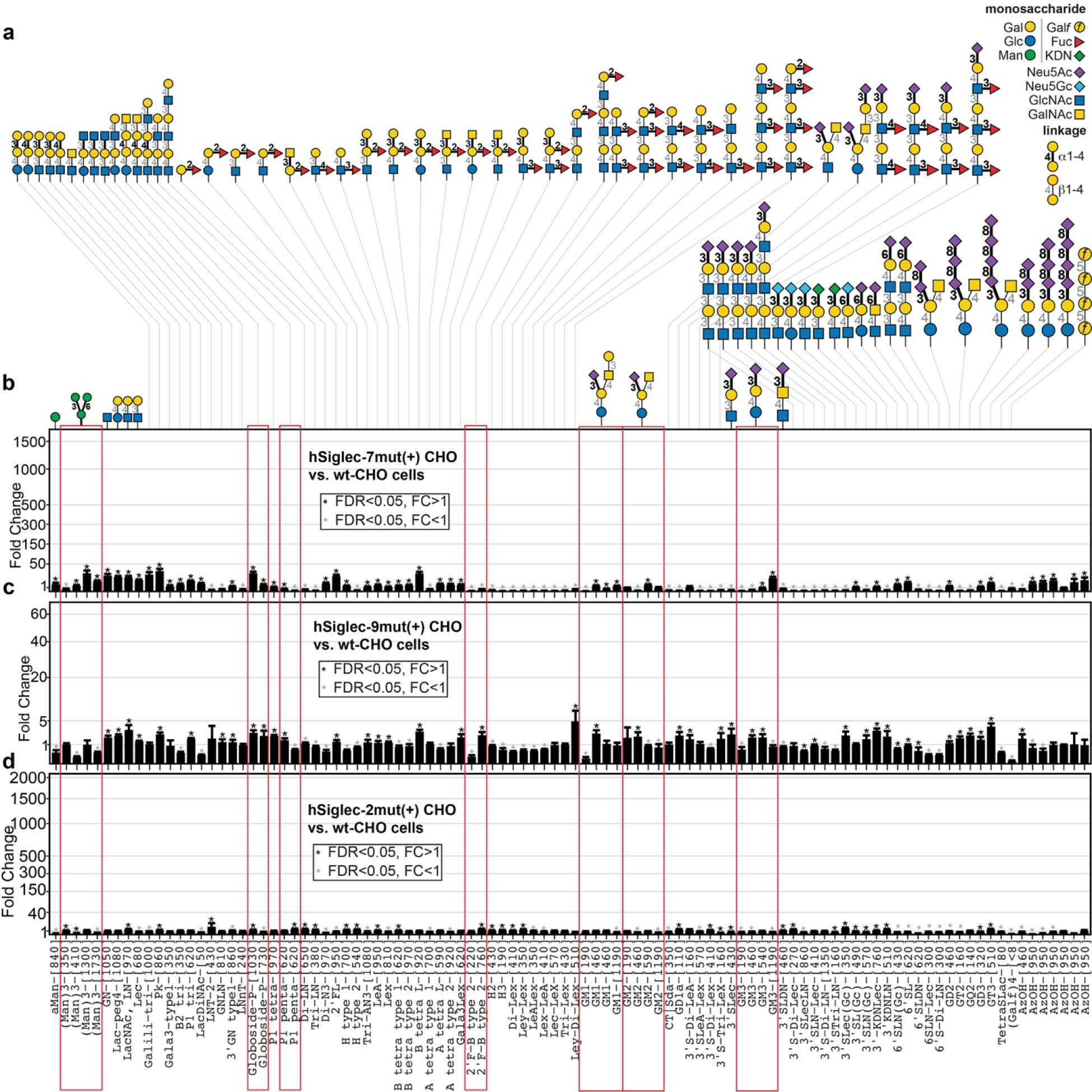
Arginine mutant of Siglec -7 -9 and hCD22. Data mirrored from Figure 3 for better visualization. **a)** Visual representation of glycans in the library. **b)** Siglec-7 R124A showed a lower enrichment compared to the wild-type Siglec on CHO cells binding. **c**) Siglec-9 R120A binding to LiGA. **d**) hCD22 R120A mutant showed a lack of binding. In **b-d**, the FC was calculated by Bioconductor edgeR DE analysis using the negative binomial model, TMM normalization, and BH correction for FDR (n= 5 for each cell type). Error bars represent s.d. propagated from the variance of the TMM-normalized sequencing data.

**Extended Data Figure 5:**
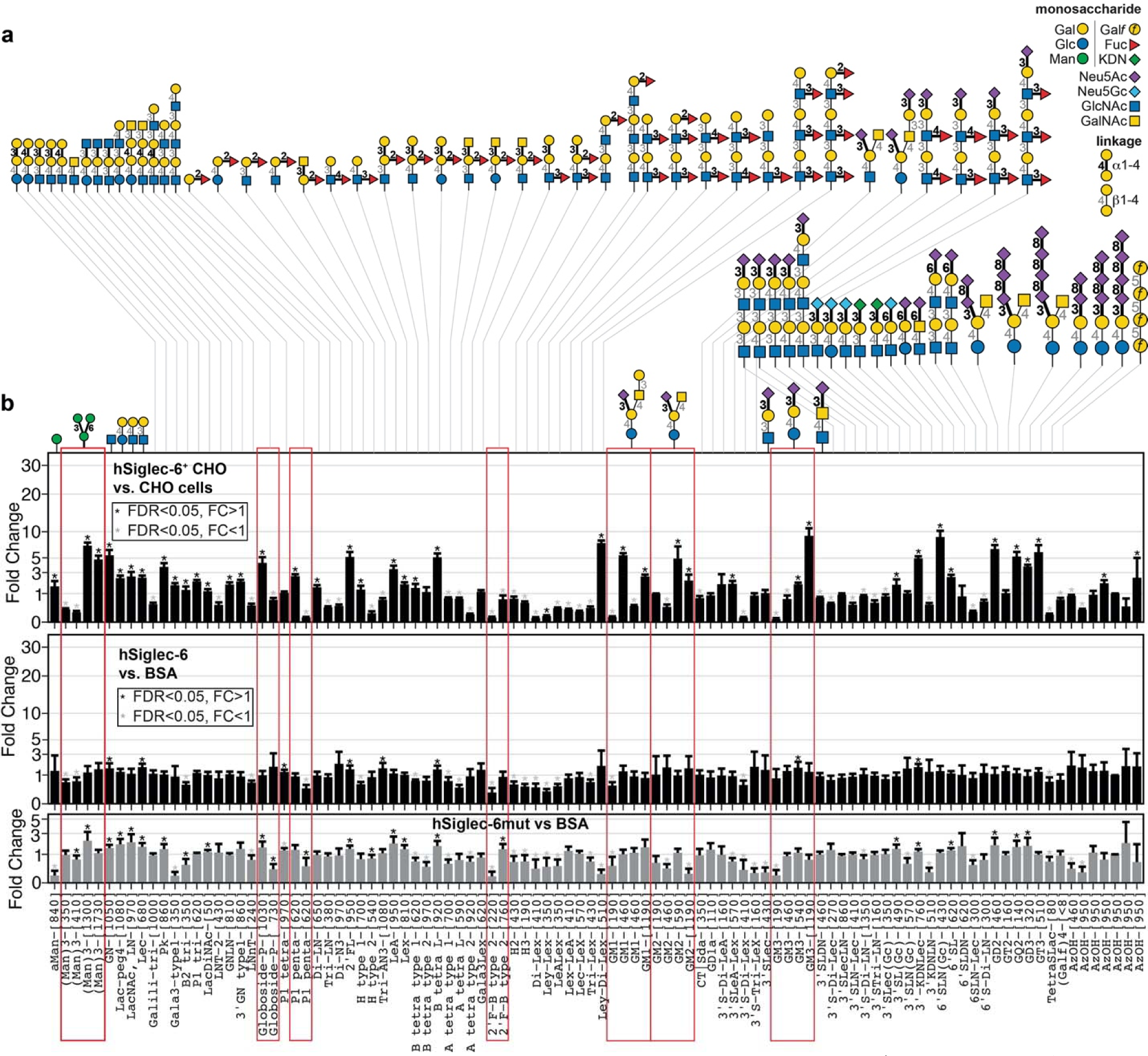
LiGA binding to purified Siglec-6 and Siglec-6+ CHO cells. **a**) Visual representation of the glycans in the LiGA. **b)** The binding pattern of LiGA on Siglec-6 expressing CHO cells showed no observable binding pattern. **c**) Binding of LiGA-100 purified Siglec 6 coated in a well. **d**) Purified Siglec-6 R122A mutant binding with LiGa-100. **e**) Differential enrichment of Siglec-6 compared to R122A mutant. The AzOH enrichment shows non-specific interaction among LiGA-100. In **b-e**, the FC was calculated by Bioconductor edgeR DE analysis using the negative binomial model, TMM normalization, and BH correction for FDR. Error bars represent s.d. propagated from the variance of the TMM-normalized sequencing data.

## Notes

### Competing Interest Statement

R.D. is a shareholders of the start-up company 48Hour Discovery Inc. that licensed the patent application (WO2018141058A1) describing LiGA technology. The remaining authors declare no competing interests.

