## Supplementary material for "Measuring carbohydrate recognition profile of lectins on live cells using liquid glycan array (LiGA)": File contains supplementary tables and figure

**This PDF includes:**

Supporting figures and table

**Supplementary Table 1:** Illumina sequencing primers. The combination of the F (forward) and R (reverse) primers are used for multiplexing of sequencing samples. The combination allows 380 samples to be analyzed in a single Illumina sequencing.

| | **Primer Sequences** |
|---|---|
| F1 | CAAGCAGAAGACGGCATACGAGATCGGTCTCGGCATTCCTGCTGAACCGCTCTTCCGATCTAAGCTTGGAGATTTTCAACGTG |
| F2 | CAAGCAGAAGACGGCATACGAGATCGGTCTCGGCATTCCTGCTGAACCGCTCTTCCGATCTACTGTTGGAGATTTTCAACGTG |
| F3 | CAAGCAGAAGACGGCATACGAGATCGGTCTCGGCATTCCTGCTGAACCGCTCTTCCGATCTAGAATTGGAGATTTTCAACGTG |
| F4 | CAAGCAGAAGACGGCATACGAGATCGGTCTCGGCATTCCTGCTGAACCGCTCTTCCGATCTTAATTTGGAGATTTTCAACGTG |
| F5 | CAAGCAGAAGACGGCATACGAGATCGGTCTCGGCATTCCTGCTGAACCGCTCTTCCGATCTTTCATTGGAGATTTTCAACGTG |
| F6 | CAAGCAGAAGACGGCATACGAGATCGGTCTCGGCATTCCTGCTGAACCGCTCTTCCGATCTTGGGTTGGAGATTTTCAACGTG |
| F8 | CAAGCAGAAGACGGCATACGAGATCGGTCTCGGCATTCCTGCTGAACCGCTCTTCCGATCTCTGTTTGGAGATTTTCAACGTG |
| F9 | CAAGCAGAAGACGGCATACGAGATCGGTCTCGGCATTCCTGCTGAACCGCTCTTCCGATCTCCACTTGGAGATTTTCAACGTG |
| F10 | CAAGCAGAAGACGGCATACGAGATCGGTCTCGGCATTCCTGCTGAACCGCTCTTCCGATCTGTAGTTGGAGATTTTCAACGTG |
| F11 | CAAGCAGAAGACGGCATACGAGATCGGTCTCGGCATTCCTGCTGAACCGCTCTTCCGATCTGCGATTGGAGATTTTCAACGTG |
| F12 | CAAGCAGAAGACGGCATACGAGATCGGTCTCGGCATTCCTGCTGAACCGCTCTTCCGATCTGGTTTTGGAGATTTTCAACGTG |
| F13 | CAAGCAGAAGACGGCATACGAGATCGGTCTCGGCATTCCTGCTGAACCGCTCTTCCGATCTAATATTGGAGATTTTCAACGTG |
| F14 | CAAGCAGAAGACGGCATACGAGATCGGTCTCGGCATTCCTGCTGAACCGCTCTTCCGATCTATTTTTGGAGATTTTCAACGTG |
| F15 | CAAGCAGAAGACGGCATACGAGATCGGTCTCGGCATTCCTGCTGAACCGCTCTTCCGATCTATGGTTGGAGATTTTCAACGTG |
| F16 | CAAGCAGAAGACGGCATACGAGATCGGTCTCGGCATTCCTGCTGAACCGCTCTTCCGATCTACCATTGGAGATTTTCAACGTG |
| F17 | CAAGCAGAAGACGGCATACGAGATCGGTCTCGGCATTCCTGCTGAACCGCTCTTCCGATCTACGTTTGGAGATTTTCAACGTG |
| F18 | CAAGCAGAAGACGGCATACGAGATCGGTCTCGGCATTCCTGCTGAACCGCTCTTCCGATCTAGTCTTGGAGATTTTCAACGTG |
| F19 | CAAGCAGAAGACGGCATACGAGATCGGTCTCGGCATTCCTGCTGAACCGCTCTTCCGATCTAGCTTTGGAGATTTTCAACGTG |
| F20 | CAAGCAGAAGACGGCATACGAGATCGGTCTCGGCATTCCTGCTGAACCGCTCTTCCGATCTTACCTTGGAGATTTTCAACGTG |
| R1 | AATGATACGGCGACCACCGAGATCTACACTCTTTCCCTACACGACGCTCTTCCGATCTAAGCACAGTTTCGGCCGA |
| R2 | AATGATACGGCGACCACCGAGATCTACACTCTTTCCCTACACGACGCTCTTCCGATCTACTGACAGTTTCGGCCGA |
| R3 | AATGATACGGCGACCACCGAGATCTACACTCTTTCCCTACACGACGCTCTTCCGATCTAGAAACAGTTTCGGCCGA |
| R4 | AATGATACGGCGACCACCGAGATCTACACTCTTTCCCTACACGACGCTCTTCCGATCTTAATACAGTTTCGGCCGA |
| R5 | AATGATACGGCGACCACCGAGATCTACACTCTTTCCCTACACGACGCTCTTCCGATCTTTCAACAGTTTCGGCCGA |
| R6 | AATGATACGGCGACCACCGAGATCTACACTCTTTCCCTACACGACGCTCTTCCGATCTTGGGACAGTTTCGGCCGA |
| R7 | AATGATACGGCGACCACCGAGATCTACACTCTTTCCCTACACGACGCTCTTCCGATCTCACGACAGTTTCGGCCGA |
| R8 | AATGATACGGCGACCACCGAGATCTACACTCTTTCCCTACACGACGCTCTTCCGATCTCTGTACAGTTTCGGCCGA |
| R9 | AATGATACGGCGACCACCGAGATCTACACTCTTTCCCTACACGACGCTCTTCCGATCTCCACACAGTTTCGGCCGA |
| R10 | AATGATACGGCGACCACCGAGATCTACACTCTTTCCCTACACGACGCTCTTCCGATCTGATGACAGTTTCGGCCGA |
| R11 | AATGATACGGCGACCACCGAGATCTACACTCTTTCCCTACACGACGCTCTTCCGATCTGCGAACAGTTTCGGCCGA |
| R12 | AATGATACGGCGACCACCGAGATCTACACTCTTTCCCTACACGACGCTCTTCCGATCTGGTTACAGTTTCGGCCGA |
| R13 | AATGATACGGCGACCACCGAGATCTACACTCTTTCCCTACACGACGCTCTTCCGATCTAATAACAGTTTCGGCCGA |
| R14 | AATGATACGGCGACCACCGAGATCTACACTCTTTCCCTACACGACGCTCTTCCGATCTATTTACAGTTTCGGCCGA |
| R15 | AATGATACGGCGACCACCGAGATCTACACTCTTTCCCTACACGACGCTCTTCCGATCTATGGACAGTTTCGGCCGA |
| R16 | AATGATACGGCGACCACCGAGATCTACACTCTTTCCCTACACGACGCTCTTCCGATCTACCAACAGTTTCGGCCGA |
| R17 | AATGATACGGCGACCACCGAGATCTACACTCTTTCCCTACACGACGCTCTTCCGATCTACGTACAGTTTCGGCCGA |
| R18 | AATGATACGGCGACCACCGAGATCTACACTCTTTCCCTACACGACGCTCTTCCGATCTAGTCACAGTTTCGGCCGA |
| R19 | AATGATACGGCGACCACCGAGATCTACACTCTTTCCCTACACGACGCTCTTCCGATCTAGCTACAGTTTCGGCCGA |
| R20 | AATGATACGGCGACCACCGAGATCTACACTCTTTCCCTACACGACGCTCTTCCGATCTTACCACAGTTTCGGCCGA |

**Supplementary Table 2:** Primer sequences used for first PCR in 2-step PCR protocol.

| | |
|---|---|
| NF10 | TTTTGGAGATTTTCAACGTG |
| -96 | CCCTCATAGTTAGCGTAACG |

**Supplementary Table 3:** Primer sequenced used for SDB-SVEK library.

| | |
|---|---|
| P1 | GAGATTTTCAACGTGAAAAAACTNCTNTTYGCNATHCCNCTNGTGGTACCTTTCTATTCTCA |
| P2 | TTAAGACTCCTTATTACGCAGTA |
| P3 | TTGCTAACATACTGCGTAATAAG |
| P4 | TTTTTTCACGTTGAAAATCTC |
| P5 | GTGGTACCTTTCTATTCTCACTCGAGYGTNGARAARAAYGAYCARAARACNTAYCAYGCNGGNGGNGGNTCG GCCGAAACTGTTGAAAG |

**Supplementary Table 4:** List of all the SDB used in this paper.

| SDB | Sequence |
|---|---|
| SDB99 | CTTCTCTTCGCGATACCGCTAAGTGTTGAGAAAAATGATCAAAAAACGTATCATGCGGGTGGTGGG |
| SDB126 | CTGCTCTTTGCAATTCCGCTGAGTGTAGAAAAAAATGATCAGAAAACATACCATGCTGGTGGCGGA |
| SDB89 | CTACTGTTCGCAATACCTCTCAGCGTGGAGAAAAATGATCAAAAAACATATCATGCAGGGGGTGGC |
| SDB125 | CTGCTATTCGCTATCCCGCTGAGTGTAGAAAAAAACGACCAGAAAACCTATCATGCCGGTGGCGGG |
| SDB87 | CTACTCTTCGCTATACCCCTCAGCGTAGAAAAAAATGACCAAAAGACCTACCATGCGGGAGGTGGG |
| SDB88 | CTGCTGTTCGCGATTCCTCTAAGTGTAGAGAAGAATGACCAGAAGACCTACCACGCGGGCGGGGGG |
| SDB54 | TTATTATTCGCAATTCCTTTAAGTGTCGAAAAAAACGATCAGAAGACCTATCACGCAGGCGGGGGC |
| SDB57 | TTATTATTCGCAATTCCTTTAAGCGTCGAGAAGAACGACCAAAAAACCTATCACGCAGGTGGCGGG |
| SDB52 | CTACTGTTTGCTATCCCGCTGAGTGTAGAGAAGAATGACCAAAAGACATATCATGCGGGAGGAGGT |
| SDB53 | CTGCTGTTTGCGATCCCGCTCAGTGTGGAGAAAAATGACCAAAAGACCTATCATGCTGGAGGGGGT |
| SDB48 | TTATTATTCGCAATTCCTTTAAGCGTAGAGAAAAACGACCAGAAGACCTATCACGCGGGAGGAGGT |
| SDB49 | CTACTGTTCGCTATCCCGCTGAGTGTGGAAAAGAATGATCAGAAGACTTACCACGCTGGTGGTGGG |
| NG04 | CTACTGTTCGCAATCCCGCTAAGTGTTGAGAAAAATGATCAAAAAACTTATCACGCCGGTGGTGGT |
| MC01 | CTTCTATTTGCTATTCCTCTAAGTGTCGAGAAGAACGACCAGAAGACGTATCATGCGGGGGGTGGT |
| SDB17 | CTACTCTTCGCGATTCCGCTTAGTGTGGAGAAGAATGATCAGAAGACTTATCATGCGGGTGGAGGT |
| SDB18 | CTGCTGTTTGCTATCCCTCTGAGTGTGGAGAAGAATGATCAGAAGACTTATCATGCGGGTGGAGGT |
| SDB15 | CTGCTGTTCGCCATACCCCTTAGTGTGGAGAAGAATGATCAGAAGACTTATCATGCGGGTGGAGGT |
| SDB16 | CTGCTGTTCGCAATCCCGCTGAGTGTGGAGAAGAATGATCAGAAGACTTATCATGCGGGTGGAGGT |
| SDB12 | CTTCTGTTCGCGATACCTCTAAGTGTGGAGAAGAATGATCAGAAGACTTATCATGCGGGTGGAGGT |
| SDB13 | CTACTTTTCGCAATTCCTCTGAGTGTGGAGAAGAATGATCAGAAGACTTATCATGCGGGTGGAGGT |
| SDB132 | CTACTGTTCGCAATACCTCTCAGCGTAGAAAAAAATGATCAAAAAACATATCACGCAGGTGGCGGA |
| SDB107 | TTATTATTCGCAATTCCTTTAAGTGTGGAGAAGAACGATCAGAAGACCTATCACGCAGGAGGCGGA |
| SDB129 | CTTCTGTTCGCGATACCGCTGAGCGTCGAAAAGAATGATCAGAAAACTTATCATGCGGGGGGCGGG |
| SDB130 | CTTCTCTTTGCGATACCGCTAAGTGTGGAAAAGAATGATCAAAAAACGTACCATGCCGGTGGGGGT |
| SDB127 | CTGCTATTTGCCATCCCACTAAGTGTTGAGAAGAACGATCAGAAAACGTATCATGCAGGTGGTGGA |
| SDB128 | TTATTATTCGCAATTCCTTTAAGCGTTGAAAAGAATGACCAGAAAACATATCACGCTGGGGGAGGG |
| SDB24 | CTACTATTCGCGATCCCGCTCAGTGTGGAGAAGAATGATCAGAAGACTTATCATGCGGGTGGAGGT |
| SDB25 | CTGCTTTTCGCAATACCTCTAAGTGTGGAGAAGAATGATCAGAAGACTTATCATGCGGGTGGAGGT |
| SDB22 | CTACTGTTTGCTATCCCACTTAGTGTGGAGAAGAATGATCAGAAGACTTATCATGCGGGTGGAGGT |
| SDB23 | CTGCTCTTTGCAATACCTCTTAGTGTGGAGAAGAATGATCAGAAGACTTATCATGCGGGTGGAGGT |
| SDB20 | CTGCTCTTTGCCATCCCGCTTAGTGTGGAGAAGAATGATCAGAAGACTTATCATGCGGGTGGAGGT |
| SDB21 | CTACTCTTTGCAATTCCCCTTAGTGTGGAGAAGAATGATCAGAAGACTTATCATGCGGGTGGAGGT |
| SDB70 | CTACTGTTCGCAATCCCGCTCAGTGTTGAAAAAAACGATCAAAAAACGTATCATGCTGGTGGAGGT |
| SDB71 | CTGCTTTTTGCTATTCCTCTGAGTGTTGAAAAAAACGATCAGAAGACTTATCACGCGGGGGGCGGG |
| SDB68 | CTACTATTCGCGATCCCCCTCAGCGTGGAGAAGAACGACCAGAAGACGTATCACGCAGGGGGGGGG |
| SDB69 | CTTCTGTTTGCGATTCCACTGAGCGTGGAGAAGAACGACCAGAAAACATACCATGCTGGTGGTGGA |
| SDB58 | CTACTCTTCGCTATACCCCTCAGCGTGGAGAAAAACGACCAAAAGACCTACCATGCAGGTGGTGGC |
| SDB67 | CTTCTGTTCGCTATACCTCTCAGCGTGGAGAAGAATGACCAGAAAACTTATCACGCAGGAGGTGGA |
| SDB84 | CTACTCTTCGCGATCCCACTGAGCGTCGAGAAGAACGATCAGAAGACTTATCACGCTGGCGGCGGC |
| SDB86 | CTTCTATTTGCTATTCCTCTAAGTGTGGAGAAGAATGATCAGAAGACATATCACGCTGGCGGGGGG |
| SDB82 | TTATTATTCGCAATTCCTTTAAGCGTGGAGAAGAACGACCAAAAGACTTATCACGCGGGCGGGGGG |
| SDB83 | CTGCTCTTCGCGATCCCTCTGAGTGTCGAAAAGAATGATCAAAAAACGTATCATGCGGGCGGTGGT |
| SDB80 | CTTCTGTTCGCGATCCCCCTAAGTGTAGAGAAGAATGATCAGAAAACGTATCATGCGGGAGGGGGT |
| SDB81 | CTACTCTTTGCTATTCCGCTTAGCGTGGAGAAGAATGATCAGAAAACGTACCACGCCGGTGGTGGC |
| SDB34 | CTTCTTTTTGCGATTCCGCTGAGTGTGGAGAAGAATGATCAGAAGACTTATCATGCGGGTGGAGGT |
| SDB35 | CTGCTCTTCGCTATTCCACTTAGTGTGGAGAAGAATGATCAGAAGACTTATCATGCGGGTGGAGGT |
| SDB28 | CTACTCTTCGCCATTCCACTGAGTGTGGAGAAGAATGATCAGAAGACTTATCATGCGGGTGGAGGT |
| SDB29 | CTGCTATTTGCGATCCCGCTGAGTGTGGAGAAGAATGATCAGAAGACTTATCATGCGGGTGGAGGT |
| SDB26 | CTGCTATTCGCTATCCCACTCAGTGTGGAGAAGAATGATCAGAAGACTTATCATGCGGGTGGAGGT |
| SDB27 | CTGCTTTTCGCTATTCCTCTCAGTGTGGAGAAGAATGATCAGAAGACTTATCATGCGGGTGGAGGT |
| SDB10 | CTACTGTTTGCTATACCGCTGAGTGTGGAGAAGAATGATCAGAAGACTTATCATGCGGGTGGAGGT |
| SDB11 | CTACTATTTGCGATTCCCCTGAGTGTGGAGAAGAATGATCAGAAGACTTATCATGCGGGTGGAGGT |
| SDB8 | CTGCTTTTTGCAATACCCCTCAGTGTGGAGAAGAATGATCAGAAGACTTATCATGCGGGTGGAGGT |
| SDB9 | CTTCTTTTTGCAATTCCTCTAAGTGTGGAGAAGAATGATCAGAAGACTTATCATGCGGGTGGAGGT |
| SDB2 | CTTCTATTCGCAATTCCGCTCAGTGTGGAGAAGAATGATCAGAAGACTTATCATGCGGGTGGAGGT |
| SDB6 | CTGCTGTTTGCGATTCCACTGAGTGTGGAGAAGAATGATCAGAAGACTTATCATGCGGGTGGAGGT |
| SDB135 | CTTCTGTTTGCTATACCTCTAAGTGTGGAGAAGAATGACCAAAAGACATATCACGCAGGTGGGGGA |
| SDB198 | CTGCTCTTTGCTATACCCCTAAGTGTGGAGAAAAATGACCAAAAAACTTATCACGCAGGAGGTGGT |
| SDB197 | CTGCTGTTTGCGATACCGCTTAGTGTTGAAAAAAATGACCAGAAAACCTACCACGCCGGTGGGGGT |

| | |
|---|---|
| SDB196 | CTGCTTTTTGCTATTCCTCTGAGTGTTGAGAAGAATGACCAGAAGACGTACCACGCGGGCGGGGGT |
| SDB199 | CTGCTCTTTGCTATACCCCTAAGCGTAGAAAAAAACGATCAGAAAACATATCATGCTGGAGGTGGT |
| SDB31 | CTGCTTTTCGCGATACCACTGAGTGTGGAGAAGAATGATCAGAAGACTTATCATGCGGGTGGAGGT |
| SDB78 | CTGCTGTTCGCGATTCCCCTAAGTGTCGAGAAAAATGATCAAAAAACTTATCATGCGGGAGGCGGG |
| SDB44 | CTGCTATTTGCGATTCCCCTAAGTGTAGAGAAAAACGATCAAAAGACGTATCATGCCGGGGGCGGT |
| SDB65 | CTGCTTTTTGCCATCCCTCTTAGTGTTGAAAAGAACGATCAAAAAACTTATCACGCAGGGGGGGGG |
| SDB73 | TTATTATTCGCAATTCCTTTAAGCGTTGAAAAAAACGATCAAAAGACATATCACGCTGGCGGAGGA |
| SDB75 | CTGCTCTTCGCGATCCCTCTAAGTGTGGAAAAAAATGATCAGAAAACGTACCATGCTGGAGGGGGT |
| SDB77 | CTGCTATTTGCCATCCCGCTGAGCGTCGAGAAAAACGATCAAAAGACATACCATGCCGGCGGAGGA |
| SDB131 | CTGCTGTTTGCGATTCCCCTGAGTGTCGAAAAGAATGACCAAAAGACGTACCATGCGGGGGGGGGG |
| SDB141 | CTTCTTTTTGCCATACCCCTGAGCGTAGAAAAAAACGACCAGAAAACATATCATGCTGGCGGCGGT |
| SDB220 | TTATTATTCGCAATTCCTTTAAGTGTGGAAAAAAATGATCAAAAGACATACCACGCGGGGGGCGGG |
| SDB201 | CTACTCTTTGCTATTCCACTAAGTGTGGAGAAAAATGATCAAAAGACGTACCATGCGGGTGGCGGT |
| SDB218 | CTGCTTTTTGCTATTCCTCTGAGTGTGGAAAAGAACGATCAGAAGACGTATCATGCCGGAGGAGGT |
| SDB96 | CTGCTGTTCGCAATCCCGCTAAGTGTTGAAAAAAATGATCAGAAAACATATCACGCTGGGGGTGGA |
| SDB62 | TTATTATTCGCAATTCCTTTAAGCGTAGAAAAGAACGATCAAAAAACGTATCACGCAGGTGGTGGA |
| SDB98 | CTGCTGTTTGCGATTCCTCTGAGTGTCGAAAAGAATGATCAAAAAACATACCATGCGGGGGGTGGG |
| SDB100 | CTACTGTTCGCAATCCCGCTCAGCGTCGAGAAGAATGACCAGAAGACGTACCATGCCGGGGGCGGG |
| SDB192 | CTACTATTTGCGATACCGCTAAGTGTGGAAAAAAACGATCAAAAAACATACCACGCTGGCGGCGGG |
| SDB181 | TTATTATTCGCAATTCCTTTAAGTGTGGAGAAAAACGACCAAAAGACCTACCATGCTGGGGGGGGT |
| SDB171 | CTTCTCTTTGCAATCCCGCTGAGCGTCGAGAAGAATGACCAGAAGACATACCACGCTGGTGGCGGC |
| SDB162 | TTATTATTCGCAATTCCTTTAAGTGTTGAGAAAAACGACCAGAAGACTTATCATGCTGGGGGGGGA |
| SDB97 | CTGCTGTTTGCAATCCCCCTAAGTGTCGAAAAGAACGATCAGAAAACATACCATGCGGGTGGGGGT |
| SDB92 | TTATTATTCGCAATTCCTTTAAGTGTGGAGAAGAATGACCAGAAAACGTACCACGCGGGGGGTGGG |
| SDB91 | CTGCTGTTTGCGATTCCCCTGAGCGTCGAGAAGAATGACCAAAAAACGTACCATGCGGGGGGTGGG |
| SDB39 | CTGCTGTTTGCGATCCCTCTAAGTGTCGAAAAGAACGACCAAAAAACTTACCTTGCAGGGGGGGGT |
| SDB161 | CTGCTATTCGCTATCCCCCTGAGTGTTGAGAAGAATGACCAAAAGACATATCACGCCGGGGGAGGG |
| SDB150 | CTGCTCTTTGCTATCCCTCTAAGTGTTGAAAAAAACGACCAGAAGACTTATCACGCAGGTGGTGGG |
| SDB117 | TTATTATTCGCAATTCCTTTAAGCGTTGAAAAGAATGACCAGAAAACGTACCACGCGGGGGGCGGA |
| SDB118 | CTACTCTTCGCGATACCCCTCAGTGTCGAAAAGAATGATCAGAAGACTTATCATGCAGGTGGTGGT |
| SDB111 | CTGCTGTTTGCTATCCCGCTAAGTGTAGAAAAAAATGACCAGAAAACGTACCATGCGGGCGGTGGG |
| SDB119 | CTGCTTTTTGCCATCCCTCTTAGTGTCGAGAAAAACGACCAGAAAACATACCACGCCGGAGGGGGT |
| SDB120 | CTGCTGTTTGCTATTCCACTAAGTGTTGAAAAAAACGATCAGAAGACTTATCATGCTGGGGGAGGA |
| SDB122 | CTGCTTTTTGCTATACCCCTGAGCGTGGAGAAGAACGATCAAAAGACCTATCACGCGGGGGGAGGC |
| SDB123 | NTGCTATTTGCCATCCCACTAAGCGTGGAAAAGAATGATCAGAAGACATATCATGCCGGCGGAGGT |
| SDB133 | CTACTCTTTGCTATTCCACTAAGCGTTGAAAAAAACGATCAGAAGACGTACCACGCTGGGGGCGGG |
| SDB136 | CTTCTTTTTGCGATTCCGCTCAGTGTAGAGAAGAACGATCAAAAGACGTATCATGCTGGAGGAGGT |
| SDB138 | CTACTATTCGCTATACCCCTAAGCGTGGAGAAAAATGACCAGAAGACGTATCACGCGGGGGGGGGG |
| SDB139 | CTGCTTTTTGCTATTCCTCTGAGCGTTGAGAAAAATGATCAAAAGACTTATCATGCTGGAGGGGGT |
| SDB140 | CTGCTGTTTGCGATCCCGCTTAGCGTAGAGAAGAACGACCAGAAGACGTACCACGCCGGCGGGGGG |
| SDB143 | CTGCTGTTCGCGATTCCCCTAAGTGTCGAAAAAAATGATCAAAAAACCTATCACGCAGGAGGCGGT |
| SDB147 | CTACTGTTCGCAATCCCGCTAAGCGTTGAGAAGAACGATCAAAAAACATACCACGCAGGTGGGGGA |
| SDB148 | CTTCTATTTGCTATCCCGCTAAGCGTTGAGAAAAATGACCAGAAAACGTACCATGCGGGCGGTGGG |
| SDB149 | CTACTCTTTGCAATCCCGCTTAGCGTGGAGAAGAACGACCAGAAGACTTACCACGCGGGTGGTGGG |
| SDB155 | TTATTATTCGCAATTCCTTTAAGTGTGGAGAAGAATGATCAAAAGACGTACCATGCAGGCGGAGGT |
| SDB156 | TTATTATTCGCAATTCCTTTAAGTGTCGAGAAAAACGATCAAAAGACCTACCACGCAGGGGGTGGT |
| SDB157 | CTGCTGTTTGCTATACCCCTTAGCGTGGAAAAGAATGACCAGAAGACCTACCATGCCGGTGGGGGT |
| SDB158 | CTACTTTTTGCTATTCCGCTGAGCGTGGAGAAGAATGATCAGAAGACATACCATGCGGGTGGAGGG |
| SDB159 | CTGCTTTTTGCTATTCCTCTGAGTGTCGAAAAGAACGATCAAAAGACTTATCATGCAGGGGGGGGG |
| SDB163 | CTGCTGTTTGCGATACCCCTGAGCGTGGAGAAAAATGACCAAAAGACTTACCATGCTGGAGGAGGA |
| SDB164 | CTGCTATTCGCGATCCCCCTGAGCGTTGAAAAAAATGATCAAAAAACTTATCATGCAGGCGGAGGG |
| SDB165 | CTGCTGTTTGCTATTCCACTCAGCGTGGAAAAAAATGACCAAAAGACCTACCACGCAGGCGGAGGG |
| SDB166 | CTGCTTTTTGCTATTCCTCTTAGCGTGGAAAAAAACGACCAAAAAACGTACCATGCAGGTGGCGGG |
| SDB170 | TTATTATTCGCAATTCCTTTAAGCGTGGAGAAAAATGACCAAAAAACGTATCATGCGGGGGGGGGT |
| SDB174 | CTACTGTTCGCTATTCCGCTTAGCGTGGAGAAAAATGATCAAAAAACCTACCACGCTGGGGGAGGT |
| SDB183 | CTACTATTTGCGATCCCCCTAAGTGTAGAGAAAAATGATCAGAAAACCTACCACGCCGGAGGCGGT |
| SDB61 | CTGCTCTTTGCTATTCCCCGGAGTGTTGAGAAGAATGATCAGAAGACGTATCATGNGGGGGGGGGG |
| SDB64 | CTGCTCTTTGCTATACCCCTAAGCGTCGAGAAGAACGACCAGAAGACCTACCACGCTGGAGGGGGA |
| SDB66 | TTATTATTCGCAATTCCTTTAAGTGTCGAGAAGAACGATCAGAAGACCTACCACGCAGGGGGGGGT |
| SDB74 | CTACTCTTTGCGATCCCTCTGAGTGTCGAAAAAAATGACCAAAAGACCTATCACGCGGGCGGGGGA |
| SDB76 | CTGCTTTTTGCCATACCGCTAAGTGTTGAGAAGAATGATCAGAAGACCTACCACGCAGGAGGTGGC |

| | |
|---|---|
| SDB79 | CTACTGTTCGCAATACCTCTCAGCGTGGAGAAAAACGACCAGAAAACGTATCATGCTGGTGGTGGA |
| SDB90 | CTTCTATTTGCGATTCCGCTGAGTGTTGAAAAAAACGATCAAAAAACTTACCACGCCGGTGGGGGC |
| SDB93 | TTATTATTCGCAATTCCTTTAAGTGTAGAAAAGAACGATCAAAAGACTTATCATGCGGGGGGGGGA |
| SDB94 | CTACTCTTTGCTATCCCTCTAAGTGTAGAAAAGAATGATCAAAAAACATATCATGCGGGTGGGGGT |
| SDB95 | CTTCTCTTTGCGATACCGCTAAGTGTAGAAAAGAACGACCAAAAGACCTATCACGCGGGTGGTGGG |
| SDB101 | CTACTTTTTGCTATTCCTCTCAGTGTTGAAAAAAACGATCAGAAGACTTACCATGCGGGTGGGGGT |
| SDB102 | CTACTCTTTGCTATCCCTCTAAGTGTCGAGAAGAATGATCAGAAAACATACCATGCCGGGGGGGGT |
| SDB103 | TTATTATTCGCAATTCCTTTAAGTGTTGAAAAAAACGACCAGAAGACTTATCACGCGGGTGGGGGG |
| SDB104 | CTGCTATTTGCCATCCCACTAAGTGTGGAGAAAAATGATCAGAAAACTTATCATGCCGGAGGTGGG |
| SDB105 | CTGCTTTTTGCTATTCCTCTTAGTGTCGAGAAGAATGATCAAAAAACTTACCACGCGGGTGGTGGA |
| SDB106 | CTACTGTTCGCAATCCCGCTAAGTGTAGAAAAAAACGATCAGAAGACTTACCATGCTGGTGGTGGG |
| SDB108 | CTTCTGTTCGCGATCCCCCTAAGTGTGGAGAAAAATGACCAAAAGACTTATCATGCTGGGGGAGGT |
| SDB109 | CTTCTCTTCGCGATCCCACTGAGTGTCGAGAAGAATGACCAGAAGACATACCACGCGGGCGGAGGC |
| SDB110 | CTGCTGTTTGCGATCCCGCTAAGCGTGGAAAAGAACGATCAGAAGACGTACCATGCGGGAGGGGGG |
| SDB113 | CTACTGTTTGCCATTCCCCTGAGTGTCGAAAAAAATGATCAGAAAACTTATCACGCAGGAGGGGGG |
| SDB115 | CTTCTGTTCGCAATTCCGCTGAGCGTAGAGAAGAATGATCAAAAGACGTATCATGCGGGTGGTGGA |
| SDB116 | CTGCTTTTTGCTATACCTCTTAGCGTGGAGAAGAACGATCAGAAAACGTATCATGCTGGTGGTGGG |
| SDB1 | CTGCTGTTCGCAATACCACTCAGTGTGGAGAAGAATGATCAGAAGACTTATCATGCGGGTGGAGGT |
| SDB3 | CTGCTTTTCGCAATTCCGCTTAGTGTGGAGAAGAATGATCAGAAGACTTATCATGCGGGTGGAGGT |
| SDB36 | CTGCTCTTCGCGATCCCTCTGAGTGTGGAGAAGAATGATCAGAAGACTTATCATGCGGGTGGAGGT |
| SDB37 | CTTCTTTTTGCCATCCCGCTCAGTGTGGAGAAGAATGATCAGAAGACTTATCATGCGGGTGGAGGT |
| SDB38 | CTGCTGTTTGCGATACCGCTAAGTGTGGAGAAGAATGATCAGAAGACTTATCATGCGGGTGGAGGT |
| SDB40 | CTTCTGTTCGCTATTCCGCTTAGCGTGGAAAAGAACGATCAAAAGACCTATCACGCCGGGGGAGGG |
| SDB41 | CTTCTCTTTGCGATTCCCCTGAGTGTAGAAAAAAATGATCAGAAGACATATCATGCGGGAGGTGGG |
| SDB42 | CTTCTGTTTGCAATACCCCTGAGCGTGGAGAAGAATGATCAAAAGACGTACCATGCGGGGGGAGGG |
| SDB43 | CTTCTGTTTGCTATTCCGCTCAGTGTGGAGAAGAATGATCAAAAAACCTACCATGCCGGGGGAGGG |
| SDB45 | CTGCTGTTTGCGATTCCTCTGAGCGTGGAAAAAAATGACCAAAAAACCTACCATGCAGGGGGGGGA |
| SDB46 | CTGCTATTTGCCATCCCGCTGAGCGTTGAAAAAAACGACCAGAAAACGTATCACGCGGGGGGGGGG |
| SDB50 | CTGCTCTTTGCGATTCCGCTGAGTGTCGAGAAGAACGACCAGAAAACATATCACGCGGGGGGGGGA |
| SDB51 | CTGCTCTTTGCTATACCCCTAAGCGTCGAGAAAAATGATCAAAAAACCTATCACGCCGGGGGGGGA |
| SDB55 | CTGCTCTTTGCAATTCCGCTGAGTGTCGAAAAAAATGACCAAAAGACTTATCACGCTGGTGGTGGA |
| SDB60 | CTACTGTTCGCCATCCCTCTGAGTGTAGAAAAGAACGACCAAAAGACTTACCATGCTGGTGGGGGG |
| SDB184 | CTGCTATTTGCCATCCCGCTGAGCGTGGAGAAGAACGATCAAAAGACGTACCATGCTGGAGGTGGG |
| SDB185 | CTTCTTTTTGCAATTCCACTCAGCGTGGAGAAAAACGATCAGAAAACATATCATGCGGGTGGAGGG |
| SDB187 | CTACTATTTGCGATTCCGCTAAGCGTAGAGAAAAATGACCAGAAGACTTACCACGCAGGAGGTGGG |
| SDB188 | CTACTGTTCGCCATCCCGCTTAGCGTGGAAAAAAACGATCAGAAGACATACCATGCGGGGGGAGGG |
| SDB189 | TTATTATTCGCAATTCCTTTAAGCGTGGAAAAGAATGATCAAAAGACGTACCATGCCGGCGGGGGC |
| SDB190 | CTGCTGTTCGCTATTCCACTAAGTGTAGAAAAGAATGATCAGAAGACTTATCATGCAGGTGGGGGG |
| SDB191 | TTATTATTCGCAATTCCTTTAAGCGTAGAGAAGAATGATCAAAAAACATATCACGCTGGTGGTGGT |
| SDB193 | CTGCTGTTCGCTATCCCGCTAAGTGTTGAGAAAAATGATCAAAAGACTTACCATGCAGGAGGTGGA |
| SDB194 | CTTCTCTTTGCGATACCGCTCAGTGTTGAAAAAAACGATCAAAAGACCTACCACGCGGGTGGTGGC |
| SDB195 | CTTCTTTTCGCCATACCGCTAAGCGTTGAAAAAAACGATCAAAAAACTTATCATGCTGGAGGGGGT |
| SDB200 | CTTCTCTTTGCTATTCCTCTTAGTGTCGAGAAAAACGACCAAAAGACCTACCATGCGGGGGGAGGG |
| SDB202 | CTGCTCTTTGCTATTCCCCTGAGCGTGGAGAAGAATGACCAGAAAACGTACCATGCAGGAGGCGGG |
| SDB203 | CTACTCTTCGCTATACCCCTCAGTGTTGAGAAAAACGATCAAAAGACCTATCATGCCGGAGGGGGT |
| SDB204 | CTGCTGTTTGCGATACCGCTTAGTGTTGAAAAAAATGATCAAAAAACCTATCACGCCGGTGGAGGG |
| SDB205 | CTTCTTTTTGCTATCCCACTGAGCGTCGAGAAGAATGACCAAAAGACTTACCACGCGGGAGGTGGA |
| SDB206 | CTTCTATTTGCTATTCCTCTAAGTGTTGAAAAAAACGACCAAAAAACATATCATGCGGGGGGCGGA |
| SDB207 | CTACTCTTCGCGATCCCACTGAGTGTTGAAAAAAATGATCAAAAAACCTACCATGCTGGCGGGGGT |
| SDB208 | TTATTATTCGCAATTCCTTTAAGTGTTGAGAAAAATGACCAGAAAACTTACCATGCGGGAGGAGGA |
| SDB209 | CTGCTATTTGCGATCCCCCTTAGTGTGGAAAAAAATGATCAGAAGACGTATCACGCGGGAGGAGGA |
| SDB210 | CTACTGTTCGCTATTCCGCTTAGTGTTGAAAAGAATGACCAGAAAACGTATCATGCCGGTGGCGGA |
| SDB211 | CTACTGTTTGCTATCCCGCTTAGCGTTGAGAAGAATGATCAAAAGACGTATCATGCGGGCGGCGGA |
| SDB212 | CTACTGTTCGCGATCCCACTGAGTGTTGAAAAGAATGACCAGAAAACATATCACGCGGGTGGGGGT |
| SDB213 | CTTCTCTTCGCCATACCGCTCAGTGTTGAAAAAAATGATCAGAAGACTTATCATGCTGGTGGAGGC |
| SDB214 | CTGCTGTTTGCGATACCGCTTAGCGTAGAAAAAAATGATCAAAAAACTTACCACGCCGGTGGCGGG |
| SDB215 | CTTCTTTTCGCCATTCCACTAAGTGTTGAGAAAAACGACCAAAAAACGTACCATGCCGGGGGTGGG |
| SDB219 | CTTCTCTTTGCAATACCCCTTAGTGTAGAAAAAAATGACCAGAAAACATACCATGCCGGTGGAGGT |
| SDB221 | TTATTATTCGCAATTCCTTTAAGCGTGGAGAAAAATGACCAAAAAACGTACCACGCTGGGGGGGGG |
| SDB222 | CTGCTATTTGCTATCCCGCTGAGTGTTGAGAAAAACGATCAAAAAACGTACCACGCTGGAGGTGGT |
| SDB223 | CTACTCTTTGCTATTCCACTAAGCGTTGAAAAAAATGACCAGAAAACTTATCATGCCGGAGGGGGA |

```
SDB224  TTATTATTCGCAATTCCTTTAAGTGTGGAGAAGAACGATCAGAAGACCTATCACGCCGGGGGTGGA
SDB228  CTGCTGTTTGCGATTCCCCTAAGTGTAGAGAAAAACGATCAAAAAACATATCATGCAGGAGGCGGA
SDB226  CTGCTGTTTGCGATCCCTCTAAGTGTCGAGAAGAACGATCAGAAGACTTATCATGCCGGAGGGGGT
SDB217  TTATTATTCGCAATTCCTTTAAGTGTGGAAAAGAACGATCAAAAAACTTATCACGCGGGTGGGGGC
SDB137  CTGCTGTTTGCGATCCCTCTAAGTGTGGAAAAAAACGATCAGAAAACCTATCACGCTGGCGGGGGT
SDB121  CTGCTGTTTGCGATACCGCTTAGTGTGGAGAAAAACGATCAGAAGACCTATCATGCTGGGGGTGGA
SDB167  CTGCTTTTTGCTATTCCTCTGAGTGTCGAAAAAAACGATCAGAAGACGTATCACGCAGGTGGTGGT
SDB146  CTGCTGTTTGCTATACCCCTTAGTGTCGAGAAAAATGACCAAAAGACGTACCATGCGGGAGGAGGG
SDB145  CTGCTTTTTGCCATTCCACTTAGTGTTGAGAAGAACGACCAGAAAACGTACCACGCGGGGGGTGGG
SDB134  CTGCTTTTTGCTATACCCCTGAGTGTAGAGAAGAATGACCAAAAAACTTACCACGCTGGTGGGGGC
SDB30   CTGCTTTTTGCGATTCCCCTTAGTGTGGAGAAGAATGATCAGAAGACTTATCATGCGGGTGGAGGT
SDB153  CTACTATTCGCTATTCCTCTGAGTGTCGAGAAAAACGATCAAAAGACCTATCATGCGGGTGGGGGT
SDB33   CTGCTTTTTGCCATACCGCTGAGTGTGGAGAAGAATGATCAGAAGACTTATCATGCGGGTGGAGGT
SDB59   TTATTATTCGCAATTCCTTTAAGTGTCGAAAAGAATGACCAAAAAACTTATCATGCAGGTGGGGGA
SDB154  CTACTGTTCGCTATTCCGCTAAGTGTAGAAAAAAACGATCAAAAGACGTATCACGCCGGTGGGGGA
SDB169  CTGCTGTTCGCGATTCCCCTAAGCGTCGAAAAGAATGACCAAAAGACGTATCACGCTGGCGGGGGG
SDB142  CTGCTTTTTGCTATACCCCTGAGCGTTGAGAAGAATGACCAGAAGACTTATCACGCAGGTGGAGGG
SDB72   TTATTATTCGCAATTCCTCTAAGCGTCGAGAAGAACGATCAAAAAACTTATCACGCGGGAGGTGGA
SDB144  CTTCTTTTTGCCATACCCCTGAGCGTAGAGAAGAATGATCAAAAGACTTATCATGCTGGGGGAGGA
SDB151  CTGCTGTTTGCGATCCCGCTTAGCGTAGAAAAGAATGACCAGAAGACATACCACGCGGGAGGGGGG
SDB160  TTATTATTCGCAATTCCTTTAAGTGTCGAAAAAAATGACCAGAAAACTTACCATGCTGGAGGGGGC
SDB56   CTACTGTTTGCTATACCTCTTAGCGTTGAAAAAAATGACCAGAAAACTTACCATGCGGGTGGAGGT
SDB168  TTATTATTCGCAATTCCTTTAAGCGTGGAAAAAAATGATCAAAAGACATATCACGCTGGTGGTGGG
SDB152  CTGCTTTTTGCTATACCTCTCAGCGTAGAAAAGAATGACCAGAAGACGTACCATGCCGGTGGGGGG
SDB114  CTACTGTTTGCGATCCCACTCAGCGTGGAGAAGAACGACCAGAAGACCTACCACGCGGGGGGAGGG
SDB186  CTACTGTTCGCAATCCCGCTAAGTGTTGAGAAAAACGATCAAAAGACTTATCATGCAGGCGGAGGA
SDB112  CTACTGTTCGCGATCCCACTGAGTGTAGAAAAAAATGACCAGAAGACGTATCACGCAGGCGGAGGC
SDB85   TTATTATTCGCAATTCCTTTAAGTGTGGAGAAGAACGACCAGAAAACGTATCACGCAGGTGGGGGT
```

### **Supplementary Table 5:** LiGA-YY

| **SDB** | **Modification** |
|---|---|
| SDB99 | Manb-S0 |
| SDB126 | Manb-S0 |
| SDB89 | Manb-S0 |
| SDB125 | Manb-S0 |
| SDB87 | Manb-S0 |
| SDB88 | Manb-S0 |
| SDB54 | Mana-S6 |
| SDB57 | Mana-S6 |
| SDB52 | Mana-S6 |
| SDB53 | Mana-S6 |
| SDB48 | Mana-S6 |
| SDB49 | Mana-S6 |
| **NG04** | Mana-S6 |
| SDB17 | Mana1-6[Mana1-3]Mana-S6 |
| SDB18 | Mana1-6[Mana1-3]Mana-S6 |
| SDB15 | Mana1-6[Mana1-3]Mana-S6 |
| SDB16 | Mana1-6[Mana1-3]Mana-S6 |
| SDB12 | Mana1-6[Mana1-3]Mana-S6 |
| SDB13 | Mana1-6[Mana1-3]Mana-S6 |
| SDB132 | Galb-P4 |
| SDB107 | Galb-P4 |
| SDB129 | Galb-P4 |
| SDB130 | Galb-P4 |
| SDB127 | Galb-P4 |
| SDB128 | Galb-P4 |
| SDB24 | Galb1-4Glcb-Sp |
| SDB25 | Galb1-4Glcb-Sp |
| SDB22 | Galb1-4Glcb-Sp |
| SDB23 | Galb1-4Glcb-Sp |
| SDB20 | Galb1-4Glcb-Sp |
| SDB21 | Galb1-4Glcb-Sp |
| MC01 | Galb1-4Glcb-Sp |
| SDB70 | Galb1-3GlcNAcb1-3Galb1-4GlcNAcb-Sp |
| SDB71 | Galb1-3GlcNAcb1-3Galb1-4GlcNAcb-Sp |
| SDB68 | Galb1-3GlcNAcb1-3Galb1-4GlcNAcb-Sp |
| SDB69 | Galb1-3GlcNAcb1-3Galb1-4GlcNAcb-Sp |
| SDB58 | Galb1-3GlcNAcb1-3Galb1-4GlcNAcb-Sp |
| SDB67 | Galb1-3GlcNAcb1-3Galb1-4GlcNAcb-Sp |
| SDB84 | Galfb-S8 |
| SDB86 | Galfb-S8 |
| SDB82 | Galfb-S8 |
| SDB83 | Galfb-S8 |
| SDB80 | Galfb-S8 |
| SDB81 | Galfb-S8 |
| SDB34 | Galfb1-5Galfb1-5Galfb-S8 |
| SDB35 | Galfb1-5Galfb1-5Galfb-S8 |
| SDB28 | Galfb1-5Galfb1-5Galfb-S8 |
| SDB29 | Galfb1-5Galfb1-5Galfb-S8 |
| SDB26 | Galfb1-5Galfb1-5Galfb-S8 |
| SDB27 | Galfb1-5Galfb1-5Galfb-S8 |
| SDB10 | Galfb1-5Galfb1-5Galfb1-5Galfb-S8 |
| SDB11 | Galfb1-5Galfb1-5Galfb1-5Galfb-S8 |
| SDB8 | Galfb1-5Galfb1-5Galfb1-5Galfb-S8 |
| SDB9 | Galfb1-5Galfb1-5Galfb1-5Galfb-S8 |

| | |
|---|---|
| SDB2 | Galfb1-5Galfb1-5Galfb1-5Galfb-S8 |
| SDB6 | Galfb1-5Galfb1-5Galfb1-5Galfb-S8 |

**Supplementary Table 6:** LiGA-100 (EF.xlsx)

| SDB | Modification | Axis Name |
|---|---|---|
| SDB123 | Galb1-4Glcb-P4 | Lac-peg4-[1080] |
| SDB217 | Mana-S6 | aMan-[840] |
| SDB20 | Mana1-6[Mana1-3]Mana-S6 | (Man)3-[350] |
| SDB137 | Mana1-6[Mana1-3]Mana-S6 | (Man)3-[410] |
| SDB121 | Mana1-6[Mana1-3]Mana-S6 | (Man)3-[1300] |
| SDB122 | Mana1-6[Mana1-3]Mana-S6 | (Man)3-[1730] |
| SDB167 | Fuca1-2Galb1-4Glcb-Sp | 2'FL-[950] |
| SDB146 | Fuca1-2Galb1-3GlcNAcb-Sp | H type 1-[700] |
| SDB35 | Fuca1-2(Galb1-4GlcNAcb1-3)2b-Sp | H2-[430] |
| SDB145 | Fuca1-2(Galb1-4GlcNAcb1-3)3b-Sp | H3-[190] |
| SDB64 | Fuca1-2Galb1-4GlcNAcb-Sp | H-type 2-[540] |
| SDB34 | GalNAca1-3[Fuca1-2]Galb1-3GlcNAcb-Sp | A tetra type 1-[700] |
| SDB134 | GalNAca1-3[Fuca1-2]Galb1-4GlcNAcb-Sp | A tetra type 2-[920] |
| SDB15 | GalNAca1-3[Fuca1-2]Galb1-4Glcb-Sp | A tetra L-[590] |
| SDB30 | Gala1-3[Fuca1-2]Galb1-4Glcb-Sp | B tetra L-[920] |
| SDB153 | GlcNAcb1-3Galb1-4GlcNAcb-Sp | GNLN-[810] |
| SDB33 | GalNAcb1-3Gala1-4Galb1-4Glcb-Sp | Globoside-P-[1030] |
| SDB59 | GalNAcb1-3Gala1-4Galb1-4Glcb-Sp | Globoside-P-[730] |
| SDB154 | GlcNAcb-Sp | GN-[1050] |
| SDB89 | GalNAcb1-3Gala1-4Galb1-4GlcNAcb-Sp | P1 tetra-[970] |
| SDB53 | Gala1-4Galb1-4GlcNAcb1-3Galb1-4Glcb-Sp | P1 penta-[620] |
| SDB23 | Galb1-3GalNAcb1-3Gala1-4Galb1-4GlcNAcb-Sp | P1 penta-[620] |
| SDB58 | GalNAcb1-4GlcNAcb-Sp | LacDiNAc-[50] |
| SDB169 | GlcNAcb1-3Galb1-3GlcNAcb-Sp | 3'GN type1-[860] |
| SDB8 | Galb1-4GlcNAcb1-3Galb1-4Glcb-Sp | LNnT-[240] |
| SDB142 | Gala1-3[Fuca1-2]Galb1-3GlcNAcb-Sp | B tetra type 1-[620] |
| SDB72 | Gala1-3[Fuca1-2]Galb1-4[Fuca1-3]GlcNAcb-Sp | 2'F-B type 2-[220] |
| SDB60 | Gala1-3[Fuca1-2]Galb1-4[Fuca1-3]GlcNAcb-Sp | 2'F-B type 2-[760] |
| SDB144 | Gala1-3[Fuca1-2]Galb1-4GlcNAcb-Sp | B tetra type 2-[970] |
| SDB151 | Galb1-4[Fuca1-3]GlcNAcb-Sp | Lex-[810] |
| SDB111 | (Galb1-4[Fuca1-3]GlcNAcb1-3)2b-Sp | Di-Lex-[410] |
| SDB69 | Galb1-3GlcNAcb1-3Galb1-4[Fuca1-3]GlcNAcb-Sp | Lec-LeX-[570] |
| SDB130 | Galb1-3[Fuca1-4]GlcNAcb1-3Galb1-4[Fuca1-3]GlcNAcb-Sp | LeALex-[350] |
| SDB84 | Galb1-4[Fuca1-3]GlcNAcb1-3Galb1-3[Fuca1-4]GlcNAcb-Sp | Lex-LeA-[410] |
| SDB115 | GlcNAcb1-3Galb1-4Glcb-Sp | LNT-2-[430] |
| SDB26 | Fuca1-2Galb1-4[Fuca1-3]GlcNAcb1-3Galb1-4[Fuca1-3]GlcNAcb-Sp | Ley-Lex-[350] |
| SDB160 | Fuca1-2Galb1-4[Fuca1-3]GlcNAcb1-3(Galb1-4[Fuca1-3]GlcNAcb1-3)2b-Sp | Ley-Di-Lex-[510] |
| SDB56 | Gala1-3Galb1-4[Fuca1-3]GlcNAcb-Sp | Gala3Lex-[620] |
| SDB117 | KDNa2-3Galb1-4GlcNAcb-Sp | 3'KDNLN-[510] |
| SDB168 | Gala1-3Galb1-3GlcNAcb-Sp | Gala3-type1-[350] |
| SDB152 | Gala1-4Galb1-4Glcb-Sp | Pk-[860] |
| SDB55 | Gala1-4Galb1-4GlcNAcb-Sp | P1 tri-[620] |
| SDB143 | Gala1-3Galb1-4GlcNAcb-Sp | B2 tri-[350] |
| SDB113 | Gala1-3Galb1-4Glcb-Sp | Galili-tri-[1000] |
| SDB125 | Fuca1-2Galb-Sp | Di-N3-[970] |
| SDB126 | GalNAca1-3[Fuca1-2]Galb-Sp | Tri-AN3-[1080] |
| SDB52 | Galb1-3GlcNAcb-Sp | Lec-[680] |
| SDB79 | Galb1-4GlcNAcb-Sp | LacNAc, LN-[970] |
| SDB114 | Galb1-3[Fuca1-4]GlcNAcb-Sp | LeA-[950] |
| SDB135 | (Galb1-4[Fuca1-3]GlcNAcb1-3)3b-Sp | Tri-Lex-[430] |
| SDB186 | (Galb1-4GlcNAcb1-3)2b-Sp | Di-LN-[650] |
| SDB112 | (Galb1-4GlcNAcb1-3)3b-Sp | Tri-LN-[380] |
| SDB73 | Neu5Aca2-3Galb1-3GlcNAcb-Sp | 3'SLec-[430] |
| SDB220 | Neu5Aca2-3GalNAcb1-4GlcNAcb-Sp | 3'SLDN-[460] |
| SDB181 | Neu5Aca2-3Galb1-4Glcb-Sp | GM3-[190] |
| SDB171 | Neu5Aca2-3Galb1-4Glcb-Sp | GM3-[460] |
| SDB192 | Neu5Aca2-3Galb1-4Glcb-Sp | GM3-[1190] |
| SDB98 | Neu5Aca2-3[GalNAcb1-4]Galb1-4Glcb-Sp | GM2-[590] |
| SDB97 | Neu5Aca2-3[GalNAcb1-4]Galb1-4Glcb-Sp | GM2-[190] |
| SDB92 | Neu5Aca2-3[GalNAcb1-4]Galb1-4Glcb-Sp | GM2-[460] |
| SDB162 | Neu5Aca2-3[GalNAcb1-4]Galb1-4Glcb-Sp | GM2-[1190] |
| SDB39 | Neu5Aca2-3[Galb1-3GalNAcb1-4]Galb1-4Glcb-Sp | GM1-[190] |
| SDB99 | Neu5Aca2-3[Galb1-3GalNAcb1-4]Galb1-4Glcb-Sp | GM1-[460] |
| SDB10 | Neu5Aca2-3[Galb1-3GalNAcb1-4]Galb1-4Glcb-Sp | GM1-[460] |
| SDB91 | Neu5Aca2-3[Galb1-3GalNAcb1-4]Galb1-4Glcb-Sp | GM1-[1190] |

| | | |
|---|---|---|
| SDB65 | Neu5Aca2-3[GalNAcb1-4]Galb1-4GlcNAcb-Sp | CT\|Sda-[350] |
| SDB31 | Neu5Aca2-3[Neu5Aca2-3Galb1-3GalNAcb1-4]Galb1-4Glcb-Sp | GD1a-[110] |
| SDB96 | Neu5Aca2-3Galb1-3[Fuca1-4]GlcNAcb1-3Galb1-4[Fuca1-3]GlcNAcb-Sp | 3'SLeA-Lex-[570] |
| SDB88 | Neu5Aca2-3Galb1-3GlcNAcb1-3Galb1-4GlcNAcb-Sp | 3'SLecLN-[860] |
| SDB80 | Neu5Aca2-3Galb1-4GlcNAcb1-3Galb1-3GlcNAcb-Sp | 3'SLN-Lec-[410] |
| SDB118 | Neu5Gca2-3Galb1-3GlcNAcb-Sp | 3'SLec (Gc)-[350] |
| SDB198 | Neu5Aca2-3(Galb1-4GlcNAcb1-3)2b-Sp | 3'S-Di-LN-[1350] |
| SDB196 | Neu5Aca2-3(Galb1-4GlcNAcb1-3)3b-Sp | 3'STri-LN-[160] |
| SDB218 | Neu5Aca2-3(Galb1-4[Fuca1-3]GlcNAcb1-3)3b-Sp | 3'S-Tri-LeX-[160] |
| SDB132 | Neu5Aca2-3Galb1-4Glcb-Sp | GM3-[540] |
| SDB161 | Neu5Gca2-6Galb1-4GlcNAcb-Sp | 6'SLN (Gc)-[430] |
| SDB131 | Neu5Aca2-6GalNAcb1-4GlcNAcb-Sp | 6'SLDN-[620] |
| SDB75 | Neu5Aca2-6Galb1-4Glcb-Sp | 6'SL-[620] |
| SDB199 | Neu5Aca2-6Galb1-4GlcNAcb1-3Galb1-3GlcNAcb-Sp | 6SLN-Lec-[300] |
| SDB197 | Neu5Aca2-6(Galb1-4GlcNAcb1-3)2b-Sp | 6'S-Di-LN-[300] |
| SDB141 | Neu5Gca2-3Galb1-4GlcNAcb-Sp | 3'SLN (Gc)-[570] |
| SDB77 | Neu5Gca2-3Galb1-4Glcb-Sp | 3'SL (Gc)-[490] |
| SDB68 | Neu5Aca2-8Neu5Aca2-3Galb1-4Glcb-Sp | GD3-[320] |
| SDB201 | Neu5Aca2-8Neu5Aca2-8Neu5Aca2-3Galb1-4Glcb-Sp | GT3-[510] |
| SDB71 | Neu5Aca2-8Neu5Aca2-8Neu5Aca2-3[GalNAcb1-4]Galb1-4Glcb-Sp | GT2-[160] |
| SDB44 | Neu5Aca2-8Neu5Aca2-8Neu5Aca2-8Neu5Aca2-3[GalNAcb1-4]Galb1-4Glcb-Sp | GQ2-[140] |
| SDB78 | Neu5Aca2-8Neu5Aca2-8Neu5Aca2-8Neu5Aca2-3Galb1-4Glcb-Sp | TetraSLac-[80] |
| SDB100 | Neu5Aca2-8Neu5Aca2-3[GalNAcb1-4]Galb1-4Glcb-Sp | GD2-[460] |
| SDB150 | KDNa2-3Galb1-3GlcNAcb-Sp | 3'-KDNLec-[760] |
| SDB83 | Neu5Aca2-3Galb1-3GlcNAcb1-3Galb1-3GlcNAcb-Sp | 3'S-Di-Lec-[270] |
| SDB62 | Neu5Aca2-3(Galb1-3[Fuca1-4]GlcNAcb1-3)2b-Sp | 3'S-Di-LeA-[160] |
| SDB13 | Neu5Aca2-3(Galb1-4[Fuca1-3]GlcNAcb1-3)2b-Sp | 3'S-Di-Lex-[410] |
| SDB1 | Galfb1-5Galfb1-5Galfb1-5Galfb-S8 | (Galf)4-[<8] |
| SDB85 | AzOH | AzOH-[460] |
| SDB66 | AzOH | AzOH-[950] |
| SDB74 | AzOH | AzOH-[950] |
| SDB40 | AzOH | AzOH-[950] |
| SDB41 | AzOH | AzOH-[950] |
| SDB46 | AzOH | AzOH-[950] |
| SDB51 | AzOH | AzOH-[950] |

### **Supplementary Table 7:** LiGA-g (EH.xlsx)

| SDB | Modification | Axis Name |
|---|---|---|
| SDB135 | (Galb1-4[Fuca1-3]GlcNAcb1-3)3b-Sp | Tri-Lex-[430] |
| SDB198 | Neu5Aca2-3(Galb1-4GlcNAcb1-3)2b-Sp | 3'S-Di-LN-[1350] |
| SDB197 | Neu5Aca2-6(Galb1-4GlcNAcb1-3)2b-Sp | 6'S-Di-LN-[300] |
| SDB196 | Neu5Aca2-3(Galb1-4GlcNAcb1-3)3b-Sp | 3'STri-LN-[160] |
| SDB199 | Neu5Aca2-6Galb1-4GlcNAcb1-3Galb1-3GlcNAcb-Sp | 6SLN-Lec-[300] |
| SDB31 | Neu5Aca2-3[Neu5Aca2-3Galb1-3GalNAcb1-4]Galb1-4Glcb-Sp | GD1a-[110] |
| SDB78 | Neu5Aca2-8Neu5Aca2-8Neu5Aca2-8Neu5Aca2-3Galb1-4Glcb-Sp | TetraSLac-[80] |
| SDB44 | Neu5Aca2-8Neu5Aca2-8Neu5Aca2-8Neu5Aca2-3[GalNAcb1-4]Galb1-4Glcb-Sp | GQ2-[140] |
| SDB65 | Neu5Aca2-3[GalNAcb1-4]Galb1-4GlcNAcb-Sp | CT\|Sda-[350] |
| SDB68 | Neu5Aca2-8Neu5Aca2-3Galb1-4Glcb-Sp | GD3-[320] |
| SDB71 | Neu5Aca2-8Neu5Aca2-8Neu5Aca2-3[GalNAcb1-4]Galb1-4Glcb-Sp | GT2-[160] |
| SDB73 | Neu5Aca2-3Galb1-3GlcNAcb-Sp | 3'SLec-[430] |
| SDB75 | Neu5Aca2-6Galb1-4Glcb-Sp | 6'SL-[620] |
| SDB77 | Neu5Gca2-3Galb1-4Glcb-Sp | 3'SL (Gc)-[490] |
| SDB80 | Neu5Aca2-3Galb1-4GlcNAcb1-3Galb1-3GlcNAcb-Sp | 3'SLN-Lec-[410] |
| SDB83 | Neu5Aca2-3Galb1-3GlcNAcb1-3Galb1-3GlcNAcb-Sp | 3'S-Di-Lec-[270] |
| SDB88 | Neu5Aca2-3Galb1-3GlcNAcb1-3Galb1-4GlcNAcb-Sp | 3'SLecLN-[860] |
| SDB131 | Neu5Aca2-6GalNAcb1-4GlcNAcb-Sp | 6'SLDN-[620] |
| SDB132 | Neu5Aca2-3Galb1-4Glcb-Sp | GM3-[540] |
| SDB141 | Neu5Gca2-3Galb1-4GlcNAcb-Sp | 3'SLN (Gc)-[570] |
| SDB220 | Neu5Aca2-3GalNAcb1-4GlcNAcb-Sp | 3'SLDN-[460] |
| SDB201 | Neu5Aca2-8Neu5Aca2-8Neu5Aca2-3Galb1-4Glcb-Sp | GT3-[510] |
| SDB218 | Neu5Aca2-3(Galb1-4[Fuca1-3]GlcNAcb1-3)3b-Sp | 3'S-Tri-LeX-[160] |
| SDB96 | Neu5Aca2-3Galb1-3[Fuca1-4]GlcNAcb1-3Galb1-4[Fuca1-3]GlcNAcb-Sp | 3'SLeA-Lex-[570] |
| SDB62 | Neu5Aca2-3(Galb1-3[Fuca1-4]GlcNAcb1-3)2b-Sp | 3'S-Di-LeA-[160] |
| SDB98 | Neu5Aca2-3[GalNAcb1-4]Galb1-4Glcb-Sp | GM2-[590] |
| SDB99 | Neu5Aca2-3[Galb1-3GalNAcb1-4]Galb1-4Glcb-Sp | GM1-[460] |
| SDB100 | Neu5Aca2-8Neu5Aca2-3[GalNAcb1-4]Galb1-4Glcb-Sp | GD2-[460] |
| SDB192 | Neu5Aca2-3Galb1-4Glcb-Sp | GM3-[1190] |
| SDB181 | Neu5Aca2-3Galb1-4Glcb-Sp | GM3-[190] |
| SDB171 | Neu5Aca2-3Galb1-4Glcb-Sp | GM3-[460] |
| SDB162 | Neu5Aca2-3[GalNAcb1-4]Galb1-4Glcb-Sp | GM2-[1190] |
| SDB97 | Neu5Aca2-3[GalNAcb1-4]Galb1-4Glcb-Sp | GM2-[190] |
| SDB92 | Neu5Aca2-3[GalNAcb1-4]Galb1-4Glcb-Sp | GM2-[460] |
| SDB91 | Neu5Aca2-3[Galb1-3GalNAcb1-4]Galb1-4Glcb-Sp | GM1-[1190] |
| SDB39 | Neu5Aca2-3[Galb1-3GalNAcb1-4]Galb1-4Glcb-Sp | GM1-[190] |
| SDB10 | Neu5Aca2-3[Galb1-3GalNAcb1-4]Galb1-4Glcb-Sp | GM1-[460] |
| SDB13 | Neu5Aca2-3(Galb1-4[Fuca1-3]GlcNAcb1-3)2b-Sp | 3'S-Di-Lex-[410] |
| SDB161 | Neu5Gca2-6Galb1-4GlcNAcb-Sp | 6'SLN (Gc)-[430] |
| SDB150 | KDNa2-3Galb1-3GlcNAcb-Sp | 3'-KDNLec-[760] |
| SDB117 | KDNa2-3Galb1-4GlcNAcb-Sp | 3'KDNLN-[510] |
| SDB118 | Neu5Gca2-3Galb1-3GlcNAcb-Sp | 3'SLec (Gc)-[350] |
| SDB111 | (Galb1-4[Fuca1-3]GlcNAcb1-3)2b-Sp | Di-Lex-[410] |
| SDB119 | Neu5Aca2-3Galb1-4Glcb-Sp | GM3-[160] |
| SDB120 | Neu5Aca2-3Galb1-4Glcb-Sp | GM3-[160] |
| SDB122 | Neu5Aca2-3Galb1-4Glcb-Sp | GM3-[160] |
| SDB123 | Neu5Aca2-3Galb1-4Glcb-Sp | GM3-[160] |
| SDB125 | Neu5Aca2-3Galb1-4Glcb-Sp | GM3-[160] |
| SDB126 | Neu5Aca2-3Galb1-4Glcb-Sp | GM3-[380] |
| SDB127 | Neu5Aca2-3Galb1-4Glcb-Sp | GM3-[380] |
| SDB128 | Neu5Aca2-3Galb1-4Glcb-Sp | GM3-[380] |
| SDB129 | Neu5Aca2-3Galb1-4Glcb-Sp | GM3-[380] |
| SDB133 | Neu5Aca2-3Galb1-4Glcb-Sp | GM3-[380] |
| SDB136 | Neu5Aca2-3Galb1-4Glcb-Sp | GM3-[680] |

| | | |
|---|---|---|
| SDB138 | Neu5Aca2-3Galb1-4Glcb-Sp | GM3-[680] |
| SDB139 | Neu5Aca2-3Galb1-4Glcb-Sp | GM3-[680] |
| SDB140 | Neu5Aca2-3Galb1-4Glcb-Sp | GM3-[680] |
| SDB143 | Neu5Aca2-3Galb1-4Glcb-Sp | GM3-[680] |
| SDB147 | Neu5Aca2-3Galb1-4Glcb-Sp | GM3-[810] |
| SDB148 | Neu5Aca2-3Galb1-4Glcb-Sp | GM3-[810] |
| SDB149 | Neu5Aca2-3Galb1-4Glcb-Sp | GM3-[810] |
| SDB155 | Neu5Aca2-3Galb1-4Glcb-Sp | GM3-[810] |
| SDB156 | Neu5Aca2-3Galb1-4Glcb-Sp | GM3-[810] |
| SDB157 | Neu5Aca2-3Galb1-4Glcb-Sp | GM3-[950] |
| SDB158 | Neu5Aca2-3Galb1-4Glcb-Sp | GM3-[950] |
| SDB159 | Neu5Aca2-3Galb1-4Glcb-Sp | GM3-[950] |
| SDB163 | Neu5Aca2-3Galb1-4Glcb-Sp | GM3-[950] |
| SDB164 | Neu5Aca2-3Galb1-4Glcb-Sp | GM3-[950] |
| SDB165 | Neu5Aca2-3Galb1-4Glcb-Sp | GM3-[1300] |
| SDB166 | Neu5Aca2-3Galb1-4Glcb-Sp | GM3-[1300] |
| SDB170 | Neu5Aca2-3Galb1-4Glcb-Sp | GM3-[1300] |
| SDB174 | Neu5Aca2-3Galb1-4Glcb-Sp | GM3-[1300] |
| SDB183 | Neu5Aca2-3Galb1-4Glcb-Sp | GM3-[1300] |
| SDB61 | Neu5Aca2-3[GalNAcb1-4]Galb1-4Glcb-Sp | GM2-[160] |
| SDB64 | Neu5Aca2-3[GalNAcb1-4]Galb1-4Glcb-Sp | GM2-[160] |
| SDB66 | Neu5Aca2-3[GalNAcb1-4]Galb1-4Glcb-Sp | GM2-[160] |
| SDB67 | Neu5Aca2-3[GalNAcb1-4]Galb1-4Glcb-Sp | GM2-[160] |
| SDB70 | Neu5Aca2-3[GalNAcb1-4]Galb1-4Glcb-Sp | GM2-[160] |
| SDB74 | Neu5Aca2-3[GalNAcb1-4]Galb1-4Glcb-Sp | GM2-[380] |
| SDB76 | Neu5Aca2-3[GalNAcb1-4]Galb1-4Glcb-Sp | GM2-[380] |
| SDB79 | Neu5Aca2-3[GalNAcb1-4]Galb1-4Glcb-Sp | GM2-[380] |
| SDB81 | Neu5Aca2-3[GalNAcb1-4]Galb1-4Glcb-Sp | GM2-[380] |
| SDB82 | Neu5Aca2-3[GalNAcb1-4]Galb1-4Glcb-Sp | GM2-[380] |
| SDB86 | Neu5Aca2-3[GalNAcb1-4]Galb1-4Glcb-Sp | GM2-[680] |
| SDB87 | Neu5Aca2-3[GalNAcb1-4]Galb1-4Glcb-Sp | GM2-[680] |
| SDB89 | Neu5Aca2-3[GalNAcb1-4]Galb1-4Glcb-Sp | GM2-[680] |
| SDB90 | Neu5Aca2-3[GalNAcb1-4]Galb1-4Glcb-Sp | GM2-[680] |
| SDB93 | Neu5Aca2-3[GalNAcb1-4]Galb1-4Glcb-Sp | GM2-[680] |
| SDB94 | Neu5Aca2-3[GalNAcb1-4]Galb1-4Glcb-Sp | GM2-[810] |
| SDB95 | Neu5Aca2-3[GalNAcb1-4]Galb1-4Glcb-Sp | GM2-[810] |
| SDB101 | Neu5Aca2-3[GalNAcb1-4]Galb1-4Glcb-Sp | GM2-[810] |
| SDB102 | Neu5Aca2-3[GalNAcb1-4]Galb1-4Glcb-Sp | GM2-[810] |
| SDB103 | Neu5Aca2-3[GalNAcb1-4]Galb1-4Glcb-Sp | GM2-[810] |
| SDB104 | Neu5Aca2-3[GalNAcb1-4]Galb1-4Glcb-Sp | GM2-[950] |
| SDB105 | Neu5Aca2-3[GalNAcb1-4]Galb1-4Glcb-Sp | GM2-[950] |
| SDB106 | Neu5Aca2-3[GalNAcb1-4]Galb1-4Glcb-Sp | GM2-[950] |
| SDB107 | Neu5Aca2-3[GalNAcb1-4]Galb1-4Glcb-Sp | GM2-[950] |
| SDB108 | Neu5Aca2-3[GalNAcb1-4]Galb1-4Glcb-Sp | GM2-[950] |
| SDB109 | Neu5Aca2-3[GalNAcb1-4]Galb1-4Glcb-Sp | GM2-[1300] |
| SDB110 | Neu5Aca2-3[GalNAcb1-4]Galb1-4Glcb-Sp | GM2-[1300] |
| SDB113 | Neu5Aca2-3[GalNAcb1-4]Galb1-4Glcb-Sp | GM2-[1300] |
| SDB115 | Neu5Aca2-3[GalNAcb1-4]Galb1-4Glcb-Sp | GM2-[1300] |
| SDB116 | Neu5Aca2-3[GalNAcb1-4]Galb1-4Glcb-Sp | GM2-[1300] |
| SDB1 | Neu5Aca2-3[Galb1-3GalNAcb1-4]Galb1-4Glcb-Sp | GM1-[160] |
| SDB2 | Neu5Aca2-3[Galb1-3GalNAcb1-4]Galb1-4Glcb-Sp | GM1-[160] |
| SDB3 | Neu5Aca2-3[Galb1-3GalNAcb1-4]Galb1-4Glcb-Sp | GM1-[160] |
| SDB6 | Neu5Aca2-3[Galb1-3GalNAcb1-4]Galb1-4Glcb-Sp | GM1-[160] |
| SDB9 | Neu5Aca2-3[Galb1-3GalNAcb1-4]Galb1-4Glcb-Sp | GM1-[380] |
| SDB16 | Neu5Aca2-3[Galb1-3GalNAcb1-4]Galb1-4Glcb-Sp | GM1-[380] |
| SDB25 | Neu5Aca2-3[Galb1-3GalNAcb1-4]Galb1-4Glcb-Sp | GM1-[380] |

| | | |
|---|---|---|
| SDB27 | Neu5Aca2-3[Galb1-3GalNAcb1-4]Galb1-4Glcb-Sp | GM1-[380] |
| SDB34 | Neu5Aca2-3[Galb1-3GalNAcb1-4]Galb1-4Glcb-Sp | GM1-[680] |
| SDB35 | Neu5Aca2-3[Galb1-3GalNAcb1-4]Galb1-4Glcb-Sp | GM1-[680] |
| SDB36 | Neu5Aca2-3[Galb1-3GalNAcb1-4]Galb1-4Glcb-Sp | GM1-[680] |
| SDB37 | Neu5Aca2-3[Galb1-3GalNAcb1-4]Galb1-4Glcb-Sp | GM1-[680] |
| SDB38 | Neu5Aca2-3[Galb1-3GalNAcb1-4]Galb1-4Glcb-Sp | GM1-[680] |
| SDB40 | Neu5Aca2-3[Galb1-3GalNAcb1-4]Galb1-4Glcb-Sp | GM1-[810] |
| SDB41 | Neu5Aca2-3[Galb1-3GalNAcb1-4]Galb1-4Glcb-Sp | GM1-[810] |
| SDB42 | Neu5Aca2-3[Galb1-3GalNAcb1-4]Galb1-4Glcb-Sp | GM1-[810] |
| SDB43 | Neu5Aca2-3[Galb1-3GalNAcb1-4]Galb1-4Glcb-Sp | GM1-[810] |
| SDB45 | Neu5Aca2-3[Galb1-3GalNAcb1-4]Galb1-4Glcb-Sp | GM1-[810] |
| SDB46 | Neu5Aca2-3[Galb1-3GalNAcb1-4]Galb1-4Glcb-Sp | GM1-[950] |
| SDB48 | Neu5Aca2-3[Galb1-3GalNAcb1-4]Galb1-4Glcb-Sp | GM1-[950] |
| SDB49 | Neu5Aca2-3[Galb1-3GalNAcb1-4]Galb1-4Glcb-Sp | GM1-[950] |
| SDB50 | Neu5Aca2-3[Galb1-3GalNAcb1-4]Galb1-4Glcb-Sp | GM1-[950] |
| SDB51 | Neu5Aca2-3[Galb1-3GalNAcb1-4]Galb1-4Glcb-Sp | GM1-[950] |
| SDB53 | Neu5Aca2-3[Galb1-3GalNAcb1-4]Galb1-4Glcb-Sp | GM1-[1300] |
| SDB54 | Neu5Aca2-3[Galb1-3GalNAcb1-4]Galb1-4Glcb-Sp | GM1-[1300] |
| SDB55 | Neu5Aca2-3[Galb1-3GalNAcb1-4]Galb1-4Glcb-Sp | GM1-[1300] |
| SDB57 | Neu5Aca2-3[Galb1-3GalNAcb1-4]Galb1-4Glcb-Sp | GM1-[1300] |
| SDB60 | Neu5Aca2-3[Galb1-3GalNAcb1-4]Galb1-4Glcb-Sp | GM1-[1300] |
| SDB184 | Neu5Aca2-3[Neu5Aca2-3Galb1-3GalNAcb1-4]Galb1-4Glcb-Sp | GD1a-[160] |
| SDB185 | Neu5Aca2-3[Neu5Aca2-3Galb1-3GalNAcb1-4]Galb1-4Glcb-Sp | GD1a-[160] |
| SDB187 | Neu5Aca2-3[Neu5Aca2-3Galb1-3GalNAcb1-4]Galb1-4Glcb-Sp | GD1a-[160] |
| SDB188 | Neu5Aca2-3[Neu5Aca2-3Galb1-3GalNAcb1-4]Galb1-4Glcb-Sp | GD1a-[160] |
| SDB189 | Neu5Aca2-3[Neu5Aca2-3Galb1-3GalNAcb1-4]Galb1-4Glcb-Sp | GD1a-[160] |
| SDB190 | Neu5Aca2-3[Neu5Aca2-3Galb1-3GalNAcb1-4]Galb1-4Glcb-Sp | GD1a-[380] |
| SDB191 | Neu5Aca2-3[Neu5Aca2-3Galb1-3GalNAcb1-4]Galb1-4Glcb-Sp | GD1a-[380] |
| SDB193 | Neu5Aca2-3[Neu5Aca2-3Galb1-3GalNAcb1-4]Galb1-4Glcb-Sp | GD1a-[380] |
| SDB194 | Neu5Aca2-3[Neu5Aca2-3Galb1-3GalNAcb1-4]Galb1-4Glcb-Sp | GD1a-[380] |
| SDB195 | Neu5Aca2-3[Neu5Aca2-3Galb1-3GalNAcb1-4]Galb1-4Glcb-Sp | GD1a-[380] |
| SDB200 | Neu5Aca2-3[Neu5Aca2-3Galb1-3GalNAcb1-4]Galb1-4Glcb-Sp | GD1a-[680] |
| SDB202 | Neu5Aca2-3[Neu5Aca2-3Galb1-3GalNAcb1-4]Galb1-4Glcb-Sp | GD1a-[680] |
| SDB203 | Neu5Aca2-3[Neu5Aca2-3Galb1-3GalNAcb1-4]Galb1-4Glcb-Sp | GD1a-[680] |
| SDB204 | Neu5Aca2-3[Neu5Aca2-3Galb1-3GalNAcb1-4]Galb1-4Glcb-Sp | GD1a-[680] |
| SDB205 | Neu5Aca2-3[Neu5Aca2-3Galb1-3GalNAcb1-4]Galb1-4Glcb-Sp | GD1a-[680] |
| SDB206 | Neu5Aca2-3[Neu5Aca2-3Galb1-3GalNAcb1-4]Galb1-4Glcb-Sp | GD1a-[810] |
| SDB207 | Neu5Aca2-3[Neu5Aca2-3Galb1-3GalNAcb1-4]Galb1-4Glcb-Sp | GD1a-[810] |
| SDB208 | Neu5Aca2-3[Neu5Aca2-3Galb1-3GalNAcb1-4]Galb1-4Glcb-Sp | GD1a-[810] |
| SDB209 | Neu5Aca2-3[Neu5Aca2-3Galb1-3GalNAcb1-4]Galb1-4Glcb-Sp | GD1a-[810] |
| SDB210 | Neu5Aca2-3[Neu5Aca2-3Galb1-3GalNAcb1-4]Galb1-4Glcb-Sp | GD1a-[810] |
| SDB211 | Neu5Aca2-3[Neu5Aca2-3Galb1-3GalNAcb1-4]Galb1-4Glcb-Sp | GD1a-[950] |
| SDB212 | Neu5Aca2-3[Neu5Aca2-3Galb1-3GalNAcb1-4]Galb1-4Glcb-Sp | GD1a-[950] |
| SDB213 | Neu5Aca2-3[Neu5Aca2-3Galb1-3GalNAcb1-4]Galb1-4Glcb-Sp | GD1a-[950] |
| SDB214 | Neu5Aca2-3[Neu5Aca2-3Galb1-3GalNAcb1-4]Galb1-4Glcb-Sp | GD1a-[950] |
| SDB215 | Neu5Aca2-3[Neu5Aca2-3Galb1-3GalNAcb1-4]Galb1-4Glcb-Sp | GD1a-[950] |
| SDB219 | Neu5Aca2-3[Neu5Aca2-3Galb1-3GalNAcb1-4]Galb1-4Glcb-Sp | GD1a-[1300] |
| SDB221 | Neu5Aca2-3[Neu5Aca2-3Galb1-3GalNAcb1-4]Galb1-4Glcb-Sp | GD1a-[1300] |
| SDB222 | Neu5Aca2-3[Neu5Aca2-3Galb1-3GalNAcb1-4]Galb1-4Glcb-Sp | GD1a-[1300] |
| SDB223 | Neu5Aca2-3[Neu5Aca2-3Galb1-3GalNAcb1-4]Galb1-4Glcb-Sp | GD1a-[1300] |
| SDB224 | Neu5Aca2-3[Neu5Aca2-3Galb1-3GalNAcb1-4]Galb1-4Glcb-Sp | GD1a-[1300] |

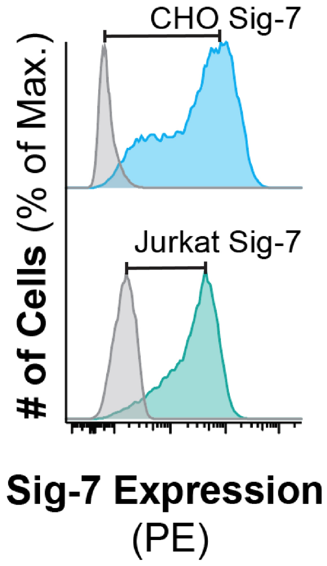


**Supplementary Figure 1:** Anti-Siglec-7 antibody staining of Siglec-7$^+$ Jurkat and Siglec-7$^+$ CHO cells, and respective wild-type cells.
