## Supplementary material for "Measuring carbohydrate recognition profile of lectins on live cells using liquid glycan array (LiGA)": Zipped folder containing all the software, MALDI TOF spectra and analyzed files: 00-install-prerequisites.R

```
# Install missing packages used by the other R scripts in this directory.
#
# WARNING: If many package dependancies need installation this can take ~30 minutes.
# WARNING: If you mantain specific versisons of libraries, review this script
#          before overwriting your existing install.

# choose the install library - default to first user library
install.lib <- sub(":.*$", "", Sys.getenv("R_LIBS_USER"))

# create the install library directories if they do not already exist
dir.create(path = install.lib, recursive = TRUE, showWarnings = FALSE)

# edgeR is distributed through BioConductor - install it if necessary, then get edgeR
# https://bioconductor.org/packages/release/bioc/html/edgeR.html
if (!requireNamespace("BiocManager", quietly = TRUE)){
    install.packages("BiocManager", lib = install.lib)
}
if(!"edgeR" %in% installed.packages()[,"Package"]){
    BiocManager::install("edgeR", lib = install.lib)
}
if(!"statmod" %in% installed.packages()[,"Package"]){
    install.packages("statmod", lib = install.lib)  # apparently edgeR may require statmod functions
}
# knitr/Rmarkdown, install it after tinytex (used of knitr PDF output)
if(!"tinytex" %in% installed.packages()[,"Package"]){
    install.packages("tinytex", lib = install.lib)
}
tinytex::install_tinytex()  # to uninstall TinyTeX, run tinytex::uninstall_tinytex() 
if(!"rmarkdown" %in% installed.packages()[,"Package"]){
    install.packages("rmarkdown", lib = install.lib)    # used for report generation
}
# ggplot2 - widely used plot package
if(!"dplyr" %in% installed.packages()[,"Package"]){
    install.packages("dplyr", lib = install.lib)      # common graphing package
}
# dose-response curve package
if(!"stringr" %in% installed.packages()[,"Package"]){
    install.packages("stringr", lib = install.lib)
}

#install.packages("tidyverse", lib = install.lib)
```
