## Supplementary material for "Measuring carbohydrate recognition profile of lectins on live cells using liquid glycan array (LiGA)": Zipped folder containing all the software, MALDI TOF spectra and analyzed files: DE.rmd

````
---
params:
        TableName: "/media/Data/48HD/LiGA Script/TEST.txt"
        TestCols: "9,10,11,12,13,14,15,16"
        ControlCols: "1,2,3,4,5,6,7,8"
        saveName: "DE_TEST.txt"
---

---
title: "DE Analysis"
output: html_document
---

```{r setup, include=FALSE}
knitr::opts_chunk$set(echo = TRUE)
```

## Loading Required Libraries
```{r echo=TRUE}
# This is a test script for DE analysis, the input data are table files
# The output of this script is a set of pair-wise comparison tables, including original reads, logFoldChanges, pValue and adjusted-pValue
library(edgeR)
#library(dplyr)
#library(tidyr)
library(ggplot2)
library(knitr)
#library(stringr)
```

```{r getPrams, echo=TRUE}
if (params$TableName == ".txt"){
         knit_exit("Dataset is not defined!")
         write.table(data.frame(), params$saveName,quote=FALSE, sep ="\t", row.names=FALSE)
 }

TestCols = as.numeric(unlist(strsplit(params$TestCols,',')))
ControlCols = as.numeric(unlist(strsplit(params$ControlCols, ',')))
```

## Loading Files
This section loads the dataset table.
```{r loading, echo=TRUE, message = FALSE, warning = FALSE}
# group the files
FileGroup = factor(1:2)

# load the table files and process the data
counts = read.table(params$TableName, header = 1, check.names = FALSE)

totalNcol = ncol(counts)-1
if(totalNcol!= length(TestCols)+length(ControlCols)){
        knit_exit("number of control and test columns does not match the dataset!")  
        write.table(data.frame(), params$saveName,quote=FALSE, sep ="\t", row.names=FALSE)
}
group = rep(1, totalNcol)
if(1 %in% TestCols){
        colNamesDesign = c("test", "control")
}else{
        colNamesDesign = c("control", "test")
}

for(j in 1:length(TestCols)){
        group[TestCols[j]] = 2
}
contrastLevels = c("test-control")

# assign seq to row names 
row.names(counts) = counts[,1]
counts = counts[,-1]
colgroup = group
group = factor(group)
```
There are total `r dim(counts)[1]` sequences to be tested. And the total reads of each columns are show as follows.

```{r echo=TRUE, results='asis'}
counts[is.na(counts)]=0

barplot(colSums(counts), col = group,ylab = "Library sizes", main="")
```


## DE Analysis Using edgeR library
This section performs DE analysis using edgeR library. TMM normalization methods and Quasi-likelihood F-Test are used. The results are saved in file "finalDE.txt".
```{r DE, echo=FALSE, results='asis', warning=FALSE, message=FALSE}
# create DGEList
counts[is.na(counts)]=0
RG = DGEList(counts = counts, group = group)

# the following is from the tutorial, needs to be further modified
RG = calcNormFactors(RG)

tmm_data <- cpm(RG, normalized.lib.sizes = TRUE)
sd_control <- apply(tmm_data[,TestCols], 1, sd)    # handle reversed inputs
sd_test    <- apply(tmm_data[,ControlCols], 1, sd)

design = model.matrix(~0+group, data = RG$samples)
colnames(design) = colNamesDesign
tmp = try(estimateDisp(RG, design, robust=TRUE), TRUE)
if(inherits(tmp, "try-error")){
        RG = estimateCommonDisp(RG)
        RG = estimateTagwiseDisp(RG)
}else{
        RG = estimateDisp(RG, design, robust=TRUE)
        plotMDS(RG,top=dim(RG$counts)[1],col=colgroup)
}

# use QLFit to generate the model
fit = glmQLFit(RG, design, robust = TRUE)

coefTbl = data.frame(fit$coefficients)
names(coefTbl) = paste0(names(coefTbl), '_coef')
finalTblAll = data.frame(sequence=row.names(fit$coefficients), RG$counts, coefTbl, check.names = FALSE)
remove(coefTbl)
for(i in names(data.frame(fit$coefficients))){
        finalTblAll[[paste0(i,"_CPM")]] = exp(finalTblAll[[paste0(i,"_coef")]])*1E6
}

#colNames = names(finalTblAll)
cn = makeContrasts(contrasts = contrastLevels[1], levels=design)
qlfi = glmQLFTest(fit, contrast = cn)

    qlfiTbl <- data.frame(sequence = rownames(RG$counts),
                 sd_test = sd_test, sd_control = sd_control,
                 #var.post = qlfi$table$var.post, # not the dispersion I want 
                 logFC = qlfi$table$logFC,
                 PValue = qlfi$table$PValue,
                 QValue = p.adjust(qlfi$table$PValue, method = "BH"))

#qlfiTbl = data.frame(sequence=rownames(RG$counts),qlfi$table[,c('logFC','PValue')])
#names(qlfiTbl) = c('sequence', names(qlfiTbl)[2:ncol(qlfiTbl)])
finalTblAll = merge(finalTblAll, qlfiTbl, all = FALSE, by.x = 'sequence', by.y = 'sequence', sort = FALSE)
finalTblAll[is.na(finalTblAll)] = 0
#colNames = c(colNames,  'logFC','PValue')


#names(finalTblAll) = colNames
# finalTblAll$sequence = separate(finalTblAll, sequence, c('Mod', 'sequence'), sep = '-' )
write.table(finalTblAll, params$saveName,
            quote=FALSE, sep ="\t", row.names=FALSE)


```
````
