## Supplementary material for "Measuring carbohydrate recognition profile of lectins on live cells using liquid glycan array (LiGA)": Zipped folder containing all the software, MALDI TOF spectra and analyzed files: differential-test.R

```
############  Clears All Data  ############
if(!is.null(dev.list())) dev.off() # Clear plots
cat("\014") # Clear console
rm(list=ls())# Clea n workspace
#############  Library  ##############

library(rmarkdown)
library(dplyr)
library(stringr)


wd <- setwd("/Users/mirat/Library/CloudStorage/Dropbox/Derda Lab/Glycan Array/differential-testing")

LiGA.data.dir <- "LiGA-data"
dict_dir <- "dictionaries"
order_table_dir <- "figure data tables/axes"
campaign_dir <- "figure data tables/new requests"
scriptDir <- wd	# location script to run in
input_filename_list <- list.files(path = campaign_dir, pattern = "*.csv")
input_filename_list
#input_filename_list <- list.files(path = campaign_dir, pattern = ".*.csv")


# extract the description fields from the LiGA datafile names
# inputs: a dataframe and the index of a column of the filenames
# results are additional columns prepended to the result frame
parse_LiGA_filenames <- function(files, index) {
    date_regex <- "(2[0-9]{3})(0[1-9]|1[012])(0[1-9]|[12][0-9]|3[01])"
    chem_regex <- "-([0-9]+)([A-Z]{2,})([a-z]{2,})([A-Z]{2,})-([A-Z]+)"
    parts <- str_match(basename(as.character(files[[index]])), paste0(date_regex, chem_regex))
    parsed_names <- data.frame(parts[,2:9], files, stringsAsFactors = FALSE)
    colnames(parsed_names)[1:8] <- c("Year", "Month", "Day", "Library",
                                     "Modification", "Target", "Postprocess", "ID")
    parsed_names
}


### read in the dictionary file, keep (non-blank) SDB and Axis.name pairs
### remove leading and trailing periods in the column names, duplicate SDB entries
read_dictionary_tables <- function(dictionary_basename, order_table_basename, label_column = "Axis.name",
                                   known_SDB_list, dict_dir = "dictionaries", order_table_dir = "figure data tables/axes") {
    # if more than one file is found just use the first
    dict_filenames <- list.files(path = dict_dir,
                                 pattern = paste0(dictionary_basename, ".txt"))
    
    dictionary <- read.delim(file.path(dict_dir, dict_filenames[1]), header = TRUE,
                             sep = "\t", stringsAsFactors = FALSE)
    colnames(dictionary) <- sub("^\\.*(.*?)\\.*$", "\\1", colnames(dictionary))
    
    # ignore unneeded columns for simplicity
    colnames(dictionary)[colnames(dictionary) == label_column] <- "Axis.name"
    dictionary <- dictionary[,colnames(dictionary) %in% c("SDB", "Axis.name", "Alphanum")]
    
    # handle duplicated SDB entries
    dupl_dictionary_SDBs <- duplicated(dictionary$SDB)
    dictionary <- dictionary[!dupl_dictionary_SDBs,]
    dictionary$Axis.name[dictionary$SDB %in% dupl_dictionary_SDBs] = "mix"
    
    # drop unknown SDBs and blank entries from the dictionary
    dictionary <- dictionary[dictionary$SDB %in% known_SDB_list,]
    dictionary <- dictionary[dictionary$Axis.name != "" & dictionary$SDB != "",]
    if(length(dictionary) != 3 | length(dictionary$SDB) < 1){
        stop(paste0("The dictionary file appears to be empty or lacks the SDB, Alphanum, and/or Label columns: ",
                    dict_filenames[1], " ", input_columns$Label[i,]))
    }
    
    # read in the order table - drop extra columns and duplicate Alphanum entries
    order_table <- read.delim(file.path(order_table_dir, paste0(order_table_basename, ".csv")), 
                              header = TRUE, sep = ",", stringsAsFactors = FALSE)
    colnames(order_table) <- sub("^\\.*(.*?)\\.*$", "\\1", colnames(order_table))
    order_table <- order_table[,colnames(order_table) %in% c("Alphanum", "Order")]
    
    # remove all copies of duplicate Alphanum entries - multiples propagate through joins
    # TODO: could rescue repeated Alphanum's where all entries agree on the same order
    dupl_order_table_Alphanum <- order_table$Alphanum[duplicated(order_table$Alphanum)]
    order_table <- order_table[!order_table$Alphanum %in% dupl_order_table_Alphanum,]
    
    if(length(order_table) != 2 | length(order_table$Order) < 1){
        stop(paste0("The Order Table file appears to be empty or lacks the Order column: ",
                    order_table_basename))
    }
    
    dictionary <- dictionary %>% left_join(order_table, by = "Alphanum")
    dictionary <- dictionary[,colnames(dictionary) %in% c("SDB", "Axis.name", "Order")]
    
    # make NA in the Order a special position at the end of the list
    # and any duplicated SDBs a special order position at the front of the list
    dictionary$Order[is.na(dictionary$Order)] <- max(dictionary$Order, na.rm = TRUE) + 1
    dictionary$Order[dictionary$SDB %in% dupl_dictionary_SDBs] <- min(dictionary$Order) - 1
    
    dictionary <- dictionary[order(dictionary$Order, nchar(dictionary$SDB), dictionary$SDB),]
    
    dictionary
}	

### read dataframe listing datafiles, columns to select, and matching dictionary
### return a single merged table of the requested columns
read_and_merge_columns <- function(input_columns, LiGA_dir = LiGA.data.dir,
                                   dict_dir = dict_dir, order_table_dir = order_table_dir) {
    
    # verify the inputs
    if(length(colnames(input_columns)[colnames(input_columns) %in%
                                      c("Filename", "Columns", "Dictionary", "Labels", "OrderTable")]) != 5){
        stop("One or more required columns (Filename, Columns, Dictionary, Labels, OrderTable) not found.")
    }
    # TODO check that columns are numeric values
    
    if(length(unique(input_columns$OrderTable)) != 1){
        stop("Multiple OrderTables in input .")
    }
    
    # apply the LiGA_dir prefix if given
    if(LiGA_dir != "") {
        input_columns$Filename <- file.path(LiGA_dir, paste0(trimws(input_columns$Filename), ".txt"))
    }
    
    # TODO: what if filenames do not meet the expected pattern?
    parsed_names <- parse_LiGA_filenames(input_columns, "Filename")
    parsed_names$root <- paste0(parsed_names$Year, parsed_names$Month,
                                parsed_names$Day, "-", parsed_names$Library, parsed_names$Modification,
                                parsed_names$Target, parsed_names$Postprocess)
    
    # if an ExcludedSDBs column is not present create an empty one
    if(!"ExcludedSDBs" %in% colnames(parsed_names)){
        parsed_names$ExcludedSDBs <- ""
    }
    if(!"Type" %in% colnames(parsed_names)){
        parsed_names$Type <- ""
    }
    
    
    # read in the table of pre-labelled SDB sequences
    identified_seqs <- read.table('identified-sequence-table2.txt', header = TRUE,
                                  stringsAsFactors = FALSE);
    
    # create data structures to store the results
    merged_data <- data.frame(SDB = character(), Barcode = character(),
                              stringsAsFactors = FALSE)
    column_types <- c("Barcode", "Barcode")
    
    # loop over the listed files, merging columns into the output data frame
    for(i in 1:length(parsed_names$Filename)){
        # read in the file
        file_data <- read.table(parsed_names$Filename[i], header = TRUE,
                                skip = 0, sep = "\t", quote = "\"", stringsAsFactors = FALSE)
        # discard the sum total line (Nuc == XX) - can be recalculated later
        file_data <- file_data[file_data$Nuc != "XX",]
        
        # match each sequence with known SDBs
        file_data <- right_join(identified_seqs, file_data, by = "Nuc")
        
        # read in the dictionary file and order table
        dictionary <- read_dictionary_tables(parsed_names$Dictionary[i], parsed_names$OrderTable[i],
                                             parsed_names$Label[i], known_SDB_list = identified_seqs$SDB,
                                             dict_dir = dict_dir, order_table_dir = order_table_dir)
        
        # use Axis.name labels where the SDBs match, fallback to the SDB if unknown
        # and drop data for any excluded cases
        file_data <- full_join(file_data, dictionary, by = "SDB")
        file_data$Barcode <- file_data$Axis.name
        unknown_barcode <- is.na(file_data$Axis.name)
        file_data$Barcode[unknown_barcode] <- "" #file_data$SDB[unknown_barcode]
        file_data <- file_data[!file_data$SDB %in% parsed_names$ExcludedSDBs[[i]],]
        #file_data <- file_data[order(file_data$Order, nchar(file_data$SDB), file_data$SDB, na.last = TRUE),]
        
        # list the indices of columns containing the read counts
        # exclude columns named "Mod", "Nuc", "AA", etc.
        data_columns <- (1:NCOL(file_data))[!colnames(file_data) %in% c("SDB",
                                                                        "Distance", "index", "mindex", "Primer", "Nuc", "Mod", "AA",
                                                                        "Axis.name", "Order", "Barcode")]
        
        # choose the requested subset of the data columns
        active_columns <- data_columns[as.vector(parsed_names$Columns[[i]])]
        for(col in 1:length(active_columns)){
            column_types <- c(column_types, as.character(parsed_names$Type[i]))
            # new column name: filename root followed by the column number
            column_name <- paste0(parsed_names$root[i], col)
            colnames(file_data)[active_columns[col]] <- column_name
            
            # extract a data_frame with the barcode and one column
            # of read counts, summing read counts over barcodes
            data_column <- file_data %>% group_by(SDB, Barcode) %>%
                summarize(count_sum = sum(!!as.name(column_name)))
            
            # fix the column name and merge it to the result frame
            colnames(data_column)[3] <- column_name
            merged_data <- full_join(merged_data, data_column, by = c("SDB", "Barcode"))
            
            # replace any NAs in the count data with zeros - restore NA SDBs
            sdb_names <- merged_data$SDB
            merged_data[is.na(merged_data)] <- 0
            merged_data$SDB <- sdb_names
        }
    }
    ordering_frame <- merged_data %>% left_join(dictionary, by = c("SDB", "Barcode" = "Axis.name"))
    ordering <- order(ordering_frame$Order,			# primary sort by provided Order
                      ordering_frame$Barcode == "",		# empty labels at the end
                      sub("-\\[[0-9<]*]$", "",  ordering_frame$Barcode),   # break ties by glycan (alphabetic)
                      suppressWarnings(as.integer(
                          sub(".*-\\[<?0*([1-9][0-9]*)\\]$", "\\1", ordering_frame$Barcode))),
                      ordering_frame$Barcode,			# finally just use the raw string
                      nchar(ordering_frame$SDB),			# then use the SDB
                      ordering_frame$SDB, na.last = TRUE)
    merged_data <- merged_data[ordering,]
    list(table = merged_data, types = column_types)
}


# routine to run the edgeR comparison process on data described in the campaign_dir
for(i in 1:length(input_filename_list)){
    # drop the extension (if any) from the filename to get the root used for naming new outputs
    # note: special handling of indexing at length == 1 - does not work for all dot filenames
    filename_root <- unlist(strsplit(basename(input_filename_list[i]), split = "[.]"))
    filename_root <- paste(filename_root[1:length(filename_root)-1], collapse = ".")
    
    # read the campaign file
    testing <- read.table(file.path(campaign_dir, input_filename_list[i]),
                          sep = ",", quote = "\"", header = TRUE, stringsAsFactors = FALSE)
    
    # check for the expected columns and skip any file without
    if(length(unique(colnames(testing)[colnames(testing) %in%
                                       c("Type", "Filename", "Columns", "Dictionary", "Labels", "OrderTable")])) == 6){
        
        print(paste0("Processing: ", filename_root))
        testing$Type <- factor(testing$Type)
        testing$Columns <- lapply(strsplit(testing$Columns, ","), as.integer)
        
        # R converts blank strings into NA on input so reverse this before extracting the list
        testing$ExcludedSDBs[is.na(testing$ExcludedSDBs)] <- ""
        testing$ExcludedSDBs <- lapply(strsplit(testing$ExcludedSDBs, ","), trimws)
        
        testing_data <- read_and_merge_columns(testing, LiGA_dir = LiGA.data.dir,
                                               dict_dir = dict_dir, order_table_dir = order_table_dir)
        
        # Jessica's DE script does not allow NA row entries so delete them now
        # similarly only test and control columns are allowed
        reduced_types <- testing_data$types[-2]
        subset_indices <- reduced_types %in% c("Barcode", "Test", "Control")
        test_columns <- (0:(length(reduced_types[subset_indices])-1))[reduced_types[subset_indices] == "Test"]
        control_columns <- (0:(length(reduced_types[subset_indices])-1))[reduced_types[subset_indices] == "Control"]
        
        # label (first) column must be named sequence
        subset_table <- testing_data$table[-2]
        colnames(subset_table)[1] <- "sequence"
        subset_table$sequence <- gsub("[ \t]", ".",
                                      paste0(testing_data$table$Barcode, ".", testing_data$table$SDB))
        
        # consider different subsets of the known SDBs
        for(case in 3:3){
            if(case == 1){
                analysis_root = paste0(filename_root, "-all")
            } 
            if(case == 2){
                analysis_root = paste0(filename_root, "-knownSDBs")
                subset_table <- subset_table[subset_table$sequence != ".NA",]
            } 
            if(case == 3){
                analysis_root = paste0(filename_root, "-knownGlycans")
                subset_table <- subset_table[subset_table != paste0(".", testing_data$table$SDB),]
            }
            data_table_filename <- file.path(".", paste0(analysis_root, ".tsv"))
            write.table(subset_table[!is.na(subset_table$sequence),subset_indices],
                        file = data_table_filename, sep = "\t", quote = FALSE, row.names = FALSE)
            
            # # use edgeR, apply TMM and calculate the mean & st. dev. for each condition
            saveName = paste0("compare_", analysis_root, ".txt")
            htmlName = paste0("compare_", analysis_root, ".html")
            
            # test and control seem backwards here, but this is to label the final outputs correctly
            render(file.path(scriptDir, "auxiliary/DE.rmd"),
                   params = list("TableName" = file.path("..", data_table_filename),
                                 "TestCols" = paste(test_columns, collapse = ","),
                                 "ControlCols" = paste(control_columns, collapse = ","),
                                 "saveName" = saveName))
            file.remove(data_table_filename)
            
            print("Warning message:")
            file.copy(file.path(scriptDir,"auxiliary", saveName), scriptDir, overwrite = TRUE)
            file.remove(file.path(scriptDir,"auxiliary", saveName))
            #file.copy(file.path(scriptDir,"auxiliary/DE.html"), scriptDir, overwrite = TRUE)
            #file.remove(file.path(scriptDir,"auxiliary/DE.html"))
            tmp = try(file.rename("auxiliary/DE.html", htmlName), TRUE)
            if(inherits(tmp, "try-error")){
                log = paste0(log, "\nWarning: DE in ", scriptDir, " failed!")
            }
        }
    }
}

# 
# parse_LiGA_filenames <- function(files, index) {
#   date_regex <- "(2[0-9]{3})(0[1-9]|1[012])(0[1-9]|[12][0-9]|3[01])"
#   chem_regex <- "-([0-9]+)([A-Z]{2,})([a-z]{2,})([A-Z]{2,})-([A-Z]+)"
#   parts <- str_match(basename(as.character(files[[index]])), paste0(date_regex, chem_regex))
#   parsed_names <- data.frame(parts[,2:9], files, stringsAsFactors = FALSE)
#   colnames(parsed_names)[1:8] <- c("Year", "Month", "Day", "Library",
#                                    "Modification", "Target", "Postprocess", "ID")
#   parsed_names
# }
# 
# 
# ### read in the dictionary file, keep (non-blank) SDB and Axis.name pairs
# ### remove leading and trailing periods in the column names, duplicate SDB entries
# read_dictionary_tables <- function(dictionary_basename, order_table_basename, label_column = "Axis.name",
#                                    known_SDB_list, dict_dir = "dictionaries", order_table_dir = "figure data tables/axes") {
#   # if more than one file is found just use the first
#   dict_filenames <- list.files(path = dict_dir,
#                                pattern = paste0(dictionary_basename, ".txt"))
#   
#   dictionary <- read.delim(file.path(dict_dir, dict_filenames[1]), header = TRUE,
#                            sep = "\t", stringsAsFactors = FALSE)
#   colnames(dictionary) <- sub("^\\.*(.*?)\\.*$", "\\1", colnames(dictionary))
#   
#   # ignore unneeded columns for simplicity
#   colnames(dictionary)[colnames(dictionary) == label_column] <- "Axis.name"
#   dictionary <- dictionary[,colnames(dictionary) %in% c("SDB", "Axis.name", "Alphanum")]
#   
#   # handle duplicated SDB entries
#   dupl_dictionary_SDBs <- duplicated(dictionary$SDB)
#   dictionary <- dictionary[!dupl_dictionary_SDBs,]
#   dictionary$Axis.name[dictionary$SDB %in% dupl_dictionary_SDBs] = "mix"
#   
#   # drop unknown SDBs and blank entries from the dictionary
#   dictionary <- dictionary[dictionary$SDB %in% known_SDB_list,]
#   dictionary <- dictionary[dictionary$Axis.name != "" & dictionary$SDB != "",]
#   if(length(dictionary) != 3 | length(dictionary$SDB) < 1){
#     stop(paste0("The dictionary file appears to be empty or lacks the SDB, Alphanum, and/or Label columns: ",
#                 dict_filenames[1], " ", input_columns$Label[i,]))
#   }
#   
#   # read in the order table - drop extra columns and duplicate Alphanum entries
#   order_table <- read.delim(file.path(order_table_dir, paste0(order_table_basename, ".csv")), 
#                             header = TRUE, sep = ",", stringsAsFactors = FALSE)
#   colnames(order_table) <- sub("^\\.*(.*?)\\.*$", "\\1", colnames(order_table))
#   order_table <- order_table[,colnames(order_table) %in% c("Alphanum", "Order")]
#   
#   # remove all copies of duplicate Alphanum entries - multiples propagate through joins
#   # TODO: could rescue repeated Alphanum's where all entries agree on the same order
#   dupl_order_table_Alphanum <- order_table$Alphanum[duplicated(order_table$Alphanum)]
#   order_table <- order_table[!order_table$Alphanum %in% dupl_order_table_Alphanum,]
#   
#   if(length(order_table) != 2 | length(order_table$Order) < 1){
#     stop(paste0("The Order Table file appears to be empty or lacks the Order column: ",
#                 order_table_basename))
#   }
#   
#   dictionary <- dictionary %>% left_join(order_table, by = "Alphanum")
#   dictionary <- dictionary[,colnames(dictionary) %in% c("SDB", "Axis.name", "Order")]
#   
#   # make NA in the Order a special position at the end of the list
#   # and any duplicated SDBs a special order position at the front of the list
#   dictionary$Order[is.na(dictionary$Order)] <- max(dictionary$Order, na.rm = TRUE) + 1
#   dictionary$Order[dictionary$SDB %in% dupl_dictionary_SDBs] <- min(dictionary$Order) - 1
#   
#   dictionary <- dictionary[order(dictionary$Order, nchar(dictionary$SDB), dictionary$SDB),]
#   
#   dictionary
# }	
# 
# ### read dataframe listing datafiles, columns to select, and matching dictionary
# ### return a single merged table of the requested columns
# read_and_merge_columns <- function(input_columns, LiGA_dir = LiGA.data.dir,
#                                    dict_dir = dict_dir, order_table_dir = order_table_dir) {
#   
#   # verify the inputs
#   if(length(colnames(input_columns)[colnames(input_columns) %in%
#                                     c("Filename", "Columns", "Dictionary", "Labels", "OrderTable")]) != 5){
#     stop("One or more required columns (Filename, Columns, Dictionary, Labels, OrderTable) not found.")
#   }
#   # TODO check that columns are numeric values
#   
#   if(length(unique(input_columns$OrderTable)) != 1){
#     stop("Multiple OrderTables in input .")
#   }
#   
#   # apply the LiGA_dir prefix if given
#   if(LiGA_dir != "") {
#     input_columns$Filename <- file.path(LiGA_dir, paste0(trimws(input_columns$Filename), ".txt"))
#   }
#   
#   # TODO: what if filenames do not meet the expected pattern?
#   parsed_names <- parse_LiGA_filenames(input_columns, "Filename")
#   parsed_names$root <- paste0(parsed_names$Year, parsed_names$Month,
#                               parsed_names$Day, "-", parsed_names$Library, parsed_names$Modification,
#                               parsed_names$Target, parsed_names$Postprocess)
#   
#   # if an ExcludedSDBs column is not present create an empty one
#   if(!"ExcludedSDBs" %in% colnames(parsed_names)){
#     parsed_names$ExcludedSDBs <- ""
#   }
#   if(!"Type" %in% colnames(parsed_names)){
#     parsed_names$Type <- ""
#   }
#   
#   
#   # read in the table of pre-labelled SDB sequences
#   identified_seqs <- read.table('identified-sequence-table.gz', header = TRUE,
#                                 stringsAsFactors = FALSE);
#   
#   # create data structures to store the results
#   merged_data <- data.frame(SDB = character(), Barcode = character(),
#                             stringsAsFactors = FALSE)
#   column_types <- c("Barcode", "Barcode")
#   
#   # loop over the listed files, merging columns into the output data frame
#   for(i in 1:length(parsed_names$Filename)){
#     # read in the file
#     file_data <- read.table(parsed_names$Filename[i], header = TRUE,
#                             skip = 0, sep = "\t", quote = "\"", stringsAsFactors = FALSE)
#     # discard the sum total line (Nuc == XX) - can be recalculated later
#     file_data <- file_data[file_data$Nuc != "XX",]
#     
#     # match each sequence with known SDBs
#     file_data <- right_join(identified_seqs, file_data, by = "Nuc")
#     
#     # read in the dictionary file and order table
#     dictionary <- read_dictionary_tables(parsed_names$Dictionary[i], parsed_names$OrderTable[i],
#                                          parsed_names$Label[i], known_SDB_list = identified_seqs$SDB,
#                                          dict_dir = dict_dir, order_table_dir = order_table_dir)
#     
#     # use Axis.name labels where the SDBs match, fallback to the SDB if unknown
#     # and drop data for any excluded cases
#     file_data <- full_join(file_data, dictionary, by = "SDB")
#     file_data$Barcode <- file_data$Axis.name
#     unknown_barcode <- is.na(file_data$Axis.name)
#     file_data$Barcode[unknown_barcode] <- "" #file_data$SDB[unknown_barcode]
#     file_data <- file_data[!file_data$SDB %in% parsed_names$ExcludedSDBs[[i]],]
#     #file_data <- file_data[order(file_data$Order, nchar(file_data$SDB), file_data$SDB, na.last = TRUE),]
#     
#     # list the indices of columns containing the read counts
#     # exclude columns named "Mod", "Nuc", "AA", etc.
#     data_columns <- (1:NCOL(file_data))[!colnames(file_data) %in% c("SDB",
#                                                                     "Distance", "index", "mindex", "Primer", "Nuc", "Mod", "AA",
#                                                                     "Axis.name", "Order", "Barcode")]
#     
#     # choose the requested subset of the data columns
#     active_columns <- data_columns[as.vector(parsed_names$Columns[[i]])]
#     for(col in 1:length(active_columns)){
#       column_types <- c(column_types, as.character(parsed_names$Type[i]))
#       # new column name: filename root followed by the column number
#       column_name <- paste0(parsed_names$root[i], col)
#       colnames(file_data)[active_columns[col]] <- column_name
#       
#       # extract a data_frame with the barcode and one column
#       # of read counts, summing read counts over barcodes
#       data_column <- file_data %>% group_by(SDB, Barcode) %>%
#         summarize(count_sum = sum(!!as.name(column_name)))
#       
#       # fix the column name and merge it to the result frame
#       colnames(data_column)[3] <- column_name
#       merged_data <- full_join(merged_data, data_column, by = c("SDB", "Barcode"))
#       
#       # replace any NAs in the count data with zeros - restore NA SDBs
#       sdb_names <- merged_data$SDB
#       merged_data[is.na(merged_data)] <- 0
#       merged_data$SDB <- sdb_names
#     }
#   }
#   ordering_frame <- merged_data %>% left_join(dictionary, by = c("SDB", "Barcode" = "Axis.name"))
#   ordering <- order(ordering_frame$Order,			# primary sort by provided Order
#                     ordering_frame$Barcode == "",		# empty labels at the end
#                     sub("-\\[[0-9<]*]$", "",  ordering_frame$Barcode),   # break ties by glycan (alphabetic)
#                     suppressWarnings(as.integer(
#                       sub(".*-\\[<?0*([1-9][0-9]*)\\]$", "\\1", ordering_frame$Barcode))),
#                     ordering_frame$Barcode,			# finally just use the raw string
#                     nchar(ordering_frame$SDB),			# then use the SDB
#                     ordering_frame$SDB, na.last = TRUE)
#   merged_data <- merged_data[ordering,]
#   list(table = merged_data, types = column_types)
# }
# 
# 
# # routine to run the edgeR comparison process on data described in the campaign_dir
# for(i in 1:length(input_filename_list)){
#   # drop the extension (if any) from the filename to get the root used for naming new outputs
#   # note: special handling of indexing at length == 1 - does not work for all dot filenames
#   filename_root <- unlist(strsplit(basename(input_filename_list[i]), split = "[.]"))
#   filename_root <- paste(filename_root[1:length(filename_root)-1], collapse = ".")
#   filename_root
#   # read the campaign file
#   testing <- read.table(file.path(campaign_dir, input_filename_list[i]),
#                         sep = ",", quote = "\"", header = TRUE, stringsAsFactors = FALSE)
#   
#   testing
#   # check for the expected columns and skip any file without
#   if(length(unique(colnames(testing)[colnames(testing) %in%
#                                      c("Type", "Filename", "Columns", "Dictionary", "Labels", "OrderTable")])) == 6){
#     
#     print(paste0("Processing: ", filename_root))
#     testing$Type <- factor(testing$Type)
#     testing$Columns <- lapply(strsplit(testing$Columns, ","), as.integer)
#     
#     # R converts blank strings into NA on input so reverse this before extracting the list
#     testing$ExcludedSDBs[is.na(testing$ExcludedSDBs)] <- ""
#     testing$ExcludedSDBs <- lapply(strsplit(testing$ExcludedSDBs, ","), trimws)
#     
#     testing_data <- read_and_merge_columns(testing, LiGA_dir = LiGA.data.dir,
#                                            dict_dir = dict_dir, order_table_dir = order_table_dir)
#     
#     # Jessica's DE script does not allow NA row entries so delete them now
#     # similarly only test and control columns are allowed
#     reduced_types <- testing_data$types[-2]
#     subset_indices <- reduced_types %in% c("Barcode", "Test", "Control")
#     test_columns <- (0:(length(reduced_types[subset_indices])-1))[reduced_types[subset_indices] == "Test"]
#     control_columns <- (0:(length(reduced_types[subset_indices])-1))[reduced_types[subset_indices] == "Control"]
#     
#     # label (first) column must be named sequence
#     subset_table <- testing_data$table[-2]
#     colnames(subset_table)[1] <- "sequence"
#     subset_table$sequence <- gsub("[ \t]", ".",
#                                   paste0(testing_data$table$Barcode, ".", testing_data$table$SDB))
#     
#     # consider different subsets of the known SDBs
#     for(case in 3:3){
#       if(case == 1){
#         analysis_root = paste0(filename_root, "-all")
#       } 
#       if(case == 2){
#         analysis_root = paste0(filename_root, "-knownSDBs")
#         subset_table <- subset_table[subset_table$sequence != ".NA",]
#       } 
#       if(case == 3){
#         analysis_root = paste0(filename_root, "-knownGlycans")
#         subset_table <- subset_table[subset_table != paste0(".", testing_data$table$SDB),]
#       }
#       data_table_filename <- file.path(".", paste0(analysis_root, ".tsv"))
#       write.table(subset_table[!is.na(subset_table$sequence),subset_indices],
#                   file = data_table_filename, sep = "\t", quote = FALSE, row.names = FALSE)
#       
#       # # use edgeR, apply TMM and calculate the mean & st. dev. for each condition
#       saveName = paste0("compare_", analysis_root, ".txt")
#       htmlName = paste0("compare_", analysis_root, ".html")
#       
#       # test and control seem backwards here, but this is to label the final outputs correctly
#       render(file.path(scriptDir, "auxiliary/DE.rmd"),
#              params = list("TableName" = file.path("..", data_table_filename),
#                            "ControlCols" = paste(control_columns, collapse = ","),
#                            "TestCols" = paste(test_columns, collapse = ","),
#                            "saveName" = saveName))
#       file.remove(data_table_filename)
#       
#       print("Warning message:")
#       file.copy(file.path(scriptDir,"auxiliary", saveName), scriptDir, overwrite = TRUE)
#       file.remove(file.path(scriptDir,"auxiliary", saveName))
#       #file.copy(file.path(scriptDir,"auxiliary/DE.html"), scriptDir, overwrite = TRUE)
#       #file.remove(file.path(scriptDir,"auxiliary/DE.html"))
#       tmp = try(file.rename("auxiliary/DE.html", htmlName), TRUE)
#       if(inherits(tmp, "try-error")){
#         log = paste0(log, "\nWarning: DE in ", scriptDir, " failed!")
#       }
#     }
#   }
# }
```
