## Supplementary material for "Measuring carbohydrate recognition profile of lectins on live cells using liquid glycan array (LiGA)": Zipped folder containing all the software, MALDI TOF spectra and analyzed files: EqualMonomers.m

```
function [I, REASON] = EqualMonomers(obj, val)
%FIND   Calculate the molecular weight of the glycan object
%   I = MW(obj) returns the molecular weight as floating point number

glycans1 = obj.glycans;

if isa(val, 'GlycanLeaf')
    glycans2 = val.glycans;
else
    I = nan;
    REASON = 'passed arguments are not glycan objects';
    A = whos('obj');
    B = whos('val');
    warning(['Function is comparing ' A.class ...
             ' class of inputs to '  B.class '!!']);
    return
end

if size(glycans1) ~= size(glycans2)
    I = 0;
    REASON = 'different number of monomers';
    return;
end

if numel(glycans1)==0 && numel(glycans2)==0
    I = nan;
    REASON = 'no recognizable monomers in either glycan';
    return;
end

[S1, IX1] = sort(glycans1);
[S2, IX2] = sort(glycans2);

for i = 1:numel(S1)   
    test(i) = ~strcmp(S1{i}, S2{i});
end

if sum(test) == 0
    I = 1;
    REASON = ['monomers ' num2str(IX1) ' in glycan 1 are equal to '...
              char(10)...
              'monomers ' num2str(IX2) ' in glycan 2 '];
else
    I = 0;
    REASON = ['monomer(s) ' num2str(IX1(test)) ' in glycan 1 are not equal to '...
              char(10)...
              'monomer(s) ' num2str(IX2(test)) ' in glycan 2 '];
end
    

end
```
