## Supplementary material for "Measuring carbohydrate recognition profile of lectins on live cells using liquid glycan array (LiGA)": Zipped folder containing all the software, MALDI TOF spectra and analyzed files: GlycanLeaf.m

```
    classdef GlycanLeaf
    
properties
    % the only property of glycan that matters at construction
    % is the name of the glycan in a string. The rest, like coordinates
    % will be requested upon drawing, if drawing is called. Things like
    % topology, MW, sequence is calculated from the name.
    % Things like linkages and anomers are dispensable in most cases.
    % Even the name, technically, can be built after construction, but
    % there has to be some indication that it wasn't the proper name passed
    % and it needs to be created later.
      String
      
      origin      % the X,Y origin of the leaf, if not defined it is 0,0 
                  % it can be passed upon construction or in obj.draw
      angle       % ange at which the leaf is drawn, passed in obj.draw
      
      newGlycans
      newLinkages
      spacing

end
   
   properties (Constant)
      
      DISPLAY = 0;
      
        temp1 = {'sp', 'Glc', 'GalNAc', 'Neu5Ac', 'Fuc'};
        temp2 = {'1-2', '', '', '', '', '', ''};
        temp3 = {'a', 'b',   'a',   'a', 'a'};
        
        excel = 'monomers.xls';
        MWcol = 5;
        Monomercol = 1;

        % definitions of shapes for each monosacharide
        CFGnotation = {
            'Man',      'circle',   'green',    '';
            'Glc',      'circle',   'blue',     '';
            'Glu',      'circle',   'blue',     '';
            'Gal',      'circle',   'yellow',   '';
            'Rha',      'triangle', 'grey',     '';
            'Galf',     'circle',   'yellow',   'f';
            'GlcNAc',   'square',   'blue',     '';
            'GlcN(Gc)', 'square',   'turquoise','';
            'ManNAc',   'square',   'green',    '';
            'GalNAc',   'square',   'yellow',    '';
            'Fuc',      'triangle', 'red',      '';
            'Sp',       'circle',   'grey',     'Sp';
            'Thr',      'circle',	'grey',     'T';
            'Neu5Ac',   'diamond',  'magenta',  '';
            'KDN'       'diamond',  'grey',     '';
            'Neu5Gc',   'diamond',  'turquoise','';
            'Neu5,9Ac2', 'diamond',  'magenta',  '9Ac';
            'unkonwn',  'circle',   'white',    '?';
            'Sp0',      'circle',   'white',    'sp0';
            'Sp1',      'circle',   'white',    'sp1';
            'Sp8',      'circle',   'white',    'sp8';
            'Sp9',      'circle',   'white',    'sp8';
            'Sp10',     'circle',   'white',    'sp10';
            'Sp11',     'circle',   'white',    'sp11';
            'Sp12',     'circle',   'white',    'sp12';
            'Sp13',     'circle',   'white',    'sp13';
            'Sp14',     'circle',   'white',    'sp14';
            'Sp15',     'circle',   'white',    'sp15';
            'Sp19',     'circle',   'white',    'sp19';
            'Sp20',     'circle',   'white',    'sp20';
            'Sp21',     'circle',   'white',    'sp21';
            'Sp22',     'circle',   'white',    'sp22';
            'Sp23',     'circle',   'white',    'sp23';
            'Sp24',     'circle',   'white',    'sp24';
            'Sp25',     'circle',   'white',    'sp25';
            'p4',       'circle',   'white',    'P4';
            'P4',       'circle',   'white',    'P4';
            '3S',       'circle',   'white',    '3S';
            '6S',       'circle',   'white',    '6S';
            '6P',       'circle',   'white',    '6P';
            'S8',       'circle',   'white',    'S8';
            'S6',       'circle',   'white',    'S8';
            'S0',       'circle',   'white',    'S0';
            'Ss1',      'circle',   'white',    'Ss1';
            'DBCO',     'octagon',  'grey',     'DBCO';
        };
    
        % list of all the wrong names observed in glycans and ways to
        % resolve them. This list will just get bigger and bigger. 
%         replacement = {
%         
%             'Lac',      'Galb1-4Glc';
%             'LN',       'Galb1-4GlcNAc';
%             'NeuAc',    'Neu5Ac';
%             'NeuAc(9Ac)', 'Neu5,9Ac';
%             'Neu5Ac(9Ac)','Neu5,9Ac';
%             'sp',       'Sp';
%             'GlcNac',   'GlcNAc';
%             };
%         
        replacement = {
            'Lac',              'Lac',              'Galb1-4Glc';
            'LN',               'LN',               'Galb1-4GlcNAc';
            'NeuAc',            'NeuAc',            'Neu5Ac';
            'NeuAc(9Ac)',      'NeuAc(9Ac)a',      'Neu5,9Ac2a';
            'Neu5Ac(9Ac)a',     'Neu5Ac\(9Ac\)a',   'Neu5,9Ac2a';
            'Neu5Aca2,8',       'Neu5Aca2,8',       'Neu5Aca2-8';
            'Neu5Aca(2-3)',     'Neu5Aca\(2-3\)',   'Neu5Aca2-3';
            'GlcNac',           'GlcNac',           'GlcNAc';
            'sp',               'sp',               'Sp';
            };

        % color definitions
        COLORS = {
            'red',          [1 0 0];
            'green',        [0 1 0];
            'blue',         [0 0 1];
            'yellow',       [1 1 0];
            'grey',         [1 1 1]*0.8;
            'magenta',      [1 0 1];
            'turquoise',    [0 0.3 0];
            'white',        [1 1 1];
            };
   end
   
   properties (Dependent)
      Area
      
        glycans     % name of glycans defined from obj.string
        linkage     % linkage of glycans defined from obj.string
        anomers     % anomers of glycans defined from obj.string
        color       % color of each shape defined via obj.CFGnotation
        RGB         % RGB gamma of each shape defined via obj.COLORS 
        shape       % shape of glycan defined via obj.CFGnotation
        txt         % text written inside the shape

        % The angle is with + direction of
        % the X-axis, i.e., 0 means leaf is growin in the same direction as X-axis

        % topology of the leaf; either linear 0 or angular, where angles at each
        % joint point are defined as local rotation
        topology    % defined from obj.string
        
        % coordinates of each joint point of the leaf. This part needs to be
        % completed base on predefined topology of the leaf and angles at joints if
        % any. This variable can be passed out to the next leaf contructor to builf
        % leafs from the joint points outwards. 
        XY
   end
   
   methods    
      function obj = GlycanLeaf(name)
         % this is constructor where all dependent properties are set
         if obj.DISPLAY 
            disp('im inside constructor');
         end 
         
         if nargin > 0
            obj.String = name;
            obj.origin = [0,0];
            obj.angle  = 0;
            
            % clear the constant property at construction. It can be used
            % later to pass glycan structure from another object as 
            % sugar2.glycans = sugar1.glycans(end-1:end)
            obj.newGlycans = {};
         end
         
         if obj.DISPLAY 
            disp('glycan object is constructed');
         end
      end
      
      function [val, link] = DecodeName(obj)
          
        string = obj.String;
        
        %OligomerResolver(input, 1)
        string = OligomerResolver(string, 1);
          
        mono = struct;
        separator = '-[0-9][a-b]';
        string=[ deblank(string) '-'];

        %create regexp token expressions from CFG monomer names
        type{1} = '[ab]\d{1}-\d{1}';
        type{2} = '[ab]\(\d{1}-\d{1}\)';
        type{3} = '[ab]-';
        type{4} = '\d{1}-\d{1}';
        type{5} = '\(\d{1}-\d{1}\)';
        type{6} = '\)';
        type{7} = '[ab]\d{1},\d{1}';
        type{7} = '[ab]';
        
        linkEXP = '(';
        for i=1:numel(type)
            linkEXP = [linkEXP type{i} '|'];
        end
        linkEXP = [linkEXP '-)'];
        
        for i =1:size(obj.CFGnotation,1)
            mono(i).name = obj.CFGnotation{i,1};
            mono(i).token = ['(?<' mono(i).name '>' mono(i).name linkEXP ')'];
            %mono(i).expression = mono(i).token;
            mono(i).expression = [mono(i).name linkEXP];
            %mono(i).expression = [ mono(i).name '[' separator ']' ];
        end
if strcmp(string,'(3S)')
    sdfsd
end
        % cycle through every CGF monomer and search for it in the name
        for i = 1:numel(mono)

            %mono(i).found = regexp(string,mono(i).token,'match');
            mono(i).found = regexp(string, mono(i).expression,'match');
            
            temp = regexp(string,mono(i).expression,'start');
            % if the monomer is found, record its location and linkage!
            
            if ~isempty(temp)

                mono(i).location = temp;
                mono(i).number = numel(temp);
            end
        end
        
          
        % if location is not defined then no sugars have been recognized
        % abort the program.
        if ~isfield(mono,'location')
            disp(['There are no recognizable monomers in ' string]);
            obj.glycans = {};
            val = {};
            link = {};
            return;
        end

        k=1;
        monosacharide = {};
        position = [];


        for i=1:numel(mono)
            if mono(i).location
                for j = 1:numel(mono(i).location)
                    position(k) = mono(i).location(j);
                    monosacharide{k} = char(mono(i).name);
                    
                    % find the glycan name itself in the found expression
                    % and mark its end. The linkage, is everything after
                    % that end. 
                    temp = regexp( mono(i).found{j}, mono(i).name, 'end');
                    glycanLinkage{k} = mono(i).found{j}(temp(1)+1:end);
                    k=k+1;
                end
            end
        end

    % extract all positions into numerical array

        [~,IX] = sort(position,'descend');

        for i=1:numel( IX )
            if obj.DISPLAY
                disp(['Monomer #' num2str(i)...
                      ' is ' monosacharide{ IX(i) } ]);
            end
            obj.newGlycans{i} = monosacharide{ IX(i) };  
            obj.newLinkages{i} = glycanLinkage{ IX(i) };  
        end

        val = obj.newGlycans;
        link = obj.newLinkages;
        obj.glycans = obj.newGlycans;
        obj.linkage = obj.newLinkages;
      end
      
      function val = get.glycans(obj)
          
         % this function has to resolve the name and extract glycans
         % they are returned as cell array of strings
         
         % if glycans were redefined after constructing the object, this
         % line of code will pass the new value through constant property
         if ~isempty(obj.newGlycans)
             val = obj.newGlycans;
             return
         end
         
         if isempty(obj.String) | strcmp(obj.String,'internal')
             %disp('name is blank, call the glycan builder app');
             % I guess it can also call for some prompt where glycans are
             % entered by the user after the fact
             val = obj.temp1;
         elseif strcmp ( obj.String, 'image')
             disp('image recognition from supplied image');
             % it can be image
             % recognition from shapes, yeah!
             val = obj.temp1;
         else
             val = obj.String;
         end
         
      end
      
      function val = get.linkage(obj) 
         % this function has to resolve the name and extract glycans
         % they are returned as cell array of strings
         
         % if glycans were redefined after constructing the object, this
         % line of code will pass the new value through constant property
         if ~isempty(obj.newLinkages)
             val = obj.newLinkages;
             return
         end
      
          temp = []; % assign blank for now until the function is ready

          if ~isempty(temp)
             obj.glycans = temp;
          else
             obj.glycans = [0,0];
          end
      end  
      
      function obj = set.glycans(obj, val)
          
          obj.newGlycans = val;
          
      end

      
      function obj = set.linkage(obj, val)
          
          obj.newLinkages = val;
          
      end

      function obj = get.anomers(obj) 
      % extract anomers from name, either a or b 
      % if not defined, set to ?
      
          temp = obj.String;
          temp = []; % assign blank for now until the function is ready
      
          if ~iesmpty(temp)
             obj.glycans = val;
          else
             obj.glycans = '?';
          end
      end
      
      function val = get.color(obj) 
      % color of each glycan, assuming glycan names are correct 
      
        tempGlycans = obj.glycans; % avoid repeating calling  
        val = cell(size(tempGlycans)); % preallocatio
      
         for i=1:numel(tempGlycans)

            IX = find ( strcmp( tempGlycans{i}, obj.CFGnotation(:, 1) ));
            if ~isempty(IX)
                val{i} = obj.CFGnotation{IX, 3};
            else
                if obj.DISPLA 
                    fprintf(['glycan ' num2str(i) ...
                             '(' char(tempGlycans{i})...
                             ') is  not recognized.']);
                end
                IX = find ( strcmp( 'unkonwn', obj.CFGnotation(:, 1) ));
                val{i} = obj.CFGnotation{IX, 3};
                
                if obj.DISPLAY 
                    disp([' Assigned ' char (val{i}) ' color']); 
                end
            end
         end
        
      end
      
      function val = get.RGB(obj) 
      % and its RGB gamma of each shape for drawing 
        
        tempColor = obj.color; % avoid repeating calling
        val = cell(size(tempColor)); % preallocation
        
          
        for i=1:numel(tempColor)

            IX = find ( strcmp( tempColor{i}, obj.COLORS(:,1) ));
            if ~isempty(IX)
                val{i} = obj.COLORS{IX, 2};
            else
                if obj.DISPLAY 
                    fprintf(['color ' num2str(i)...
                             '(' char(tempColor{i})...
                             ') is  not recognized.']);
                end
                IX = find ( strcmp( 'white', tempColor(:, 1) ));
                val{i} = obj.COLORS{IX, 2};
                if obj.DISPLAY 
                    disp([' Assigned ' char (val{i}) ' RGB gamma']);
                end
            end
        end
        
      end 
        
      function val = get.shape(obj)
          % shape of the glycan, assuming glycan names are correct
          
        tempGlycans = obj.glycans; % avoid repeating calling  
        val = cell(size(tempGlycans)); % preallocation
          
        for i=1:numel(tempGlycans)

            IX = find ( strcmp( tempGlycans{i}, obj.CFGnotation(:, 1) ));
            if ~isempty(IX)
                val{i} = obj.CFGnotation{IX, 2};
            else
                if obj.DISPLAY 
                    fprintf(['glycan ' num2str(i)...
                             '(' char(tempGlycans{i})...
                             ') is  not recognized.']);
                end
                IX = find ( strcmp( 'unkonwn', obj.CFGnotation(:, 1) ));
                val{i} = obj.CFGnotation{IX, 2};
                
                if obj.DISPLAY 
                    disp([' Assigned ' char (val{i}) ' shape']); 
                end
            end
        end
        
      end 
      
      function val = get.txt(obj)
          % shape of the glycan, assuming glycan names are correct
          
        tempGlycans = obj.glycans; % avoid repeating calling  
        val = cell(size(tempGlycans)); % preallocation
          
        for i=1:numel(tempGlycans)

            IX = find ( strcmp( tempGlycans{i}, obj.CFGnotation(:, 1) ));
            if ~isempty(IX)
                val{i} = obj.CFGnotation{IX, 4};
            else
                if obj.DISPLAY 
                    fprintf(['glycan ' num2str(i)...
                             '(' char(tempGlycans{i})...
                             ') is  not recognized.']);
                end
                IX = find ( strcmp( 'unkonwn', obj.CFGnotation(:, 1) ));
                val{i} = obj.CFGnotation{IX, 4};
                
                if obj.DISPLAY 
                    disp([' Assigned ' char (val{i}) ' txt']); 
                end
            end
        end
        
      end

      function val = get.XY(obj) 
        % coordinates of each joint point of the leaf. This part needs to be
        % completed based on predefined topology of the leaf and angles at joints if
        % any. This variable can be passed out to the next leaf contructor to builf
        % leafs from the joint points outwards.
        
        tempGlycans = obj.glycans; % avoid repeating calling  
        N = numel(tempGlycans);
        val = zeros(N,2); % preallocation
        
        for i=1:N
          val(i,1) = obj.origin(1)+obj.spacing*(i-1)*cos(obj.angle);
          val(i,2) = obj.origin(2)+obj.spacing*(i-1)*sin(obj.angle);
        end
          % calculation of the next joint based on geometry
      end
        
      function obj = set.angle(obj,val)
          
      % ange at which the leaf is drawn. The angle is with + direction of
      % the X-axis, i.e., 0 means leaf is in the same direction as X-axis
        
          if ~isempty(val)
              obj.angle = val;
          else
              obj.angle = 0;
          end
      end          
    
      function obj = set.origin(obj,val) 
          % the X,Y origin of the linear stretch of glycans
          if obj.DISPLAY 
             disp('im inside origin');
          end
          if ~isempty(val)
              obj.origin = val;
          else
              obj.origin = [0, 0];
          end
          if obj.DISPLAY 
            disp(['origin is ' char(obj.origin) ]);
          end
      end

      function obj = get.topology(obj) 
          
        % this function will return an error because its 
        % attempting to change a property!
        % WORK IN PROGRESS. DO NOT USE
        % topology of the leaf is a dependent value defined from
        % the name of the glycan. It is either linear 0 or angular,
        % where angles at each
        % joint point are defined as local rotation    
        
          for i = 1:nume(obj.glycans)
            obj.topology(i) = 0;
          end
      end
         
      function obj = set.String(obj,val)
         if ~ischar(val)
            error('must be a string of characters!')
         end
         obj.String = val;

      end
      
      function draw(obj, varargin)

            X0 = obj.origin(1);
            Y0 = obj.origin(2);
            
            w=0.7*obj.spacing;
            L=0;
            reshape = 0;
            HOLD = 1;
            
            % by default, do not skip the first glycan / linker
            SKIP1 = 0;
            
            if exist('varargin','var')
                M = length(varargin);
                if rem(M,2) ~= 0, error('Parameters/Values must come in pairs'); 
                end

                % read input variables
                for ni = 1:2:M
                    switch lower(varargin{ni})
                        case 'reshape',    reshape = varargin{ni+1};
                        case 'hold',       HOLD    = varargin{ni+1};
                        case 'skip1',      SKIP1   = varargin{ni+1};
    
                    end
                end
            end 

            figH = gcf; %figure(1); 
            
            
            H = 100;
            
            if reshape
                set(figH, 'name', obj.String, ...
                   'position', [200,200,H*numel(obj.glycans) H]);
            end
            
            if HOLD
                hold on;
            end
          
          for i=1:numel(obj.glycans)

              if obj.DISPLAY 
                disp(  [char(obj.shape{i}) '  segment #=' num2str(L) ] );
              end

            if i<numel(obj.glycans)
                ThickLine(obj, X0, Y0, L, obj.spacing*0.8, obj.angle);
            end
            
            if i==1 && SKIP1
                L=L+obj.spacing;
                continue
            end
            
            DrawShape(obj, i, X0, Y0, L)

%             switch char(obj.shape{i})
% 
%                 case 'circle',   DrawShape(obj,obj.txt{i}, 20, X0, Y0, L, w, obj.angle, obj.RGB{i})
%                 case 'square',   DrawShape(obj.txt{i}, 4,  X0, Y0, L, 1.3*w, obj.angle, obj.RGB{i})
%                 case 'diamond',  DrawShape(obj.txt{i}, -4, X0, Y0, L, 1.3*w, obj.angle, obj.RGB{i})
%                 case 'triangle', DrawShape(obj.txt{i}, 3,  X0, Y0, L, w, obj.angle, obj.RGB{i})
%                 case 'line',     DrawShape(obj.txt{i}, 3,  X0, Y0, L, w, obj.angle, obj.RGB{i})
%             end
            L=L+obj.spacing;
          end
          
      end
 
      function ThickLine(obj, X,Y,L,w, rotate)

        obj.glycans;
        % % calculate shift from origin accounting for rotation.

        moveX = cos(rotate)*(L+w/2);
        moveY = sin(rotate)*(L+w/2);

        % get coordinates of a a skinny rectangle
        N=20;
         t = (1/(2*N):1/N:1)'*2*pi;

         t = t([end,1,10,11]);

            trX = moveX+(w/2)*cos(t+rotate);
            trY = moveY+(w/2)*sin(t+rotate);

            fill(trX+X, trY+Y, [0 0 0]);


      end
      
      function DrawShape(obj, i, X, Y, L)
          
           COLOR  = obj.RGB{i};
           TXT    = obj.txt{i};
           rotate = obj.angle;
           
           switch char(obj.shape{i})

                case 'circle',   N=20;  w = obj.spacing * 0.7;
                case 'square',   N= 4;  w = obj.spacing;
                case 'diamond',  N=-4;  w = obj.spacing * 0.8;
                case 'triangle', N= 3;  w = obj.spacing;
                case 'line',     N= 4;  w = obj.spacing; %WRONGGG
                case 'octagon',  N= 8;  w = obj.spacing * 0.7;
            end
          
            moveX = cos(rotate)*L;
            moveY = sin(rotate)*L;

            if N>0
                t = (1/(2*N):1/N:1)'*2*pi;
                trX = moveX+(w/2)*cos(t+rotate);
                trY = moveY+(w/2)*sin(t+rotate);
            else
                N=-N;
                t = (1/(2*N):1/N:1)'*2*pi;
                trX = moveX+(w/2)*cos(t+rotate+pi/4);
                trY = moveY+(w/2)*sin(t+rotate+pi/4);
            end

                fill(trX+X, trY+Y, COLOR);
                %text(moveX, moveY, TXT)
                text(X+moveX, Y+moveY, TXT, 'HorizontalAlignment', 'center', ...
           'fontsize', 7, 'fontname', 'arial');
      end
      
      function disp(obj)
         disp(['Glycan with name: ', obj.String])
      end
      
      function CalculateOrigins(obj)
          
          % I'm not sure if I need it
      end
          
   end
   methods (Static)
      function leaf = createObj(name)

          if nargin == 0
            name = 'internal'; 
          end

        T = GlycanLeaf();
        oldname = name;
        
        for i = 1:size(T.replacement,1)

            [START,END] = regexp(name, T.replacement{i,2}, 'start','end');
            MATCH = regexp(name, T.replacement{i,2}, 'match');
            
            while numel(START)>0
                fprintf(['found problematic term ' T.replacement{i,1}...
                      ' in locaiton ' num2str(START(1)) ] );

                name = [name(1:START(1)-1)...
                        T.replacement{i,3}...
                        name(END(1)+1:end)];
                    
                disp([' and replaced with ' T.replacement{i,3}]);
                
                [START,END] = regexp(name, T.replacement{i,2},'start','end');
            end
%             if ~isempty(START)
% rr
%                 fprintf(['found problematic term ' T.replacement{i,1}...
%                       ' in locaitons ' num2str(START) ] );
%                 for j=1:numel(START)
%                     name = [name( 1 : START(j)-1) T.replacement{i,3}...
%                             name( START(j)+numel(T.replacement{i,1}) : end)];
%                 end
%                 disp([' and replaced with ' T.replacement{i,2}]);
%             end
        end
        
        clear T;
        
        leaf = GlycanLeaf(name);
        
        [t,l] = DecodeName(leaf);
        
        % set the spacinf between glycans to predefined value
        % this value can be changed later by user
        leaf.spacing = 1;
        %xy = CalculateOrigins(leaf);
        
        if ~isempty(t)
            leaf.glycans = t;
        else
            leaf.glycans = {'unkonwn'};
        end
        
        if ~isempty(l)
            leaf.linkage = l;
        else
            leaf.linkage = {'unkonwn'};
        end
        
            leaf.shape;
            leaf.color;
            leaf.RGB;

      end  
   end
    end
```
