## Supplementary material for "Measuring carbohydrate recognition profile of lectins on live cells using liquid glycan array (LiGA)": Zipped folder containing all the software, MALDI TOF spectra and analyzed files: MW.m

```
function [I, WARNING, mw] = MW(obj)
%FIND   Calculate the molecular weight of the glycan object
%   I = MW(obj) returns the molecular weight as floating point number
%   
%   [I, WARNING] = MW(obj) returns the molecular weight and the WARNING if
%   some glycans were not recognized
%
%   [I, ~, mw] = MW(obj) returns the molecular weight and molecular weights
%   of the individual monomers recognized in the glycan

    water = 18.02;

    val = obj.glycans;
    mw = zeros(size(val));
    WARNING = cell(size(val));
    
    [~,~,C] = xlsread(obj.excel);
    for i = 1: numel(val)
        ix = find ( strcmp(val{i}, C(:, obj.Monomercol)) );
        if ~isempty(ix)
            if numel(ix)>2
                % found two instances of the same name
                WARNING{i} = [num2str(numel(ix))...
                              ' redundant entries '...
                               C(ix(1), obj.Monomercol)...
                              ' are found in MW table.'];
                disp([ WARNING{i} 'Molecular weights are:']);
                
                for j = 1:numel(ix)
                    disp(num2str(C(ix(i), obj.MWcol)));
                end
                disp('the first value is used to calculate the MW');
                disp('final mass is possibly wrong');
                mw(i) = C(ix(1), obj.MWcol);

            else
                % only one istance exists; extract the mass
                mw(i) = C{ix(1), obj.MWcol};
                
                if mw(i)==0
                    WARNING{i} = ['Molecular Weight of '...
                                   C(ix(1), obj.Monomercol)...
                                   ' is set to 0 in the table'];
                else
                    WARNING{i} = 'ok';
                end
            end
        else
            WARNING{i} = 'could not find the monomer';
            % did not find the monomer
            mw(i) = nan;
        end
    end
    
    I = sum(mw( ~isnan(mw) )) - water * (numel( ~isnan(mw) ) - 1 );
end
```
