## Supplementary material for "Measuring carbohydrate recognition profile of lectins on live cells using liquid glycan array (LiGA)": Zipped folder containing all the software, MALDI TOF spectra and analyzed files: drawGlycan.m

```
function [branch] = drawGlycan(varargin)

TEXT = 0;
DISP = 0;
branch_rotation = pi/2;
spacing = 1;

input = 'Neu5Aca2-6Galb1-4GlcNAcb-Sp';
XY = [1,1];
angle = 0;

    if exist('varargin','var')
        L = length(varargin);
        if rem(L,2) ~= 0 
            error('Parameters/Values must come in pairs.'); 
        end

        % read input variables
        for ni = 1:2:L
            switch lower(varargin{ni})
                case 'input',          	input =     varargin{ni+1};
                case 'xy',              XY =        varargin{ni+1};
                case 'angle',           angle =     varargin{ni+1}; 
                case 'branch_rotation', branch_rotation = varargin{ni+1}; 
                case 'text',            TEXT =      varargin{ni+1};
                case 'disp',            DISP =      varargin{ni+1}; 
                case 'spacing',         spacing =   varargin{ni+1}; 
            end
        end
    end
% this script will fail with names that contain linkage in brackets
% Neu5Aca(2-3)(Galb(1-4)[Fuca(1-3)]GlcNAc)3b-Sp
% quick fix is do detect this pattern and replace it

badLinkage = '\(\d{1}-\d{1}\)';
[badS,badE] = regexp(input,badLinkage,'start','end');

while numel(badS)>0
    [badS,badE] = regexp(input,badLinkage,'start','end');
    if numel(badS)>0
        input = [ input(1:badS(1)-1) ...
                 input(badS(1)+1:badE(1)-1) ...
                 input(badE(1)+1:end)];
    end
end

% any oligomers will confuse the script and need to be resolved
input = OligomerResolver(input, 1);

% step 1, detect linear portion and branching pattern

[linear_string, branch_flag] = DetectLinearAndBranches(input, DISP);

% exist if there is a problem with brackets or branch detection
if branch_flag(end) > '0'
    disp('there is a problem with brackets and branches; skipping!');
    branch = struct;
    return
end

% create linear sugar object
linear_sugar = GlycanLeaf.createObj(linear_string);  

% set the origins and orientation of the linear glycan object
linear_sugar.origin = XY;
linear_sugar.angle = angle;
linear_sugar.spacing = spacing;


if DISP
    
    linearN = numel(linear_sugar.glycans);
    
    disp(['The number of glycans in linear portion is ' ...
          num2str(linearN) ]);
    fprintf('The glycans are : ');
    for i = 1:linearN
        fprintf(['-' linear_sugar.glycans{i}]);   
    end
    disp(char(10));
end

% % create an structure array of branches
if sum(find(branch_flag>'0'))>1
    branch = StructureArrayofBranches(input, linear_sugar, branch_flag);
else
    branch = [];
end

% go through every branch and either draw it if its linear or deconstruct
% it again if its further branched

for i = 1:numel(branch)

    % is branch composed of linear glycans or further branched?
    IX = find(branch(i).branch_flag > branch(i).branch_flag(1));
    
    if isempty(IX)
        
        % Branch is linear: create a glycan object and draw the branch
        if DISP
            disp(['Branch ' branch(i).glycan ' # ' num2str(i) ' is linear']);
        end
        object = GlycanLeaf.createObj( branch(i).branchANDcore );
        
        object.origin = linear_sugar.XY( branch(i).position_in_linear, :);
        object.angle =  linear_sugar.angle + branch_rotation;
        object.spacing = linear_sugar.spacing;


        draw(object, 'reshape', 0, 'skip1', 1);  % draw the branch
    else
        % deconstruct it further
        if DISP
            disp(['Branch ' branch(i).glycan ' # ' num2str(i) ' is branched']);
        end
        
        subglycan = branch(i).branchANDcore;
        
        % this is dynamic programming: reuse the function itself!!
        drawGlycan('input', subglycan,...
                   'XY', linear_sugar.XY(branch(i).position_in_linear,:),...
                   'angle',linear_sugar.angle + branch_rotation,...
                   'spacing', spacing,...
                   'branch_rotation', branch_rotation,...
                   'disp',DISP);

    end
    
end


% draw the linear portion last
draw(linear_sugar, 'reshape', 0); hold on;


if TEXT
    text(XY(1),XY(2), [linear_string '     '],...
           'HorizontalAlignment', 'right', ...
           'fontsize', 7, 'fontname', 'arial');
end

end

  
function [linear_string, branch_flag, MAX_branch] = ...
            DetectLinearAndBranches(input,DISP)


    % extract the linear portion of the glycan and convert it to an object
    branch = 0;
    branch_flag = '';
    linear_string = '';
    closeBracket = {};
    

    if DISP
        disp([char(10) char(10) char(10)]);
        disp([  'Input glycan is ' input ]);
        fprintf('Branch flag is  ');
    end

    for i= 1:numel(input)


        if (input(i) == '(' || input(i) ==  '[')

            % turn on the branch flag, or add a branching point
            branch = branch + 1;
            branch_flag = [branch_flag num2str(branch)];
            
            if DISP, fprintf(num2str(branch)); end

            % designate the close bracket ignore everything
            if input(i) == '('
                closeBracket{branch} = ')';
            elseif input(i) == '['
                closeBracket{branch} = ']';
            end

            continue
        end

        if branch 
            if input(i) ==  closeBracket{branch}

                % decrease the branching point flag 
                % if branch was = 1, it will turn off the branch flag
                branch_flag = [branch_flag num2str(branch)];
                
                if DISP, fprintf(num2str(branch)); end
                
                branch = branch - 1;
                continue
            else
                % if the symbol is not a close bracket, abort the cycle
                branch_flag = [branch_flag num2str(branch)];
                
                if DISP, fprintf(num2str(branch)); end
                
                continue
            end
        end

        % if cycle wasn't aborted by this point, the cycle is outside the
        % branching point and linear portion can be recorded

        branch_flag = [branch_flag num2str(branch)];
        if DISP
            fprintf(num2str(branch));
        end
        linear_string = [linear_string input(i)];

    end
    
    % find the maximum nested branching
    MAX_branch = 0;
    for i = 1:numel(branch_flag)
        temp = str2num(branch_flag(i));
        if temp>MAX_branch
            MAX_branch = temp;
        end
    end
    
    if DISP
        disp([char(10) 'Branch flag is  ' branch_flag]);
        disp([char(10) 'linear glycan is ' linear_string]);
    end


end

function [branch] = StructureArrayofBranches(input, linear_sugar, branch_flag)

    N = 0;
    linear = '';
    branch = struct;
    %branch(1).flag = 0;
    i=1;
    linearN = numel(linear_sugar.glycans);
    DEBUG = 0;

    while i<=numel(input)


        if str2num(branch_flag(i))>0
            if DEBUG
                disp(['detected a branching point #' branch_flag(i)]);
                disp(['linear glycan so far is ' linear]);
            end

            temp = GlycanLeaf.createObj(linear);

            tempN = linearN - numel(temp.glycans) ;
            if DEBUG
                disp(['The position of branching glycan is ' num2str(tempN) ]);
                disp(['The glycan from which branching occurs is ' temp.glycans{1} ]);
            end

            % mark the beginning of a branch
            N = N + 1;
            branch(N).flag = 1;
            branch(N).start = i;
            branch(N).glycan = '';
            branch(N).position_in_linear = tempN;
            branch(N).glycan_in_linear = [temp.glycans{1} temp.linkage{1}];
            branch(N).branch_flag = '';
            branch(N).position = numel(temp.glycans);

            while branch(N).flag

                if round(str2num(branch_flag(i))) == 0
                    branch(N).flag = 0;
                    branch(N).end = i;

                    if DEBUG
                        disp(['end of branch i=' num2str(i)...
                              ' input is ' input(i)])
                    end
                else

                    % mark the end of the branch
                    branch(N).glycan = [branch(N).glycan  input(i)];
                    branch(N).branch_flag = [ branch(N).branch_flag ...
                                                        branch_flag(i) ];
                    i=i+1;
                end
                
                branch(N).branchANDcore = [branch(N).glycan(2:end-1)...
                                           branch(N).glycan_in_linear];

            end

        else
            if DEBUG
                disp(['Im out i=' num2str(i) ' input is ' input(i)]);
            end
                linear = [linear input(i)];
                i=i+1;
        end

    end

end
```
