## Supplementary material for "Measuring carbohydrate recognition profile of lectins on live cells using liquid glycan array (LiGA)": Zipped folder containing all the software, MALDI TOF spectra and analyzed files: Maldi.pdf

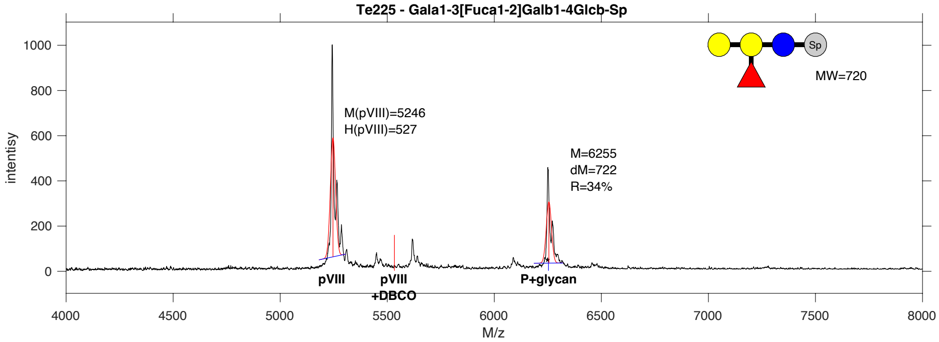


**1: MALDI-TOF spectra of glycosylated phage: B tetra L-[920]**

**SDB Number:** SDB30

**Sequencing File:** http://ligacloud.ca/searchLibInfo?f=0&b=0&d=NA

**Barcode:**
CTGCTTTTTGCGATTCCCCTTAGTGTGGAGAAGAATGATCAGAAGACTTAT
CATGCGGGTGGAGGT

**Axis Name:** B tetra L-[920]

**IUPAC:** Gal(b1-4)Glc(b1-Sp

**IUPAC (CFG):** Gala1-3[Fuca1-2]Galb1-4Glcb-Sp

**Common Name:** B tetra L

**Glytoucan ID:** NA

**Compound Number:** Te225

**Maldi File:** 386.txt

**Density:** 34

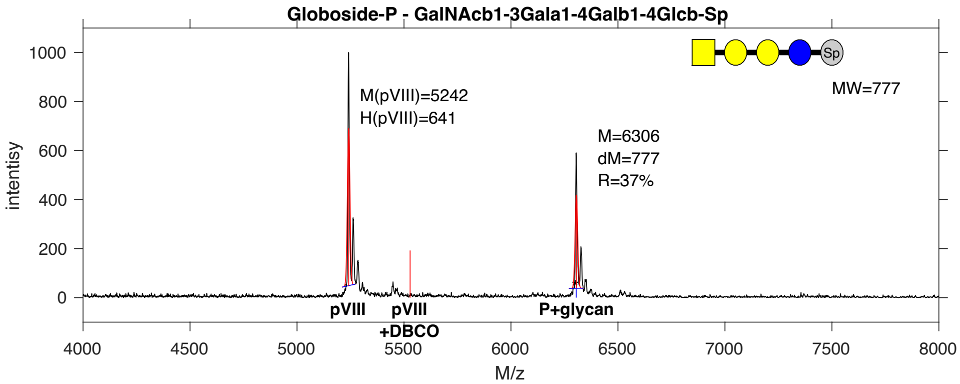


**2: MALDI-TOF spectra of glycosylated phage: Globoside-P-[1030]**

**SDB Number:** SDB33

**Sequencing File:** http://ligacloud.ca/searchLibInfo?f=0&b=0&d=NA

**Barcode:**
CTGCTTTTTGCCATACCGCTGAGTGTGGAGAAGAATGATCAGAAGACTTA
TCATGCGGGTGGAGGT

**Axis Name:** Globoside-P-[1030]

**IUPAC:** GalNAc(b1-3)Gal(a1-4)Gal(b1-4)Glc(b1-Sp

**IUPAC (CFG):** GalNAcb1-3Gala1-4Galb1-4Glcb-Sp

**Common Name:** Globoside-P

**Glytoucan ID:** G24499XE

**Compound Number:** Te272

**Maldi File:** 387.txt

**Density:** 38

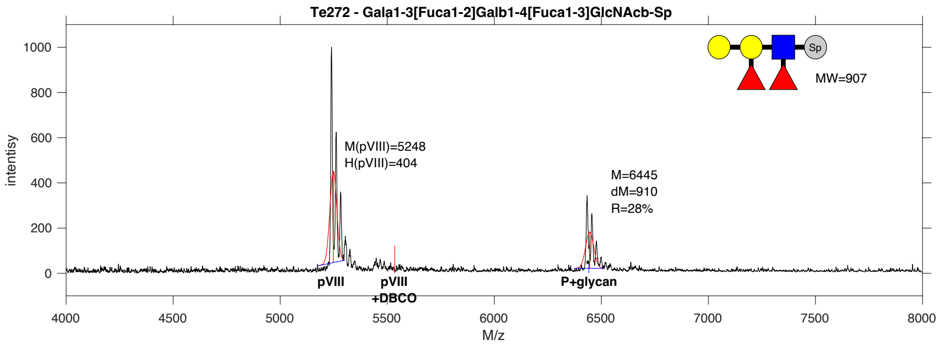


**3: MALDI-TOF spectra of glycosylated phage: 2’F-B type 2-[760]**

**SDB Number:** SDB60

**Sequencing File:** http://ligacloud.ca/searchLibInfo?f=0&b=0&d=20210330-87CLooDK-MS

**Barcode:**
CTACTGTTCGCCATCCCTCTGAGTGTAGAAAAGAACGACCAAAAGACTTACCATGCTGGTGGGGGG

**Axis Name:** 2’F-B type 2-[760]

**IUPAC:** Gal(a1-3)[Fuc(a1-2)]Gal(b1-4)[Fuc(a1-3)]GlcNAc(b1-Sp

**IUPAC (CFG):** Gala1-3[Fuca1-2]Galb1-4[Fuca1-3]GlcNAcb-Sp

**Common Name:** 2’F-B type 2

**Glytoucan ID:** G04624DG

**Compound Number:** Te262

**Maldi File:** 388.txt

**Density:** 28

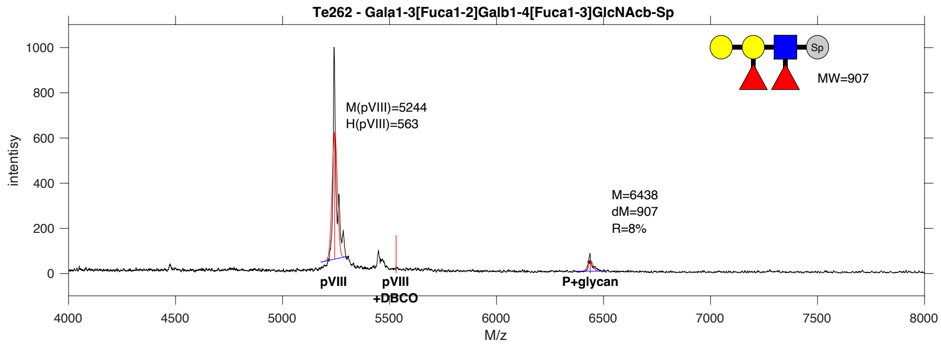


#### 4: MALDI-TOF spectra of glycosylated phage: 2’F-B type 2-[220]

**SDB Number:** SDB72

**Sequencing File:** http://ligacloud.ca/searchLibInfo?f=0&b=0&d=NA

**Barcode:**
TTATTATTCGCAATTCCTCTAAGCGTCGAGAAGAACGATCAAAAAACTTA
TCACGCGGGAGGTGGA

**Axis Name:** 2’F-B type 2-[220]

**IUPAC:** Gal(a1-3)[Fuc(a1-2)]Gal(b1-4)[Fuc(a1-3)]GlcNAc(b1-Sp

**IUPAC (CFG):** Gala1-3[Fuca1-2]Galb1-4[Fuca1-3]GlcNAcb-Sp

**Common Name:** 2’F-B type 2

**Glytoucan ID:** G04624DG

**Compound Number:** Te262

**Maldi File:** 389.txt

**Density:** 8

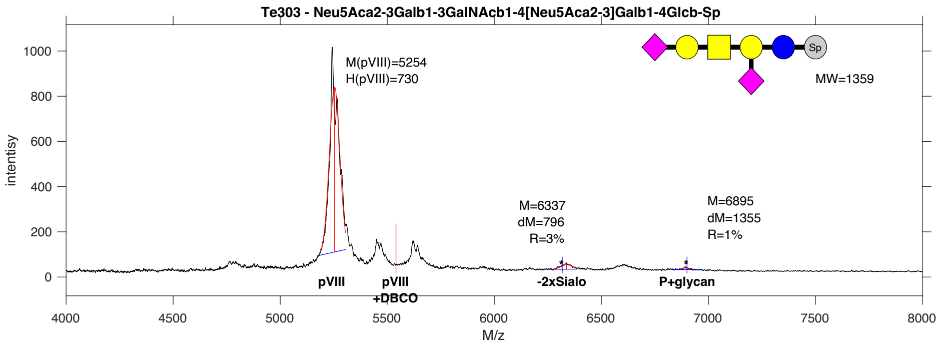


#### 5: MALDI-TOF spectra of glycosylated phage: GD1a-[110]

**SDB Number:** SDB31

**Sequencing File:** http://ligacloud.ca/searchLibInfo?f=0&b=0&d=20210319-87CLooVF-MS

**Barcode:** CTGCTTTTCGCGATACCACTGAGTGTGGAGAAGAATGATCAGAAGACTTATCATGCGGGTGGAGGT

**Axis Name:** GD1a-[110]

**IUPAC:** Neu5Ac(a2-3)[Neu5Ac(a2-3)Gal(b1-3)GalNAc(b1-4)]Gal(b1-4)Glc(b1-Sp

**IUPAC (CFG):** Neu5Aca2-3[Neu5Aca2-3Galb1-3GalNAcb1-4]Galb1-4Glcb-Sp

**Common Name:** GD1a

**Glytoucan ID:** G21404QI

**Compound Number:** Te303

**Maldi File:** 400.txt

**Density:** 4

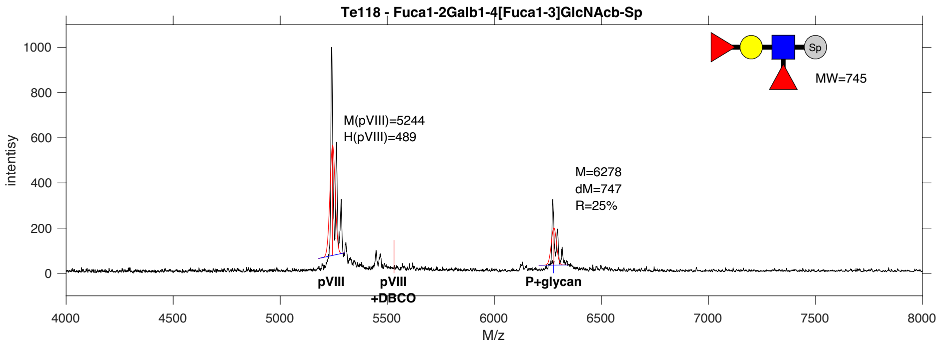


**6: MALDI-TOF spectra of glycosylated phage: 3’GN-Di-LN-[1110]**

**SDB Number:** SDB46

**Sequencing File:** http://ligacloud.ca/searchLibInfo?f=0&b=0&d=NA

**Barcode:**
CTGCTATTTGCCATCCCGCTGAGCGTTGAAAAAAACGACCAGAAAACGTA
TCACGCGGGGGGGGGG

**Axis Name:** 3’GN-Di-LN-[1110]

**IUPAC:** GlcNAc(b1-3)Gal(b1-4)GlcNAc(b1-3)Gal(b1-4)GlcNAc(b1-Sp

**IUPAC (CFG):** GlcNAcb1-3(Galb1-4GlcNAcb1-3)2b-Sp

**Common Name:** 3’GN-Di-LN

**Glytoucan ID:** G73480MV

**Compound Number:** Te99

**Maldi File:** 401.txt

**Density:** 41

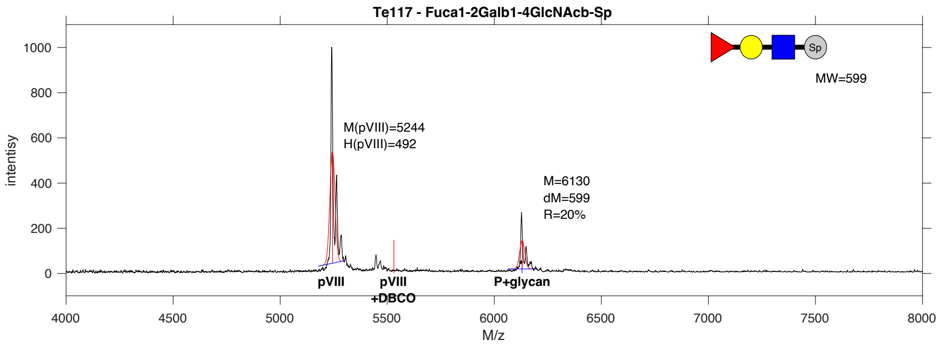


**7: MALDI-TOF spectra of glycosylated phage: H-type 2-[540]**

**SDB Number:** SDB64

**Sequencing File:** http://ligacloud.ca/searchLibInfo?f=0&b=0&d=NA

**Barcode:**
CTGCTCTTTGCTATACCCCTAAGCGTCGAGAAGAACGACCAGAAGACCTA
CCACGCTGGAGGGGGA

**Axis Name:** H-type 2-[540]

**IUPAC:** Fuc(a1-2)Gal(b1-4)GlcNAc(b1-Sp

**IUPAC (CFG):** Fuca1-2Galb1-4GlcNAcb-Sp

**Common Name:** H-type 2

**Glytoucan ID:** G17510DO

**Compound Number:** Tr117

**Maldi File:** 402.txt

**Density:** 20

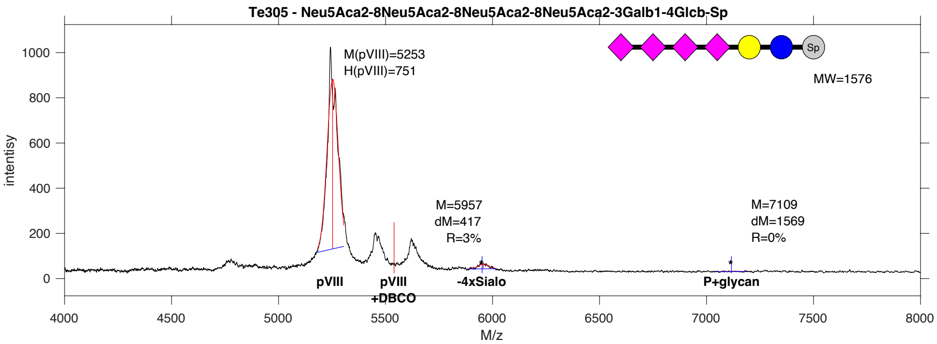


#### 8: MALDI-TOF spectra of glycosylated phage: TetraSLac-[80]

**SDB Number:** SDB78

**Sequencing File:** http://ligacloud.ca/searchLibInfo?f=0&b=0&d=20210319-87CLooZH-MS

**Barcode:** CTGCTGTTCGCGATTCCCCTAAGTGTCGAGAAAAATGATCAAAAAACTTATCATGCGGGAGGCGGG

**Axis Name:** TetraSLac-[80]

**IUPAC:** Neu5Ac(a2-8)Neu5Ac(a2-8)Neu5Ac(a2-8)Neu5Ac(a2-3)Gal(b1-4)Glc(b1-Sp

**IUPAC (CFG):** Neu5Aca2-8Neu5Aca2-8Neu5Aca2-8Neu5Aca2-3Galb1-4Glcb-Sp

**Common Name:** TetraSLac

**Glytoucan ID:** Na

**Compound Number:** Te305

**Maldi File:** 404.txt

**Density:** 3

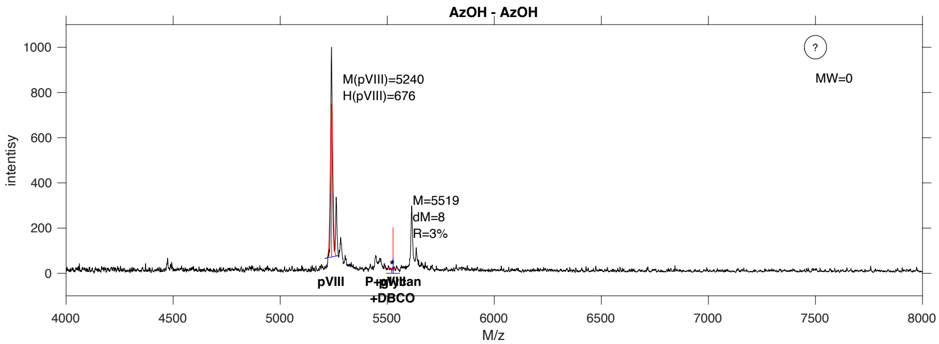


**9: MALDI-TOF spectra of glycosylated phage: AzOH-[460]**

**SDB Number:** SDB85

**Sequencing File:** http://ligacloud.ca/searchLibInfo?f=0&b=0&d=20210319-87CLooKN-MS

**Barcode:** TTATTATTCGCAATTCCTTTAAGTGTGGAGAAGAACGACCAGAAAACGTATCACGCAGGTGGGGGT

**Axis Name:** AzOH-[460]

**IUPAC:** NASp

**IUPAC (CFG):** AzOH

**Common Name:** AzOH

**Glytoucan ID:** Not Registered

**Compound Number:** AzOH

**Maldi File:** 409.txt

**Density:** 17

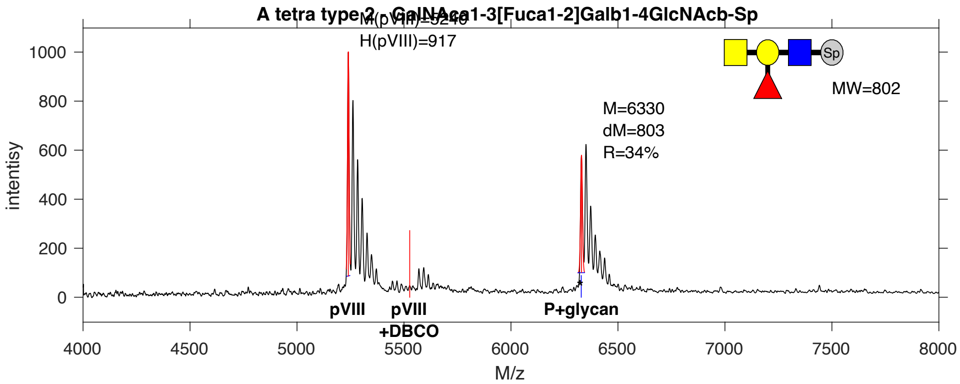


**10: MALDI-TOF spectra of glycosylated phage: A tetra type 2-[920]**

**SDB Number:** SDB134

**Sequencing File:** http://ligacloud.ca/searchLibInfo?f=0&b=0&d=NA

**Barcode:**
CTGCTTTTTGCTATACCCCTGAGTGTAGAGAAGAATGACCAAAAAACTTACCACGCTGGTGGGGGC

**Axis Name:** A tetra type 2-[920]

**IUPAC:** Gal(a1-3)Gal(b1-4)[Fuc(a1-3)]GlcNAc(b1-Sp

**IUPAC (CFG):** GalNAca1-3[Fuca1-2]Galb1-4GlcNAcb-Sp

**Common Name:** A tetra type 2

**Glytoucan ID:** G86393KI

**Compound Number:** Te222

**Maldi File:** RR-II-79.txt

**Density:** 34

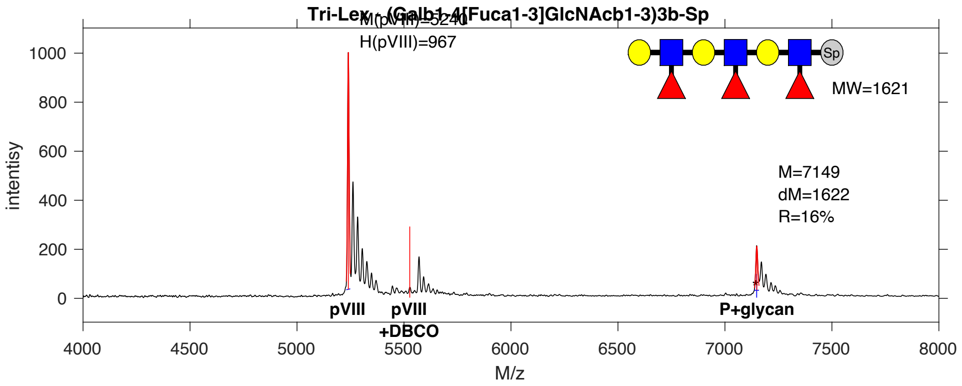


**11: MALDI-TOF spectra of glycosylated phage: Tri-Lex-[430]**

**SDB Number:** SDB135

**Sequencing File:** http://ligacloud.ca/searchLibInfo?f=0&b=0&d=NA

**Barcode:**
CTTCTGTTTGCTATACCTCTAAGTGTGGAGAAGAATGACCAAAAGACATA
TCACGCAGGTGGGGGA

**Axis Name:** Tri-Lex-[430]

**IUPAC:** Gal(b1-4)[Fuc(a1-3)]GlcNAc(b1-3)Gal(b1-4)[Fuc(a1-3)]GlcNAc(b1-3)Gal(b1-4)[Fuc(a1-3)]GlcNAc(b1-Sp

**IUPAC (CFG):** (Galb1-4[Fuca1-3]GlcNAcb1-3)3b-Sp

**Common Name:** Tri-Lex

**Glytoucan ID:** G57884AX

**Compound Number:** Te102

**Maldi File:** MS-XIV-62.txt

**Density:** 16

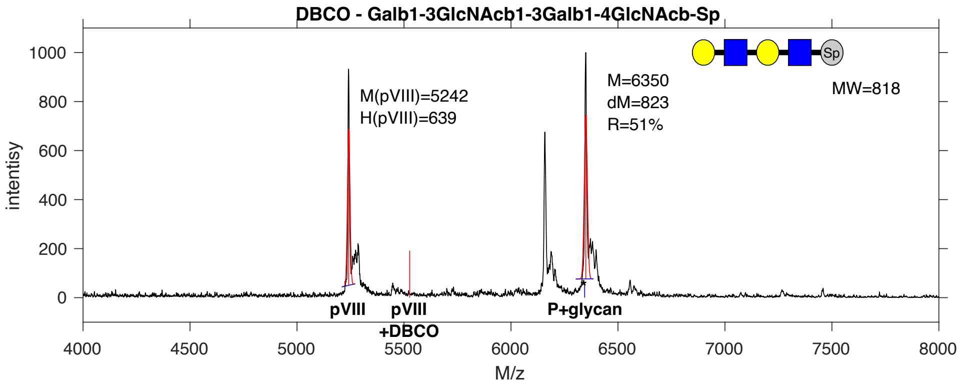


**12: MALDI-TOF spectra of glycosylated phage: (Galf)4-[1380]**

**SDB Number:** SDB138

**Sequencing File:** http://ligacloud.ca/searchLibInfo?f=0&b=0&d=NA

**Barcode:**
CTACTATTCGCTATACCCCTAAGCGTGGAGAAAAATGACCAGAAGACGTA
TCACGCGGGGGGGGGG

**Axis Name:** (Galf)4-[1380]

**IUPAC:** Galf(b1-5)Galf(b1-5)Galf(b1-5)Galf(b1-S8

**IUPAC (CFG):** Galfb1-5Galfb1-5Galfb1-5Galfb-S8

**Common Name:** (Galf)4

**Glytoucan ID:** G92890KE

**Compound Number:** Galf4

**Maldi File:** RR-II-111.txt

**Density:** 51

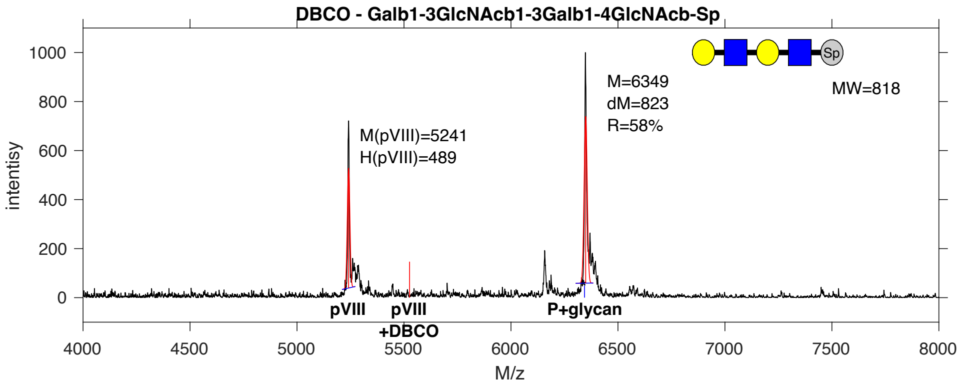


**13: MALDI-TOF spectra of glycosylated phage: (Galf)4-[1570]**

**SDB Number:** SDB138

**Sequencing File:** http://ligacloud.ca/searchLibInfo?f=0&b=0&d=NA

**Barcode:**
CTACTATTCGCTATACCCCTAAGCGTGGAGAAAAATGACCAGAAGACGTA
TCACGCGGGGGGGGGG

**Axis Name:** (Galf)4-[1570]

**IUPAC:** Galf(b1-5)Galf(b1-5)Galf(b1-5)Galf(b1-S8

**IUPAC (CFG):** Galfb1-5Galfb1-5Galfb1-5Galfb-S8

**Common Name:** (Galf)4

**Glytoucan ID:** G92890KE

**Compound Number:** Galf4

**Maldi File:** RR-II-112.txt

**Density:** 58

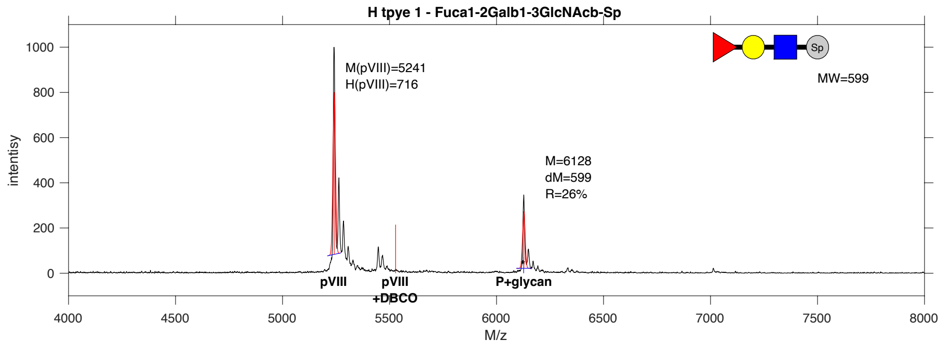


#### 14: MALDI-TOF spectra of glycosylated phage: H type 1-[700]

**SDB Number:** SDB146

**Sequencing File:** http://ligacloud.ca/searchLibInfo?f=0&b=0&d=20210319-87CLooMJ-MS

**Barcode:**
CTGCTGTTTGCTATACCCCTTAGTGTCGAGAAAAATGACCAAAAGACGTACCATGCGGGAGGAGGG

**Axis Name:** H type 1-[700]

**IUPAC:** Fuc(a1-2)Gal(b1-3)GlcNAc(b1-Sp

**IUPAC (CFG):** Fuca1-2Galb1-3GlcNAcb-Sp

**Common Name:** H type 1

**Glytoucan ID:** NA

**Compound Number:** Tr116

**Maldi File:** RR-III-84.txt

**Density:** 26

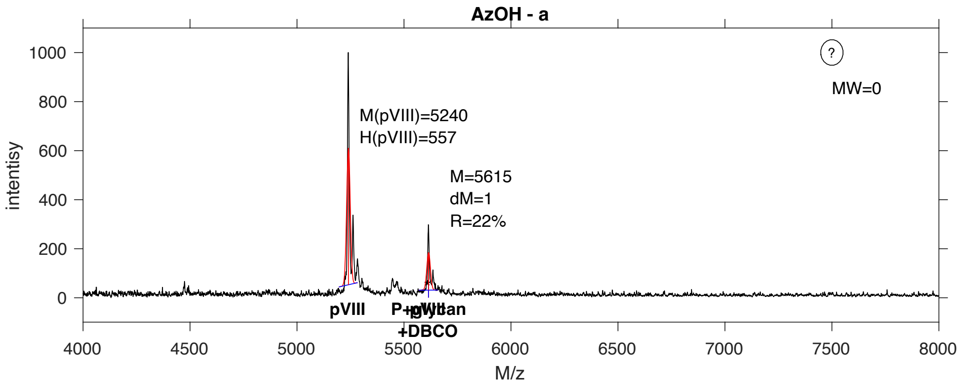


**15: MALDI-TOF spectra of glycosylated phage: AzOH-[950]**

**SDB Number:** SDB66

**Sequencing File:** http://ligacloud.ca/searchLibInfo?f=0&b=0&d=NA

**Barcode:**
TTATTATTCGCAATTCCTTTAAGTGTCGAGAAGAACGATCAGAAGACCTA
CCACGCAGGGGGGGGT

**Axis Name:** AzOH-[950]

**IUPAC:** NASp

**IUPAC (CFG):** AzOH

**Common Name:** AzOH

**Glytoucan ID:** Not Registered

**Compound Number:** AzOH

**Maldi File:** MS-XIV-173.txt

**Density:** 35

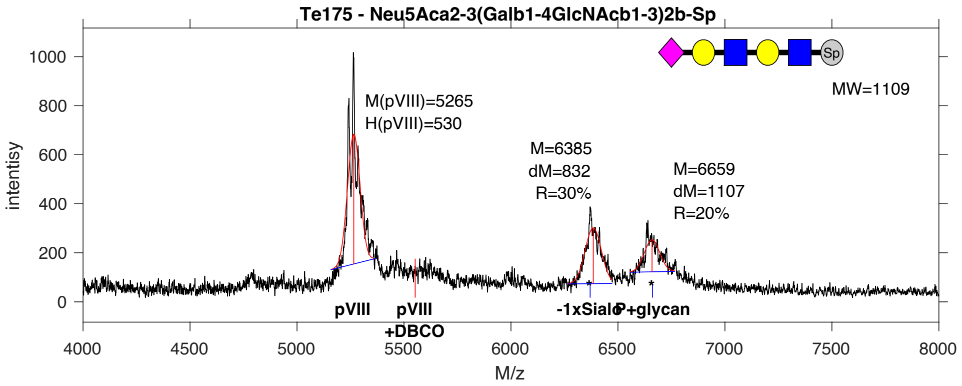


**16: MALDI-TOF spectra of glycosylated phage: 3’S-Di-LN-[1350]**

**SDB Number:** SDB198

**Sequencing File:** http://ligacloud.ca/searchLibInfo?f=0&b=0&d=NA

**Barcode:**
CTGCTCTTTGCTATACCCCTAAGTGTGGAGAAAAATGACCAAAAAACTTATCACGCAGGAGGTGGT

**Axis Name:** 3’S-Di-LN-[1350]

**IUPAC:** Neu5Ac(a2-3)Gal(b1-4)GlcNAc(b1-3)Gal(b1-4)GlcNAc(b1-Sp

**IUPAC (CFG):** Neu5Aca2-3(Galb1-4GlcNAcb1-3)2b-Sp

**Common Name:** 3’S-Di-LN

**Glytoucan ID:** G31119YY

**Compound Number:** Te175

**Maldi File:** MS-XIV-40i.txt

**Density:** 50

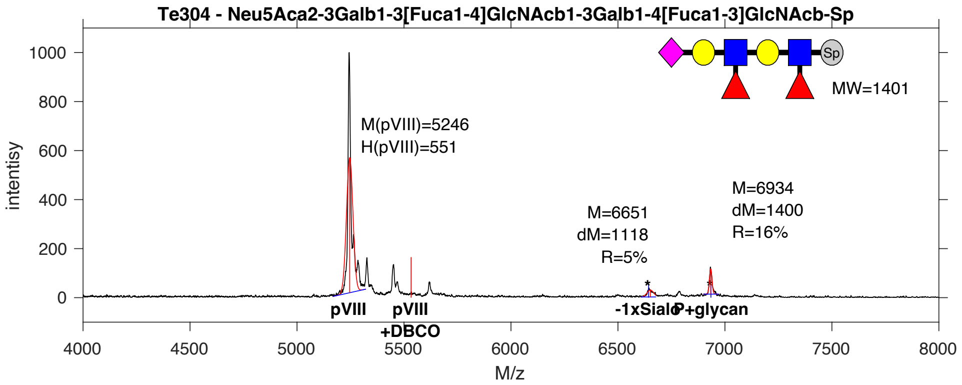


**17: MALDI-TOF spectra of glycosylated phage: 3'SLeA-Lex-[570]**

**SDB Number:** SDB96

**Sequencing File:** http://ligacloud.ca/searchLibInfo?f=0&b=0&d=20210319-87CLooOI-MS

**Barcode:** CTGCTGTTCGCAATCCCGCTAAGTGTTGAAAAAAATGATCAGAAAACATATCACGCTGGGGGTGGA

**Axis Name:** 3'SLeA-Lex-[570]

**IUPAC:** Neu5Ac(a2-3)Gal(b1-3)[Fuc(a1-4)]GlcNAc(b1-3)Gal(b1-4)[Fuc(a1-3)]GlcNAcb-SpSp

**IUPAC (CFG):** Neu5Aca2-3Galb1-3[Fuca1-4]GlcNAcb1-3Galb1-4[Fuca1-3]GlcNAcb-Sp

**Common Name:** 3'SLeA-Lex

**Glytoucan ID:** NA

**Compound Number:** Te304

**Maldi File:** MS-XIV-75i.txt

**Density:** 21

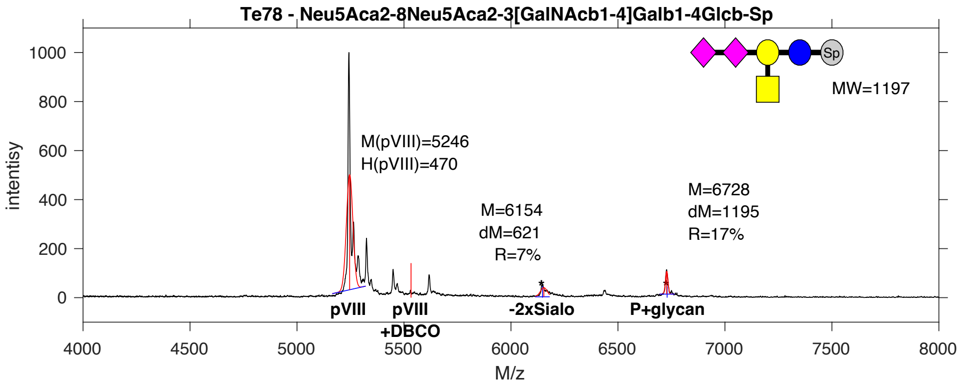


**18: MALDI-TOF spectra of glycosylated phage: GD2-[460]**

**SDB Number:** SDB100

**Sequencing File:** http://ligacloud.ca/searchLibInfo?f=0&b=0&d=20210319-87CLooEJ-MS

**Barcode:**
CTACTGTTCGCAATCCCGCTCAGCGTCGAGAAGAATGACCAGAAGACGTA
CCATGCCGGGGGCGGG

**Axis Name:** GD2-[460]

**IUPAC:** Neu5Ac(a2-8)Neu5Ac(a2-3)[GalNAc(b1-4)]Gal(b1-4)Glc(b1-Sp

**IUPAC (CFG):** Neu5Aca2-8Neu5Aca2-3[GalNAcb1-4]Galb1-4Glcb-Sp

**Common Name:** GD2

**Glytoucan ID:** G79648LW

**Compound Number:** Te78

**Maldi File:** MS-XIV-75v.txt

**Density:** 17

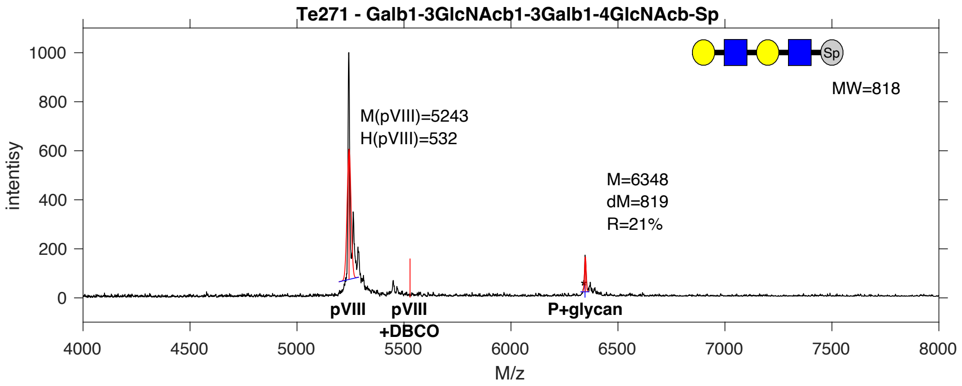


**19: MALDI-TOF spectra of glycosylated phage: LNT-Nac-[410]**

**SDB Number:** SDB180

**Sequencing File:** http://ligacloud.ca/searchLibInfo?f=0&b=0&d=NA

**Barcode:**
CTACTGTTCGCAATCCCGCTAAGTGTTGAGAAAAATGATCAAAAAACTTA
TCACGCCGGTGGTGGT

**Axis Name:** LNT-Nac-[410]

**IUPAC:** Gal(b1-3)GlcNAc(b1-3)Gal(b1-4)GlcNAc(b1-Sp

**IUPAC (CFG):** Galb1-3GlcNAcb1-3Galb1-4GlcNAcb-Sp

**Common Name:** LNT-NAc

**Glytoucan ID:** NA

**Compound Number:** LNT-NAc

**Maldi File:** MS-XIV-77i.txt

**Density:** 20

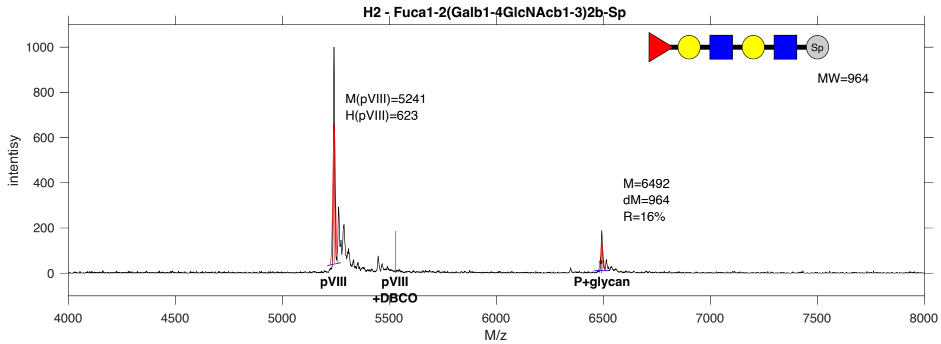


**20: MALDI-TOF spectra of glycosylated phage: H2-[430]**

**SDB Number:** SDB35

**Sequencing File:** http://ligacloud.ca/searchLibInfo?f=0&b=0&d=NA

**Barcode:**
CTGCTCTTCGCTATTCCACTTAGTGTGGAGAAGAATGATCAGAAGACTTAT
CATGCGGGTGGAGGT

**Axis Name:** H2-[430]

**IUPAC:** Fuc(a1-2)Gal(b1-4)GlcNAc(b1-3)Gal(b1-4)GlcNAc(b1-Sp

**IUPAC (CFG):** Fuca1-2(Galb1-4GlcNAcb1-3)2b-Sp

**Common Name:** H2

**Glytoucan ID:** G99268MP

**Compound Number:** Te134

**Maldi File:** RR-III-108.txt

**Density:** 16

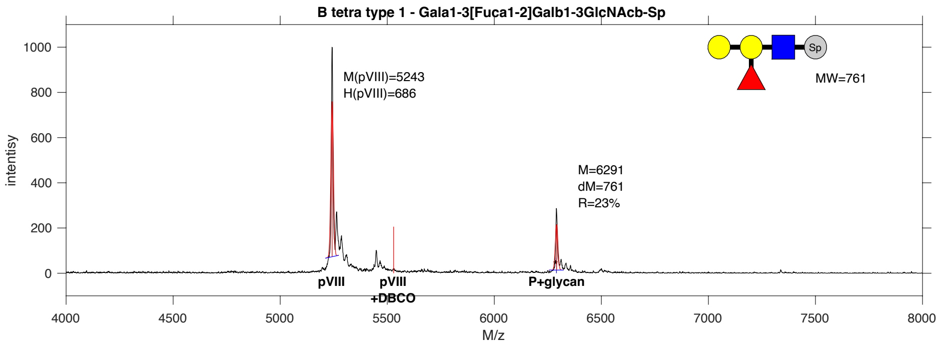


**21: MALDI-TOF spectra of glycosylated phage: B tetra type 1-[620]**

**SDB Number:** SDB142

**Sequencing File:** http://ligacloud.ca/searchLibInfo?f=0&b=0&d=20210319-87CLooPF-MS

**Barcode:** CTGCTTTTTGCTATACCCCTGAGCGTTGAGAAGAATGACCAGAAGACTTATCACGCAGGTGGAGGG

**Axis Name:** B tetra type 1-[620]

**IUPAC:** Gal(a1-3)[Fuc(a1-2)]Gal(b1-3)GlcNAc(b1-Sp

**IUPAC (CFG):** Gala1-3[Fuca1-2]Galb1-3GlcNAcb-Sp

**Common Name:** B tetra type 1

**Glytoucan ID:** G31734OS

**Compound Number:** Te258

**Maldi File:** RR-III-82i.txt

**Density:** 23

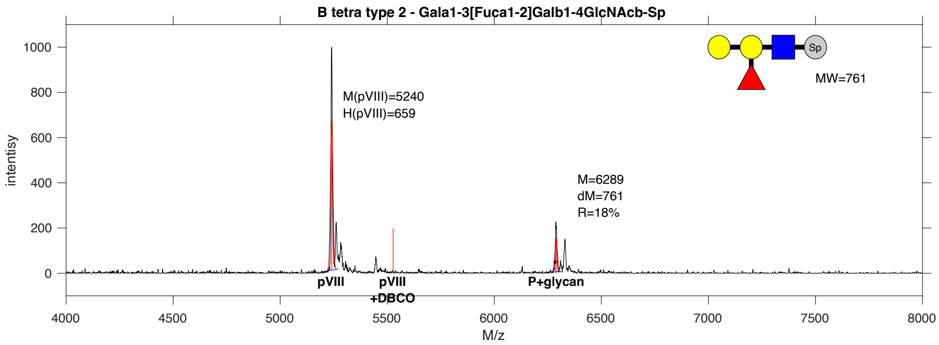


**22: MALDI-TOF spectra of glycosylated phage: B tetra type 2-[970]**

**SDB Number:** SDB144

**Sequencing File:** http://ligacloud.ca/searchLibInfo?f=0&b=0&d=20210319-87CLooLC-MS

**Barcode:** CTTCTTTTTGCCATACCCCTGAGCGTAGAGAAGAATGATCAAAAGACTTATCATGCTGGGGGAGGA

**Axis Name:** B tetra type 2-[970]

**IUPAC:** Gal(a1-3)[Fuc(a1-2)]Gal(b1-4)GlcNAc(b1-Sp

**IUPAC (CFG):** Gala1-3[Fuca1-2]Galb1-4GlcNAcb-Sp

**Common Name:** B tetra type 2

**Glytoucan ID:** G45802NQ

**Compound Number:** Te223

**Maldi File:** RR-III-83i.txt

**Density:** 36

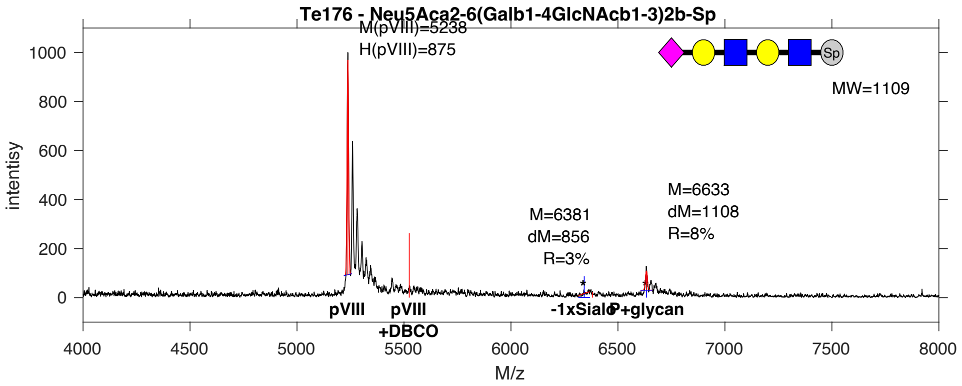


**23: MALDI-TOF spectra of glycosylated phage: 6’S-Di-LN-[300]**

**SDB Number:** SDB197

**Sequencing File:** http://ligacloud.ca/searchLibInfo?f=0&b=0&d=NA

**Barcode:**
CTGCTGTTTGCGATACCGCTTAGTGTTGAAAAAAATGACCAGAAAACCTACCACGCCGGTGGGGGT

**Axis Name:** 6’S-Di-LN-[300]

**IUPAC:** Neu5Ac(a2-6)Gal(b1-4)GlcNAc(b1-3)Gal(b1-4)GlcNAc(b1-Sp

**IUPAC (CFG):** Neu5Aca2-6(Galb1-4GlcNAcb1-3)2b-Sp

**Common Name:** 6’S-Di-LN

**Glytoucan ID:** G60430KO

**Compound Number:** Te176

**Maldi File:** MS-XIV-40ii.txt

**Density:** 11

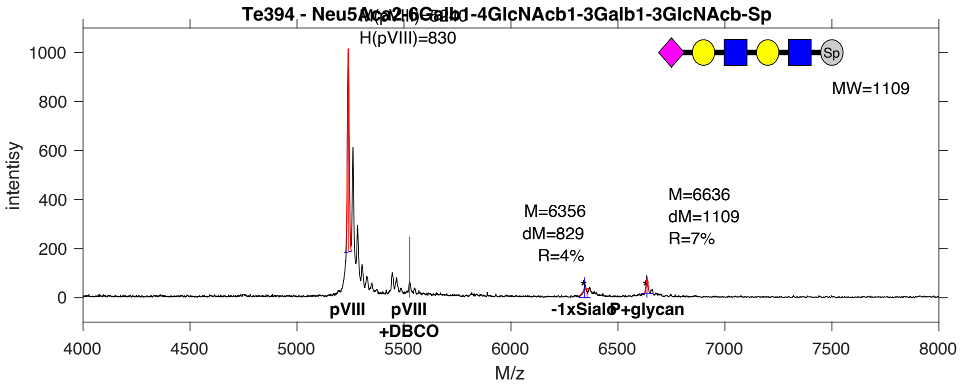


**24: MALDI-TOF spectra of glycosylated phage: 6SLN-Lec-[300]**

**SDB Number:** SDB199

**Sequencing File:** http://ligacloud.ca/searchLibInfo?f=0&b=0&d=NA

**Barcode:**
CTGCTCTTTGCTATACCCCTAAGCGTAGAAAAAAACGATCAGAAAACATA
TCATGCTGGAGGTGGT

**Axis Name:** 6SLN-Lec-[300]

**IUPAC:** Neu5Ac(a2-6)Gal(b1-4)GlcNAc(b1-3)Gal(b1-3)GlcNAc(b1-Sp

**IUPAC (CFG):** Neu5Aca2-6Galb1-4GlcNAcb1-3Galb1-3GlcNAcb-Sp

**Common Name:** 6SLN-Lec

**Glytoucan ID:** G08396BF

**Compound Number:** Te324

**Maldi File:** MS-XIV-40iv.txt

**Density:** 11

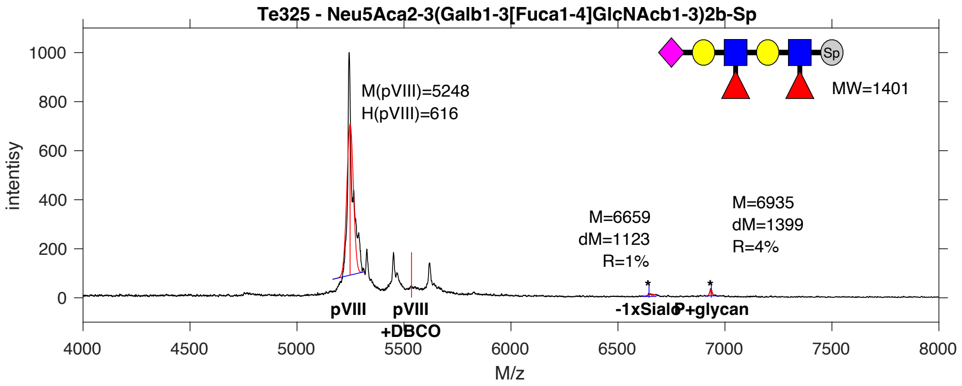


**25: MALDI-TOF spectra of glycosylated phage: 3'S-Di-LeA-[160]**

**SDB Number:** SDB97

**Sequencing File:** http://ligacloud.ca/searchLibInfo?f=0&b=0&d=NA

**Barcode:**
CTGCTGTTTGCAATCCCCCTAAGTGTCGAAAAGAACGATCAGAAAACATA
CCATGCGGGTGGGGGT

**Axis Name:** 3'S-Di-LeA-[160]

**IUPAC:** Neu5Ac(a2-3)Gal(b1-3)[Fuc(a1-4)]GlcNAc(b1-3)Gal(b1-3)[Fuc(a1-4)]GlcNAc(b1-Sp

**IUPAC (CFG):** Neu5Aca2-3(Galb1-3[Fuca1-4]GlcNAcb1-3)2b-Sp

**Common Name:** 3'S-Di-LeA

**Glytoucan ID:** G12518FG

**Compound Number:** Te325

**Maldi File:** MS-XIV-75ii.txt

**Density:** 6

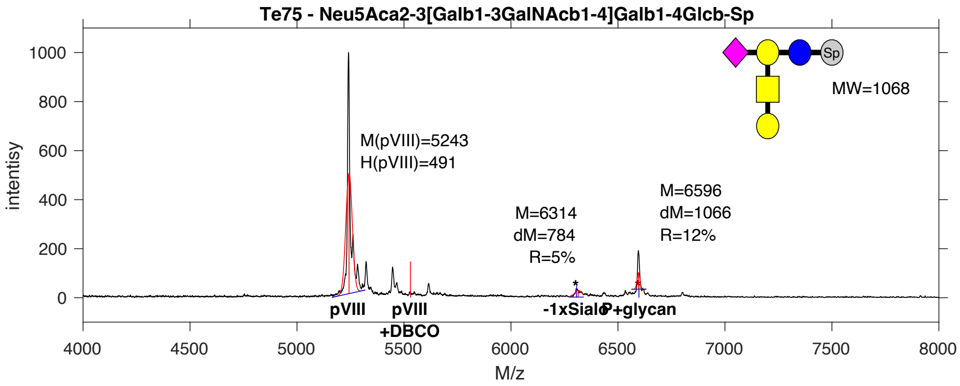


**26: MALDI-TOF spectra of glycosylated phage: GM1-[460]**

**SDB Number:** SDB99

**Sequencing File:** http://ligacloud.ca/searchLibInfo?f=0&b=0&d=NA

**Barcode:**
CTTCTCTTCGCGATACCGCTAAGTGTTGAGAAAAATGATCAAAAAACGTA
TCATGCGGGTGGTGGG

**Axis Name:** GM1-[460]

**IUPAC:** Neu5Ac(a2-3)[Gal(b1-3)GalNAc(b1-4)]Gal(b1-4)Glc(b1-Sp

**IUPAC (CFG):** Neu5Aca2-3[Galb1-3GalNAcb1-4]Galb1-4Glcb-Sp

**Common Name:** GM1

**Glytoucan ID:** G40306MD

**Compound Number:** Te75

**Maldi File:** MS-XIV-75iv.txt

**Density:** 17

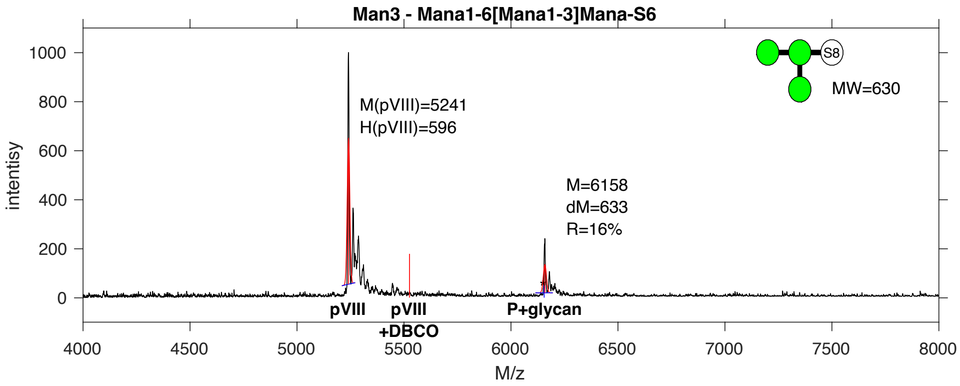


**27: MALDI-TOF spectra of glycosylated phage: (Man)3-[540]**

**SDB Number:** SDB177

**Sequencing File:** http://ligacloud.ca/searchLibInfo?f=0&b=0&d=NA

**Barcode:**
CTTCTATTTGCTATTCCTCTAAGTGTCGAGAAGAACGACCAGAAGACGTA
TCATGCGGGGGGTGGT

**Axis Name:** (Man)3-[540]

**IUPAC:** Man(a1-6)[Man(a1-3)]Man(a1-S6

**IUPAC (CFG):** Mana1-6[Mana1-3]Mana-S6

**Common Name:** (Man)3

**Glytoucan ID:** G80926OA

**Compound Number:** (Man)3

**Maldi File:** MS-XIV-77ii.txt

**Density:** 15

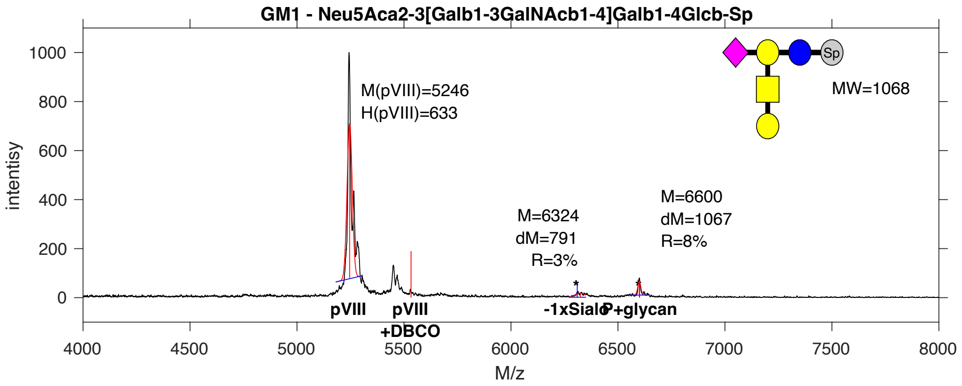


**28: MALDI-TOF spectra of glycosylated phage: GM1-[460]**

**SDB Number:** SDB10

**Sequencing File:** http://ligacloud.ca/searchLibInfo?f=0&b=0&d=NA

**Barcode:**
CTACTGTTTGCTATACCGCTGAGTGTGGAGAAGAATGATCAGAAGACTTA
TCATGCGGGTGGAGGT

**Axis Name:** GM1-[460]

**IUPAC:** Neu5Ac(a2-3)[Gal(b1-3)GalNAc(b1-4)]Gal(b1-4)Glc(b1-Sp

**IUPAC (CFG):** Neu5Aca2-3[Galb1-3GalNAcb1-4]Galb1-4Glcb-Sp

**Common Name:** GM1

**Glytoucan ID:** G40306MD

**Compound Number:** Te75

**Maldi File:** MS-XV-63_10.txt

**Density:** 17

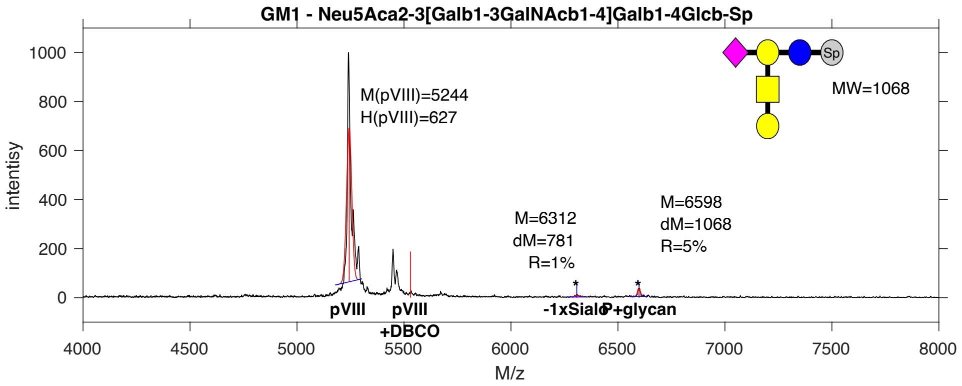


**29: MALDI-TOF spectra of glycosylated phage: GM1-[190]**

**SDB Number:** SDB39

**Sequencing File:** http://ligacloud.ca/searchLibInfo?f=0&b=0&d=NA

**Barcode:**
CTGCTGTTTGCGATCCCTCTAAGTGTCGAAAAGAACGACCAAAAAACTTACCTTGCAGGGGGGGGT

**Axis Name:** GM1-[190]

**IUPAC:** Neu5Ac(a2-3)[Gal(b1-3)GalNAc(b1-4)]Gal(b1-4)Glc(b1-Sp

**IUPAC (CFG):** Neu5Aca2-3[Galb1-3GalNAcb1-4]Galb1-4Glcb-Sp

**Common Name:** GM1

**Glytoucan ID:** G40306MD

**Compound Number:** Te75

**Maldi File:** MS-XV-63_39.txt

**Density:** 7

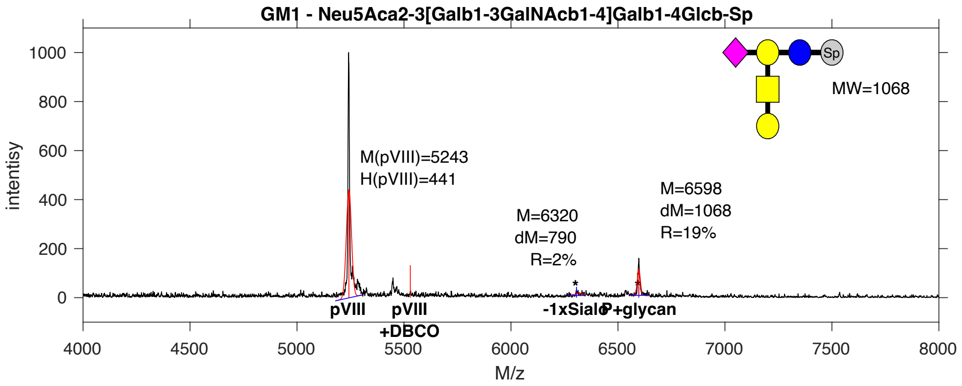


**30: MALDI-TOF spectra of glycosylated phage: GM1-[1190]**

**SDB Number:** SDB91

**Sequencing File:** http://ligacloud.ca/searchLibInfo?f=0&b=0&d=NA

**Barcode:**
CTGCTGTTTGCGATTCCCCTGAGCGTCGAGAAGAATGACCAAAAAACGTA
CCATGCGGGGGGTGGG

**Axis Name:** GM1-[1190]

**IUPAC:** Neu5Ac(a2-3)[Gal(b1-3)GalNAc(b1-4)]Gal(b1-4)Glc(b1-Sp

**IUPAC (CFG):** Neu5Aca2-3[Galb1-3GalNAcb1-4]Galb1-4Glcb-Sp

**Common Name:** GM1

**Glytoucan ID:** G40306MD

**Compound Number:** Te75

**Maldi File:** MS-XV-63_91.txt

**Density:** 44

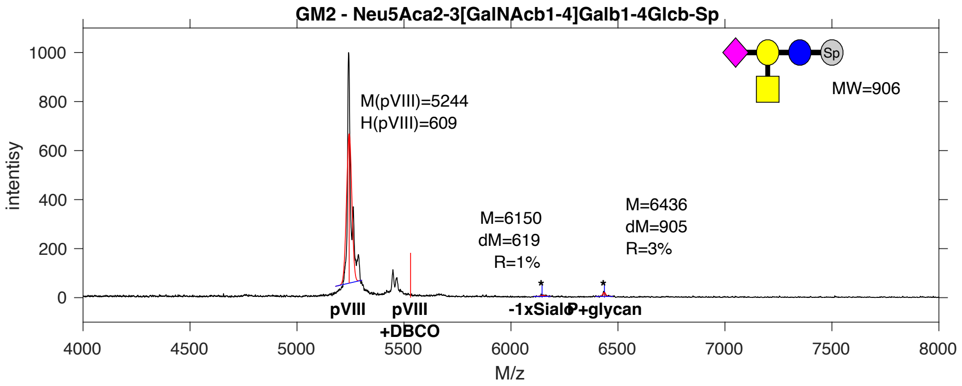


**31: MALDI-TOF spectra of glycosylated phage: GM2-[460]**

**SDB Number:** SDB92

**Sequencing File:** http://ligacloud.ca/searchLibInfo?f=0&b=0&d=NA

**Barcode:**
TTATTATTCGCAATTCCTTTAAGTGTGGAGAAGAATGACCAGAAAACGTA
CCACGCGGGGGGTGGG

**Axis Name:** GM2-[460]

**IUPAC:** Neu5Ac(a2-3)[GalNAc(b1-4)]Gal(b1-4)Glc(b1-Sp

**IUPAC (CFG):** Neu5Aca2-3[GalNAcb1-4]Galb1-4Glcb-Sp

**Common Name:** GM2

**Glytoucan ID:** G79389NT

**Compound Number:** Te74

**Maldi File:** MS-XV-63_92.txt

**Density:** 17

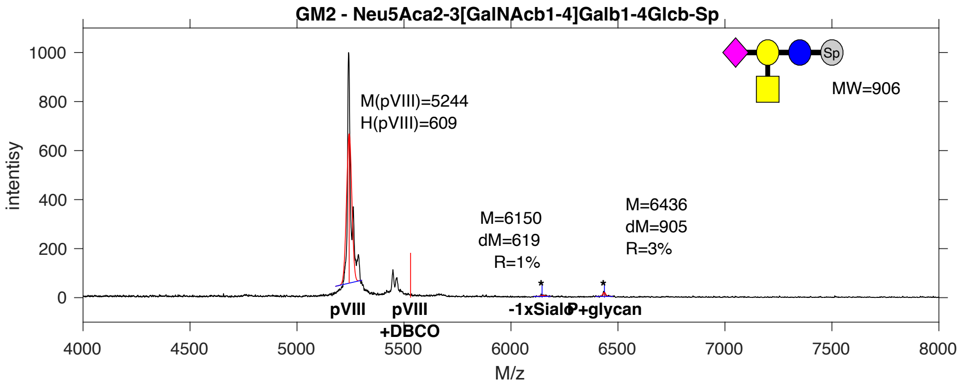


**32: MALDI-TOF spectra of glycosylated phage: GM2-[190]**

**SDB Number:** SDB97

**Sequencing File:** http://ligacloud.ca/searchLibInfo?f=0&b=0&d=NA

**Barcode:**
CTGCTGTTTGCAATCCCCCTAAGTGTCGAAAAGAACGATCAGAAAACATA
CCATGCGGGTGGGGGT

**Axis Name:** GM2-[190]

**IUPAC:** Neu5Ac(a2-3)[GalNAc(b1-4)]Gal(b1-4)Glc(b1-Sp

**IUPAC (CFG):** Neu5Aca2-3[GalNAcb1-4]Galb1-4Glcb-Sp

**Common Name:** GM2

**Glytoucan ID:** G79389NT

**Compound Number:** Te74

**Maldi File:** MS-XV-63_97.txt

**Density:** 7

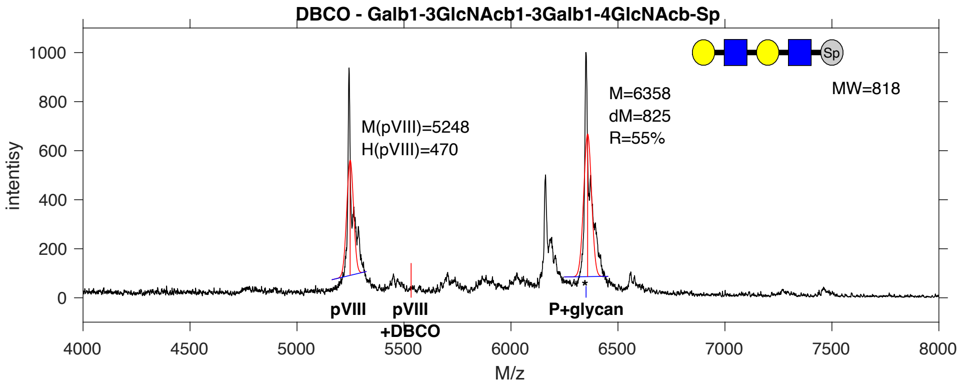


**33: MALDI-TOF spectra of glycosylated phage: (Galf)4-[1490]**

**SDB Number:** SDB138

**Sequencing File:** http://ligacloud.ca/searchLibInfo?f=0&b=0&d=NA

**Barcode:**
CTACTATTCGCTATACCCCTAAGCGTGGAGAAAAATGACCAGAAGACGTA
TCACGCGGGGGGGGGG

**Axis Name:** (Galf)4-[1490]

**IUPAC:** Galf(b1-5)Galf(b1-5)Galf(b1-5)Galf(b1-S8

**IUPAC (CFG):** Galfb1-5Galfb1-5Galfb1-5Galfb-S8

**Common Name:** (Galf)4

**Glytoucan ID:** G92890KE

**Compound Number:** Galf4

**Maldi File:** RR-II-111_2.txt

**Density:** 55

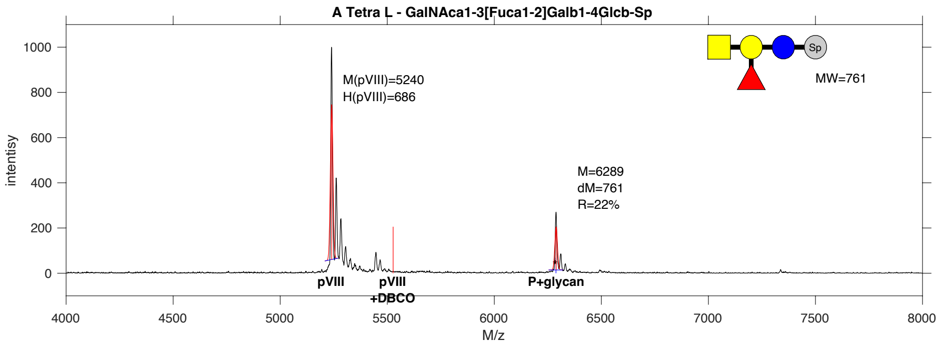


**34: MALDI-TOF spectra of glycosylated phage: A tetra L-[590]**

**SDB Number:** SDB15

**Sequencing File:** http://ligacloud.ca/searchLibInfo?f=0&b=0&d=NA

**Barcode:**
CTGCTGTTCGCCATACCCCTTAGTGTGGAGAAGAATGATCAGAAGACTTA
TCATGCGGGTGGAGGT

**Axis Name:** A tetra L-[590]

**IUPAC:** GalNAc(a1-3)[Fuc(a1-2)]Gal(b1-4)Glc(b1-Sp

**IUPAC (CFG):** GalNAca1-3[Fuca1-2]Galb1-4Glcb-Sp

**Common Name:** A tetra L

**Glytoucan ID:** G19524AS

**Compound Number:** Te224

**Maldi File:** RR-III-107i.txt

**Density:** 22

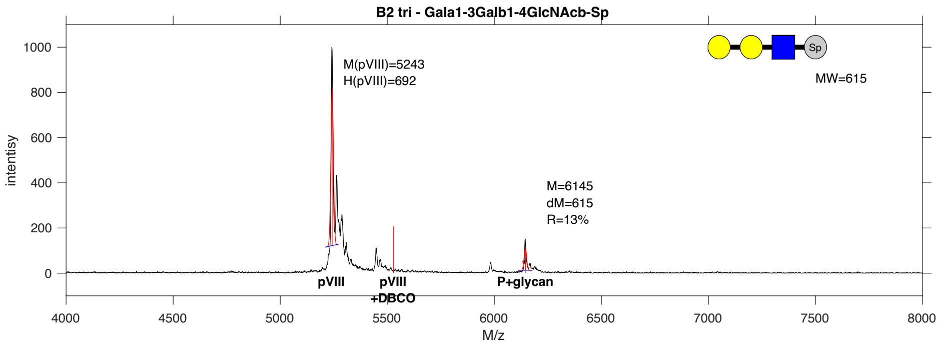


**35: MALDI-TOF spectra of glycosylated phage: B2 tri-[350]**

**SDB Number:** SDB143

**Sequencing File:** http://ligacloud.ca/searchLibInfo?f=0&b=0&d=20210319-87CLooQI-MS

**Barcode:** CTGCTGTTCGCGATTCCCCTAAGTGTCGAAAAAAATGATCAAAAAACCTATCACGCAGGAGGCGGT

**Axis Name:** B2 tri-[350]

**IUPAC:** Gal(a1-3)Gal(b1-4)GlcNAc(b1-Sp

**IUPAC (CFG):** Gala1-3Galb1-4GlcNAcb-Sp

**Common Name:** B2 tri

**Glytoucan ID:** NA

**Compound Number:** Tr60

**Maldi File:** RR-III-82ii.txt

**Density:** 13

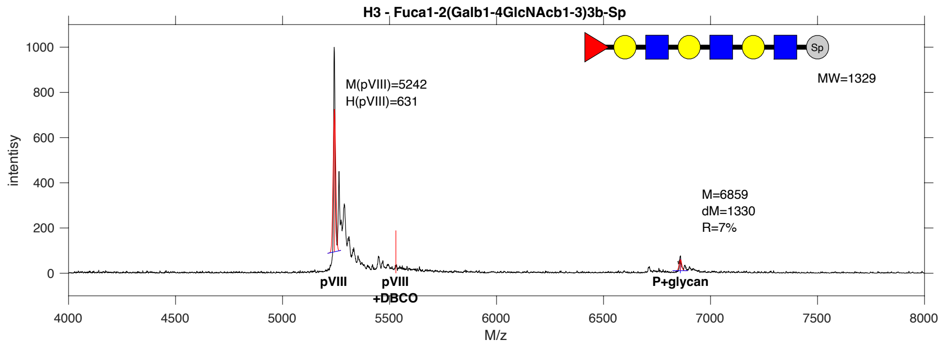


#### 36: MALDI-TOF spectra of glycosylated phage: H3-[190]

**SDB Number:** SDB145

**Sequencing File:** http://ligacloud.ca/searchLibInfo?f=0&b=0&d=20210319-87CLooIH-MS

**Barcode:** CTGCTTTTTGCCATTCCACTTAGTGTTGAGAAGAACGACCAGAAAACGTACCACGCGGGGGGTGGG

**Axis Name:** H3-[190]

**IUPAC:** Fuc(a1-2)Gal(b1-4)GlcNAc(b1-3)Gal(b1-4)GlcNAc(b1-3)Gal(b1-4)GlcNAc(b1-Sp

**IUPAC (CFG):** Fuca1-2(Galb1-4GlcNAcb1-3)3b-Sp

**Common Name:** H3

**Glytoucan ID:** G85213RR

**Compound Number:** Te135

**Maldi File:** RR-III-83ii.txt

**Density:** 7

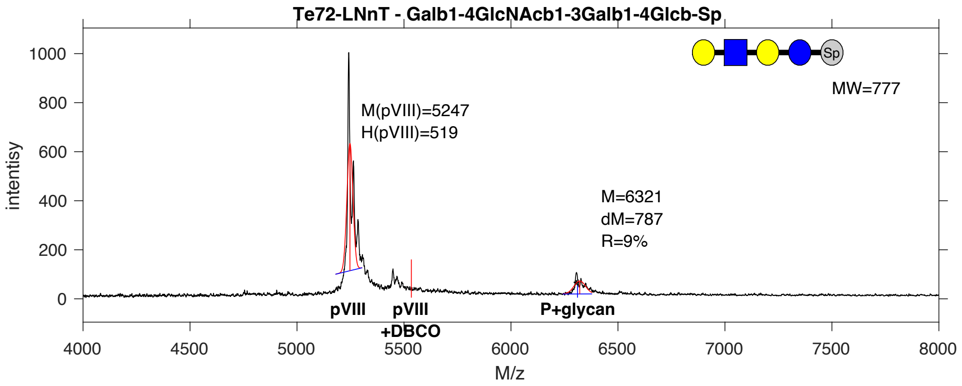


**37: MALDI-TOF spectra of glycosylated phage: LNnT-[240]**

**SDB Number:** SDB8

**Sequencing File:** http://ligacloud.ca/searchLibInfo?f=0&b=0&d=NA

**Barcode:**
CTGCTTTTTGCAATACCCCTCAGTGTGGAGAAGAATGATCAGAAGACTTA
TCATGCGGGTGGAGGT

**Axis Name:** LNnT-[240]

**IUPAC:** NANA

**IUPAC (CFG):** Galb1-4GlcNAcb1-3Galb1-4Glcb-Sp

**Common Name:** LNnT

**Glytoucan ID:** G14139DE

**Compound Number:** Te72

**Maldi File:** MS-XIV-164_8.txt

**Density:** 9

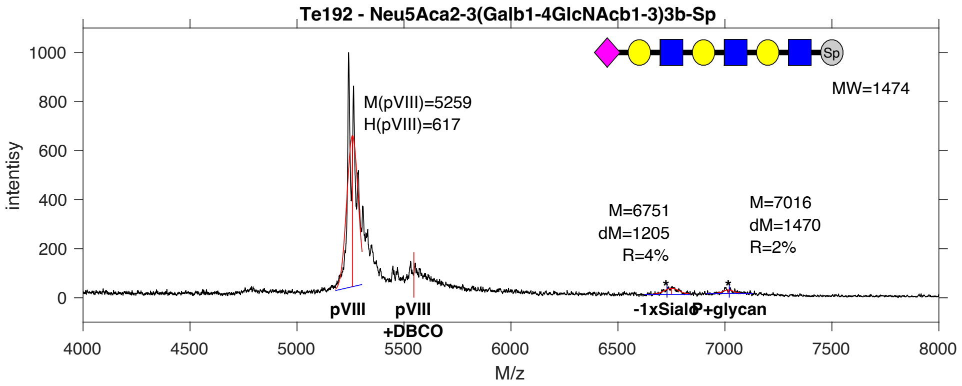


**38: MALDI-TOF spectra of glycosylated phage: 3’STri-LN-[160]**

**SDB Number:** SDB196

**Sequencing File:** http://ligacloud.ca/searchLibInfo?f=0&b=0&d=NA

**Barcode:**
CTGCTTTTTGCTATTCCTCTGAGTGTTGAGAAGAATGACCAGAAGACGTAC
CACGCGGGCGGGGGT

**Axis Name:** 3’STri-LN-[160]

**IUPAC:** NASp

**IUPAC (CFG):** Neu5Aca2-3(Galb1-4GlcNAcb1-3)3b-Sp

**Common Name:** 3’STri-LN

**Glytoucan ID:** NA

**Compound Number:** Te192

**Maldi File:** MS-XIV-40iii.txt

**Density:** 6

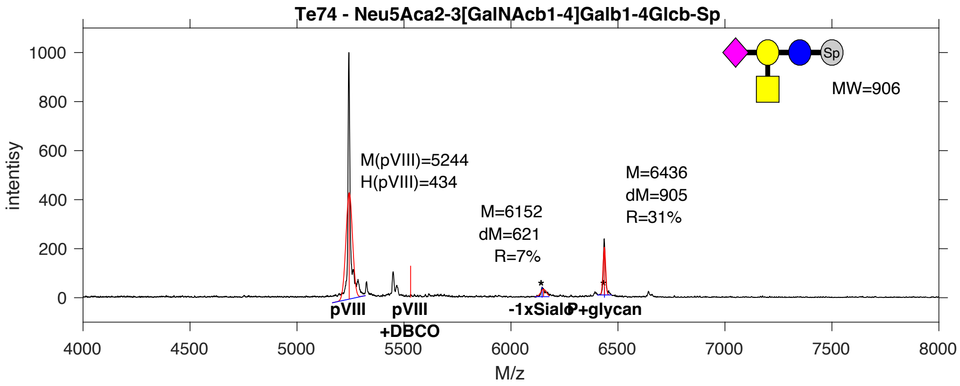


**39: MALDI-TOF spectra of glycosylated phage: GM2-[590]**

**SDB Number:** SDB98

**Sequencing File:** http://ligacloud.ca/searchLibInfo?f=0&b=0&d=NA

**Barcode:**
CTGCTGTTTGCGATTCCTCTGAGTGTCGAAAAGAATGATCAAAAAACATA
CCATGCGGGGGGTGGG

**Axis Name:** GM2-[590]

**IUPAC:** Neu5Ac(a2-3)[GalNAc(b1-4)]Gal(b1-4)Glc(b1-Sp

**IUPAC (CFG):** Neu5Aca2-3[GalNAcb1-4]Galb1-4Glcb-Sp

**Common Name:** GM2

**Glytoucan ID:** G79389NT

**Compound Number:** Te74

**Maldi File:** MS-XIV-75iii.txt

**Density:** 22

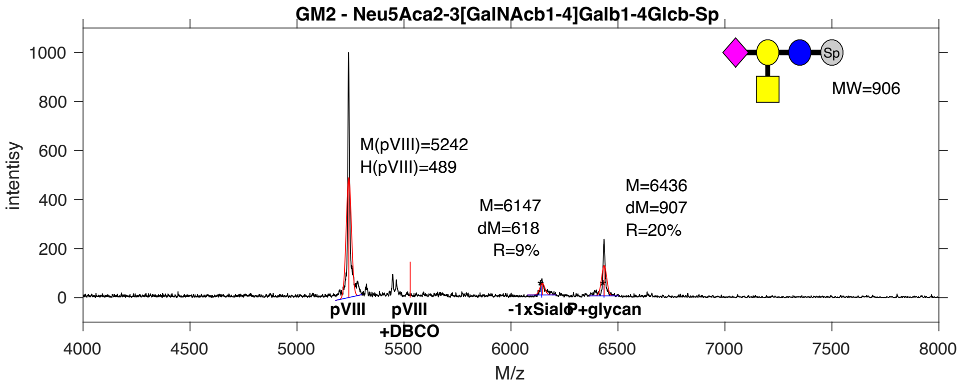


**40: MALDI-TOF spectra of glycosylated phage: GM2-[1190]**

**SDB Number:** SDB162

**Sequencing File:** http://ligacloud.ca/searchLibInfo?f=0&b=0&d=NA

**Barcode:**
TTATTATTCGCAATTCCTTTAAGTGTTGAGAAAAACGACCAGAAGACTTAT
CATGCTGGGGGGGGA

**Axis Name:** GM2-[1190]

**IUPAC:** Neu5Ac(a2-3)[GalNAc(b1-4)]Gal(b1-4)Glc(b1-Sp

**IUPAC (CFG):** Neu5Aca2-3[GalNAcb1-4]Galb1-4Glcb-Sp

**Common Name:** GM2

**Glytoucan ID:** G79389NT

**Compound Number:** Te74

**Maldi File:** MS-XV-63_162.txt

**Density:** 44

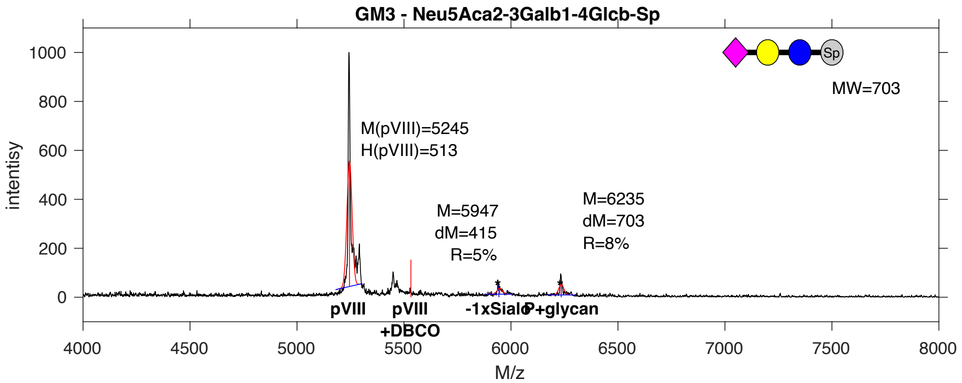


**41: MALDI-TOF spectra of glycosylated phage: GM3-[460]**

**SDB Number:** SDB171

**Sequencing File:** http://ligacloud.ca/searchLibInfo?f=0&b=0&d=NA

**Barcode:**
CTTCTCTTTGCAATCCCGCTGAGCGTCGAGAAGAATGACCAGAAGACATA
CCACGCTGGTGGCGGC

**Axis Name:** GM3-[460]

**IUPAC:** Neu5Ac(a2-3)Gal(b1-4)Glc(b1-Sp

**IUPAC (CFG):** Neu5Aca2-3Galb1-4Glcb-Sp

**Common Name:** GM3

**Glytoucan ID:** G91237TK

**Compound Number:** Tr32

**Maldi File:** MS-XV-63_171.txt

**Density:** 17

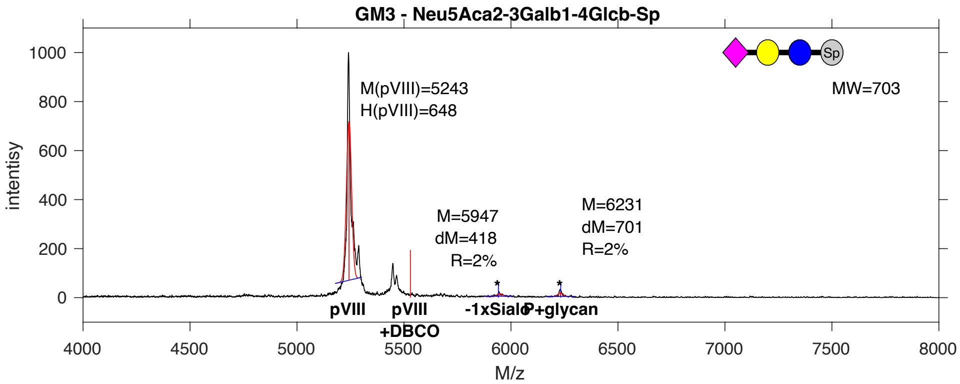


**42: MALDI-TOF spectra of glycosylated phage: GM3-[190]**

**SDB Number:** SDB181

**Sequencing File:** http://ligacloud.ca/searchLibInfo?f=0&b=0&d=NA

**Barcode:**
TTATTATTCGCAATTCCTTTAAGTGTGGAGAAAAACGACCAAAAGACCTA
CCATGCTGGGGGGGGT

**Axis Name:** GM3-[190]

**IUPAC:** Neu5Ac(a2-3)Gal(b1-4)Glc(b1-Sp

**IUPAC (CFG):** Neu5Aca2-3Galb1-4Glcb-Sp

**Common Name:** GM3

**Glytoucan ID:** G91237TK

**Compound Number:** Tr32

**Maldi File:** MS-XV-63_181.txt

**Density:** 7

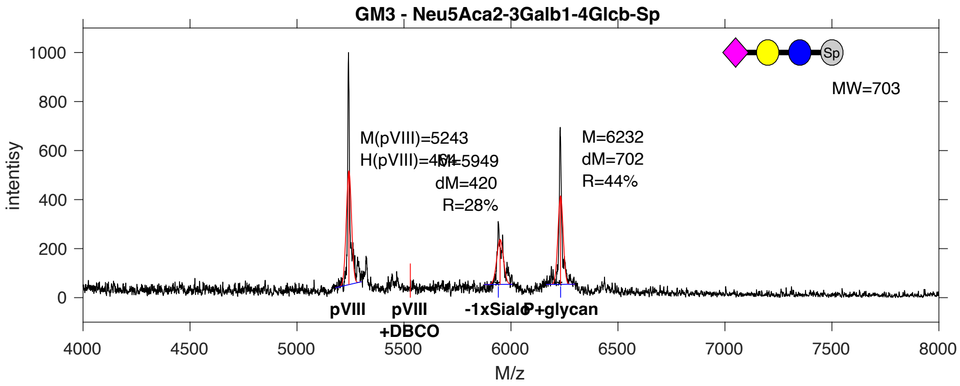


**43: MALDI-TOF spectra of glycosylated phage: GM3-[1190]**

**SDB Number:** SDB192

**Sequencing File:** http://ligacloud.ca/searchLibInfo?f=0&b=0&d=NA

**Barcode:**
CTACTATTTGCGATACCGCTAAGTGTGGAAAAAAACGATCAAAAAACATA
CCACGCTGGCGGCGGG

**Axis Name:** GM3-[1190]

**IUPAC:** Neu5Ac(a2-3)Gal(b1-4)Glc(b1-Sp

**IUPAC (CFG):** Neu5Aca2-3Galb1-4Glcb-Sp

**Common Name:** GM3

**Glytoucan ID:** G91237TK

**Compound Number:** Tr32

**Maldi File:** MS-XV-63_192.txt

**Density:** 44

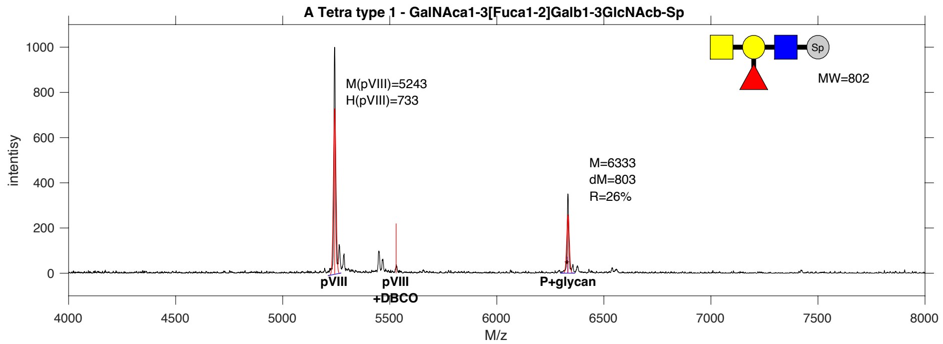


**44: MALDI-TOF spectra of glycosylated phage: A tetra type 1-[700]**

**SDB Number:** SDB34

**Sequencing File:** http://ligacloud.ca/searchLibInfo?f=0&b=0&d=NA

**Barcode:**
CTTCTTTTTGCGATTCCGCTGAGTGTGGAGAAGAATGATCAGAAGACTTAT
CATGCGGGTGGAGGT

**Axis Name:** A tetra type 1-[700]

**IUPAC:** GalNAc(a1-3)[Fuc(a1-2)]Gal(b1-3)GlcNAc(b1-Sp

**IUPAC (CFG):** GalNAca1-3[Fuca1-2]Galb1-3GlcNAcb-Sp

**Common Name:** A tetra type 1

**Glytoucan ID:** G66163TI

**Compound Number:** Te259

**Maldi File:** RR-III-107ii.txt

**Density:** 26

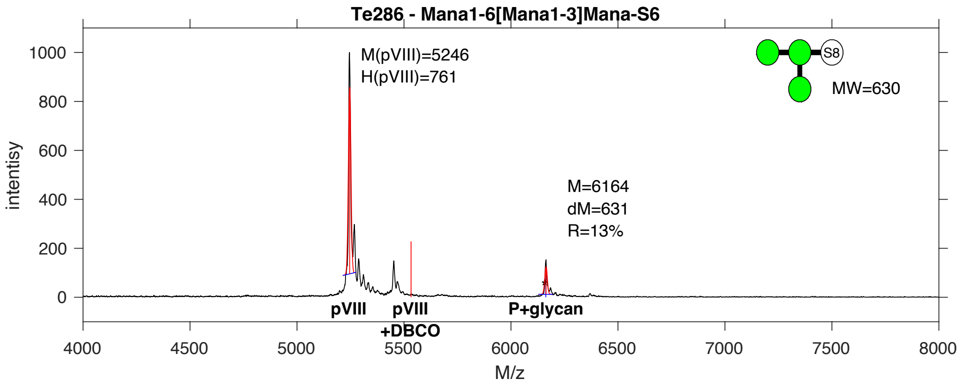


**45: MALDI-TOF spectra of glycosylated phage: (Man)3-[350]**

**SDB Number:** SDB20

**Sequencing File:** http://ligacloud.ca/searchLibInfo?f=0&b=0&d=NA

**Barcode:**
CTGCTCTTTGCCATCCCGCTTAGTGTGGAGAAGAATGATCAGAAGACTTA
TCATGCGGGTGGAGGT

**Axis Name:** (Man)3-[350]

**IUPAC:** Man(a1-6)[Man(a1-3)]Man(a1-S6

**IUPAC (CFG):** Mana1-6[Mana1-3]Mana-S6

**Common Name:** (Man)3

**Glytoucan ID:** G80926OA

**Compound Number:** PZ-8015

**Maldi File:** MS-XIV-114_20.txt

**Density:** 13

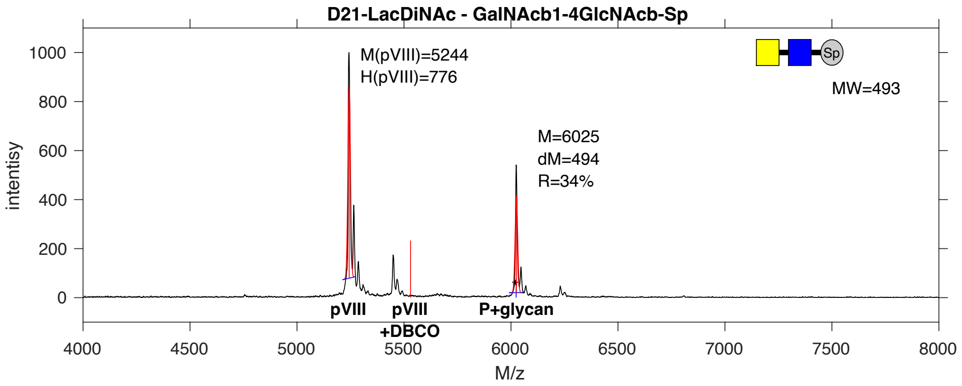


**46: MALDI-TOF spectra of glycosylated phage: LacDiNAc-[920]**

**SDB Number:** SDB28

**Sequencing File:** http://ligacloud.ca/searchLibInfo?f=0&b=0&d=NA

**Barcode:**
CTACTCTTCGCCATTCCACTGAGTGTGGAGAAGAATGATCAGAAGACTTA
TCATGCGGGTGGAGGT

**Axis Name:** LacDiNAc-[920]

**IUPAC:** GalNAc(b1-4)GlcNAc(b1-Sp

**IUPAC (CFG):** GalNAcb1-4GlcNAcb-Sp

**Common Name:** LacDiNAc

**Glytoucan ID:** G33129LQ

**Compound Number:** D21

**Maldi File:** MS-XIV-114_28.txt

**Density:** 34

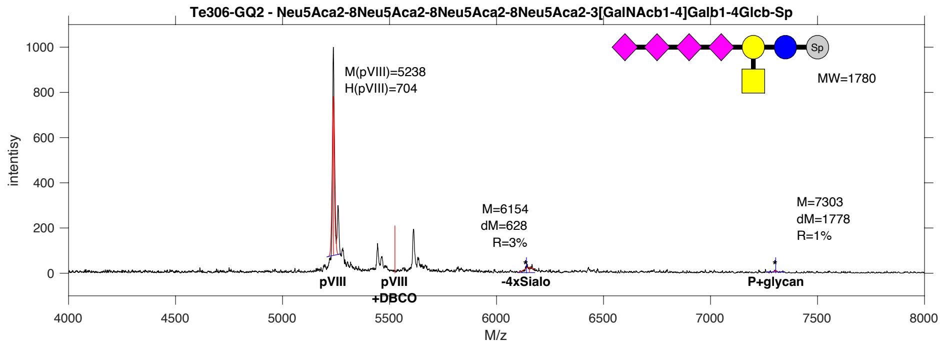


### 47: MALDI-TOF spectra of glycosylated phage: GQ2-[140]

**SDB Number:** SDB44

**Sequencing File:** http://ligacloud.ca/searchLibInfo?f=0&b=0&d=NA

**Barcode:**
CTGCTATTTGCGATTCCCCTAAGTGTAGAGAAAAACGATCAAAAGACGTA
TCATGCCGGGGGCGGT

**Axis Name:** GQ2-[140]

**IUPAC:** Neu5Ac(a2-8)Neu5Ac(a2-8)Neu5Ac(a2-8)Neu5Ac(a2-3)[GalNAc(b1-4)]Gal(b1-4)Glc(b1-Sp

**IUPAC (CFG):** Neu5Aca2-8Neu5Aca2-8Neu5Aca2-8Neu5Aca2-3[GalNAcb1-4]Galb1-4Glcb-Sp

**Common Name:** GQ2

**Glytoucan ID:** G69728IZ

**Compound Number:** Te306

**Maldi File:** MS-XIV-114_44.txt

**Density:** 5

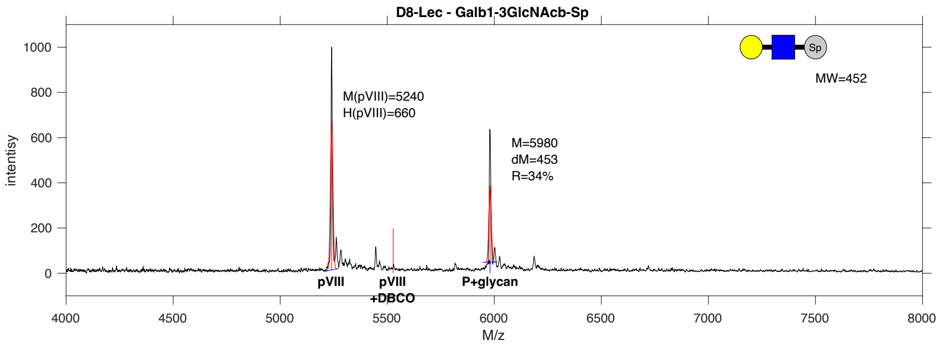


**48: MALDI-TOF spectra of glycosylated phage: Lec-[680]**

**SDB Number:** SDB52

**Sequencing File:** http://ligacloud.ca/searchLibInfo?f=0&b=0&d=NA

**Barcode:**
CTACTGTTTGCTATCCCGCTGAGTGTAGAGAAGAATGACCAAAAGACATA
TCATGCGGGAGGAGGT

**Axis Name:** Lec-[680]

**IUPAC:** Gal(b1-3)GlcNAc(b1-Sp

**IUPAC (CFG):** Galb1-3GlcNAcb-Sp

**Common Name:** Lec

**Glytoucan ID:** G16384KS

**Compound Number:** D8

**Maldi File:** MS-XIV-114_52.txt

**Density:** 25

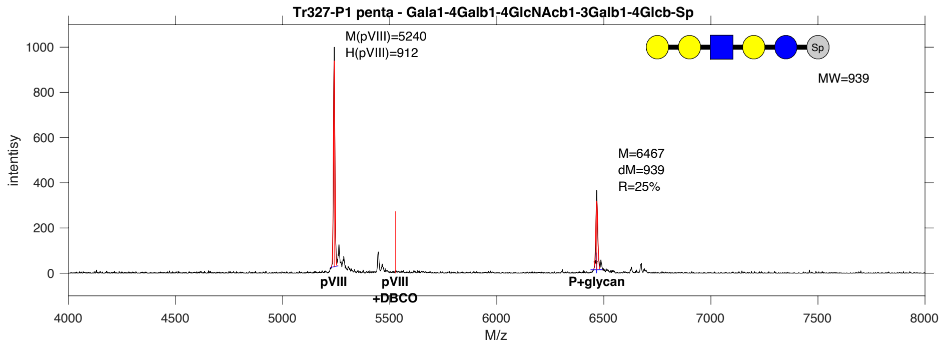


**49: MALDI-TOF spectra of glycosylated phage: P1 penta-[620]**

**SDB Number:** SDB53

**Sequencing File:** http://ligacloud.ca/searchLibInfo?f=0&b=0&d=NA

**Barcode:**
CTGCTGTTTGCGATCCCGCTCAGTGTGGAGAAAAATGACCAAAAGACCTA
TCATGCTGGAGGGGGT

**Axis Name:** P1 penta-[620]

**IUPAC:** Gal(a1-4)Gal(b1-4)GlcNAc(b1-3)Gal(b1-4)Glc(b1-Sp

**IUPAC (CFG):** Gala1-4Galb1-4GlcNAcb1-3Galb1-4Glcb-Sp

**Common Name:** P1 penta

**Glytoucan ID:** G28005AF

**Compound Number:** Te327

**Maldi File:** MS-XIV-114_53.txt

**Density:** 23

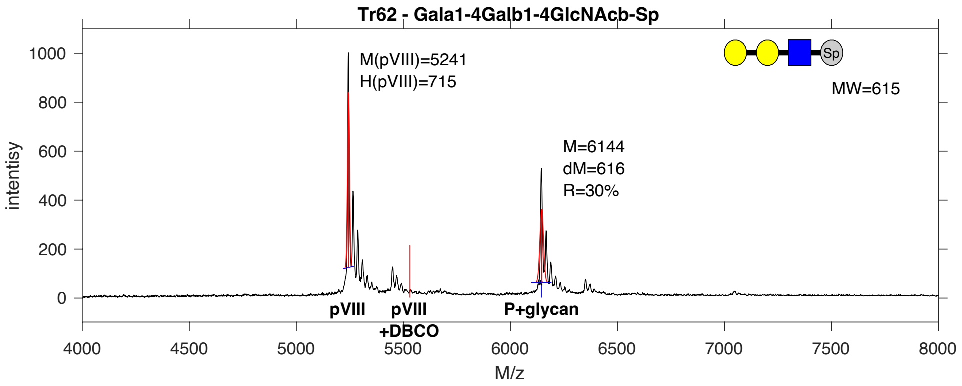


**50: MALDI-TOF spectra of glycosylated phage: P1 tri-[620]**

**SDB Number:** SDB55

**Sequencing File:** http://ligacloud.ca/searchLibInfo?f=0&b=0&d=NA

**Barcode:**
CTGCTCTTTGCAATTCCGCTGAGTGTCGAAAAAAATGACCAAAAGACTTA
TCACGCTGGTGGTGGA

**Axis Name:** P1 tri-[620]

**IUPAC:** Gal(a1-4)Gal(b1-4)GlcNAc(b1-Sp

**IUPAC (CFG):** Gala1-4Galb1-4GlcNAcb-Sp

**Common Name:** P1 tri

**Glytoucan ID:** G00076PK

**Compound Number:** Tr62

**Maldi File:** MS-XIV-114_55.txt

**Density:** 23

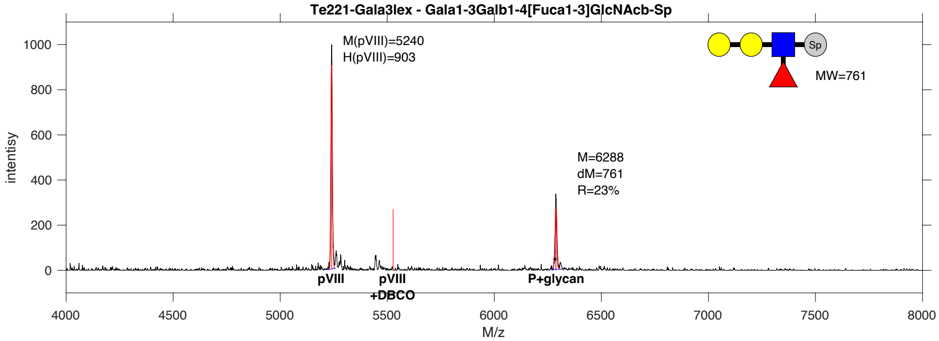


**51: MALDI-TOF spectra of glycosylated phage: Gala3Lex-[620]**

**SDB Number:** SDB56

**Sequencing File:** http://ligacloud.ca/searchLibInfo?f=0&b=0&d=NA

**Barcode:**
CTACTGTTTGCTATACCTCTTAGCGTTGAAAAAAATGACCAGAAAACTTA
CCATGCGGGTGGAGGT

**Axis Name:** Gala3Lex-[620]

**IUPAC:** Gal(a1-3)Gal(b1-4)[Fuc(a1-3)]GlcNAc(b1-Sp

**IUPAC (CFG):** Gala1-3Galb1-4[Fuca1-3]GlcNAcb-Sp

**Common Name:** Gala3Lex

**Glytoucan ID:** G86393KI

**Compound Number:** Te221

**Maldi File:** MS-XIV-114_56.txt

**Density:** 23

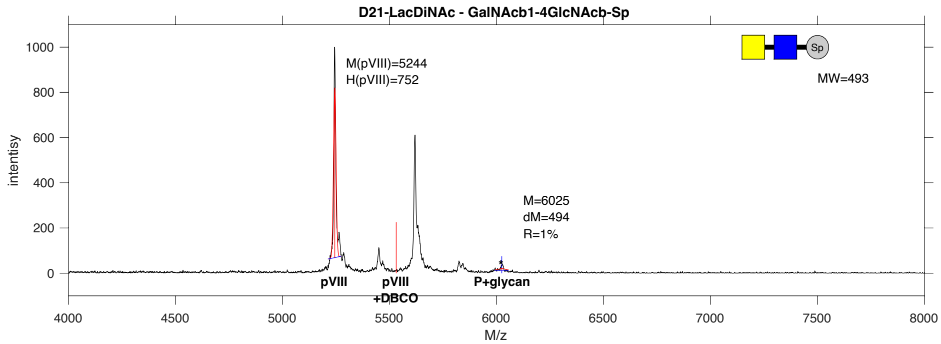


#### 52: MALDI-TOF spectra of glycosylated phage: LacDiNAc-[50]

**SDB Number:** SDB58

**Sequencing File:** http://ligacloud.ca/searchLibInfo?f=0&b=0&d=NA

**Barcode:**
CTACTCTTCGCTATACCCCTCAGCGTGGAGAAAAACGACCAAAAGACCTACCATGCAGGTGGTGGC

**Axis Name:** LacDiNAc-[50]

**IUPAC:** GalNAc(b1-4)GlcNAc(b1-Sp

**IUPAC (CFG):** GalNAcb1-4GlcNAcb-Sp

**Common Name:** LacDiNAc

**Glytoucan ID:** G33129LQ

**Compound Number:** D21

**Maldi File:** MS-XIV-114_58.txt

**Density:** 2

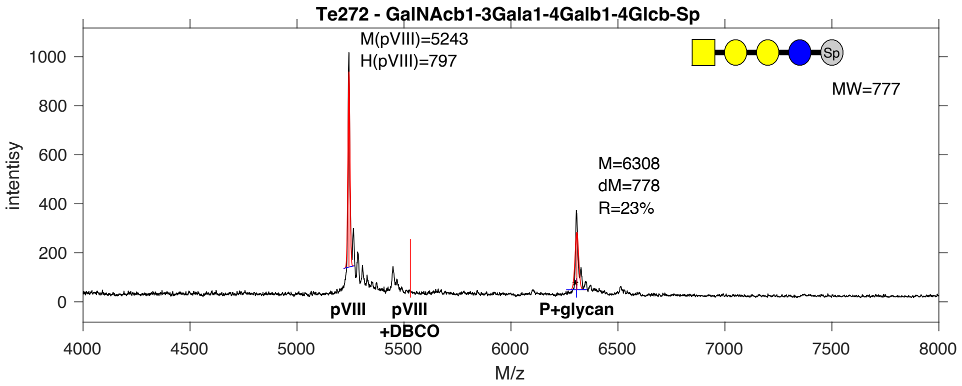


**53: MALDI-TOF spectra of glycosylated phage: Globoside-P-[730]**

**SDB Number:** SDB59

**Sequencing File:** http://ligacloud.ca/searchLibInfo?f=0&b=0&d=NA

**Barcode:**
TTATTATTCGCAATTCCTTTAAGTGTCGAAAAGAATGACCAAAAAACTTATCATGCAGGTGGGGGA

**Axis Name:** Globoside-P-[730]

**IUPAC:** GalNAc(b1-3)Gal(a1-4)Gal(b1-4)Glc(b1-Sp

**IUPAC (CFG):** GalNAcb1-3Gala1-4Galb1-4Glcb-Sp

**Common Name:** Globoside-P

**Glytoucan ID:** G24499XE

**Compound Number:** Te272

**Maldi File:** MS-XIV-114_59.txt

**Density:** 27

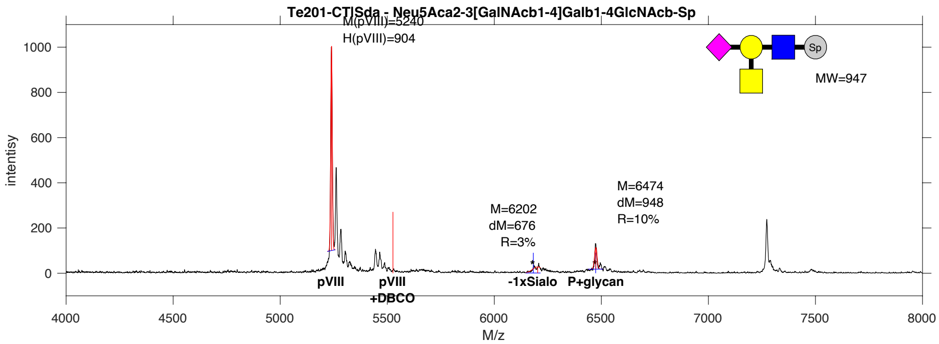


#### 54: MALDI-TOF spectra of glycosylated phage: CT|Sda-[350]

**SDB Number:** SDB65

**Sequencing File:** http://ligacloud.ca/searchLibInfo?f=0&b=0&d=NA

**Barcode:**
CTGCTTTTTGCCATCCCTCTTAGTGTTGAAAAGAACGATCAAAAAACTTAT
CACGCAGGGGGGGGG

**Axis Name:** CT|Sda-[350]

**IUPAC:** Neu5Ac(a2-3)[GalNAc(b1-4)]Gal(b1-4)GlcNAc(b1-Sp

**IUPAC (CFG):** Neu5Aca2-3[GalNAcb1-4]Galb1-4GlcNAcb-Sp

**Common Name:** CT|Sda

**Glytoucan ID:** G41052GC

**Compound Number:** Te201

**Maldi File:** MS-XIV-114_65.txt

**Density:** 13

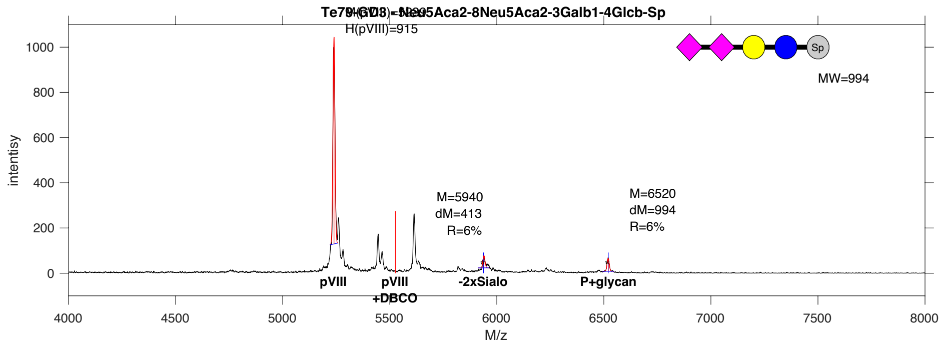


#### 55: MALDI-TOF spectra of glycosylated phage: GD3-[320]

**SDB Number:** SDB68

**Sequencing File:** http://ligacloud.ca/searchLibInfo?f=0&b=0&d=NA

**Barcode:**
CTACTATTCGCGATCCCCCTCAGCGTGGAGAAGAACGACCAGAAGACGTA
TCACGCAGGGGGGGGG

**Axis Name:** GD3-[320]

**IUPAC:** Neu5Ac(a2-8)Neu5Ac(a2-3)Gal(b1-4)Glc(b1-Sp

**IUPAC (CFG):** Neu5Aca2-8Neu5Aca2-3Galb1-4Glcb-Sp

**Common Name:** GD3

**Glytoucan ID:** G07249ZL

**Compound Number:** Te79

**Maldi File:** MS-XIV-114_68.txt

**Density:** 12

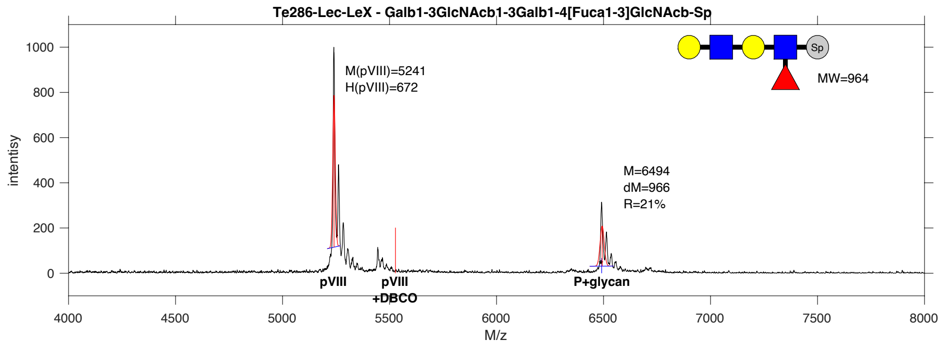


### 56: MALDI-TOF spectra of glycosylated phage: Lec-LeX-[570]

**SDB Number:** SDB69

**Sequencing File:** http://ligacloud.ca/searchLibInfo?f=0&b=0&d=NA

**Barcode:**
CTTCTGTTTGCGATTCCACTGAGCGTGGAGAAGAACGACCAGAAAACATA
CCATGCTGGTGGTGGA

**Axis Name:** Lec-LeX-[570]

**IUPAC:** Gal(b1-3)GlcNac(b1-3)Gal(b1-4)[Fuc(a1-3)]GlcNac(b1-Sp

**IUPAC (CFG):** Galb1-3GlcNAcb1-3Galb1-4[Fuca1-3]GlcNAcb-Sp

**Common Name:** Lec-LeX

**Glytoucan ID:** NA

**Compound Number:** Te286

**Maldi File:** MS-XIV-114_69.txt

**Density:** 21

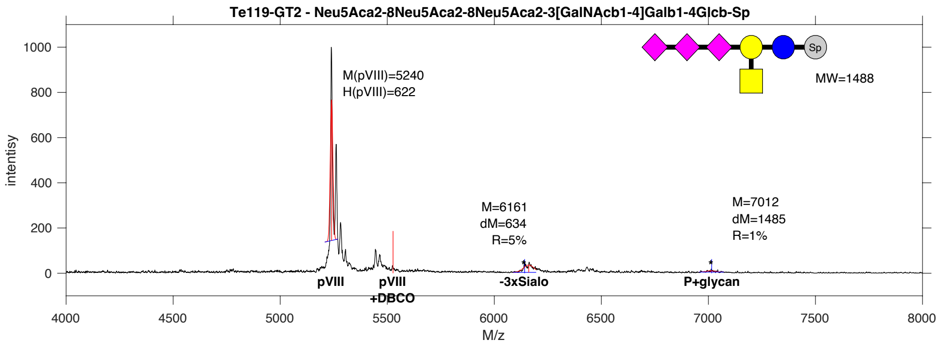


**57: MALDI-TOF spectra of glycosylated phage: GT2-[160]**

**SDB Number:** SDB71

**Sequencing File:** http://ligacloud.ca/searchLibInfo?f=0&b=0&d=NA

**Barcode:**
CTGCTTTTTGCTATTCCTCTGAGTGTTGAAAAAAACGATCAGAAGACTTAT
CACGCGGGGGGCGGG

**Axis Name:** GT2-[160]

**IUPAC:** Neu5Ac(a2-8)Neu5Ac(a2-8)Neu5Ac(a2-3)[GalNAc(b1-4)]Gal(b1-4)Glc(b1-Sp

**IUPAC (CFG):** Neu5Aca2-8Neu5Aca2-8Neu5Aca2-3[GalNAcb1-4]Galb1-4Glcb-Sp

**Common Name:** GT2

**Glytoucan ID:** G35228ZS

**Compound Number:** Te119

**Maldi File:** MS-XIV-114_71.txt

**Density:** 6

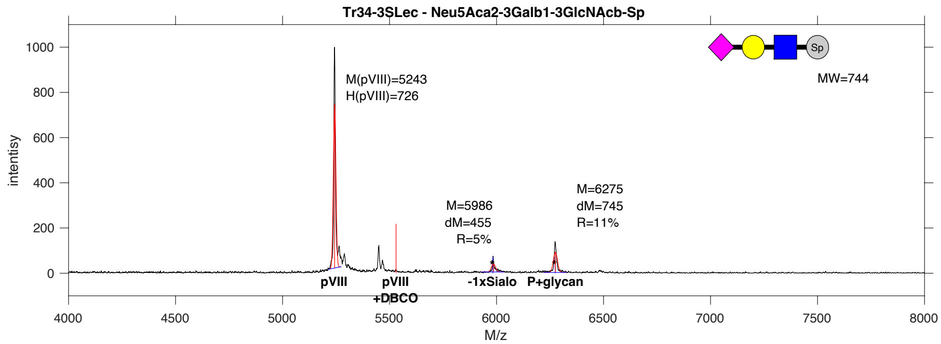


**58: MALDI-TOF spectra of glycosylated phage: 3’SLec-[430]**

**SDB Number:** SDB73

**Sequencing File:** http://ligacloud.ca/searchLibInfo?f=0&b=0&d=NA

**Barcode:**
TTATTATTCGCAATTCCTTTAAGCGTTGAAAAAAACGATCAAAAGACATA
TCACGCTGGCGGAGGA

**Axis Name:** 3’SLec-[430]

**IUPAC:** Neu5Ac(a2-3)Gal(b1-3)GlcNAcb-SpSp

**IUPAC (CFG):** Neu5Aca2-3Galb1-3GlcNAcb-Sp

**Common Name:** 3’SLec

**Glytoucan ID:** NA

**Compound Number:** Tr34

**Maldi File:** MS-XIV-114_73.txt

**Density:** 16

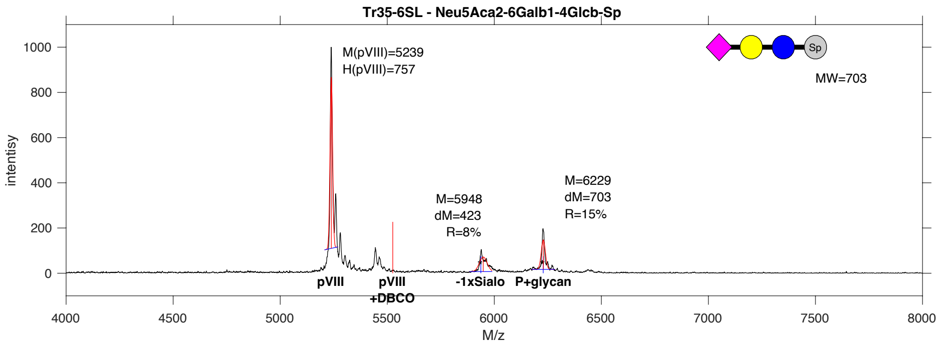


**59: MALDI-TOF spectra of glycosylated phage: 6’SL-[620]**

**SDB Number:** SDB75

**Sequencing File:** http://ligacloud.ca/searchLibInfo?f=0&b=0&d=20210319-87CLooJI-MS

**Barcode:** CTGCTCTTCGCGATCCCTCTAAGTGTGGAAAAAAATGATCAGAAAACGTACCATGCTGGAGGGGGT

**Axis Name:** 6’SL-[620]

**IUPAC:** Neu5Ac(a2-6)Gal(b1-4)Glc(b-Sp

**IUPAC (CFG):** Neu5Aca2-6Galb1-4Glcb-Sp

**Common Name:** 6’SL

**Glytoucan ID:** NA

**Compound Number:** Tr35

**Maldi File:** MS-XIV-114_75.txt

**Density:** 23

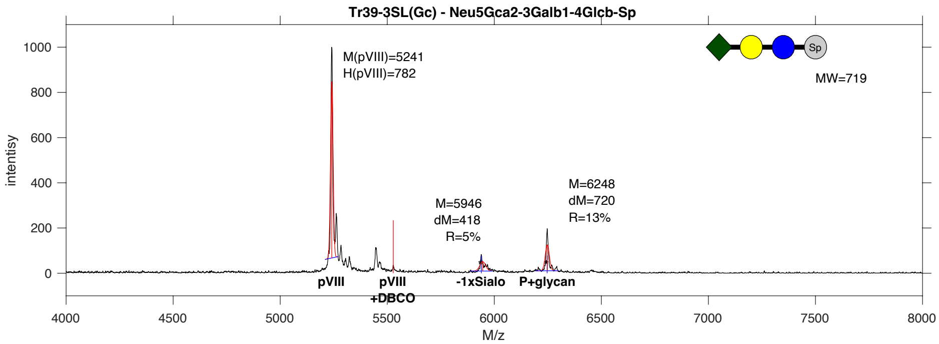


**60: MALDI-TOF spectra of glycosylated phage: 3’SL (Gc)-[490]**

**SDB Number:** SDB77

**Sequencing File:** http://ligacloud.ca/searchLibInfo?f=0&b=0&d=NA

**Barcode:**
CTGCTATTTGCCATCCCGCTGAGCGTCGAGAAAAACGATCAAAAGACATACCATGCCGGCGGAGGA

**Axis Name:** 3’SL (Gc)-[490]

**IUPAC:** Neu5Gc(a2-3)Gal(b1-4)Glc(b1-Sp

**IUPAC (CFG):** Neu5Gca2-3Galb1-4Glcb-Sp

**Common Name:** 3’SL (Gc)

**Glytoucan ID:** G61420QM

**Compound Number:** Tr39

**Maldi File:** MS-XIV-114_77.txt

**Density:** 18

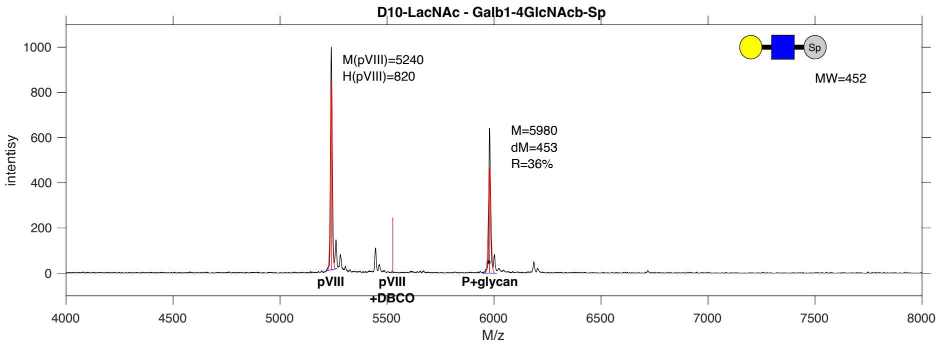


**61: MALDI-TOF spectra of glycosylated phage: LacNAc, LN-[970]**

**SDB Number:** SDB79

**Sequencing File:** http://ligacloud.ca/searchLibInfo?f=0&b=0&d=NA

**Barcode:**
CTACTGTTCGCAATACCTCTCAGCGTGGAGAAAAACGACCAGAAAACGTA
TCATGCTGGTGGTGGA

**Axis Name:** LacNAc, LN-[970]

**IUPAC:** Gal(b1-4)GlcNAc(b1-Sp

**IUPAC (CFG):** Galb1-4GlcNAcb-Sp

**Common Name:** LacNAc, LN

**Glytoucan ID:** NA

**Compound Number:** D10

**Maldi File:** MS-XIV-114_79.txt

**Density:** 36

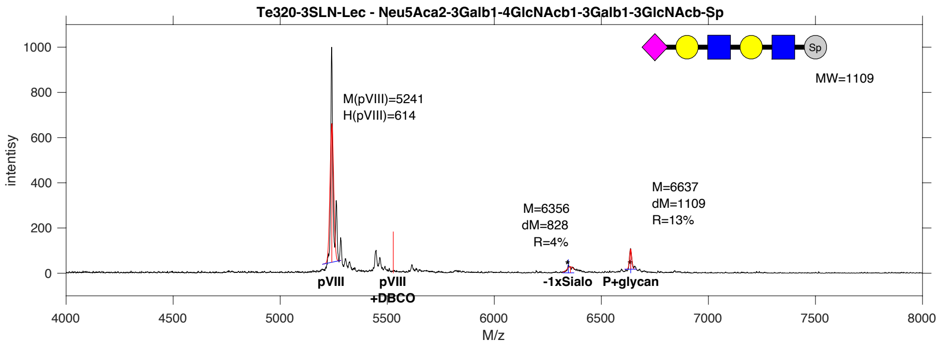


### 62: MALDI-TOF spectra of glycosylated phage: 3’SLN-Lec-[410]

**SDB Number:** SDB80

**Sequencing File:** http://ligacloud.ca/searchLibInfo?f=0&b=0&d=NA

**Barcode:**
CTTCTGTTCGCGATCCCCCTAAGTGTAGAGAAGAATGATCAGAAAACGTA
TCATGCGGGAGGGGGT

**Axis Name:** 3’SLN-Lec-[410]

**IUPAC:** Neu5Ac(a2-3)Gal(b1-4)GlcNAc(b1-3)Gal(b1-3)GlcNAc(b1-Sp

**IUPAC (CFG):** Neu5Aca2-3Galb1-4GlcNAcb1-3Galb1-3GlcNAcb-Sp

**Common Name:** 3’SLN-Lec

**Glytoucan ID:** NA

**Compound Number:** Te320

**Maldi File:** MS-XIV-114_80.txt

**Density:** 15

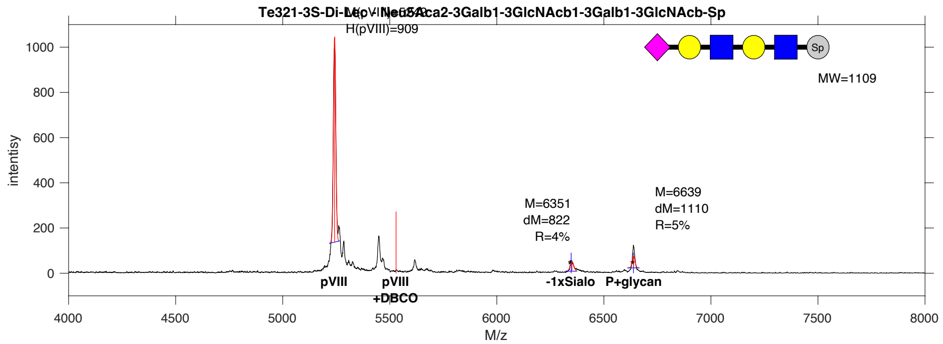


### 63: MALDI-TOF spectra of glycosylated phage: 3’S-Di-Lec-[270]

**SDB Number:** SDB83

**Sequencing File:** http://ligacloud.ca/searchLibInfo?f=0&b=0&d=NA

**Barcode:**
CTGCTCTTCGCGATCCCTCTGAGTGTCGAAAAGAATGATCAAAAAACGTA
TCATGCGGGCGGTGGT

**Axis Name:** 3’S-Di-Lec-[270]

**IUPAC:** Neu5Ac(a2-3)Gal(b1-3)GlcNAc(b1-3)Gal(b1-3)GlcNAc(b1-Sp

**IUPAC (CFG):** Neu5Aca2-3Galb1-3GlcNAcb1-3Galb1-3GlcNAcb-Sp

**Common Name:** 3’S-Di-Lec

**Glytoucan ID:** G11998FO

**Compound Number:** Te321

**Maldi File:** MS-XIV-114_83.txt

**Density:** 10

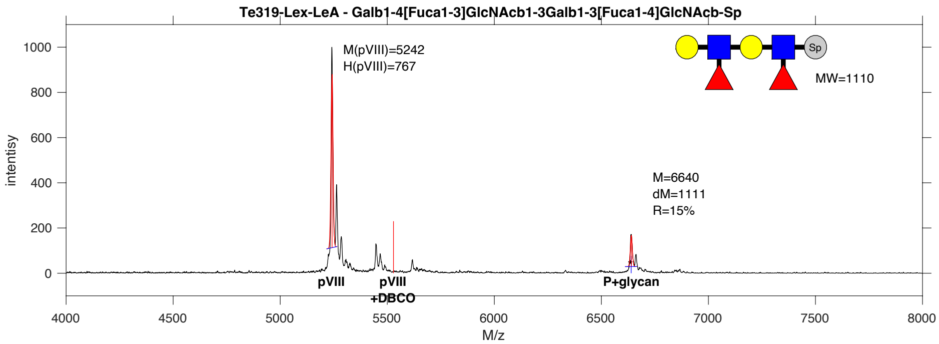


### 64: MALDI-TOF spectra of glycosylated phage: Lex-LeA-[410]

**SDB Number:** SDB84

**Sequencing File:** http://ligacloud.ca/searchLibInfo?f=0&b=0&d=NA

**Barcode:**
CTACTCTTCGCGATCCCACTGAGCGTCGAGAAGAACGATCAGAAGACTTA
TCACGCTGGCGGCGGC

**Axis Name:** Lex-LeA-[410]

**IUPAC:** Gal(b1-4)[Fuc(a1-3)]GlcNAc(b1-3)Gal(b1-3)[Fuc(a1-4)]GlcNAc(b1-Sp

**IUPAC (CFG):** Galb1-4[Fuca1-3]GlcNAcb1-3Galb1-3[Fuca1-4]GlcNAcb-Sp

**Common Name:** Lex-LeA

**Glytoucan ID:** G59754DB

**Compound Number:** Te319

**Maldi File:** MS-XIV-114_84.txt

**Density:** 15

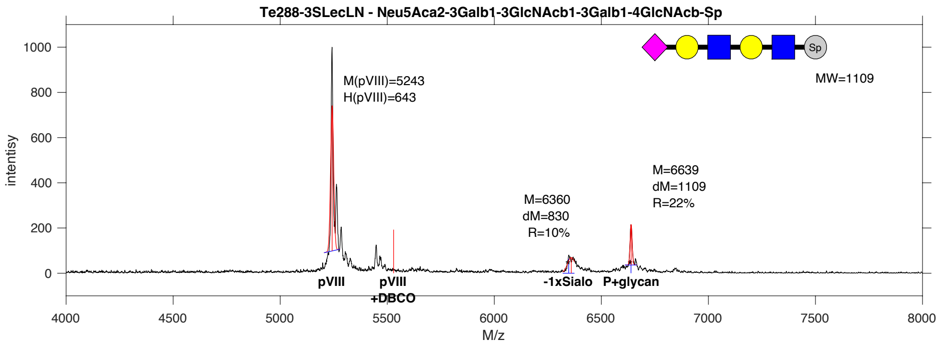


#### 65: MALDI-TOF spectra of glycosylated phage: 3’SLecLN-[860]

**SDB Number:** SDB88

**Sequencing File:** http://ligacloud.ca/searchLibInfo?f=0&b=0&d=NA

**Barcode:**
CTGCTGTTCGCGATTCCTCTAAGTGTAGAGAAGAATGACCAGAAGACCTA
CCACGCGGGCGGGGGG

**Axis Name:** 3’SLecLN-[860]

**IUPAC:** Neu5Ac(a2-3)Gal(b1-3)GlcNAc(b1-3)Gal(b1-4)GlcNAc(b1-Sp

**IUPAC (CFG):** Neu5Aca2-3Galb1-3GlcNAcb1-3Galb1-4GlcNAcb-Sp

**Common Name:** 3’SLecLN

**Glytoucan ID:** G78959US

**Compound Number:** Te288

**Maldi File:** MS-XIV-114_88.txt

**Density:** 32

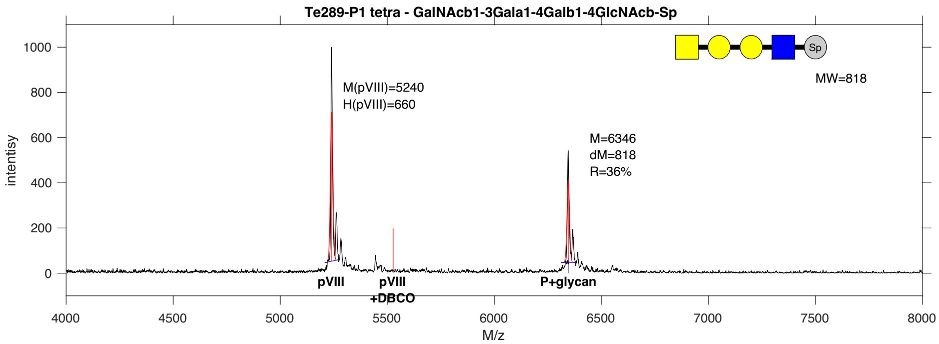


**66: MALDI-TOF spectra of glycosylated phage: P1 tetra-[970]**

**SDB Number:** SDB89

**Sequencing File:** http://ligacloud.ca/searchLibInfo?f=0&b=0&d=NA

**Barcode:**
CTACTGTTCGCAATACCTCTCAGCGTGGAGAAAAATGATCAAAAAACATA
TCATGCAGGGGGTGGC

**Axis Name:** P1 tetra-[970]

**IUPAC:** GalNAc(b1-3)Gal(a1-4)Gal(b1-4)GlcNAc(b1-Sp

**IUPAC (CFG):** GalNAcb1-3Gala1-4Galb1-4GlcNAcb-Sp

**Common Name:** P1 tetra

**Glytoucan ID:** G29548YX

**Compound Number:** Te289

**Maldi File:** MS-XIV-114_89.txt

**Density:** 36

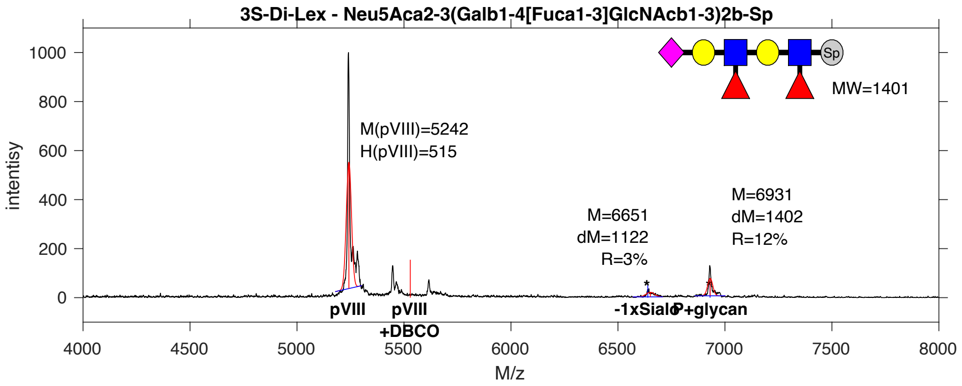


**67: MALDI-TOF spectra of glycosylated phage: 3’S-Di-Lex-[410]**

**SDB Number:** SDB13

**Sequencing File:** http://ligacloud.ca/searchLibInfo?f=0&b=0&d=NA

**Barcode:**
CTACTTTTCGCAATTCCTCTGAGTGTGGAGAAGAATGATCAGAAGACTTA
TCATGCGGGTGGAGGT

**Axis Name:** 3’S-Di-Lex-[410]

**IUPAC:** NANA

**IUPAC (CFG):** Neu5Aca2-3(Galb1-4[Fuca1-3]GlcNAcb1-3)2b-Sp

**Common Name:** 3’S-Di-Lex

**Glytoucan ID:** G19619KS

**Compound Number:** Te140

**Maldi File:** MS-XIV-157_13.txt

**Density:** 15

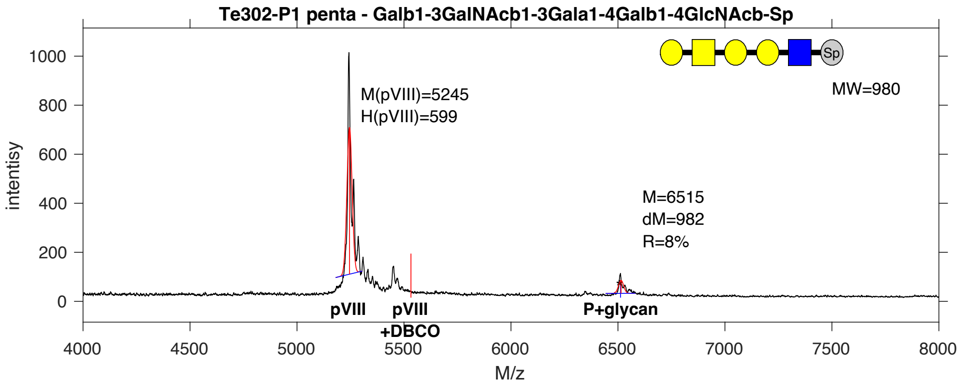


**68: MALDI-TOF spectra of glycosylated phage: P1.penta-[620]**

**SDB Number:** SDB23

**Sequencing File:** http://ligacloud.ca/searchLibInfo?f=0&b=0&d=NA

**Barcode:**
CTGCTCTTTGCAATACCTCTTAGTGTGGAGAAGAATGATCAGAAGACTTA
TCATGCGGGTGGAGGT

**Axis Name:** P1.penta-[620]

**IUPAC:** NANA

**IUPAC (CFG):** Galb1-3GalNAcb1-3Gala1-4Galb1-4GlcNAcb-Sp

**Common Name:** P1 penta

**Glytoucan ID:** G28005AF

**Compound Number:** Te302

**Maldi File:** MS-XIV-164_23.txt

**Density:** 23

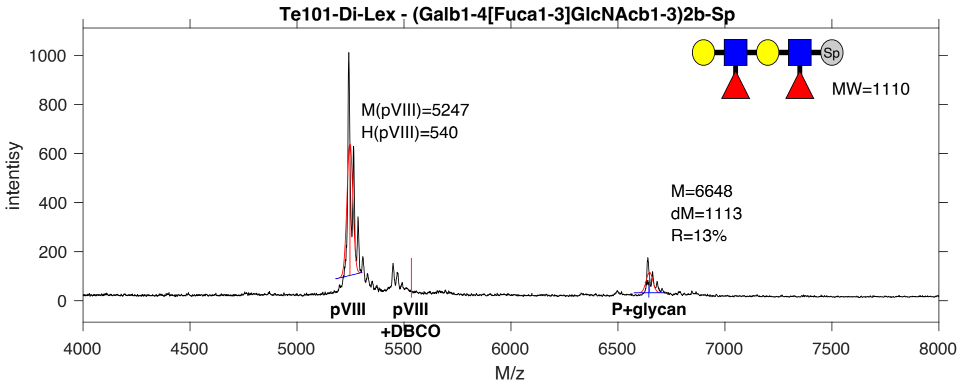


**69: MALDI-TOF spectra of glycosylated phage: Ley-Lex-[350]**

**SDB Number:** SDB26

**Sequencing File:** http://ligacloud.ca/searchLibInfo?f=0&b=0&d=NA

**Barcode:**
CTGCTATTCGCTATCCCACTCAGTGTGGAGAAGAATGATCAGAAGACTTA
TCATGCGGGTGGAGGT

**Axis Name:** Ley-Lex-[350]

**IUPAC:** NANA

**IUPAC (CFG):** Fuca1-2Galb1-4[Fuca1-3]GlcNAcb1-3Galb1-4[Fuca1-3]GlcNAcb-Sp

**Common Name:** Ley-Lex

**Glytoucan ID:** G74917JX

**Compound Number:** Te212

**Maldi File:** MS-XIV-164_26.txt

**Density:** 13

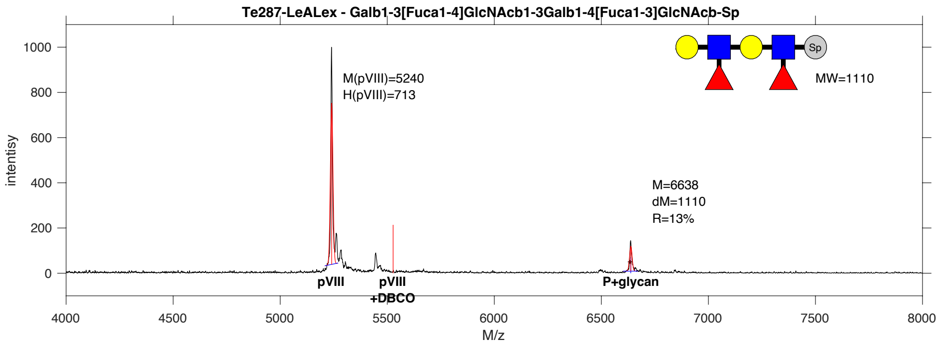


### 70: MALDI-TOF spectra of glycosylated phage: LeALex-[350]

**SDB Number:** SDB130

**Sequencing File:** http://ligacloud.ca/searchLibInfo?f=0&b=0&d=NA

**Barcode:**
CTTCTCTTTGCGATACCGCTAAGTGTGGAAAAGAATGATCAAAAAACGTA
CCATGCCGGTGGGGGT

**Axis Name:** LeALex-[350]

**IUPAC:** Gal(b1-3)[Fuc(a1-4)]GlcNAc(b1-3)Gal(b1-4)[Fuc(a1-3)]GlcNAc(b1-Sp

**IUPAC (CFG):** Galb1-3[Fuca1-4]GlcNAcb1-3Galb1-4[Fuca1-3]GlcNAcb-Sp

**Common Name:** LeALex

**Glytoucan ID:** NA

**Compound Number:** Te287

**Maldi File:** MS-XIV-114_130.txt

**Density:** 13

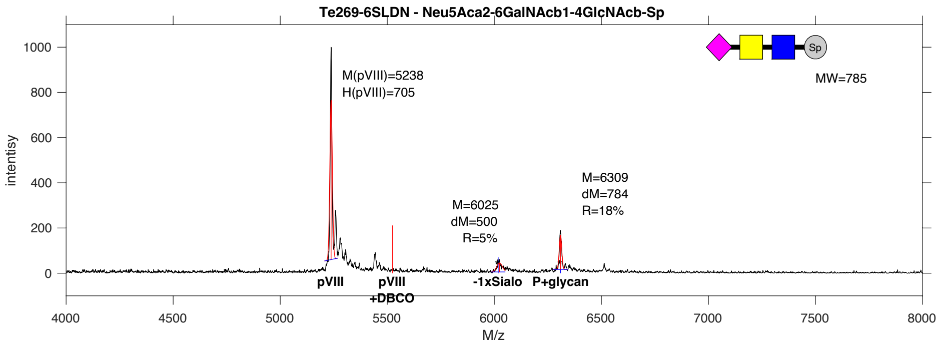


**71: MALDI-TOF spectra of glycosylated phage: 6’SLDN-[620]**

**SDB Number:** SDB131

**Sequencing File:** http://ligacloud.ca/searchLibInfo?f=0&b=0&d=NA

**Barcode:**
CTGCTGTTTGCGATTCCCCTGAGTGTCGAAAAGAATGACCAAAAGACGTA
CCATGCGGGGGGGGGG

**Axis Name:** 6’SLDN-[620]

**IUPAC:** Neu5Ac(a2-6)GalNAc(b1-4)GlcNAc(b1-Sp

**IUPAC (CFG):** Neu5Aca2-6GalNAcb1-4GlcNAcb-Sp

**Common Name:** 6’SLDN

**Glytoucan ID:** G54118CT

**Compound Number:** Tr269

**Maldi File:** MS-XIV-114_131.txt

**Density:** 23

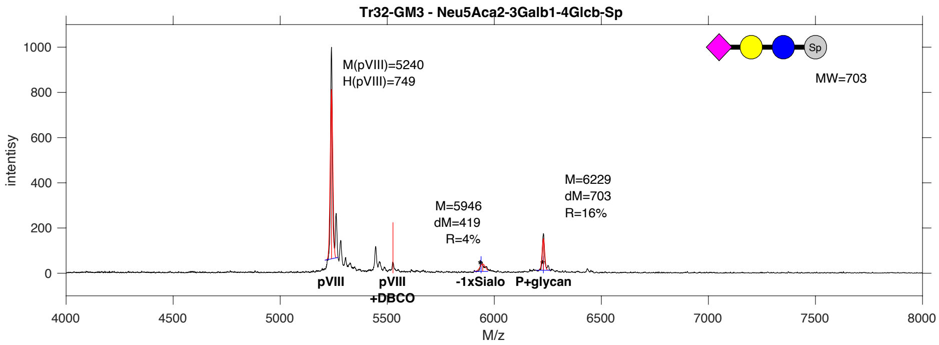


**72: MALDI-TOF spectra of glycosylated phage: GM3-[540]**

**SDB Number:** SDB132

**Sequencing File:** http://ligacloud.ca/searchLibInfo?f=0&b=0&d=NA

**Barcode:**
CTACTGTTCGCAATACCTCTCAGCGTAGAAAAAAATGATCAAAAAACATA
TCACGCAGGTGGCGGA

**Axis Name:** GM3-[540]

**IUPAC:** Neu5Ac(a2-3)Gal(b1-4)Glc(b1-Sp

**IUPAC (CFG):** Neu5Aca2-3Galb1-4Glcb-Sp

**Common Name:** GM3

**Glytoucan ID:** G91237TK

**Compound Number:** Tr32

**Maldi File:** MS-XIV-114_132.txt

**Density:** 20

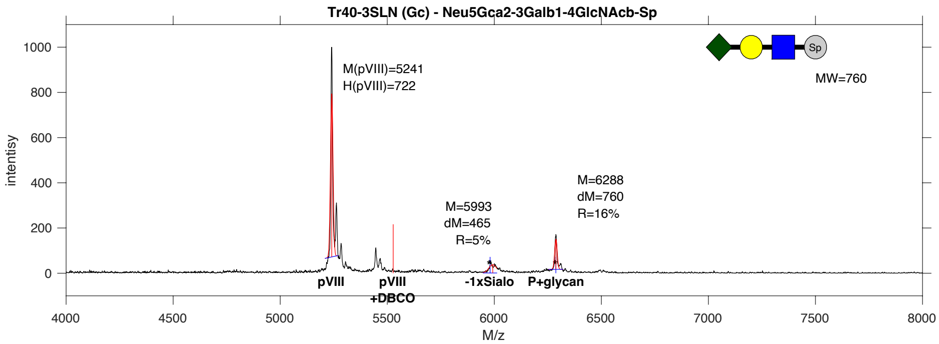


### 73: MALDI-TOF spectra of glycosylated phage: 3’SLN (Gc)-[570]

**SDB Number:** SDB141

**Sequencing File:** http://ligacloud.ca/searchLibInfo?f=0&b=0&d=20210330-87CLooJM-MS

**Barcode:** CTTCTTTTTGCCATACCCCTGAGCGTAGAAAAAAACGACCAGAAAACATATCATGCTGGCGGCGGT

**Axis Name:** 3’SLN (Gc)-[570]

**IUPAC:** Neu5Gc(a2-3)Gal(b1-4)GlcNAc(b1-Sp

**IUPAC (CFG):** Neu5Gca2-3Galb1-4GlcNAcb-Sp

**Common Name:** 3’SLN (Gc)

**Glytoucan ID:** G08169IN

**Compound Number:** Tr40

**Maldi File:** MS-XIV-114_141.txt

**Density:** 21

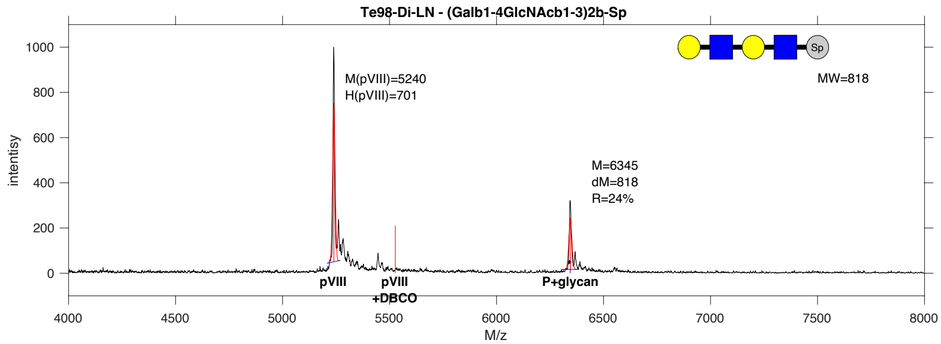


**74: MALDI-TOF spectra of glycosylated phage: Di-LN-[650]**

**SDB Number:** SDB186

**Sequencing File:** http://ligacloud.ca/searchLibInfo?f=0&b=0&d=20210330-87CLooCP-MS

**Barcode:** CTACTGTTCGCAATCCCGCTAAGTGTTGAGAAAAACGATCAAAAGACTTATCATGCAGGCGGAGGA

**Axis Name:** Di-LN-[650]

**IUPAC:** Gal(b1-4)GlcNAc(b1-3)Gal(b1-4)GlcNAc(b1-Sp

**IUPAC (CFG):** (Galb1-4GlcNAcb1-3)2b-Sp

**Common Name:** Di-LN

**Glytoucan ID:** G64789TL

**Compound Number:** Te98

**Maldi File:** MS-XIV-114_186.txt

**Density:** 24

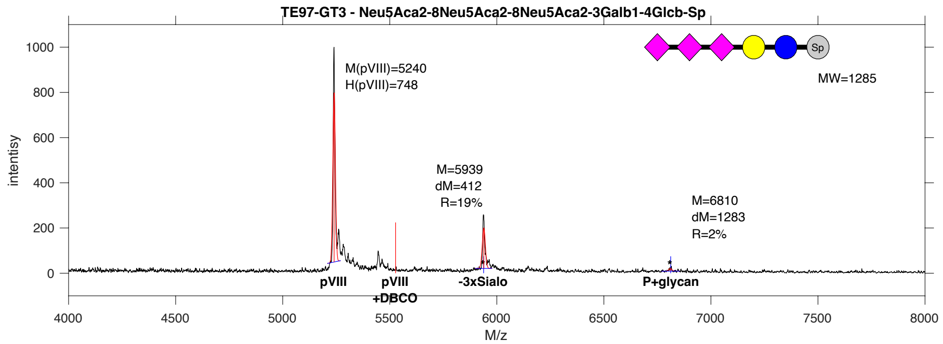


#### 75: MALDI-TOF spectra of glycosylated phage: GT3-[510]

**SDB Number:** SDB201

**Sequencing File:** http://ligacloud.ca/searchLibInfo?f=0&b=0&d=20210330-87CLooGE-MS

**Barcode:** CTACTCTTTGCTATTCCACTAAGTGTGGAGAAAAATGATCAAAAGACGTACCATGCGGGTGGCGGT

**Axis Name:** GT3-[510]

**IUPAC:** Neu5Ac(a2-8)Neu5Ac(a2-8)Neu5Ac(a2-3)Gal(b1-4)Glc(b1-Sp

**IUPAC (CFG):** Neu5Aca2-8Neu5Aca2-8Neu5Aca2-3Galb1-4Glcb-Sp

**Common Name:** GT3

**Glytoucan ID:** G93899SO

**Compound Number:** Te97

**Maldi File:** MS-XIV-114_201.txt

**Density:** 19

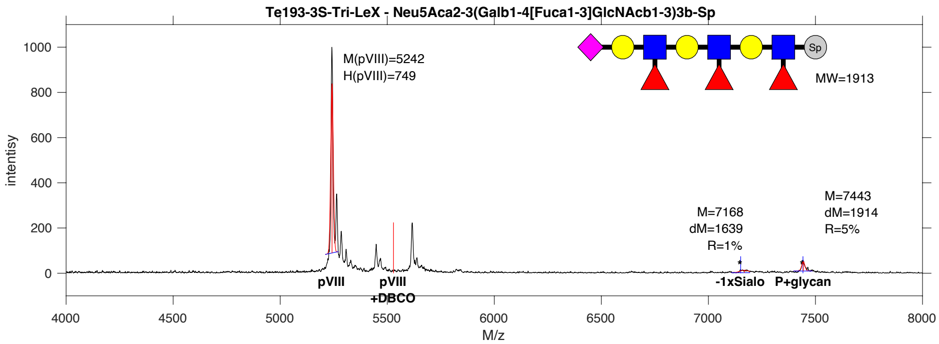


### 76: MALDI-TOF spectra of glycosylated phage: 3’S-Tri-LeX-[160]

**SDB Number:** SDB218

**Sequencing File:** http://ligacloud.ca/searchLibInfo?f=0&b=0&d=20210330-87CLooNM-MS

**Barcode:** TTATTATTCGCAATTCCTTTAAGTGTGGAAAAAAATGATCAAAAGACATACCACGCGGGGGGCGGG

**Axis Name:** 3’S-Tri-LeX-[160]

**IUPAC:** Neu5Ac(a2-3)Gal(b1-4)[Fuc(a1-3)]GlcNAc(b1-3)Gal(b1-4)[Fuc(a1-3)]GlcNAc(b1-3)Gal(b1-4)[Fuc(a1-3)]GlcNAc(b1-Sp

**IUPAC (CFG):** Neu5Aca2-3(Galb1-4[Fuca1-3]GlcNAcb1-3)3b-Sp

**Common Name:** 3’S-Tri-LeX

**Glytoucan ID:** G70868ZC

**Compound Number:** Te193

**Maldi File:** MS-XIV-114_218.txt

**Density:** 6

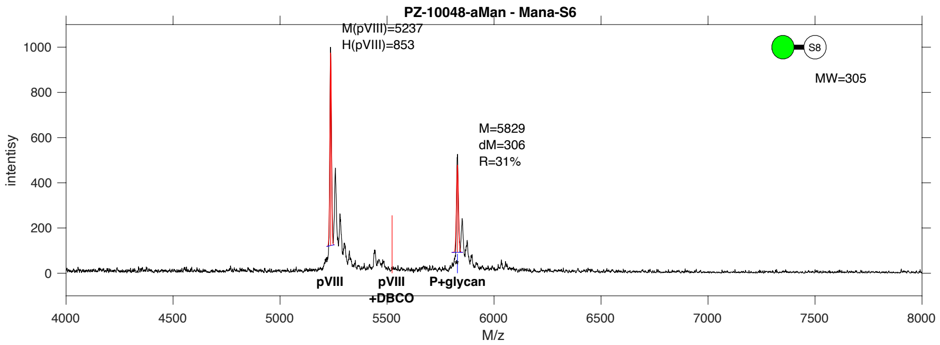


### 77: MALDI-TOF spectra of glycosylated phage: aMan-[840]

**SDB Number:** SDB217

**Sequencing File:** http://ligacloud.ca/searchLibInfo?f=0&b=0&d=20210330-87CLooEK-MS

**Barcode:** TTATTATTCGCAATTCCTTTAAGTGTGGAAAAGAACGATCAAAAAACTTATCACGCGGGTGGGGGC

**Axis Name:** aMan-[840]

**IUPAC:** Man(a1-S6

**IUPAC (CFG):** Mana-S6

**Common Name:** aMan

**Glytoucan ID:** G19165TD

**Compound Number:** PZ-10048

**Maldi File:** MS-XIV-114_219.txt

**Density:** 31

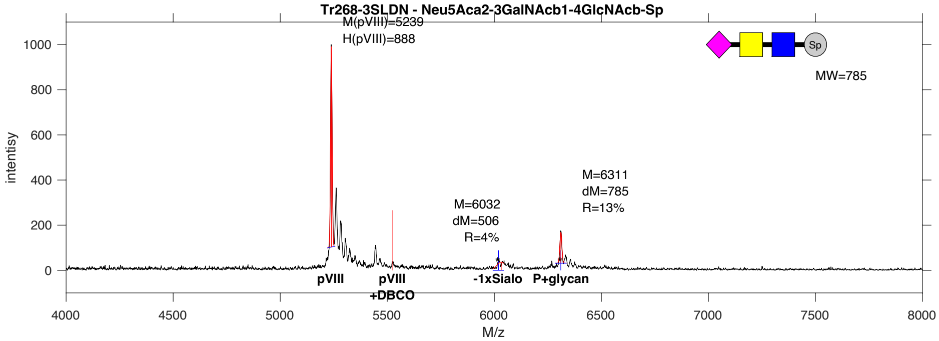


#### 78: MALDI-TOF spectra of glycosylated phage: 3’SLDN-[460]

**SDB Number:** SDB220

**Sequencing File:** http://ligacloud.ca/searchLibInfo?f=0&b=0&d=20210330-87CLooZK-MS

**Barcode:** CTGCTTTTTGCTATTCCTCTGAGTGTGGAAAAGAACGATCAGAAGACGTATCATGCCGGAGGAGGT

**Axis Name:** 3’SLDN-[460]

**IUPAC:** Neu5Ac(a2-3)GalNAc(b1-4)GlcNAc(b1-Sp

**IUPAC (CFG):** Neu5Aca2-3GalNAcb1-4GlcNAcb-Sp

**Common Name:** 3’SLDN

**Glytoucan ID:** NA

**Compound Number:** Tr268

**Maldi File:** MS-XIV-114_220.txt

**Density:** 17

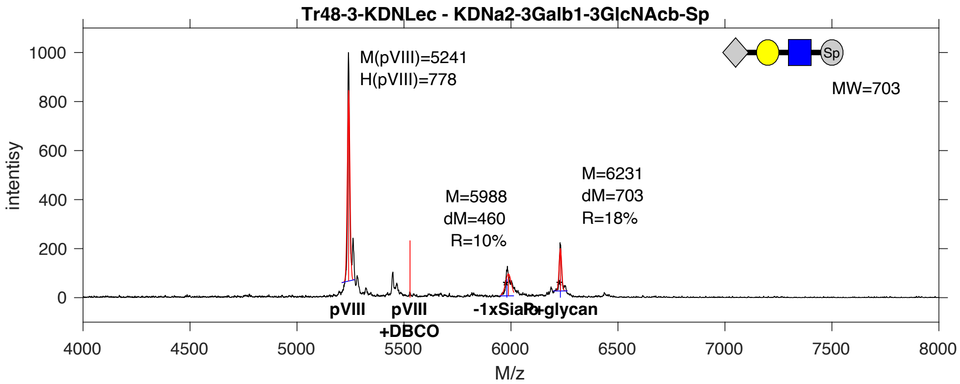


**79: MALDI-TOF spectra of glycosylated phage: 3’-KDNLec-[760]**

**SDB Number:** SDB150

**Sequencing File:** http://ligacloud.ca/searchLibInfo?f=0&b=0&d=NA

**Barcode:**
CTGCTCTTTGCTATCCCTCTAAGTGTTGAAAAAAACGACCAGAAGACTTAT
CACGCAGGTGGTGGG

**Axis Name:** 3’-KDNLec-[760]

**IUPAC:** NANA

**IUPAC (CFG):** KDNa2-3Galb1-3GlcNAcb-Sp

**Common Name:** 3’-KDNLec

**Glytoucan ID:** G13678CX

**Compound Number:** Tr48

**Maldi File:** MS-XIV-157_150.txt

**Density:** 28

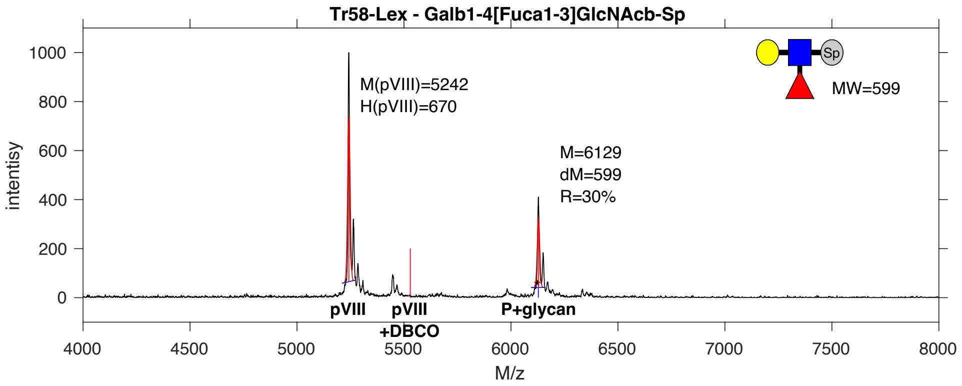


**80: MALDI-TOF spectra of glycosylated phage: Lex-[810]**

**SDB Number:** SDB151

**Sequencing File:** http://ligacloud.ca/searchLibInfo?f=0&b=0&d=NA

**Barcode:**
CTGCTGTTTGCGATCCCGCTTAGCGTAGAAAAGAATGACCAGAAGACATA
CCACGCGGGAGGGGGG

**Axis Name:** Lex-[810]

**IUPAC:** Gal(b1-4)[Fuc(a1-3)]GlcNAc(b-Sp

**IUPAC (CFG):** Galb1-4[Fuca1-3]GlcNAcb-Sp

**Common Name:** Lex

**Glytoucan ID:** NA

**Compound Number:** Tr58

**Maldi File:** MS-XIV-157_151.txt

**Density:** 30

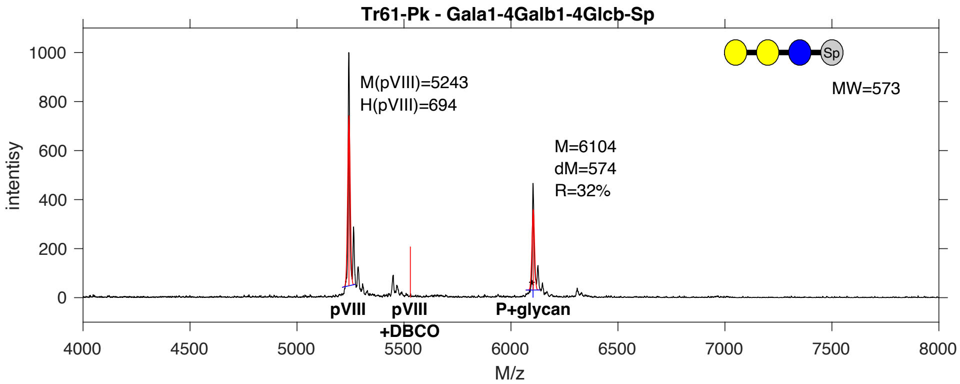


**81: MALDI-TOF spectra of glycosylated phage: Pk-[860]**

**SDB Number:** SDB152

**Sequencing File:** http://ligacloud.ca/searchLibInfo?f=0&b=0&d=NA

**Barcode:**
CTGCTTTTTGCTATACCTCTCAGCGTAGAAAAGAATGACCAGAAGACGTA
CCATGCCGGTGGGGGG

**Axis Name:** Pk-[860]

**IUPAC:** Gal(a1-4)Gal(b1-4)Glc(bSp

**IUPAC (CFG):** Gala1-4Galb1-4Glcb-Sp

**Common Name:** Pk

**Glytoucan ID:** NA

**Compound Number:** Tr61

**Maldi File:** MS-XIV-157_152.txt

**Density:** 32

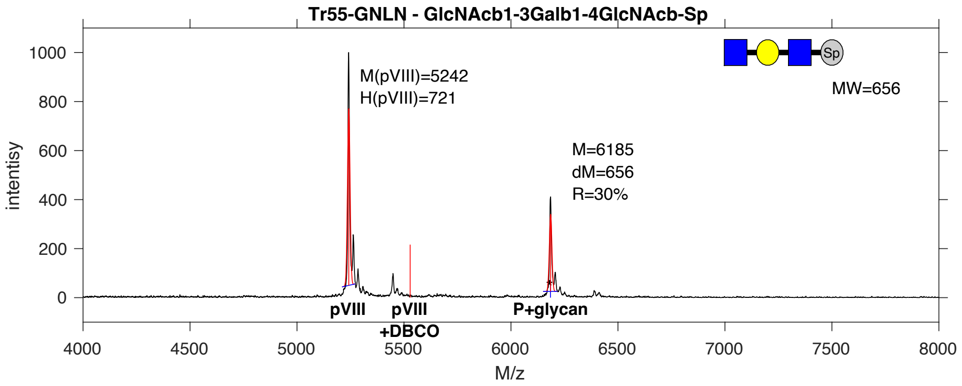


**82: MALDI-TOF spectra of glycosylated phage: GNLN-[810]**

**SDB Number:** SDB153

**Sequencing File:** http://ligacloud.ca/searchLibInfo?f=0&b=0&d=NA

**Barcode:**
CTACTATTCGCTATTCCTCTGAGTGTCGAGAAAAACGATCAAAAGACCTA
TCATGCGGGTGGGGGT

**Axis Name:** GNLN-[810]

**IUPAC:** NANA

**IUPAC (CFG):** GlcNAcb1-3Galb1-4GlcNAcb-Sp

**Common Name:** GNLN

**Glytoucan ID:** G29487JS

**Compound Number:** Tr55

**Maldi File:** MS-XIV-157_153.txt

**Density:** 30

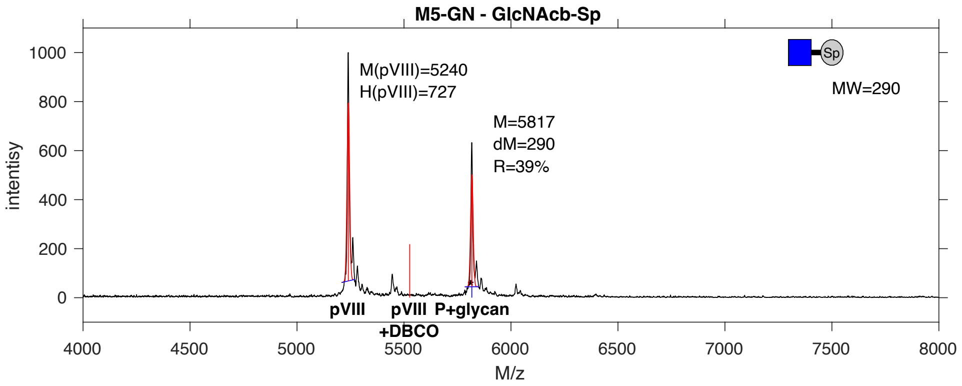


**83: MALDI-TOF spectra of glycosylated phage: GN-[1050]**

**SDB Number:** SDB154

**Sequencing File:** http://ligacloud.ca/searchLibInfo?f=0&b=0&d=NA

**Barcode:**
CTACTGTTCGCTATTCCGCTAAGTGTAGAAAAAAACGATCAAAAGACGTA
TCACGCCGGTGGGGGA

**Axis Name:** GN-[1050]

**IUPAC:** GlcNAc(bSp

**IUPAC (CFG):** GlcNAcb-Sp

**Common Name:** GN

**Glytoucan ID:** NA

**Compound Number:** M5

**Maldi File:** MS-XIV-157_154.txt

**Density:** 39

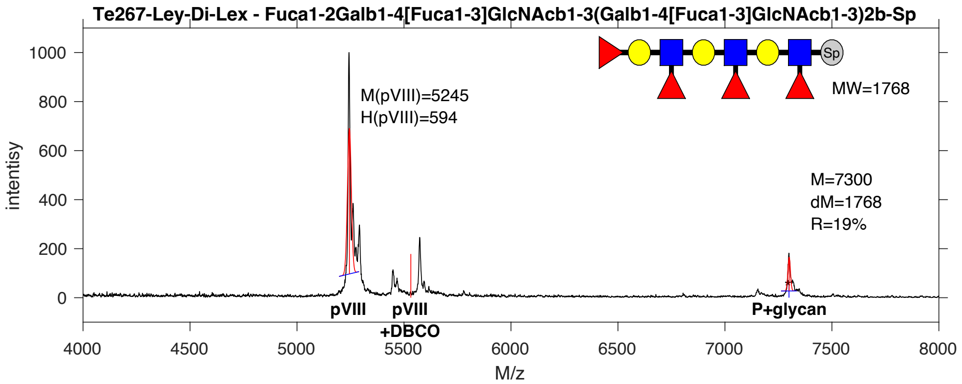


**84: MALDI-TOF spectra of glycosylated phage: Ley-Di-Lex-[510]**

**SDB Number:** SDB160

**Sequencing File:** http://ligacloud.ca/searchLibInfo?f=0&b=0&d=NA

**Barcode:**
TTATTATTCGCAATTCCTTTAAGTGTCGAAAAAAATGACCAGAAAACTTACCATGCTGGAGGGGGC

**Axis Name:** Ley-Di-Lex-[510]

**IUPAC:** NANA

**IUPAC (CFG):** Fuca1-2Galb1-4[Fuca1-3]GlcNAcb1-3(Galb1-4[Fuca1-3]GlcNAcb1-3)2b-Sp

**Common Name:** Ley-Di-Lex

**Glytoucan ID:** Na

**Compound Number:** Te267

**Maldi File:** MS-XIV-157_160.txt

**Density:** 19

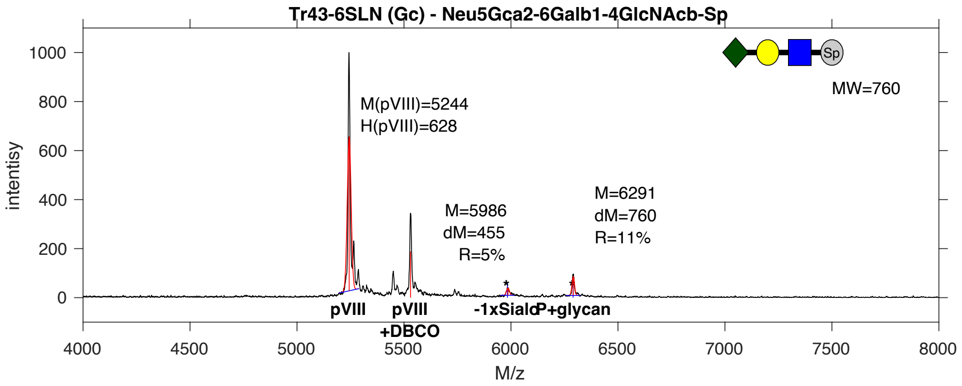


**85: MALDI-TOF spectra of glycosylated phage: 6’SLN (Gc)-[430]**

**SDB Number:** SDB161

**Sequencing File:** http://ligacloud.ca/searchLibInfo?f=0&b=0&d=NA

**Barcode:**
CTGCTATTCGCTATCCCCCTGAGTGTTGAGAAGAATGACCAAAAGACATA
TCACGCCGGGGGAGGG

**Axis Name:** 6’SLN (Gc)-[430]

**IUPAC:** NANA

**IUPAC (CFG):** Neu5Gca2-6Galb1-4GlcNAcb-Sp

**Common Name:** 6’SLN (Gc)

**Glytoucan ID:** G24684AO

**Compound Number:** Tr43

**Maldi File:** MS-XIV-157_161.txt

**Density:** 16

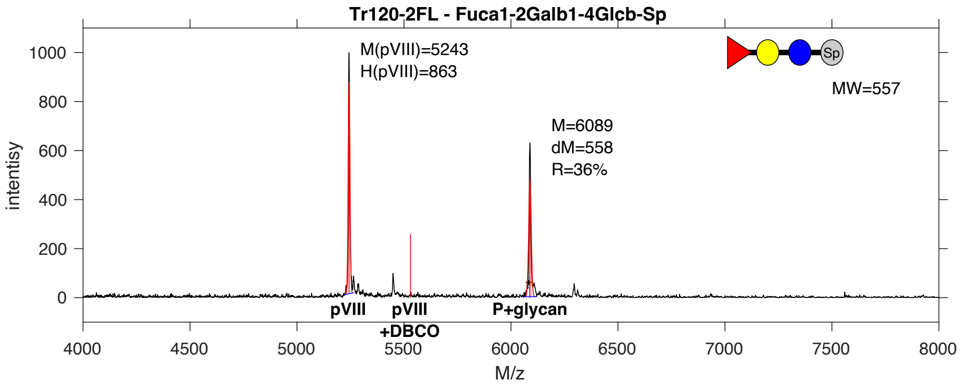


**86: MALDI-TOF spectra of glycosylated phage: 2’FL-[950]**

**SDB Number:** SDB167

**Sequencing File:** http://ligacloud.ca/searchLibInfo?f=0&b=0&d=NA

**Barcode:**
CTGCTTTTTGCTATTCCTCTGAGTGTCGAAAAAAACGATCAGAAGACGTAT CACGCAGGTGGTGGT

**Axis Name:** 2’FL-[950]

**IUPAC:** NANA

**IUPAC (CFG):** Fuca1-2Galb1-4Glcb-Sp

**Common Name:** 2’FL

**Glytoucan ID:** G50519XG

**Compound Number:** Tr120

**Maldi File:** MS-XIV-157_167.txt

**Density:** 35

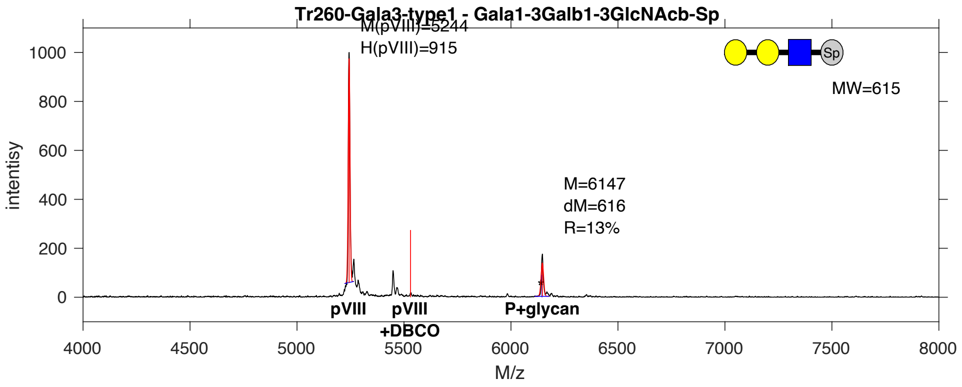


**87: MALDI-TOF spectra of glycosylated phage: Gala3-type1-[350]**

**SDB Number:** SDB168

**Sequencing File:** http://ligacloud.ca/searchLibInfo?f=0&b=0&d=NA

**Barcode:**
TTATTATTCGCAATTCCTTTAAGCGTGGAAAAAAATGATCAAAAGACATA
TCACGCTGGTGGTGGG

**Axis Name:** Gala3-type1-[350]

**IUPAC:** NANA

**IUPAC (CFG):** Gala1-3Galb1-3GlcNAcb-Sp

**Common Name:** Gala3-type1

**Glytoucan ID:** G34356AA

**Compound Number:** Tr260

**Maldi File:** MS-XIV-157_168.txt

**Density:** 13

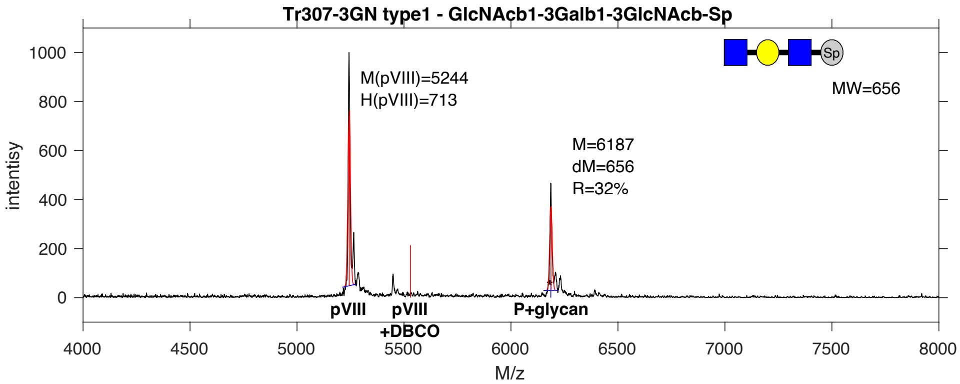


**88: MALDI-TOF spectra of glycosylated phage: 3’GN.type1-[860]**

**SDB Number:** SDB169

**Sequencing File:** http://ligacloud.ca/searchLibInfo?f=0&b=0&d=NA

**Barcode:**
CTGCTGTTCGCGATTCCCCTAAGCGTCGAAAAGAATGACCAAAAGACGTA
TCACGCTGGCGGGGGG

**Axis Name:** 3’GN.type1-[860]

**IUPAC:** GlcNAc(b1-3)Gal(b1-3)GlcNAc(b-Sp

**IUPAC (CFG):** GlcNAcb1-3Galb1-3GlcNAcb-Sp

**Common Name:** 3’GN type1

**Glytoucan ID:** NA

**Compound Number:** Tr307

**Maldi File:** MS-XIV-157_169.txt

**Density:** 32

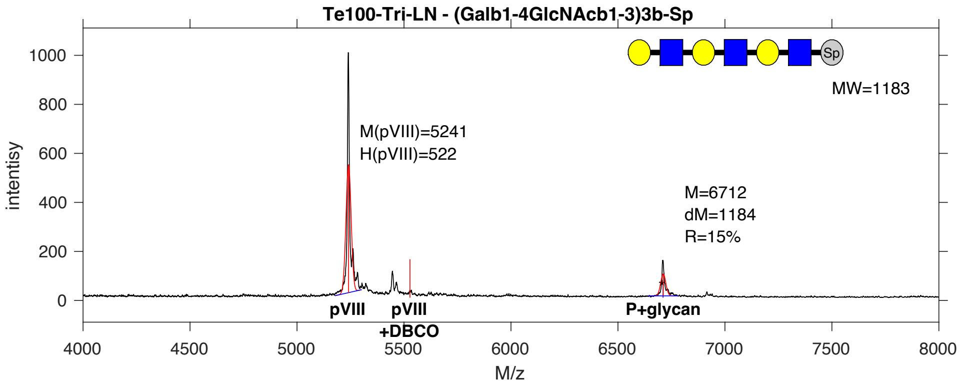


**89: MALDI-TOF spectra of glycosylated phage: Di-Lex-[410]**

**SDB Number:** SDB111

**Sequencing File:** http://ligacloud.ca/searchLibInfo?f=0&b=0&d=NA

**Barcode:**
CTGCTGTTTGCTATCCCGCTAAGTGTAGAAAAAAATGACCAGAAAACGTA
CCATGCGGGCGGTGGG

**Axis Name:** Di-Lex-[410]

**IUPAC:** NANA

**IUPAC (CFG):** (Galb1-4[Fuca1-3]GlcNAcb1-3)2b-Sp

**Common Name:** Di-Lex

**Glytoucan ID:** G79535HZ

**Compound Number:** Te101

**Maldi File:** MS-XIV-164_111.txt

**Density:** 15

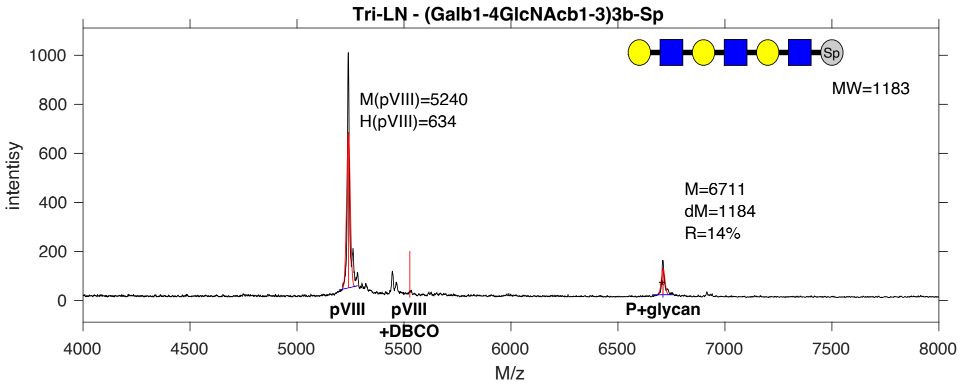


**90: MALDI-TOF spectra of glycosylated phage: Tri-LN-[380]**

**SDB Number:** SDB112

**Sequencing File:** http://ligacloud.ca/searchLibInfo?f=0&b=0&d=NA

**Barcode:**
CTACTGTTCGCGATCCCACTGAGTGTAGAAAAAAATGACCAGAAGACGTA
TCACGCAGGCGGAGGC

**Axis Name:** Tri-LN-[380]

**IUPAC:** NANA

**IUPAC (CFG):** (Galb1-4GlcNAcb1-3)3b-Sp

**Common Name:** Tri-LN

**Glytoucan ID:** G46055MA

**Compound Number:** Te100

**Maldi File:** MS-XIV-164_112.txt

**Density:** 14

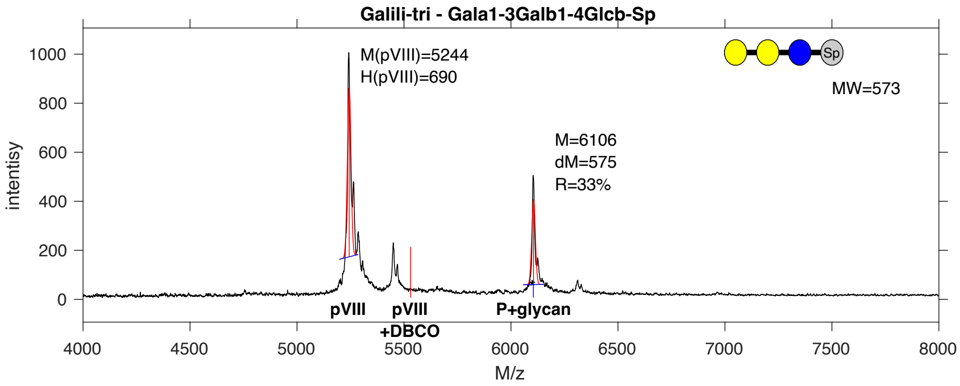


**91: MALDI-TOF spectra of glycosylated phage: Galili-tri-[1000]**

**SDB Number:** SDB113

**Sequencing File:** http://ligacloud.ca/searchLibInfo?f=0&b=0&d=NA

**Barcode:**
CTACTGTTTGCCATTCCCCTGAGTGTCGAAAAAAATGATCAGAAAACTTA
TCACGCAGGAGGGGGG

**Axis Name:** Galili-tri-[1000]

**IUPAC:** NANA

**IUPAC (CFG):** Gala1-3Galb1-4Glcb-Sp

**Common Name:** Galili-tri

**Glytoucan ID:** G78074HX

**Compound Number:** Tr59

**Maldi File:** MS-XIV-164_113.txt

**Density:** 37

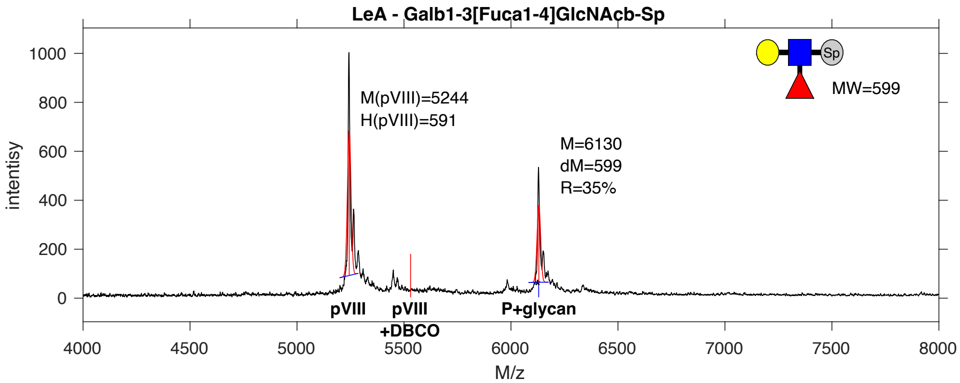


**92: MALDI-TOF spectra of glycosylated phage: LeA-[950]**

**SDB Number:** SDB114

**Sequencing File:** http://ligacloud.ca/searchLibInfo?f=0&b=0&d=NA

**Barcode:**
CTACTGTTTGCGATCCCACTCAGCGTGGAGAAGAACGACCAGAAGACCTA
CCACGCGGGGGGAGGG

**Axis Name:** LeA-[950]

**IUPAC:** NANA

**IUPAC (CFG):** Galb1-3[Fuca1-4]GlcNAcb-Sp

**Common Name:** LeA

**Glytoucan ID:** G39023AU

**Compound Number:** Tr57

**Maldi File:** MS-XIV-164_114.txt

**Density:** 35

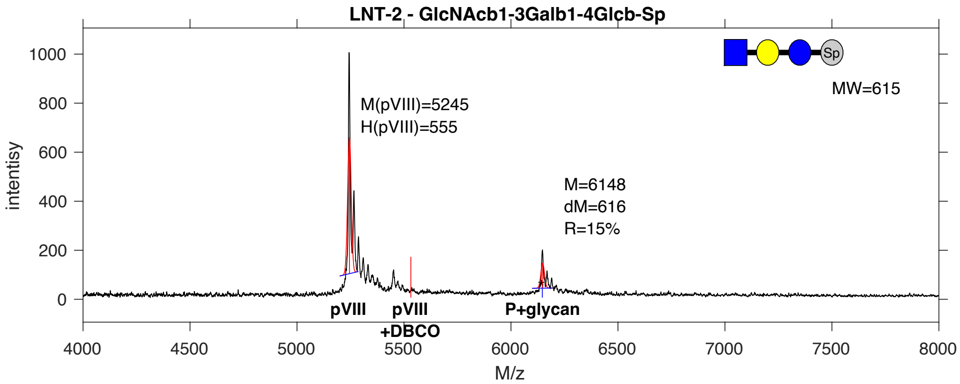


**93: MALDI-TOF spectra of glycosylated phage: LNT-2-[430]**

**SDB Number:** SDB115

**Sequencing File:** http://ligacloud.ca/searchLibInfo?f=0&b=0&d=NA

**Barcode:**
CTACTGTTCGCTATCCCGCTGAGTGTCGAAAAAAATGACCAGAAAACGTA
TCATGCGGGGGGGGGT

**Axis Name:** LNT-2-[430]

**IUPAC:** NANA

**IUPAC (CFG):** GlcNAcb1-3Galb1-4Glcb-Sp

**Common Name:** LNT-2

**Glytoucan ID:** G64979HM

**Compound Number:** Tr54

**Maldi File:** MS-XIV-164_115.txt

**Density:** 16

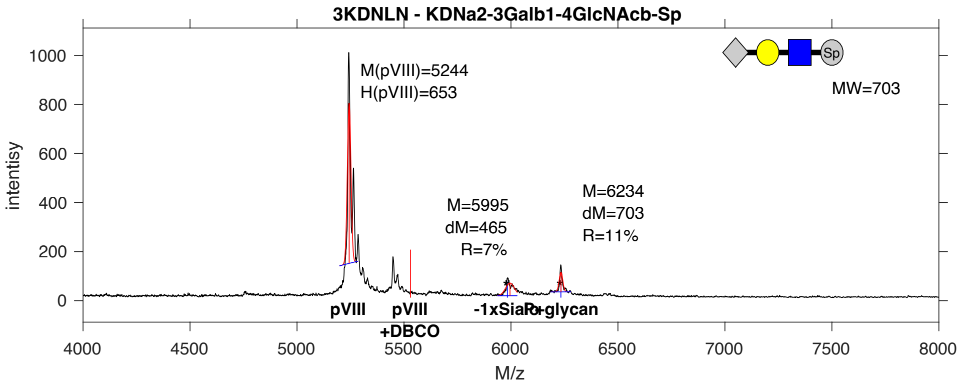


**94: MALDI-TOF spectra of glycosylated phage: 3’KDNLN-[510]**

**SDB Number:** SDB117

**Sequencing File:** http://ligacloud.ca/searchLibInfo?f=0&b=0&d=NA

**Barcode:**
TTATTATTCGCAATTCCTTTAAGCGTTGAAAAGAATGACCAGAAAACGTACCACGCGGGGGGCGGA

**Axis Name:** 3’KDNLN-[510]

**IUPAC:** NANA

**IUPAC (CFG):** KDNa2-3Galb1-4GlcNAcb-Sp

**Common Name:** 3’KDNLN

**Glytoucan ID:** G61010HE

**Compound Number:** Tr47

**Maldi File:** MS-XIV-164_117.txt

**Density:** 19

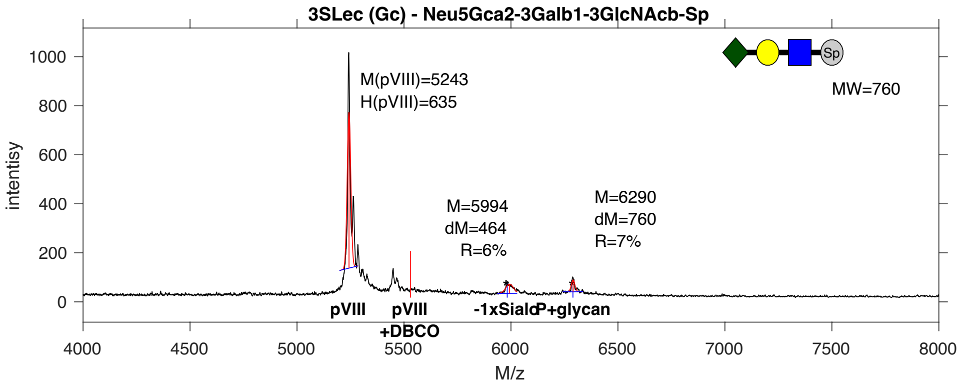


**95: MALDI-TOF spectra of glycosylated phage: 3’SLec (Gc)-[350]**

**SDB Number:** SDB118

**Sequencing File:** http://ligacloud.ca/searchLibInfo?f=0&b=0&d=NA

**Barcode:**
CTACTCTTCGCGATACCCCTCAGTGTCGAAAAGAATGATCAGAAGACTTA
TCATGCAGGTGGTGGT

**Axis Name:** 3’SLec (Gc)-[350]

**IUPAC:** NANA

**IUPAC (CFG):** Neu5Gca2-3Galb1-3GlcNAcb-Sp

**Common Name:** 3’SLec (Gc)

**Glytoucan ID:** G01245TU

**Compound Number:** Tr41

**Maldi File:** MS-XIV-164_118.txt

**Density:** 13

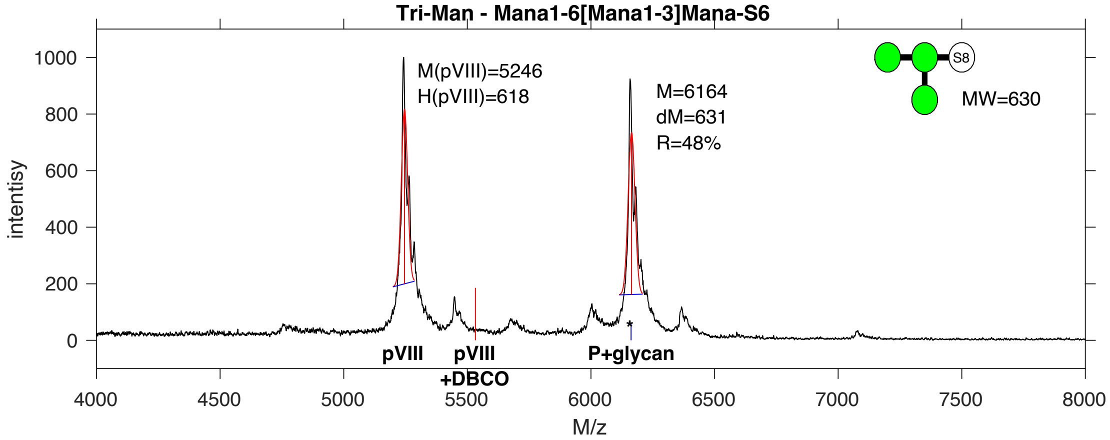


**96: MALDI-TOF spectra of glycosylated phage: (Man)3-[1300]**

**SDB Number:** SDB121

**Sequencing File:** http://ligacloud.ca/searchLibInfo?f=0&b=0&d=NA

**Barcode:**
CTGCTGTTTGCGATACCGCTTAGTGTGGAGAAAAACGATCAGAAGACCTA
TCATGCTGGGGGTGGA

**Axis Name:** (Man)3-[1300]

**IUPAC:** Man(a1-6)[Man(a1-3)]Man(a1-S6

**IUPAC (CFG):** Mana1-6[Mana1-3]Mana-S6

**Common Name:** (Man)3

**Glytoucan ID:** G80926OA

**Compound Number:** PZ-8015

**Maldi File:** MS-XIV-164_121.txt

**Density:** 48

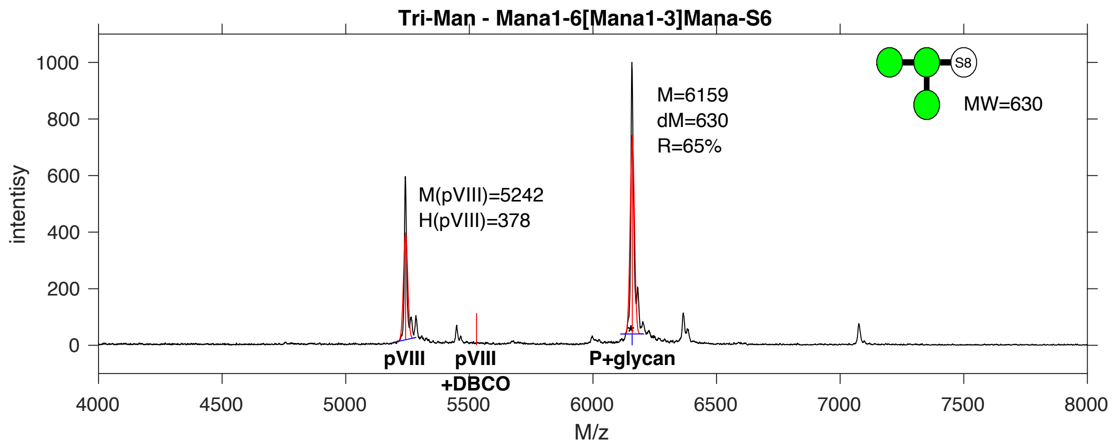


**97: MALDI-TOF spectra of glycosylated phage: (Man)3-[1730]**

**SDB Number:** SDB122

**Sequencing File:** http://ligacloud.ca/searchLibInfo?f=0&b=0&d=NA

**Barcode:**
CTGCTTTTTGCTATACCCCTGAGCGTGGAGAAGAACGATCAAAAGACCTA TCACGCGGGGGGAGGC

**Axis Name:** (Man)3-[1730]

**IUPAC:** Man(a1-6)[Man(a1-3)]Man(a1-S6

**IUPAC (CFG):** Mana1-6[Mana1-3]Mana-S6

**Common Name:** (Man)3

**Glytoucan ID:** G80926OA

**Compound Number:** PZ-8015

**Maldi File:** MS-XIV-164_122.txt

**Density:** 64

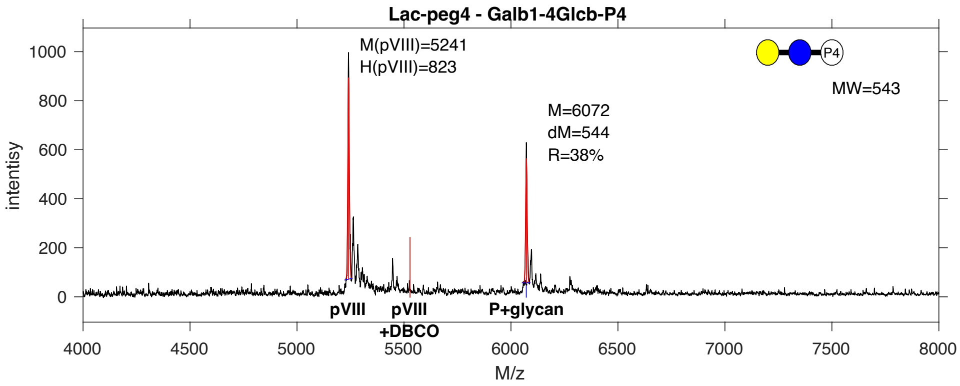


**98: MALDI-TOF spectra of glycosylated phage: Lac-peg4-[1080]**

**SDB Number:** SDB123

**Sequencing File:** http://ligacloud.ca/searchLibInfo?f=0&b=0&d=NA

**Barcode:**
NTGCTATTTGCCATCCCACTAAGCGTGGAAAAGAATGATCAGAAGACATA
TCATGCCGGCGGAGGT

**Axis Name:** Lac-peg4-[1080]

**IUPAC:** NANA

**IUPAC (CFG):** Galb1-4Glcb-P4

**Common Name:** Lac-peg4

**Glytoucan ID:** G94144EF

**Compound Number:** PZ-5080

**Maldi File:** MS-XIV-164_123.txt

**Density:** 40

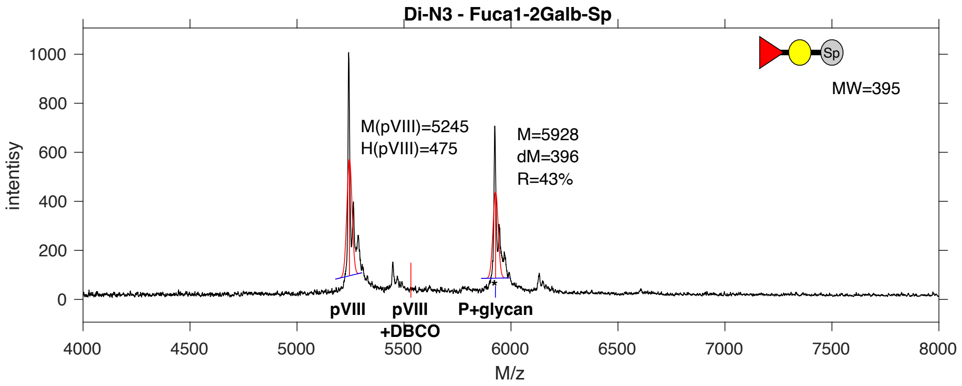


**99: MALDI-TOF spectra of glycosylated phage: Di-N3-[970]**

**SDB Number:** SDB125

**Sequencing File:** http://ligacloud.ca/searchLibInfo?f=0&b=0&d=NA

**Barcode:**
CTGCTATTCGCTATCCCGCTGAGTGTAGAAAAAAACGACCAGAAAACCTA
TCATGCCGGTGGCGGG

**Axis Name:** Di-N3-[970]

**IUPAC:** NANA

**IUPAC (CFG):** Fuca1-2Galb-Sp

**Common Name:** Di-N3

**Glytoucan ID:** G00068MO

**Compound Number:** Di-N3

**Maldi File:** MS-XIV-164_125.txt

**Density:** 36

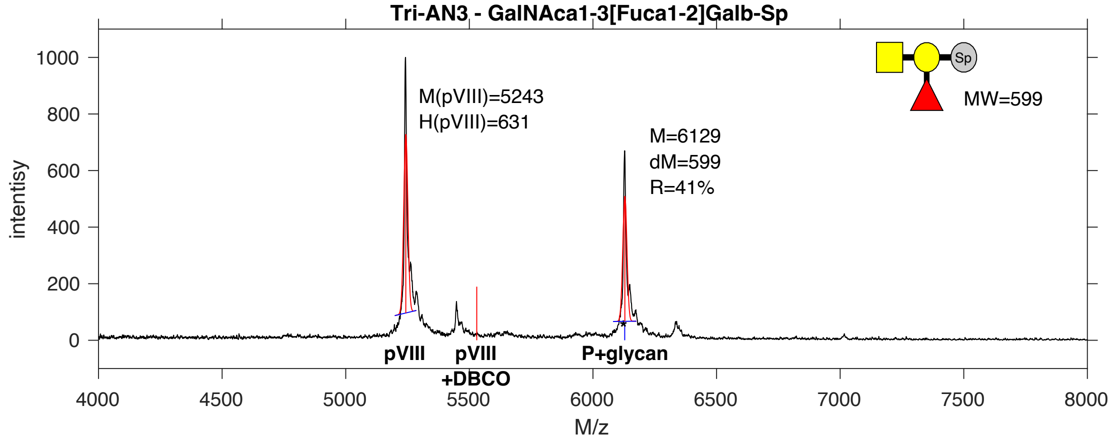


**100: MALDI-TOF spectra of glycosylated phage: Tri-AN3-[1080]**

**SDB Number:** SDB126

**Sequencing File:** http://ligacloud.ca/searchLibInfo?f=0&b=0&d=NA

**Barcode:**
CTGCTCTTTGCAATTCCGCTGAGTGTAGAAAAAAATGATCAGAAAACATA
CCATGCTGGTGGCGGA

**Axis Name:** Tri-AN3-[1080]

**IUPAC:** NANA

**IUPAC (CFG):** GalNAca1-3[Fuca1-2]Galb-Sp

**Common Name:** Tri-AN3

**Glytoucan ID:** G34704BH

**Compound Number:** Tri-AN3

**Maldi File:** MS-XIV-164_126.txt

**Density:** 40
