## Supplementary material for "Measuring carbohydrate recognition profile of lectins on live cells using liquid glycan array (LiGA)": Zipped folder containing all the software, MALDI TOF spectra and analyzed files: OligomerResolver.m

```
function [out] = OligomerResolver(in, TRIM)
    % The function resolves the oligomer and does the trimming:
    % original: Neu5Aca2-6(Galb1-4GlcNAcb1-3)2b--
    % resolved: Neu5Aca2-6Galb1-4GlcNAcb1-3Galb1-4GlcNAcb1-3b--
    % trimmed : Neu5Aca2-6Galb1-4GlcNAcb1-3Galb1-4GlcNAcb--
    % trimmed portion 'b1-3' found at the end of the bracket
    % when the flag is set to 1, the function will serach for '[ab]\d-\d'
    % and trim the last instance of it in the monomer

in0= in; % remember unchanged string;

% detect either ')n' as '\)\d' or ']n' as '\]\d+'
BRn = '(\)\d+|\]\d+)';

i=0;
[START,END] = regexp(in,BRn,'start','end');

while ~isempty(START)
    
    i=i+1;

        temp = regexp(in(START(1):END(1)),'\d+','match');
        N(i) = str2num(temp{1});
        
        % backtrack the beginning of a bracket from each BRn point
        j = START(1);
        closed = 1;
        while closed

            j= j-1;
            test = [in(j), in(START(1))];
            
            if     test == '()'
                closed = closed - 1;
            elseif test == '[]'
                closed = closed - 1;
            elseif test == '))'
                closed = closed+1;
            elseif test == ']]'
                closed = closed+1;
            end
        end

        monomer{i} = in(j+1 : START(1)-1);
        
        [S,E] =regexp(monomer{i}, '[ab]\d-\d', 'start', 'end');
        if ~isempty(S)
            trim_monomer = [monomer{i}(1 : S(end)-1) ...
                            monomer{i}(E(end)+1 : end) ];
        else
            trim_monomer = monomer{i};
        end
        
        oligomer = '';
        for k=1:N(i)
            if k == N(i) && TRIM
                oligomer = [oligomer trim_monomer];            
            else
                oligomer = [oligomer monomer{i}];
            end
        end
        
        in = [in(1: j-1) oligomer in(END+1 : end) ];
        
        [START,END] = regexp(in,BRn,'start','end');
 
end

out = in;
```
