## Supplementary material for "Measuring carbohydrate recognition profile of lectins on live cells using liquid glycan array (LiGA)": Zipped folder containing all the software, MALDI TOF spectra and analyzed files: plotONEmaldi.m

```
%%%%%%%%%%%%%%%%%%%%%%%%%%%%%%%%%%%%%%%%%%%%%%%%%%%%%%%%%%%%%%%%%%%%%%%%%%%
%The folder in which the .txt maldi spectra file is located
mydir  = pwd;
txtdir = strcat(mydir,'/TextFiles');
addpath (fullfile(txtdir));
SaveDir = fullfile(pwd,'Figures');
%%%%%%%%%%%%%%%%%%%%%%%%%%%%%%%%%%%%%%%%%%%%%%%%%%%%%%%%%%%%%%%%%%%%%%%%%%%


filename = 'MS-XIV-40i.txt.txt'; %Name of Maldi spectra file
names = '3S-Di-LN'; %Name of the Glycan
full_temp = 'Neu5Aca2-3(Galb1-4GlcNAcb1-3)2b-Sp'; %CFG name  and Spacer

isCapped = 0;  % set to 1 if azidoethanol capping was used. 
MW_DBCO = 287; % Molecular weight of DBCO linker
DBCO_label = '+DBCO' %label of DBCO peak
manual = 0; 

% clear some key variables
Nsialo = 0;
Ratio = [];

XLIM1 = 4000;
XLIM2 = 8000;

%range around the peak for gaussian fitting and baseline fit
% first range is for a larger p8 peak
RANGE0 = 70;
% second range is for the glycan peak
RANGE = 70;

fid = fopen(filename,'r');

if fid==-1
    disp(['dear user, your file ' filename ' is incorrect']);
    disp('*****************************************************');
    fprintf(FID, '%s', [filename ',' 'file not found'  char(10) ]);
end

    disp([names ' glycan is in ' filename ' ' fgetl(fid)...
           char(10) fgetl(fid)]);
    
    FORM  = '%f %f %*[^\n\r]';
    
     AllVar = textscan(fid,FORM);
    mass =  AllVar{1};
    intensity =   AllVar{2};
    
     disp(['Read ' num2str( size(mass, 1)) ' lines' ]);
    fclose(fid);
    
    IX = find(mass>XLIM1 & mass<XLIM2);
    

    plot_mass = mass(IX);
    plot_intensity = intensity(IX);
    YMIN = min(plot_intensity);
    YMAX = max(plot_intensity);
    YH = YMAX-YMIN;
    
    
    
    SCALE = 1000/YH;
    plot_intensity = plot_intensity*SCALE;
    YH = YH*SCALE;
    
    YMIN = min(plot_intensity);
    YMAX = max(plot_intensity);
    
    margin = 0.1;
    
    
    h = figure(1);
    set(h, 'Units', 'normalized', 'position', [0.2 0.6 0.45 0.25] )
    
    % find the full name based on abbreviated name

    
    %%%%%%%%% PLOT MALDI TRACE AND MAKE IT PRETTY %%%%%%%%%%%%%%%%%%%%%%%%%
    figure(1);
    hold off;
    plot(1,1);
    hold on;
    plot(plot_mass, plot_intensity, '-k');
    
    % add full name of glycan to the title
    title([names ' - ' full_temp]);
    
    set(gca,'yscale','lin','xscale','lin',...
       'TickDir','out');
   ylabel('intentisy');
   xlabel('M/z');
    xlim([4000 8000]);
    ylim([YMIN - margin*YH YMAX + margin*YH]);
    drawnow;
    hold on;
    
    
            IX2 = find ( plot_intensity == ...
                    max(plot_intensity(plot_mass>5000 & plot_mass<5500)) );
        pVIII = plot_mass(IX2(1));
        
        
        
               % define the RANGE and make sure it doesn't exceed the boundaries
        
        if IX2+RANGE0 > size(plot_intensity,1)
            H = size(plot_intensity,1);
        else
            H = IX2+RANGE0;
        end
        
        if IX2-RANGE0 < 1
            L = 1;
        else
            L = IX2-RANGE0;
        end
        
        initial_mass = pVIII;
        initial_int  = max( plot_intensity(L:H) );
        
        % set the initial width wide enough to prevent fitting to noise.
        init_width = RANGE0 / 2;
               
        % fitting starting point and lower bounds
        fo = fitoptions('Method','NonlinearLeastSquares',...
               'StartPoint',[initial_int initial_mass init_width 0 0],...
               'Lower', [0 initial_mass-2*RANGE0 init_width/3 -100 -1000]);
        ft = fittype('a1*exp(-((x-b1)/c1)^2)+e1*x+f1','options',fo);

        f0 = fit(plot_mass(L:H),plot_intensity(L:H),ft);
        
        
        
        
        X = plot_mass(L:H);
        MASS_P8 = f0.b1;
        MASS_P8_DBCO = MASS_P8 + MW_DBCO;
        
        HEIGHT_P8 = f0.a1;
        MAX_P8 = max(f0(X));
        
        plot(X, f0(X),  '-r');
        plot(X, f0.e1*X + f0.f1, '-b');
        
        % this is where thin vertical red line is drawn and the height of
        % this line is exactly the estimated height of the gaussian peak
        % after the slanted baseline correction
        
        line(MASS_P8*[1 1], [MAX_P8, MAX_P8-HEIGHT_P8], 'color','r');
        
        text(MASS_P8*1.01, MAX_P8,...
                 ['M(pVIII)=' num2str(round(MASS_P8)) char(10)...
                  'H(pVIII)=' num2str(round(HEIGHT_P8))],...
                  'VerticalAlignment', 'bottom',...
                  'HorizontalAlignment','left');
        
        text(pVIII, YMIN, 'pVIII', ...
              'FontWeight', 'bold',...
              'VerticalAlignment', 'top',...
              'HorizontalAlignment','center');

        
        line(MASS_P8_DBCO*[1 1], YMIN + [0 0.3*f0.a1], 'color', 'r');
        text(MASS_P8_DBCO, YMIN, ['pVIII' char(10) DBCO_label], ...
              'FontWeight', 'bold',...
              'VerticalAlignment', 'top',...
              'HorizontalAlignment','center');
        
        
%%%%%%%%% EXTRACT THE FULL NAME AND MAKE GLYCAN OBJECT %%%%%%%%%%%%%%%%
  
    sugar = GlycanLeaf.createObj(full_temp);
    
    hold on; plot(1,1);
    
    drawGlycan('input',sugar.String, 'XY', [7500 YMAX],...
               'spacing', 150, 'angle', pi);
    
 
    TEXT = ['MW=' num2str(round(sugar.MW)) char(10)];
    
    DEBUG = 0;
    [MW(char(10)),b,c] = sugar.MW;
    if DEBUG
    
        for j = 1:numel(sugar.glycans)
            TEXT = [TEXT sugar.glycans{j} ...
                    ': ' num2str(round(c(j) )) char(10)]; 
        end
    end
    
    text(7500, 0.90*YMAX, TEXT, 'HorizontalAlignment', 'left',...
                      'VerticalAlignment', 'top');
                  
%     set(gca,'xtick',[], 'ytick',[])
    
%%%%%%%%%%%%%%%%%%%%%%%%%%%%%%%%%%%%%%%%%%%%%%%%%%%%%%%%%%%%%%%%%%%%%%%%%%%        
        MASS_P8_DBCO_GLY = MASS_P8_DBCO + MW(char(10));
        
        line(MASS_P8_DBCO_GLY*[1 1], YMIN + [0 0.1*HEIGHT_P8], 'color', 'b');
        
        text(MASS_P8_DBCO_GLY, YMIN, '*',...
                'FontWeight', 'bold',...
                'VerticalAlignment', 'bottom',...
                'HorizontalAlignment','center');
        text(MASS_P8_DBCO_GLY, YMIN, 'P+glycan',...
                'FontWeight', 'bold',...
                'VerticalAlignment', 'top',...
                'HorizontalAlignment','center');
            
            drawnow;
        
    repeating = 1;
    j=0;
    
    while repeating>0
        if manual
            choice = questdlg('define a peak?',...
                                'define a peak?',...
                            'define a peak','no mo peaks','escape','define a peak');
            switch choice
                case 'escape', return;
                case 'no mo peaks', repeating = 0; continue;
                case 'define a peak', 

                    [x]=ginput(1); 
                    
                     j = j+1;
                    lineX(j) = x(1);
                   
            end
        else
            j = j+1;
            if j==1
                % find the main peak
                lineX(j) = MASS_P8_DBCO_GLY;
                
                IX =  (strcmp('Neu5Ac',sugar.glycans)     |...
                       strcmp('Neu5,9Ac2',sugar.glycans)  |...
                       strcmp('Neu5Gc',sugar.glycans)   |...
                       strcmp('KDN',sugar.glycans) );
                   
                repeating = sum(IX);
                
            elseif j==2 && repeating
                
                IX = ~(strcmp('Neu5Ac',sugar.glycans)    |...
                       strcmp('Neu5,9Ac2',sugar.glycans) |...
                       strcmp('Neu5Gc',sugar.glycans)  |...
                       strcmp('KDN',sugar.glycans) );
                Nsialo = sum( ~IX );
                
                [~,~,c]= sugar.MW;
                
                asialoMW = sum(c(IX)) - 18*(sum(IX)-1);

                MASS_P8_DBCO_GLY = MASS_P8_DBCO + asialoMW;
                
                lineX(j) = MASS_P8_DBCO_GLY;
                
                
                line(MASS_P8_DBCO_GLY*[1 1], YMIN + [0 0.1*HEIGHT_P8], 'color', 'b');
        
                text(MASS_P8_DBCO_GLY, YMIN, '*',...
                        'FontWeight', 'bold',...
                        'VerticalAlignment', 'bottom',...
                        'HorizontalAlignment','center');
                text(MASS_P8_DBCO_GLY, YMIN,...
                        ['-' num2str(Nsialo) 'xSialo'] ,...
                        'FontWeight', 'bold',...
                        'VerticalAlignment', 'top',...
                        'HorizontalAlignment','center');
                    
                repeating = 0;
            
            end
       
                % find the index of data nearest to the mouse click
                IX2 = max ( find( plot_mass < lineX(j)  ));

                        % define the RANGE and make sure it doesn't exceed the boundaries

                if IX2+RANGE > size(plot_intensity,1)
                    H = size(plot_intensity,1);
                else
                    H = IX2+RANGE;
                end

                if IX2-RANGE < 1
                    L = 1;
                else
                    L = IX2-RANGE;
                end

                initial_mass = lineX(j);
                initial_int  = max( plot_intensity(L:H) );
                
                % set the initial width wide enough to prevent fitting to noise.
                init_width = RANGE / 2;

                fo = fitoptions('Method','NonlinearLeastSquares',...
                       'StartPoint',[initial_int/5 initial_mass init_width 0 0],...
                       'Lower', [0 initial_mass-2*RANGE init_width/3 0 0]);
                ft = fittype('a1*exp(-((x-b1)/c1)^2)+e1*x+f1','options',fo);

                f = fit(plot_mass(L:H),...
                        plot_intensity(L:H),ft);
                    
                
                X = plot_mass(L:H);
                MASS(j) = f.b1;
                HEIGHT(j) = f.a1;
                MAX(j) = max(f(X));

                plot(X, f(X),  '-r');
                plot(X, f.e1*X + f.f1, '-b');
                line(MASS(j)*[1 1], [MAX(j), MAX(j)-HEIGHT(j)], 'color','r');
                
                Delta(j) = round(abs(MASS(j)-MASS_P8_DBCO));
                Ratio(j) = round(100*HEIGHT(j) / (HEIGHT(j) + HEIGHT_P8));
                
                if j==1
                    text(MASS(j)+100, MAX(j)*0.7 + 50,...
                        ['M=' num2str(round(MASS(j))) char(10)...
                         'dM=' num2str(Delta(j)) char(10)...
                         'R=' num2str(Ratio(j)) '%' char(10)],...
                          'VerticalAlignment', 'bottom',...
                          'HorizontalAlignment','left');
                elseif j==2
                    text(MASS(j), MAX(j) + (MAX(j)-HEIGHT(j))*0.1,...
                    ['M=' num2str(round(MASS(j))) char(10)...
                     'dM=' num2str(Delta(j)) char(10)...
                     'R=' num2str(Ratio(j)) '%' char(10)],...
                      'VerticalAlignment', 'bottom',...
                      'HorizontalAlignment','right');
                end
                                           
        end
    end
    
  
```
