## Supplementary material for "Measuring carbohydrate recognition profile of lectins on live cells using liquid glycan array (LiGA)": Zipped folder containing all the software, MALDI TOF spectra and analyzed files: ReadMe.md

# Matlab Maldi Scripts

### plotONEmaldi.m dependecies 

All the required dependent function are included in the current directory. Need to download all entire folder and sub-folders
within 'maldi' folder for the script to works. Curve Fitting Toolbox Add-in (https://www.mathworks.com/products/curvefitting.html) for Matlab is required. 

For plotting maldi, use plotONEmaldi.m and fill in the values for the variables. An example is given below:

```
filename = '1.txt'; %Name of Maldi spectra file
names = 'Te286'; %Name of the Glycan
full_temp = 'Galb1-3GlcNacb1-3Galb1-4[Fuca1-3]GlcNAcb-Sp'; %CFG name and Spacer
```
